# Dynamic assembly of the calcium hemostasis modulator 1 channel gates ATP permeation

**DOI:** 10.1101/2021.01.14.426634

**Authors:** Yue Ren, Yang Li, Yaojie Wang, Tianlei Wen, Xuhang Lu, Shenghai Chang, Xing Zhang, Yuequan Shen, Xue Yang

**Affiliations:** State Key Laboratory of Medicinal Chemical Biology, College of Life Sciences and College of Pharmacy, Nankai University, Tianjin 300350, China; Department of Biophysics and Department of Pathology of Sir Run Run Shaw Hospital, Zhejiang University School of Medicine, Hangzhou, China; Center of Cryo Electron Microscopy, Zhejiang University School of Medicine, Hangzhou, China; Synergetic Innovation Center of Chemical Science and Engineering, Tianjin 300071, China

**Author notes:** Correspondence and requests for materials should be addressed to Yuequan Shen,; Xue Yang,. equal contribution (Yue Ren, Yang Li).

**Keywords:** Calcium hemostasis modulator, CALHM1, ATP permeation, gating mechanism, assembly, Cryo-EM.

## Abstract

Calcium hemostasis modulator 1 (CALHM1) is a voltage- and Ca^2+^-gated ATP channel that plays an important role in neuronal signaling. The currently reported CALHM structures are all in an ATP-conducting state, and the gating mechanism of ATP permeation remains elusive. Here, we report three cryo-EM reconstructions of *Danio rerio* heptameric CALHM1s with ordered or flexible long C-terminal helices as well as *Danio rerio* octameric CALHM1 with flexible long C-terminal helices at resolutions of 3.2 Å, 2.9 Å, and 3.5 Å. Structural analysis revealed that the heptameric CALHM1s are in an ATP nonconducting state in which the pore diameter in the middle is approximately 6.6 Å. Compared with those inside the octameric CALHM1s, the N- helices inside heptameric CALHM1s are in the “down” position to avoid steric clash with neighboring TM1 helices. Molecular dynamic simulation shows that the pore size is significantly increased for ATP molecule permeation during the movement of the N-helix from the “down” position to the “up” position. Therefore, we proposed a mechanism in which the “piston-like” motion of the N-helix drives the dynamic assembly of the CALHM1 channel for ATP molecule permeation. Our results provide insights into the ATP permeation mechanism of the CALHM1 channel.

## Introduction

Adenosine triphosphate (ATP) release channels play an important role in various cellular signaling events (Abbracchio et al, 2009). Consequently, their malfunctions are associated with many pathophysiological processes, including neurological disorders, inflammation, and cancer progression (Taruno, 2018). Extensive research has identified five family proteins as ATP release channels so far, although some of them await further verification. These five family proteins are connexin hemichannels, pannexins (PANXs), calcium hemostasis modulators (CALHMs), volume-regulated anion channels (VRACs), and maxi-anion channels (MACs) (Taruno, 2018). Convergent evolution analysis showed that they all have four transmembrane helices (TMs) in the monomer that forms an oligomer with a pore in the middle (Siebert et al, 2013).

CALHM1 is a large-pore nonselective ion channel gated by voltage and extracellular Ca^2+^ concentration (Ma et al, 2016). In addition, CALHM1 is a voltage-gated ATP release channel that mediates purinergic neurotransmission of sweet, bitter, and umami tastes from type II taste bud cells to the taste nerve (Romanov et al, 2018; Taruno et al, 2013). Compared with the slow activation kinetics of CALHM1 homo- oligomerized channels, CALHM1/CALHM3 can form hetero-oligomerized ion channels with rapid voltage-gated activation kinetics that are closer to physiological states *in vivo* (Ma et al, 2018). Structural studies of CALHM family proteins have recently made great progress (Foskett, 2020). CALHM1 structures of several species have been reported to be octamers that do not form gap junctions due to N-glycosylation of the extracellular loop (Demura et al, 2020; Ren et al, 2020; Syrjanen et al, 2020).

CALHM2 is different from octameric CALHM1, and it forms an undecamer, with two conformations of hemichannels and gap junctions (Choi et al, 2019; Demura et al, 2020; Drozdzyk et al, 2020; Syrjanen et al, 2020). It is puzzling that different CALHM isoforms form diverse oligomeric assemblies. Recent research on the CALHM1- CALHM2 chimera structure indicated that interactions between the C-terminal helices (CTHs), and the TM4-CTH linker may determine the oligomeric state of the CALHM channels (Syrjanen et al, 2020). Ren et al. proposed that the extracellular loop 1 region within the dimer interface may contribute to oligomeric assembly (Ren et al, 2020).

In this research, we reported three cryo-EM reconstructions of the *Danio rerio* CALHM1 heptamer with ordered long C-terminal helices (drCALHM1-hepta-LCH) and flexible long C-terminal helices (drCALHM1-hepta-noLCH) as well as the *D. rerio* CALHM1 octamer with flexible long C-terminal helices (drCALHM1-octa-noLCH). The comparison between different CALHM1 oligomers from the same species suggests that the “piston-like” motion of the N-helix inside the pore may drive the dynamic assembly of the CALHM1 channel and thus regulate ATP permeation.

## Results

### Structure determination of drCALHM1 channels

The structures of CALHM1 channels were determined by cryo-electron microscopy (cryo-EM). drCALHM1 channels were purified in the presence and absence of Ca^2+^ and concentrated for cryo-EM sample preparation. All data sets were collected using a Titan Krios electron microscope operated at 300 kV accelerating voltage with a K2 Summit direct electron-counting detector. Details on the data collection and data processes are shown in Supplemental Figures 1-4. The drCALHM1- hepta-LCH structure was reconstructed from the data set in the presence of Ca^2+^. The drCALHM1-hepta-noLCH and drCALHM1-octa-noLCH structures were reconstructed from data sets in the absence of Ca^2+^. The final 3D reconstructions of drCALHM1-hepta-LCH, drCALHM1-hepta-noLCH, and drCALHM1-octa-noLCH were determined at overall resolutions of 3.2 Å, 2.9 Å, and 3.5 Å, respectively (Supplemental Figures 1 and 2). The electron densities are sufficient to resolve most amino acid side chains (Supplemental Figures 3 and 4).

### Overall structure of the drCALHM1-hepta-LCH channel

The overall structure of the heptameric drCALHM1 is shaped like a barrel and can be divided into three parts: extracellular loop, transmembrane domain, and cytoplasmic domain (Figure 1). From the top view, its extracellular region forms a ring with a diameter of approximately 98 Å, and seven small helices in the middle of the ring can be clearly observed (Figures 1A and 1D). The side view shows that the height of the drCALHM1 heptamer is approximately 92 Å. There are multiple lipid molecules that exist inside a protomer and within the interface of two protomers (Figures 1B and 1E). From the cytosolic view, it can be observed that the cytoplasmic domain mainly consists of one long helix. Seven C-terminal long helices (CTH) intercross each other (Figures 1C and 1F), resulting in a ring diameter of approximately 100 Å. The overall drCALHM1 heptamer is quite similar to that of previously reported octamer (Ren et al, 2020) except for the different oligomerization states.

**Fig. 1.**
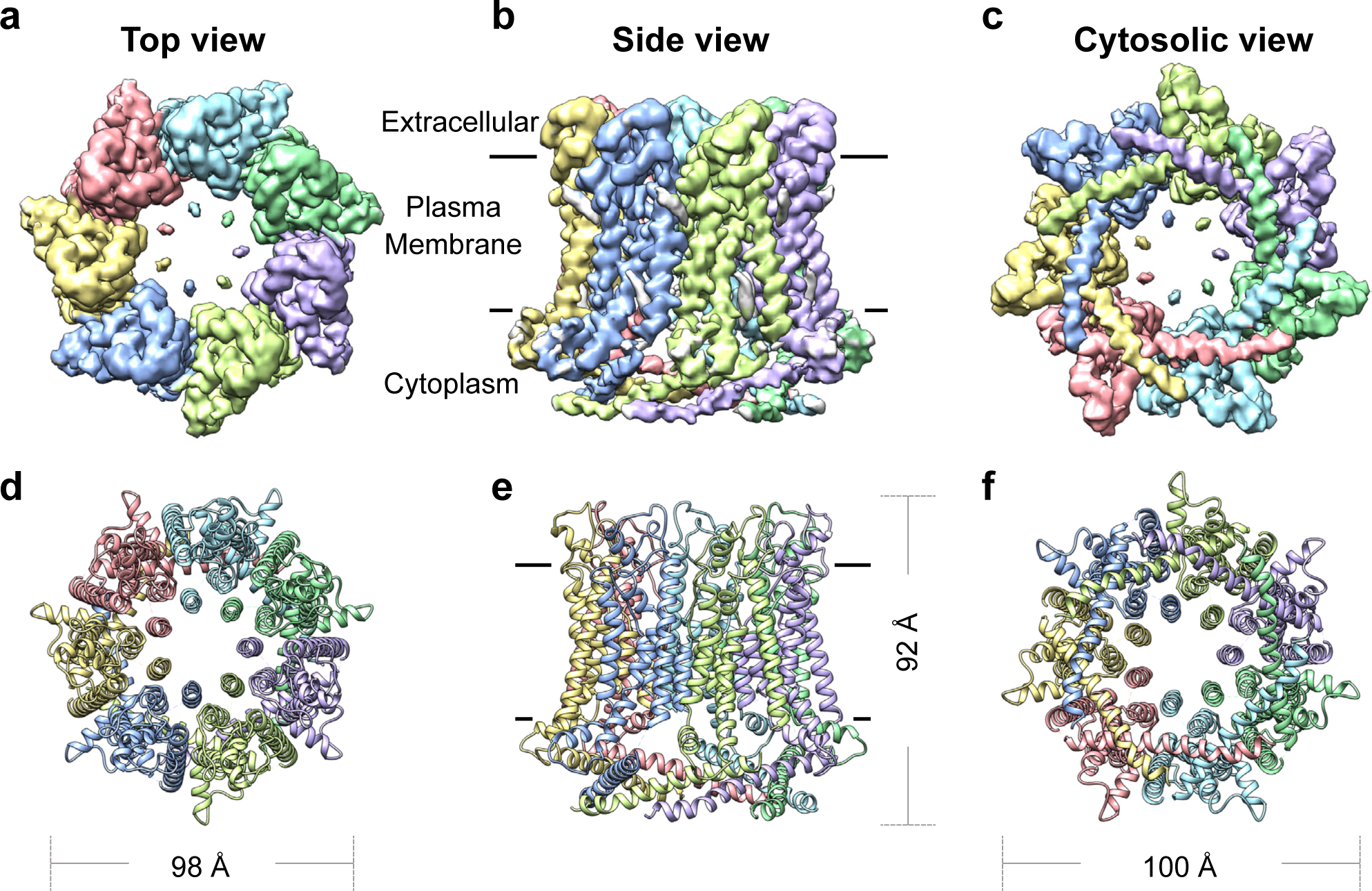
The overall architecture of drCALHM1-hepta-LCH. The surface (a) and cartoon (d) representations of the cryo-EM density map of the heptameric drCALHM1, viewed from the extracellular side of the membrane. The seven subunits are represented by different colors. The surface (b) and cartoon (e) representations of the cryo-EM density map of drCALHM1-hep- ta-LCH, viewed parallel to the membrane. The lipid-like densities are in gray. The surface (c) and cartoon (f) representations of the cryo-EM density map of drCALHM1-hepta-LCH viewed from the cytosolic side of the membrane.

### ATP nonconducting state of the drCALHM1-hepta-LCH

As shown in Figure 1, the heptamer has smaller pores than the octamer, and we further calculated the exact pore size with the N-helix having all alanines in the current model (Figure 2A). The narrowest diameter of the drCALHM1-hepta-LCH channel is approximately 14 Å (Figure 2B). If the putative side chain was built for the N-helix, the pore diameter was reduced to 6.6 Å, which blocked ATP permeation through the pore but allowed ion flux. To further validate this result, we took advantage of the molecular dynamics (MD) simulation methodology. Coarse-grained simulations of drCALHM1-hepta-LCH in the presence of 1-palmitoyl-2-oleoyl-sn-glycero-3-phosphocholine (POPC) lipids were carried out. We added hydrated Na^+^, hydrated K^+^, and Cl^−^ ions on either side of the POPC bilayer to Martini v.2.0 forcefield (Monticelli et al, 2008). Ca^2+^ ions were added on the extracellular side. During replicates of 10-μs coarse-grained MD simulations, we observed that multiple Na^+^ cations, K^+^ cations, and Cl^−^ anions crossed the membrane through the drCALHM1-hepta-LCH channel. In contrast, Ca^2+^ ions were still on the extracellular side of the POPC bilayer after simulations (Figure 2C). Ca^2+^ ions occasionally moved closer to interact with Glu79 and Glu80 in the extracellular loop and were unable to pass through the heptameric channel. These simulation results indicated that the drCALHM1-hepta-LCH channel favors the permeation of small ions (Na^+^, K^+^, and Cl^−^) but repulses Ca^2+^ ions. Moreover, its pore size is too narrow for bulky ATP molecules.

**Fig. 2.**
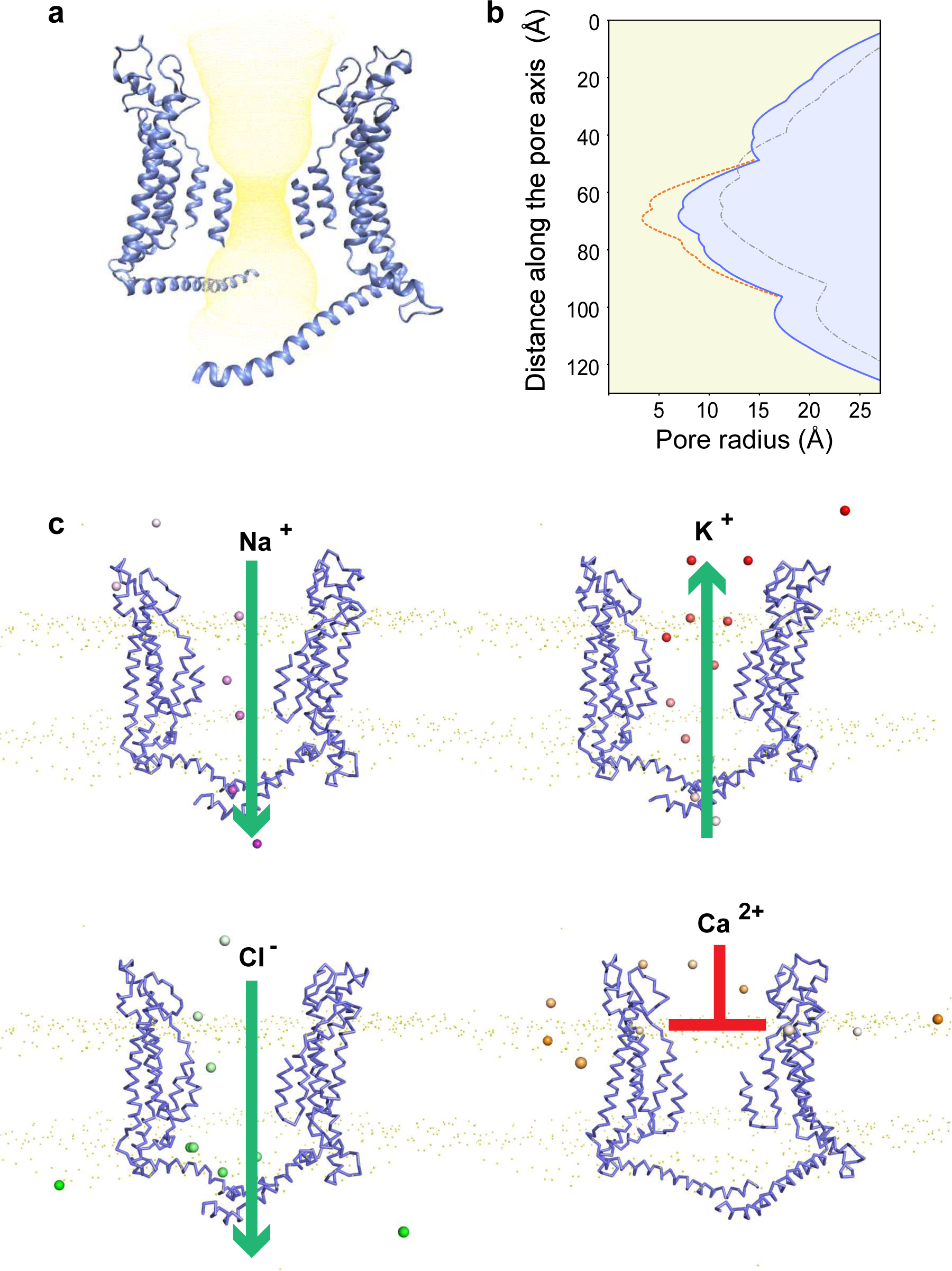
ATP nonconducting state of the central pore. (a) The permeation path of the heptamer without the N-helix side chain calculated by HOLE is marked by yellow dots. (b) The pore radii along the central axis of drCALHM1 in different oligomerization states are marked as follows: the heptamer with side chains is orange, the heptamer without side chains is blue, and the octamer without side chains is gray. (c) Positions of the ions in the heptamer channel in the coarse-grained simulations. The monomers of heptamer are shown in ribbon representation, Na^+^, K^+^, Cl^-^, and Ca^2+^ ions are shown in sphere model and colored purple, red, green, and orange, respectively. The color changes over the simulation time from transparent to opaque.

These results suggest that the drCALHM1-hepta-LCH channel is a closed conformation for ATP permeation. It has been noted that the ATP nonconducting conformation reported herein is the first to be resolved.

We also calculated the electrostatic surface potential distribution of the drCALHM1-hepta-LCH channel and found that the positive potential at the center of the pores of the heptamer is significantly stronger than that of the octamer (Supplemental Figure 5), which may be due to the change in the oligomerization state to make the heptameric channel more compact. The strong positive potential distribution inside pores may indicate the voltage-dependent characteristics of ion flux through the CALHM1 channel, as shown by electrophysiological studies (Tanis et al, 2017).

### Structural comparison of heptamer and octamer

To address the underlying mechanism of two different oligomers formed by the same CALHM1 protomer, we compared the heptamer and the octamer by the superimposition of one protomer together (Figure 3A). The entire heptamer forms a smaller circle than the octamer does. The protomer is composed of four transmembrane helices (TM1, TM2, TM3, and TM4), one extracellular helix, and one CTH. The protomers from the heptamer and octamer are extremely similar except for the position of the N-helix. Compared with the N-helix of the octamer, the N-helix of the heptamer moves towards the cytosolic direction by approximately 6 Å (Figure 3B) (defined as the “down” position in the heptamer and the “up” position in the octamer hereafter), resulting in “piston-like” motion inside the pore. In the heptamer, the N-helix aligns parallelly with TM1 and forms an interaction with TM1 (Figure 3C). The side chain of Leu28 has hydrophobic interactions with Ala10. Moreover, the N-helix from one protomer has multiple interactions with TM1 from the neighboring protomer. One hydrogen bond was formed between the side chain of Gln32 and the main chain carboxyl oxygen atom of Ala9. Additionally, the side chain of Ile25 has hydrophobic interactions with Ala16. Due to the density quality, only an alanine model was built for the N-helix. We anticipated more interactions between the N-helix and TM1 if the side chain of the N-helix could be built.

**Fig. 3.**
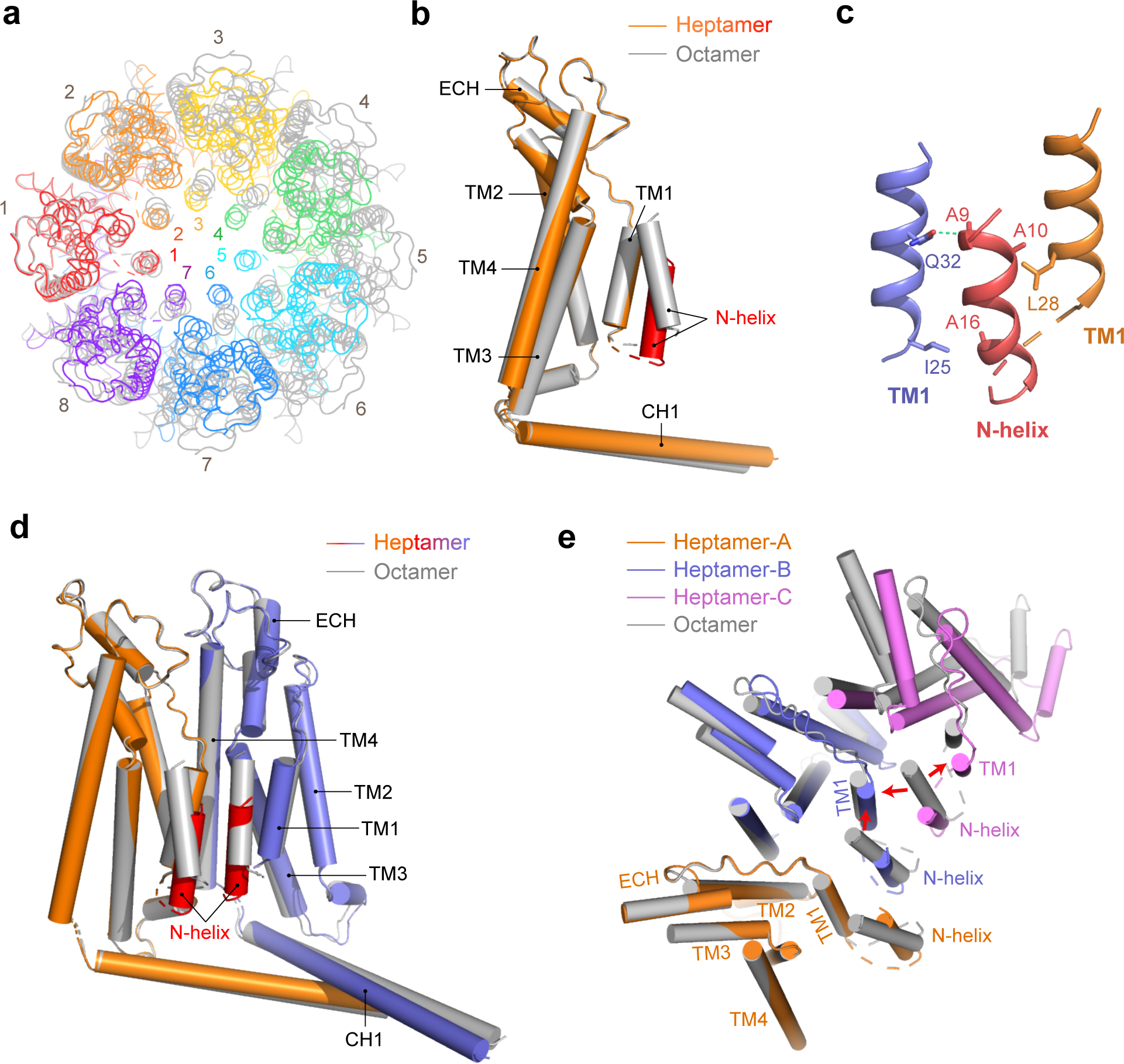
Structural comparison of the heptamer and the octamer. (a) Cartoon representation of the drCALHM1 heptamer (rainbow) and octamer (gray) viewed from the top of the membrane after superimposing a protomer from each oligomer. (b) Superimposition of the TMDs of the drCALHM1 heptamer (orange) and octamer (gray) viewed from the side of the membrane. The N-helix of the heptamer is marked in red for better contrast. (c) The interaction between the N-helix and the TM1. Orange TM1 and red N-helices are in the same protomer, and blue TM1 is on another adjacent protomer. The dashed line represents the hydrogen bond. (d) Comparison with dimers of different oligomeric states of drCALHM1 by superimposition of a protomer. The protomer of the heptamer is superimposed in orange, the other is blue, and the N-helix is marked red for contrast. The octamer is gray. (e) Comparison with trimers of the drCALHM1 heptamer and octamer (gray) by superimposition of a protomer, viewed from the top of the membrane. The three protomers of the heptamer are represented by orange, blue and magenta. The red arrows indicate the steric clashes between the N-helix of the octamer and the adjacent TM1s of the hep- tamer.

We then superimposed the dimers from the heptamer and the octamer. These two dimers are similar except for two N-helices (Figure 3D). Similar to the protomer comparison result, two N-helices in the dimer adapted from the heptamer obviously move down. After superimposing one protomer together, the neighboring protomer from the heptamer occupies a closer position to the pore than that from the octamer (Figure 3E), resulting in the N-helix from the octamer having a steric clash with TM1 from the heptamer (as shown by the red arrow). Therefore, the N-helix is mandatory in the “down” position in the heptamer, suggesting that the “up-down” motion of the N-helix may regulate the assembly of the octamer and the heptamer to gate ATP permeation.

### Up-down motion of the N-helix increases the pore diameter

To exclude the possibility of artifacts in which the drCALHM1 channel forms a heptamer, we transfected DNA encoding GFP-tagged full-length drCALHM1 into HEK293T cells and carried out electrophysiological studies. Large voltage-dependent outward currents were observed using the whole-cell configuration (Supplemental Figure 6), indicating that drCALHM1 expression in HEK293T cells forms a channel, as previously reported for the *Homo sapiens* CALHM1 channel (Ma et al, 2012). Next, we used two different crosslinking reagents, disuccinimidyl suberate (DSS) and formaldehyde, to assess the oligomeric state of drCALHM1 and *H. sapiens* CALHM1 expressed in the plasma membrane of HEK293T cells by western blots. Heptamers and octamers of CALHM1 s were observed by increasing the concentration of both crosslinking reagents (Supplemental Figure 7). These results validate our hypothesis that CALHM1 can form both the heptamer and the octamer in cells. To further test the possibility that the “up-down” motion of the N-helix can open the gate of ATP permeation, we carried out MD simulations. With the help of unbiased supervised MD and biased targeted MD simulations for drCALHM1-hepta-LCH, we accelerated the “up-down” motion of the N-helix in one protomer of the heptamer from the “down” position to the “up” position on a time scale accessible to MD simulations. During the supervised MD simulations, we monitored the root mean square deviation (RMSD) of the N-helix of protomer 1 (P1) in the current structure compared with that of P1 in the octamer to represent the movement of the N-helix. To monitor the pore size, we calculated the minimum distance of any atom between two N-helices from P1 and P4 protomers (denoted as P1−P4 thereafter), from P2 and P5 protomers (P2−P5), from P3 and P6 protomers (P3−P6), and from P4 and P7 protomers (P4−P7) (Supplemental Figure 8A).

There was obvious deformation of the 7-fold symmetry structure in the N-helix region caused by the “up-down” motion of the N-helix in P1 during the supervised MD simulations. Then, we calculated the PMF for the RMSD of the N-helix vs. the P4−P7 distance (Figure 4A). The PMF map shows the transition process of the heptamer and clearly depicts three different conformational states (state 1, state 2, and state 3) of the heptamer starting from the initial state (state 0) (Figure 4B). In initial state 0, the RMSD of the N-helix is approximately 6.66 Å, and the P4-P7 distance is approximately 9.57 Å. In states 1, 2, and 3 transformed from state 0, the RMSD of the N-helix is 5.63, 4.80, and 4.70 Å, and the P4−P7 distances are approximately 9.79, 11.89, and 13.26 Å, respectively. The conformational state of the N-helix moves closer to that of P1 in the octamer. Meanwhile, the pore size gradually increased from 9.79 Å to 13.26 Å. Furthermore, the targeted MD also observed that the pose size increased due to the “up-down” motion of the N-helix (Supplemental Figure 8B). Conventional MD simulations were performed as controls, and the position of the N-helix and the pore size of the heptamer remained similar to those in the initial state throughout the simulation time (Supplemental Figure 8C).

**Fig. 4.**
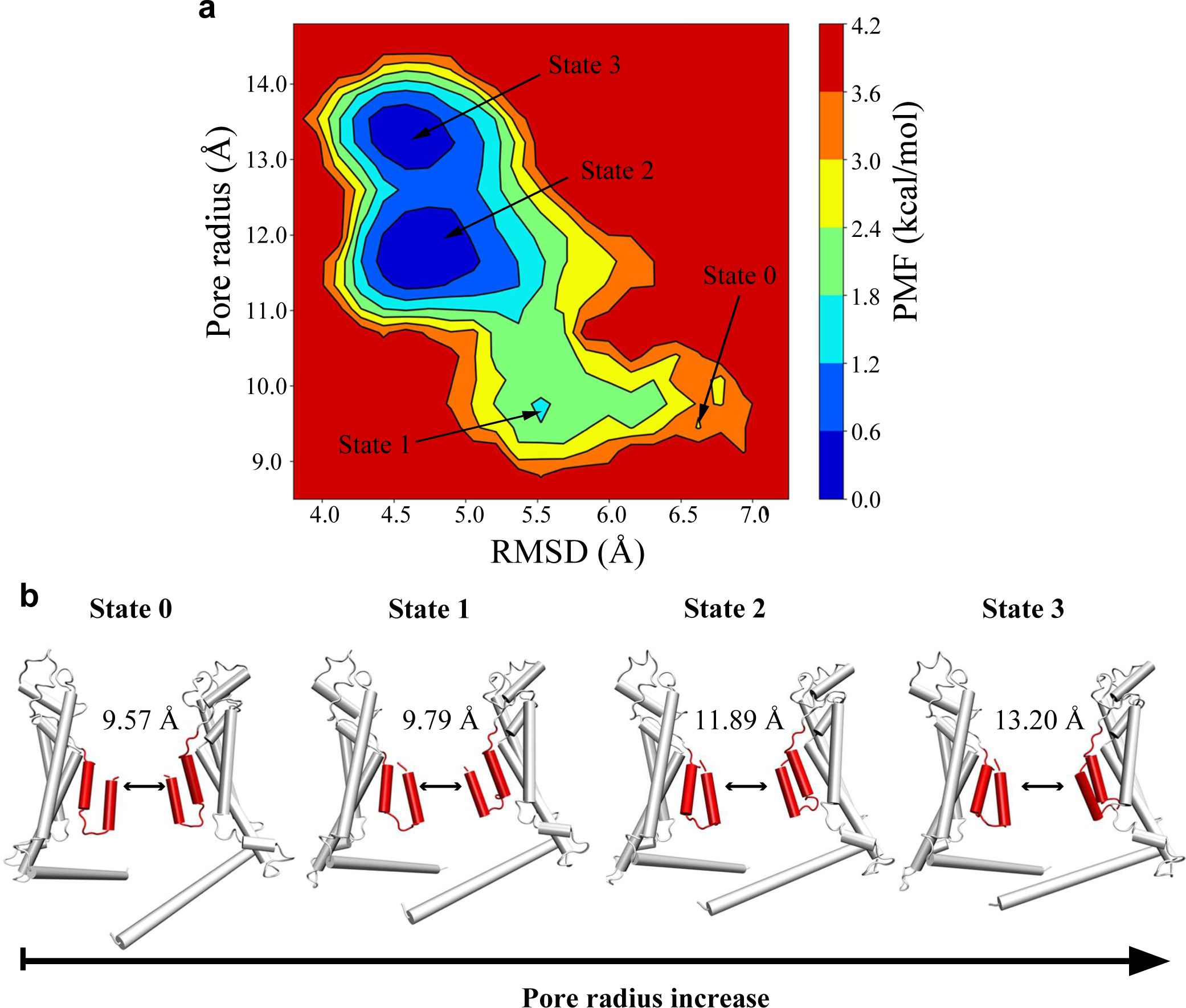
Up-down motion of the N-helix increases the pore diameter. (a) PMF calculated for the RMSD of the P_1_ N-helix in the heptamer versus the P_4_ –P_7_ distance. (b) Representative structures of P_4_ and P_7_ in state 0, state 1, state 2 and state 3 of the heptameric drCALHM1 channel. Two protomers of the heptamer are shown in cartoon.

### Structures of drCALHM1-hepta-noLCH and drCALHM1-octa-noLCH

When processing the data sets in the absence of Ca^2+^, we identified different conformation states for both heptameric (Figures 5A-5F) and octameric (Figures 5G-5 L) drCALHM1 channels in which long C-terminal helices are disordered and cannot be modeled. The overall structures of drCALHM1-hepta-noLCH versus drCALHM1-hepta-LCH and drCALHM1-octa-noLCH versus drCALHM1-octa-LCH are highly similar except for long C-terminal helices (Supplemental Figures 9A and 9B). Moreover, the pore diameters remained similar in these structures with or without long C-terminal helices (Supplemental Figures 9C and 9D), indicating that the long C-terminal helix did not regulate the pore size. We also compared the buried surface area of the protomer in different conformational states. With the long C-terminal helix, the buried surface areas are 948.6 Å^2^ and 959.6 Å^2^ for the monomer in the drCALHM1-hepta-LCH structure and drCALHM1-octa-LCH structure, respectively. However, without the long C-terminal helix, the buried surface areas are 627.6 Å^2^ and 603.3 Å^2^ for the protomer in the drCALHM1-hepta-noLCH structure and drCALHM1-octa-noLCH structure, respectively. These results suggest that the long C-terminal helix may serve as a scaffold to stabilize oligomerized CALHM1 channels.

**Fig. 5.**
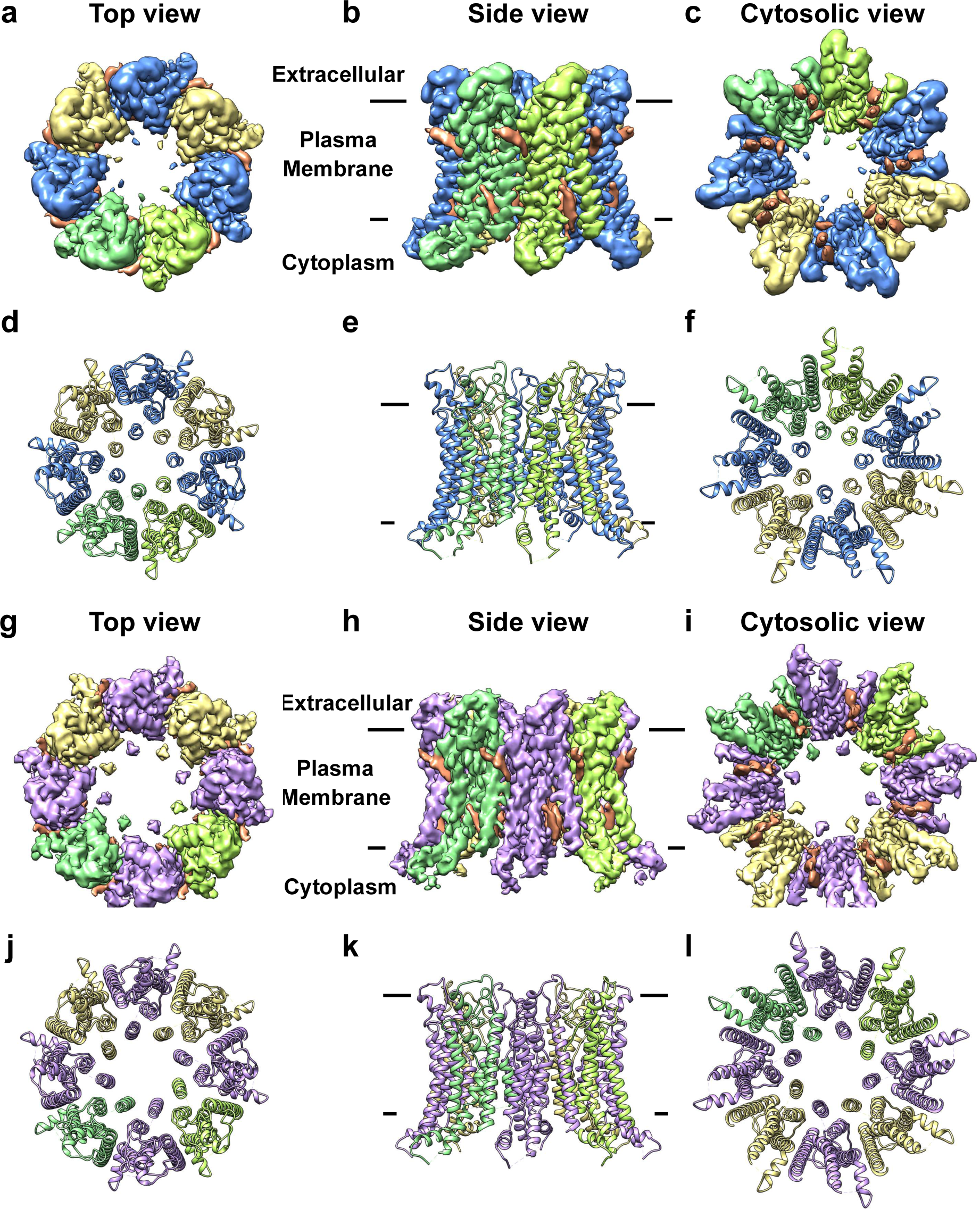
The overall structure of drCALHM1-hepta-noLCH and drCALHM1-octa-noLCH. Density map (a-c) and cartoon (d-f) of drCALHM1-hepta-noLCH in different directions. Blue and yellow are interlaced between adjacent subunits. The surface (g-i) and cartoon (j-l) of different sides of drCALHM1-octa-noLCH. The adjacent subunits are interlaced with purple and yellow. To clearly show the changes in subunit assembly during the transition of the oligo- meric state, the key subunits are marked with different degrees of green. The lipid-like densi- ties are in orange.

**Fig. 6.**
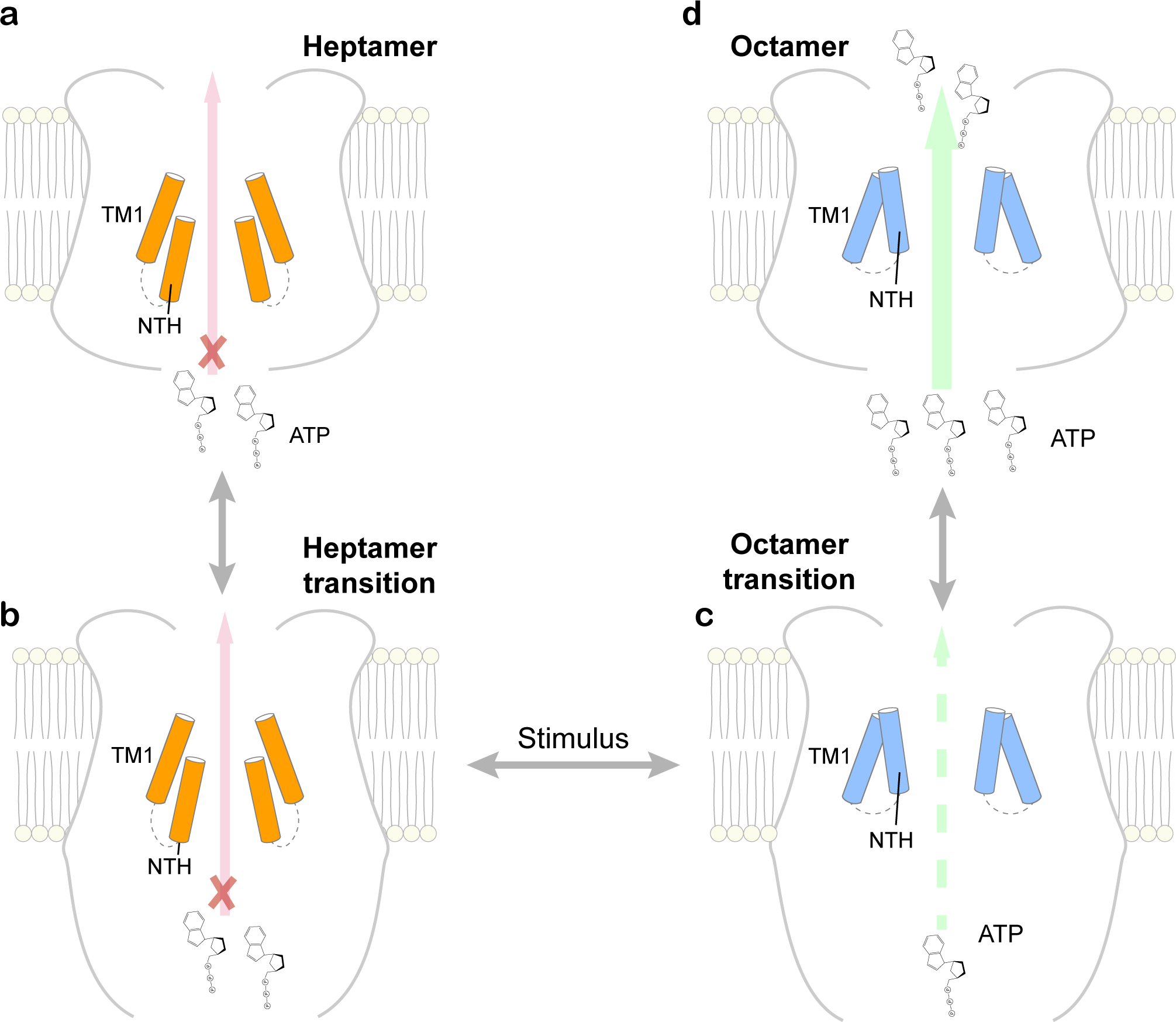
Proposed mechanism of ATP permeation in the CALHM1 channel. In the resting state, the CALHM1 channel may form a heptamer that prevents the permeation of ATP but allows ion flux. There is an equilibrium between the ordered long C-terminal helix and the disordered long C-terminal helix. Upon sensing stimulus, N-helix movement triggers the reassembly of the protomer to the octameric channel, eliciting an increase in the pore radius to permeate the ATP molecule. Adenosine triphosphate (ATP) is shown in the form of a chemical structural formula, and the green arrow represents the permeation of ATP.

## Discussion

In this study, we reported a heptameric drCALHM1 channel with an ATP nonconducting conformation. Compared with the octameric drCALHM1 channel, the heptameric drCALHM1 channel possesses two major changes. One is a much smaller channel ring, leading to the smaller pore and thus possibly blocking ATP permeation. Since the narrowest pore diameter is approximately 6.6 Å, it is likely that the ions can pass through the pores of the heptamer. Indeed, MD results showed that Na^+^, K^+^, and Cl^-^ can pass through the pore. However, Ca^2+^ was precluded from the pore, which may be caused by the positively charged environment inside the pore. Nevertheless, such narrow pores will unlikely allow ATP molecule permeation. The other difference is that the N-helix moves towards the cytosolic direction by approximately 6 Å to avoid the steric clash with neighboring TM1 helices, suggesting that the change in the oligomer state may be caused by the movement of the N-helix; in other words, the movement of the N-helix may drive dynamic assembly of the CALHM1 channel. Due to technical limitations, MD simulation from the heptamer to the octamer is unfeasible. Instead, we used supervised MD simulations and targeted MD simulations to monitor the potential impact of the movement of the N-helix in the heptameric drCALHM1 channel. As we expected, the movement of the N-helix to the “up” position led to the deformation of the 7-fold symmetric structure and significantly increased pore size.

We also determined two conformational states of heptameric and octameric drCALHM1 channels with flexible long C-terminal helices. Although the current structures of drCALHM1-hepta-noLCH and drCALHM1-octa-noLCH were reconstructed from the Ca^2+^-free data set, these two conformational states can be observed from the data set in the presence of Ca^2+^. Moreover, it has been noted that the final models of drCALHM1-hepta-noLCH and drCALHM1-octa-noLCH do not include long C-terminal helices due to the intrinsic flexibility in these specific conformational states instead of protein degradation. The purified protein sample did not show obvious degraded bands from the SDS-PAGE gel. Therefore, it is reasonable to hypothesize that for the same type of oligomer, the drCALHM1 channel may exhibit equilibrium between two conformational states (ordered versus flexible long C-terminal helices). Furthermore, from the buried surface area calculation of the protomer, it will be much easier to convert from the heptamer to the octamer for drCALHM1 with flexible long C-terminal helices than that with ordered long C-terminal helices.

Together with previously published results, we proposed a molecular mechanism of ATP permeation of the CALHM1 channel. In the resting state, the CALHM1 channel may form a heptamer that blocks ATP permeation but allows ion flux. The heptameric CALHM1 channel may exhibit equilibrium between ordered long C-terminal helices and flexible long C-terminal helices. Upon sensing, the N-helix inside the pore moves up towards the extracellular direction, driving the heptamer with flexible long C-terminal helices to form the octamer with flexible long C-terminal helices. At this stage, the pore is wide enough to permeate the ATP molecule. ATP molecules may enter the pore and induce the formation of ordered long C-terminal helices, which stabilize the octameric drCALHM1 channel in an open state and allows ATP molecule permeation. Compared with the published ATP permeation mechanism of another ATP release channel, Pannexin 1 (Ruan et al, 2020), our proposed mechanism of ATP permeation has a common point: both use the C-terminal tail to regulate ATP permeation. The major difference is that the C-terminal tail has to be cleaved during ATP permeation in Pannexin 1, while the C-terminal part in CALHM1 may switch to the disordered region for further assembly into higher-order oligomers to permeate ATP molecules.

To date, several CALHM subtype structures have been reported to form various oligomers (Choi et al, 2019; Demura et al, 2020; Drozdzyk et al, 2020; Ren et al, 2020; Syrjanen et al, 2020). Consequently, the “protein gate” or “lipid gate” model has been proposed to explain ATP permeation (Choi et al, 2019; Drozdzyk et al, 2020; Syrjanen et al, 2020). Our results are unfavorable for these two models. We did not observe any TM1 movement when compared with the closed and open states of drCALHM1 channels. Additionally, MD simulations did not support that lipids enter the pore. During the 10-μs coarse-grained MD simulations, the lipids assembled into clearly defined upper and lower leaflets around both proteins could not be accommodated within both channel pores of proteins (Supplemental Figure 10). Our new “piston-like” model is consistent with published functional data and provides a reasonable gating mechanism of ATP permeation for these unique CALHM channels.

## Acknowledgments

We are grateful to thank Ms. K. Tang in the Center of Cryo-Electron Microscopy (CCEM), Zhejiang University for their technical assistance on Cryo-EM data acquisition and Dr. A. Li and Dr. J. Lu for the support of electron microscopy at Nankai University. This work was supported by National Key Research and Development Program of China (grant 2017YFA0504801 to Y.S.; 2017YFA0504803 and 2018YFA0507700 to X.Z.), National Natural Science Foundation of China (grants 91954119 and 31870736 to X.Y.; grants 32071231 and 31870834 to Y.S.) and the Fundamental Research Funds for the Central Universities (2018XZZX001-13 to X.Z; 035-63201110 To X.Y. and 035-63201109 to Y.S.).

## Author Contributions

Y.R. and T.W. did protein expression, purification, sample preparation and cryo-EM data collection; Y.L. did MD simulations. X.L., S.C., X.Z. and X.Y. did cryo-EM data collection; X.Y. conducted cryo-EM reconstruction; Y.R., Y.L., X.Y. and Y.S. analyzed the data, designed the study and wrote the paper. All authors discussed the results and commented on the manuscript.

## Materials and Methods

### Expression of drCALHM1 channel

The full-length *Danio rerio* CALHM1 (UniProtKB number: E7F2J4, synthesized by Genewiz Inc.) was ligated into the EcoRI-NotI restriction sites of pEG-BacMam vector, with TEV protease cleavage site and enhanced green fluorescent protein (eGFP) at its N-terminal. The BacMam viruses were generated and amplified follow the standard Bac-to-Mam Baculovirus expression system (Invitrogen) in *Spodoptera frugiperda* (Sf9) cells. The bacmid produced by DH10Bac cells was transfected into Sf9 cells using X-tremeGENE HP DNA Transfection Reagent (Roche), and then cultured at 27 °C. HEK293S GnTi^-^ cells were cultured in Freestyle 293 expression medium (Thermo Fisher Scientific) at 37 °C with 5% CO2. When cell density reached 2 x10^6^ cells per mL, transfected with the second-generation virus (P2). After transfection, the cells were incubated at 37 °C for 24h, and then sodium butyrate was added to a final concentration of 10 mM to facilitate protein expression. The cells were cultured at 30 °C for another 60 h before harvest.

### Purification of drCALHM1 channel

The cells were collected by centrifugation at 800 ×g, and resuspended in lysis buffer consisting of 20 mM Tris-HCl pH 7.5, 200 mM NaCl, 1 mM PMSF, and then lysed through sonication for 15 min. The membrane pellets were collected by ultracentrifugation at 180,000 ×g for 1 h, and then homogenized in the buffer consisting of 20 mM Tris-HCl pH 7.5, 200 mM NaCl, 1× protease inhibitor cocktail (Roche), and solubilized in 1% (w/v) n-dodecyl-β-D-maltoside (DDM, Anatrace), 0.2% (w/v) cholesteryl hemisuccinate (CHS, Sigma-Aldrich) at 4 °C for 3 h. Insoluble membrane debris was removed by centrifugation at 110,000 ×g for 40 min. The supernatant was applied to agarose beads conjugated with anti-GFP nanobody and rotated at 4°C for 2.5 h. The beads were rinsed three times in 20 mM Tris-HCl pH 7.5, 500 mM NaCl, 0.025% (w/v) DDM and 0.005% (w/v) CHS, then rinsed three times in 10 mM adenosine triphosphate (ATP, Macklin), 10 mM MgCl2, 20 mM Tris-HCl pH 7.5, 200 mM NaCl, 0.025% (w/v) DDM and 0.005% (w/v) CHS. After that, the beads were washed with W buffer, which contains 20 mM Tris-HCl pH 7.5, 200 mM NaCl, and 0.0063% (w/v) glycol-diosgenin (GDN, Anatrace). drCALHM1 protein was eluted with W buffer containing TEV protease overnight at 4 °C, and concentrated for further purified by size-exclusion chromatography on a Superose 6 Increase 10/300 GL column (GE Healthcare) in 20 mM Tris-HCl pH 7.5, 200 mM NaCl, 0.0063% GDN with 2 mM Ca^2+^. The peak fractions were pooled, concentrated to 6 mg/ml by a 100-kDa cut-off Centricon (Merck Millipore), and flash frozen for further cryo-EM grid preparation.

### Cryo-EM sample preparation and imaging

An aliquot of 2.5 microliters of purified drCALHM1 sample was applied onto a glow discharged holey carbon film grid (200 mesh, R2/1, Quantifoil). The grid was blotted and flash-frozen in liquid ethane with FEI Vitrobot Mark IV. The grid was loaded onto an FEI Titan Krios electron microscope operated at 300 kV accelerating voltage. Image stacks were recorded on a K2 Summit direct electron counting detector (Gatan) set in counting mode using SerialEM (http://bio3d.colorado.edu/SerialEM/) with a defocus range of 1.5 and 2.5 μm at a pixel size of 1.014 Å/pixel. A total of 40 frames were acquired in 8 seconds for each image stack, giving a total electron dose of 51.2 e^-^/A^2^. Finally, 2292 image stacks were acquired in an imaging session of 24 hours.

### Image processing

The recorded image stacks were processed by MotionCor2 (Zheng et al, 2017) for a 5 x 5 patch drift correction with dose weighting. The non-dose weighted images were used for CTF estimation by CTFFIND 4 (Rohou & Grigorieff, 2015) Images of poor quality were removed before particle picking. A total number of 801,380 particles were semi-automatically picked from dose weighted images by Gautomatch (https://www.mrc-lmb.cam.ac.uk/kzhang/) and extracted by RELION-3 (Fernandez-Leiro & Scheres, 2017) in a box size of 260 pixels. A round of 2D classification were performed in RELION-3 to remove contaminations, ice, and bad particles, yielding 482,632 good particles. The selected particles were then used to generate the initial model with an ab initio method with C1 symmetry in CryoSPARC-2 (https://cryosparc.com/). The initial model was pass-filtered to 30 Å resolution and applied to the first round of 3D classification in RELION-3. The particles were divided into 3 subsets, only 1 of the 3 subsets showed a 7-fold symmetry structure. Particles in this subset were selected for 3D auto-refine with C7 symmetry. Two more rounds of 3D classification by local angular search with C7 symmetry yielded 23,073 particles in well-defined class that showed CTH domain in the map. Another round of 3D auto-refine was performed with C7 symmetry, and generated reconstruction at 3.4 Å resolution. Per-particle motion correction was carried out using Bayesian polishing in RELION-3. The shiny, polished particles were then transferred cryosparc-2 for 3D classification using Heterogenous Refinement with C7 symmetry, 12,671 particles were selected and then refined to 3.2 Å resolution in CryoSPARC-2 using non-uniform refinement. The stated resolutions were evaluated using the “gold-standard” FSC = 0.143 criterion. The local resolution was calculated by ResMap (Swint-Kruse & Brown, 2005) or using two cryoEM maps independently refined from halves of data. For structures of drCALHM1-hepta-noLCH and drCALHM1-octa-noLCH, details of data process were listed in Supplemental Figure 2. Data collection and reconstruction statistics are presented in Supplemental Table 1.

### Model building and refinement

Initially the monomer structure of drCALHM1 octamer was rigid body fitted into the map obtained at 3.2 Å resolution. The fitted model was rebuilt manually using COOT optimizing the fit and subjected to global refinement and minimization in real space using the module “real_space_refinement” in PHENIX (Adams et al, 2010). The quality of the model was assessed with MolProbity (Adams et al, 2010). The final model exhibited good geometry, as indicated by the Ramachandran plot. The pore radius was calculated using HOLE (http://www.holeprogram.org/). Refinement statistics of three drCALHM1 structures are shown in Table S1.

### Electrophysiology

HEK293T cells were transiently transfected with GFP-tagged drCALHM1 for 16 to 24 h at 37°C with 5% CO2. Transfected cells were then digested by trypsin and plated onto 35-mm dishes for cultivation for at least 3 h before electrophysiology. The glass pipettes were pulled to a suitable shape using a P-97 glass microelectrode puller (Sutter Instrument, Novato) and polished with an MF-830 (Narishige, Tokyo, Japan). For the whole-cell configuration of recording drCALHM1-3C-GFP currents, all internal pipette solutions were 140 mM CsF, 6 mM MgCl2, 1 mM CaCl2, 11 mM EGTA, 2 mM TEA^+^, and 10 mM HEPES (pH adjusted to 7.3 with NaOH and methanesulfonic acid, ∼310 mOsm). The resistance of the pipette was 3 to 5 MΩ after being filled with the internal recording solution. For the whole-cell configuration of recording currents, the external bath solution was 145 mM NaCl, 5.4 mM KCl 10 mM TEA^+^, 10 mM glucose, 1.5 mM CaCl2, 1 mM MgCl2, and 10 HEPES (pH adjusted to 7.4 with NaOH and methanesulfonic acid, ∼330 mOsm). All experiments were conducted at room temperature with the stimulation voltage: 25 ms voltage step to −100 mV from a potential of 0 mV, followed by a ramp voltage increasing from −100 to +60 mV in 5 s. Whole-cell currents were amplified with an Axopatch 700B and digitized with a Digidata 1550A system (Molecular Devices, Sunnyvale). All currents were sampled at 10 kHz and low-pass filtered at 2 kHz through pCLAMP software (Molecular Devices, Sunnyvale). Origin 9.0 software (OriginLab Corp., Northampton) was also used for data analysis.

### In cell crosslinking experiment

HEK293T cells were grown in Dulbecco’s modified Eagle’s medium (DMEM) with 10% FBS at 37 ℃ in 5% CO2. When the confluence of cells reached 70-80%, 6 μg plasmid with a 2× FLAG tag at the C-terminus was transfected per 10 cm dish. After 24 h, HEK293T cells were washed three times with cold PBS.

To carry out a cross-linking experiment using disuccinimidyl suberate (DSS, Thermo), 4 mL of different DSS concentrations in PBS were added to each 10 cm dish for 30 min at room temperature. After that, 5 mL PBS containing 20 mM Tris-HCl (pH 7.5) was added at 4°C for 15 min to quench crosslinking. The cells were then collected and suspended in 500 μL of lysis buffer containing 20 mM Tris-HCl pH 7.5, 200 mM NaCl, and 1× protease inhibitor cocktail (Roche) and solubilized in 0.5% (w/v) lauryl maltose neopentyl glycol (LMNG, Anatrace) at 4°C for 2 h. The lysates were centrifuged at 18,000 ×g for 30 min at 4°C. Then, the supernatants were mixed with the anti-DYKDDDDK tag affinity gel (Biolegend) at 4°C for 4 h. The resin was rinsed three times with wash buffer containing 20 mM Tris-HCl pH 7.5, 200 mM NaCl, and 0.003% (w/v) LMNG. The protein was eluted overnight with wash buffer supplemented with 500-600 μg/mL FLAG peptide. The eluted protein was denatured with a final concentration of 2% SDS and heated to 65°C for 10 min, resuspended in 1× LDS sample buffer and 100 mM DTT, and heated at 65°C for another 10 min.

The method of formaldehyde crosslinking started to add 4 mL of different concentrations of formaldehyde in PBS to each 10 cm dish to crosslink the protein for 30 min at room temperature. After that, 5 mL buffer containing 100 mM Tris-HCl (pH 8.0) and 150 mM NaCl was added at room temperature for 10 min to quench crosslinking. After completely draining the liquid in the dish, the cells were collected and suspended in 500 μL lysis buffer containing 10 mM sodium phosphate pH 7.4, 100 mM NaCl, 25 mM TCEP (Aladdin), 50 mM N-ethylmaleimide (Sigma-Aldrich), and 1× protease inhibitor cocktail (Roche), lysed by adding 0.5% (w/v) lauryl maltose neopentyl glycol (LMNG, Anatrace), and rotated gently at room temperature. After 2 h, the lysate was heated at 37°C for 30 min and then centrifuged at 17,000 ×g for 10 min at room temperature. Then, the supernatants were mixed with the anti-DYKDDDDK tag affinity gel (Biolegend) at room temperature for 4 h. The resin was rinsed three times with wash buffer containing 10 mM sodium phosphate pH 7.4, 100 mM NaCl, 25 mM TCEP, 50 mM N-ethylmaleimide, and 0.003% (w/v) LMNG. Then, the protein was eluted overnight with wash buffer supplemented with 500-600 μg/mL FLAG peptide. The eluted protein was added to a final concentration of 2% SDS and heated to 65°C for 10 min. Then, the protein was resuspended in 1× LDS sample buffer and 25 mM TCEP and heated at 65°C for another 10 min.

Finally, 15 μL of each sample was loaded in a 4-12% Bis-Tris SurePAGE gel (GenScript) using MOPS running buffer and then transferred onto a PVDF membrane (Millipore). The primary antibody was anti-FLAG (Sigma-Aldrich), and anti-mouse (Abcam) was used as the secondary antibody.

### Molecular dynamics simulations

The molecular structure of the heptamer was prepared for the MD simulations. The missing residues (21-24, 93-94, and 205-207) were built by homology modeling using Modeler 9.24 (Webb & Sali, 2016). The residues are protonated at neutral pH. The heptamer was inserted into 160 Å × 160 Å POPC bilayers with their pore axis aligned parallel to the z axis through visualized operations in VMD (Humphrey et al, 1996).

The system was then solvated in TIP3P water boxes (Jorgensen et al, 1983) and neutralized by 0.15 M NaCl. The final system of the heptamer consisted of 343,897 atoms. Simulations were performed with Amber 2018 (D.A. Case et al, 2018) by using AMBER force field FF14SB (Maier et al, 2015) for proteins and LIPID14 (Dickson et al, 2014) for POPC. A 12 Å cut-off was set for the nonbonded interactions. The SHAKE algorithm (Ryckaert et al, 1977) integration was used to constrain the covalent bonds involving hydrogen atoms, and the Particle Mesh Ewald (PME) algorithm (Darden et al, 1993) was applied to treat long-range electrostatic interactions. The time step was set to 2 fs.

The initial energy minimization, thermalization, and a series of equilibrations were performed for the systems. First, each system was minimized for 10,000 steps. Second, the thermalization of each system heating from 0 K to 310 K was carried out in 500 ps using the Langevin thermostat (Pastor et al, 1988). The proteins and lipid head groups were fixed with a constraint of 50 kcal·mol^−1^·Å^−2^. Third, a series of equilibrations were performed for each system.: The POPC bilayer was equilibrated for 30 ns with the proteins constrained (50 kcal·mol^−1^·Å^−2^). After that, the missing residues were optimized for 30 ns, and the other residues of proteins were constrained (50 kcal·mol^−1^·Å^−2^). Finally, all atoms in the heptamer system were released and equilibrated for 20 ns with no constraints. The frames were saved every 500 steps for analysis.

Supervised MD simulation incorporates a tabu-like supervision algorithm on the reaction coordinate into a conventional MD simulation (Deganutti et al, 2020; Salmaso et al, 2017), accelerating the process of the “up-down” motion of the N-helix. Protomer 1 in the heptamer system was replaced with the monomer of the octamer (Target0) to obtain the reference structure. Then, we calculated the RMSD of the N-helix of protomer 1 in the current structure compared with Target0. The in-house Python script was employed to monitor the RMSD and process the MD simulation with the Amber engine (D.A. Case et al, 2018). The supervised MD was carried out in three main steps: (i) the 600-ps conventional MD simulations were performed and arrived at the checkpoint; (ii) 8 snapshots in 75-ps intervals were extracted from the trajectory, and the RMSD of each frame was calculated and collected; and (iii) if the slope of 8 values of RMSD was negative, the next 600-ps simulation was performed to reach the next checkpoint. Otherwise, the supervised MD simulation restarts from the checkpoint using the velocities randomly assigned. We performed an 80-ns supervised MD simulation, and then the supervised MD procedure was stopped. Only the productive steps were saved for analysis.

Targeted MD simulations (Schlitter et al, 1994; Xiao et al, 2015) can also accelerate the transition process by using a constraint. We carried out the targeted MD in the isothermal-isobaric (NPT) ensemble. Here, the coordinates of the N-helix in the “up” and “down” positions of protomer 1 were used as the initial and target positions, respectively. The targeted MD simulations were performed for 1 ns with force constants of 0.02 kcal·mol^−1^·Å^−2^. The frames were saved every 500 steps for analysis. As controls, a 100-ns conventional MD simulation was performed.

The coarse-grained MARTINI model (Marrink et al, 2007; Monticelli et al, 2008) for the heptamer and the octamer were built, respectively, with the ElNeDyn elastic network (Periole et al, 2009) applied to proteins. The CG protein coordinates of the heptamer and the octamer were separately positioned in the center of the simulation box of size 16 x 16 x 18 nm^3^ and 17 x 17 x 18 nm^3^ with their pore axis aligned parallel to the z axis and embedded in a POPC bilayer using the insane script (Wassenaar et al, 2015). Each system was solvated at 0.15 M NaCl concentration on either side and 0.02 M CaCl2 concentration on extracellular side. Coarse-grained simulations were carried out using Gromacs v.2020.2. The Martini v.2.2 force field was used for protein (Marrink et al, 2007; Monticelli et al, 2008) and the Martini v.2.0 force field was used for POPC and ions (Marrink et al, 2004). The simulations were run in the NPT ensemble. The time step was set to 20 fs. By using the velocity-rescaling thermostat (Bussi et al, 2007) with coupling constants of τT = 1.0 ps, the temperature was maintained at 310 K. The pressure was controlled at 1 bar by a semi-isotropic Parrinello-Rahman barostat (Parrinello & Rahman, 1982) with a coupling constant of τP=12.0 ps. The type of constrain applied to bonds was the LINCS algorithm (Hess et al, 1998). A Verlet cut- off scheme was used, and the PME method (Darden et al, 1993) was applied to calculate long-range electrostatic interactions. Periodic boundary condition was used in x and y axis. Ten microliters of equilibrium simulation trajectory data were collected for each system. The all-atom and coarse-grained MD simulations in this work were repeated 3 times.

### Accession Numbers

Atomic coordinates have been deposited in the Protein Data Bank under accession number 7DSE, 7DSD and 7DSC for drCALHM1-hepta-LCH, drCALHM1- hepta-noLCH and drCALHM1-octa-noLCH, respectively. The cryo-EM density maps have been deposited in the Electron Microscopy Data Bank under accession number EMD-30832, EMD-30831 and EMD-30830 for drCALHM1-hepta-LCH, drCALHM1- hepta-noLCH and drCALHM1-octa-noLCH, respectively. Additional data related to this paper may be requested from the authors.

## 1 Overall quality at a glance

The following experimental techniques were used to determine the structure:*ELECTRON MICROSCOPY*

The reported resolution of this entry is 2.90 Å.

Percentile scores (ranging between 0-100) for global validation metrics of the entry are shown in the following graphic. The table shows the number of entries on which the scores are based.

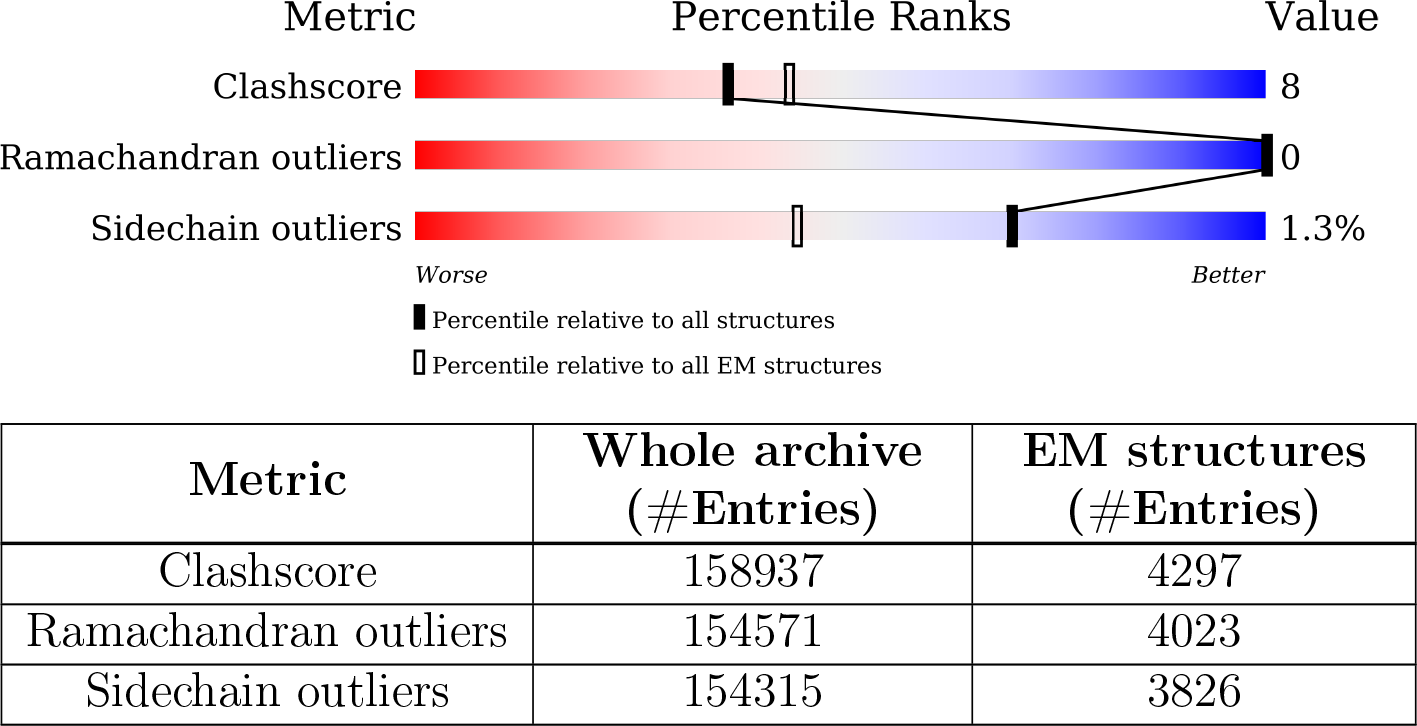

The table below summarises the geometric issues observed across the polymeric chains and their fit to the map. The red, orange, yellow and green segments of the bar indicate the fraction of residues that contain outliers for *>*=3, 2, 1 and 0 types of geometric quality criteria respectively. A grey segment represents the fraction of residues that are not modelled. The numeric value for each fraction is indicated below the corresponding segment, with a dot representing fractions *<*=5% The upper red bar (where present) indicates the fraction of residues that have poor fit to the EM map (all-atom inclusion *<* 40%). The numeric value is given above the bar.

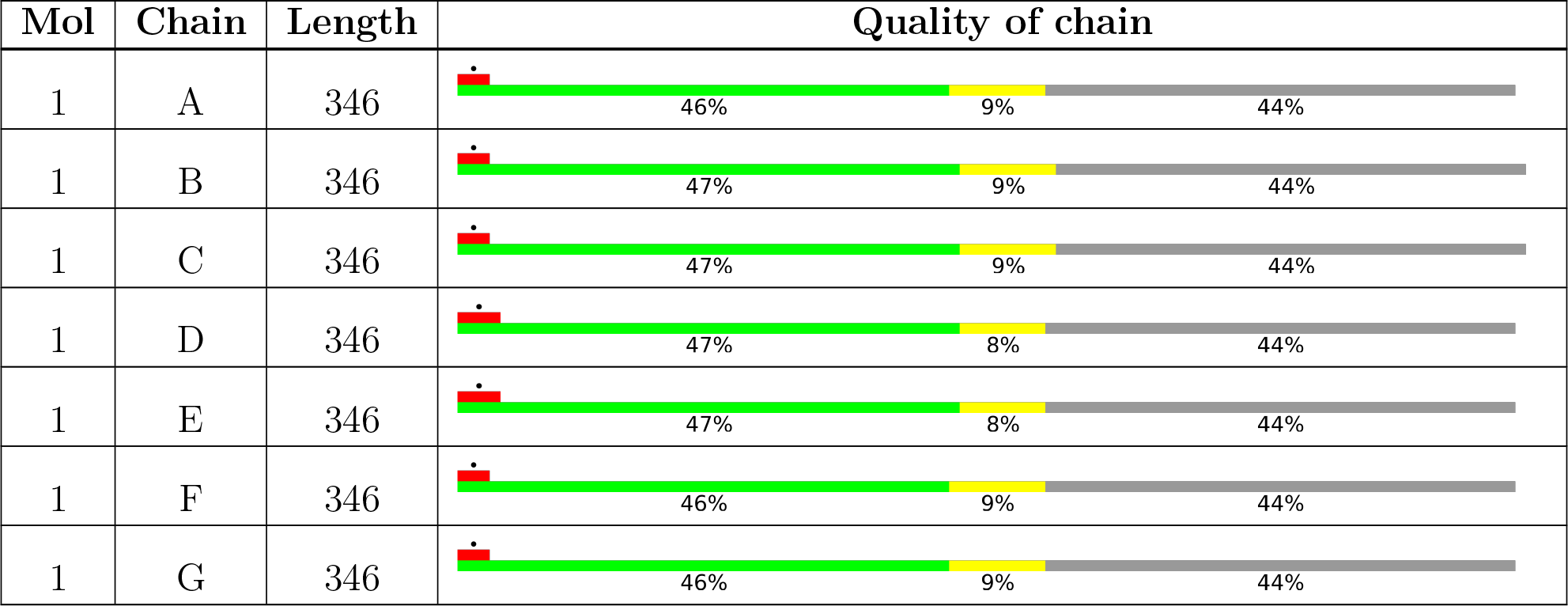

## 2 Entry composition

There are 2 unique types of molecules in this entry. The entry contains 10542 atoms, of which 0 are hydrogens and 0 are deuteriums.

In the tables below, the AltConf column contains the number of residues with at least one atom in alternate conformation and the Trace column contains the number of residues modelled with at most 2 atoms.

• Molecule 1 is a protein called Calcium homeostasis modulator 1.

| Mol | Chain | Residues | Atoms |  |  |  |  | AltConf | Trace |
| --- | --- | --- | --- | --- | --- | --- | --- | --- | --- |
|  |  |  | Total | C | N | O | S |  |  |
| 1 | A | 193 | Total<br>1492 | 974 | 241 | 264 | 13 | 0 | 0 |
| 1 | B | 193 | Total<br>1492 | 974 | 241 | 264 | 13 | 0 | 0 |
| 1 | C | 193 | Total<br>1492 | 974 | 241 | 264 | 13 | 0 | 0 |
| 1 | D | 193 | Total<br>1492 | 974 | 241 | 264 | 13 | 0 | 0 |
| 1 | E | 193 | Total<br>1492 | 974 | 241 | 264 | 13 | 0 | 0 |
| 1 | F | 193 | Total<br>1492 | 974 | 241 | 264 | 13 | 0 | 0 |
| 1 | G | 193 | Total<br>1492 | 974 | 241 | 264 | 13 | 0 | 0 |

• Molecule 2 is 2-acetamido-2-deoxy-beta-D-glucopyranose (three-letter code: NAG) (formula: C_8_H_15_NO_6_).

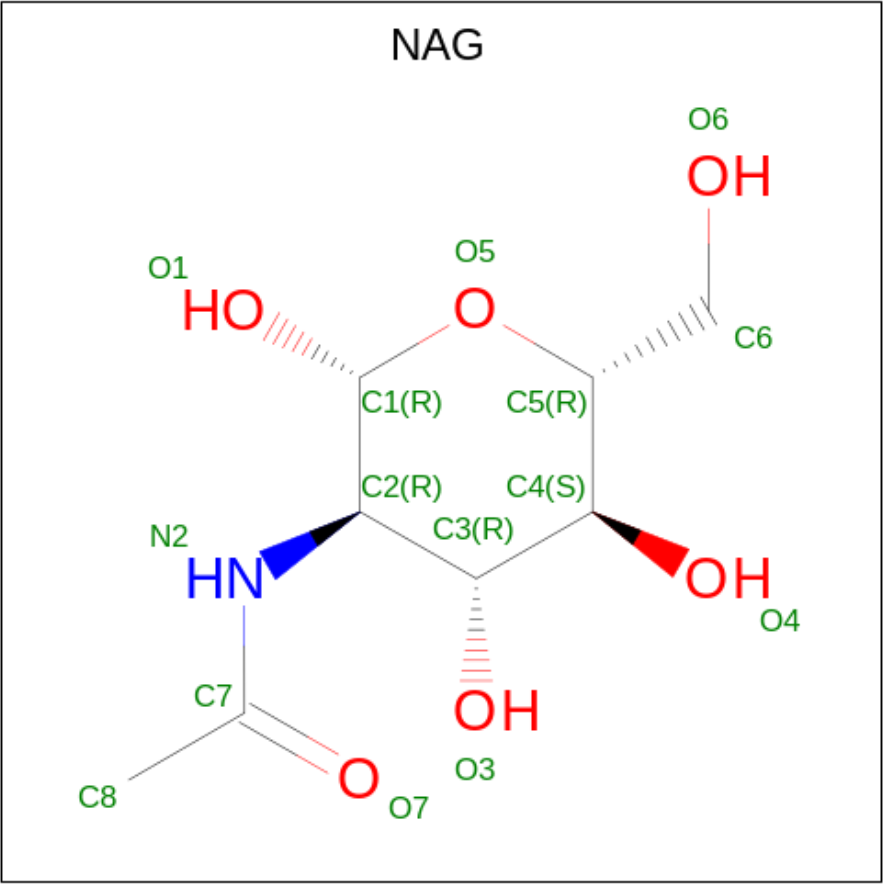

| Mol | Chain | Residues | Atoms |  |  |  | AltConf |
| --- | --- | --- | --- | --- | --- | --- | --- |
| 2 | A | 1 | Total | C | N | O | 0 |
|  |  |  | 14 | 8 | 1 | 5 |  |
| 2 | B | 1 | Total | C | N | O | 0 |
|  |  |  | 14 | 8 | 1 | 5 |  |
| 2 | C | 1 | Total | C | N | O | 0 |
|  |  |  | 14 | 8 | 1 | 5 |  |
| 2 | D | 1 | Total | C | N | O | 0 |
|  |  |  | 14 | 8 | 1 | 5 |  |
| 2 | E | 1 | Total | C | N | O | 0 |
|  |  |  | 14 | 8 | 1 | 5 |  |
| 2 | F | 1 | Total | C | N | O | 0 |
|  |  |  | 14 | 8 | 1 | 5 |  |
| 2 | G | 1 | Total | C | N | O | 0 |
|  |  |  | 14 | 8 | 1 | 5 |  |

## 3 Residue-property plots

These plots are drawn for all protein, RNA, DNA and oligosaccharide chains in the entry. The first graphic for a chain summarises the proportions of the various outlier classes displayed in the second graphic. The second graphic shows the sequence view annotated by issues in geometry and atom inclusion in map density. Residues are color-coded according to the number of geometric quality criteria for which they contain at least one outlier: green = 0, yellow = 1, orange = 2 and red = 3 or more. A red diamond above a residue indicates a poor fit to the EM map for this residue (all-atom inclusion < 40%). Stretches of 2 or more consecutive residues without any outlier are shown as a green connector. Residues present in the sample, but not in the model, are shown in grey.

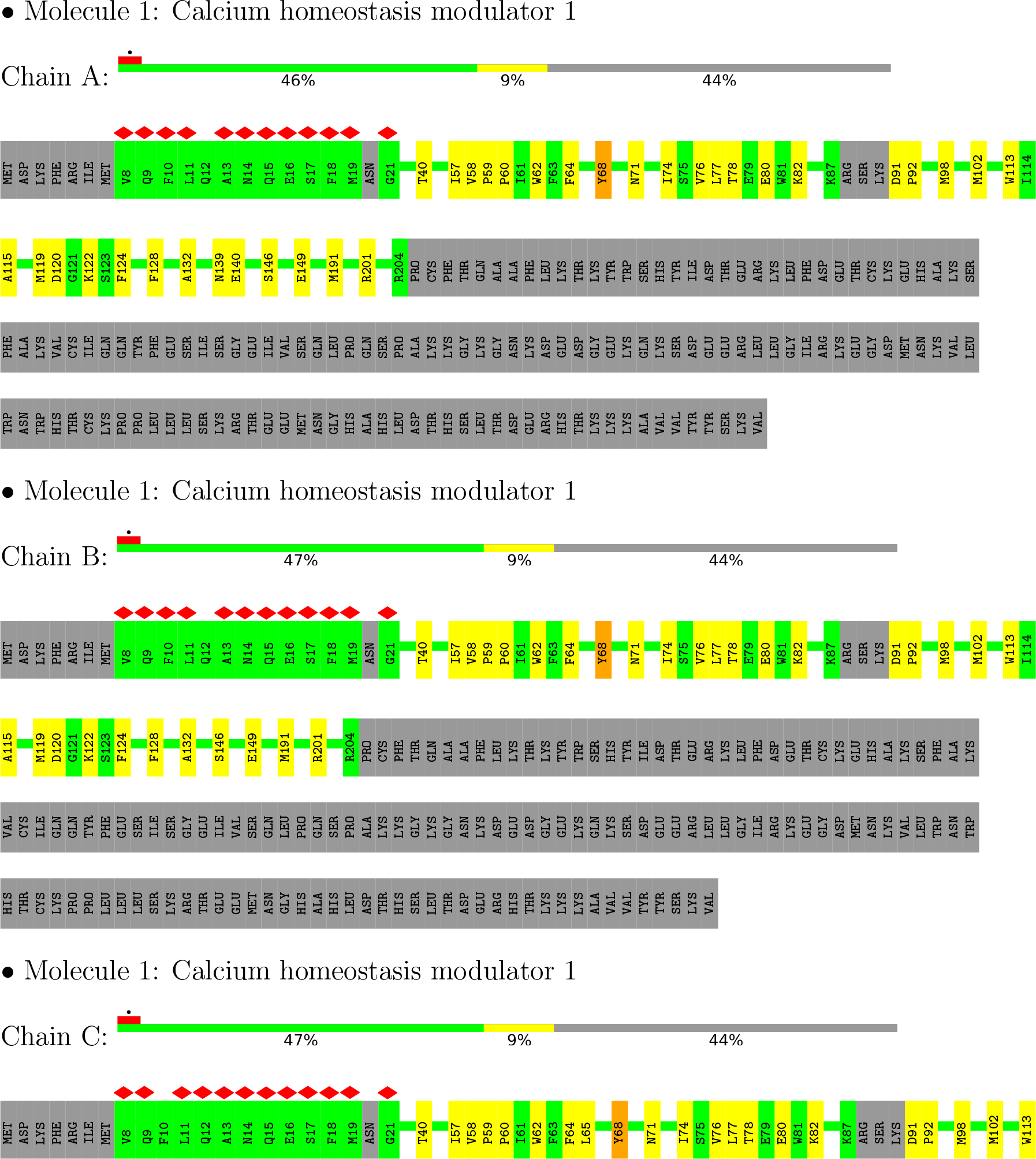
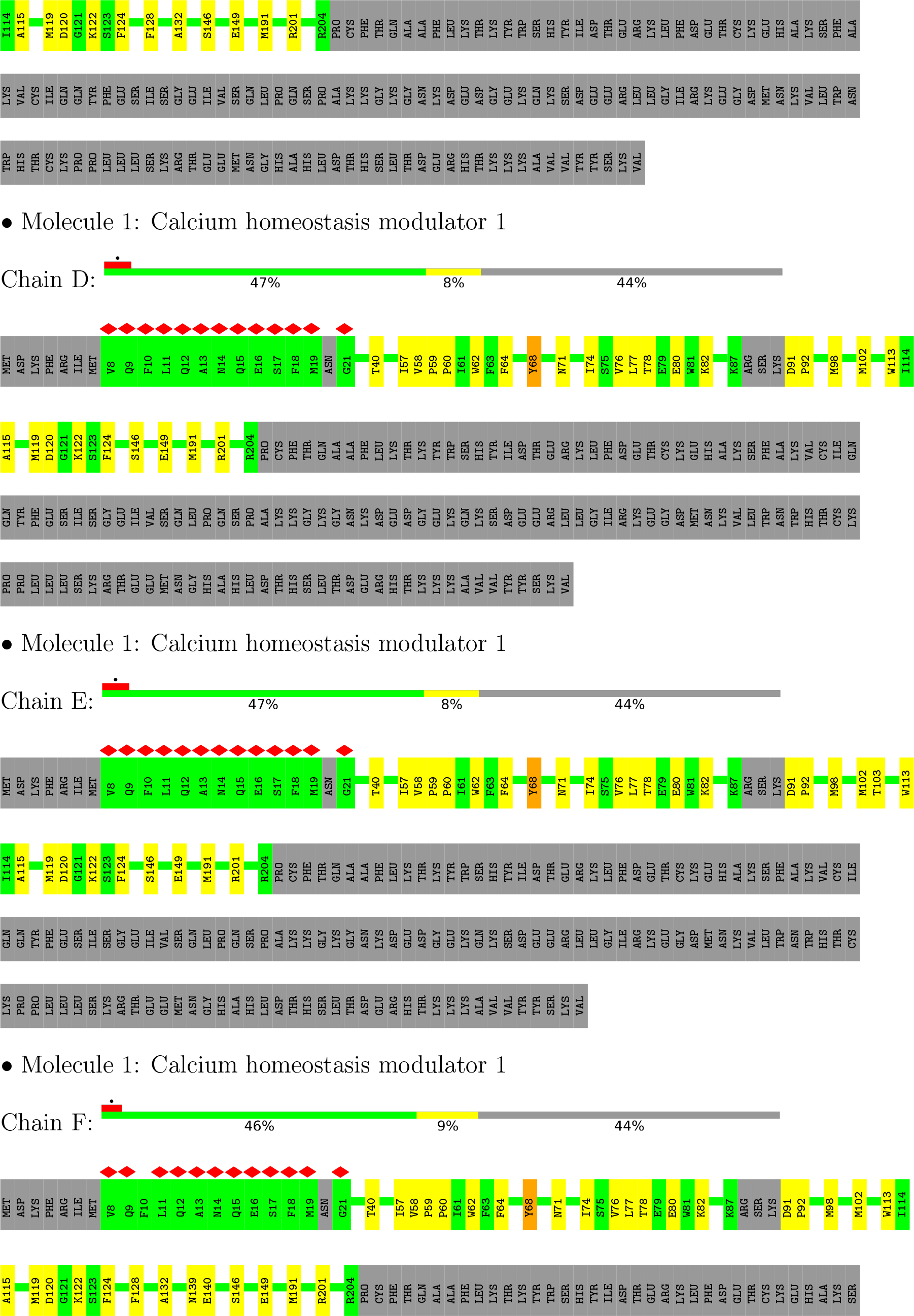
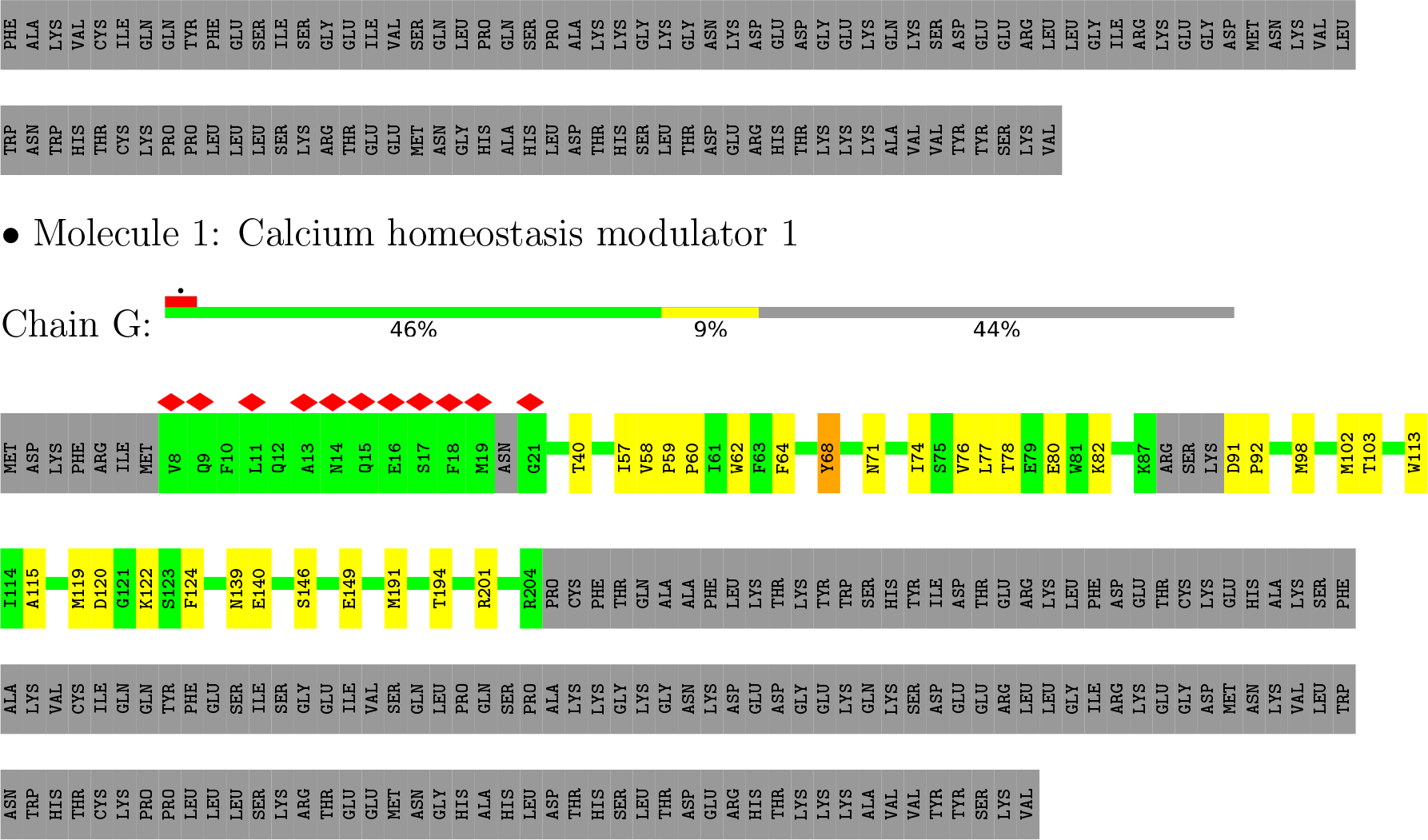

## 4 Experimental information

| Property | Value | Source |
| --- | --- | --- |
| EM reconstruction method | SINGLE PARTICLE | Depositor |
| Imposed symmetry | POINT, Not provided |  |
| Number of particles used | 56801 | Depositor |
| Resolution determination method | FSC 0.143 CUT-OFF | Depositor |
| CTF correction method | NONE | Depositor |
| Microscope | FEI TITAN KRIOS | Depositor |
| Voltage (kV) | 300 | Depositor |
| Electron dose ($e^-/\text{\AA}^2$) | 50 | Depositor |
| Minimum defocus (nm) | Not provided |  |
| Maximum defocus (nm) | Not provided |  |
| Magnification | Not provided |  |
| Image detector | GATAN K2 SUMMIT (4k x 4k) | Depositor |
| Maximum map value | 1.390 | Depositor |
| Minimum map value | -0.580 | Depositor |
| Average map value | -0.001 | Depositor |
| Map value standard deviation | 0.045 | Depositor |
| Recommended contour level | 0.154 | Depositor |
| Map size (Å) | 263.64, 263.64, 263.64 | wwPDB |
| Map dimensions | 260, 260, 260 | wwPDB |
| Map angles (°) | 90.0, 90.0, 90.0 | wwPDB |
| Pixel spacing (Å) | 1.014, 1.014, 1.014 | Depositor |

## 5 Model quality

### 5.1 Standard geometry

Bond lengths and bond angles in the following residue types are not validated in this section: NAG

The Z score for a bond length (or angle) is the number of standard deviations the observed value is removed from the expected value. A bond length (or angle) with *|Z| >* 5 is considered an outlier worth inspection. RMSZ is the root-mean-square of all Z scores of the bond lengths (or angles).

| Mol | Chain | Bond lengths |  | Bond angles |  |
| --- | --- | --- | --- | --- | --- |
| | | RMSZ | $\#\vert Z\vert > 5$ | RMSZ | $\#\vert Z\vert > 5$ |
| 1 | A | 0.28 | 0/1526 | 0.41 | 0/2072 |
| 1 | B | 0.28 | 0/1526 | 0.41 | 0/2072 |
| 1 | C | 0.28 | 0/1526 | 0.41 | 0/2072 |
| 1 | D | 0.28 | 0/1526 | 0.41 | 0/2072 |
| 1 | E | 0.29 | 0/1526 | 0.41 | 0/2072 |
| 1 | F | 0.28 | 0/1526 | 0.41 | 0/2072 |
| 1 | G | 0.28 | 0/1526 | 0.41 | 0/2072 |
| All | All | 0.28 | 0/10682 | 0.41 | 0/14504 |

There are no bond length outliers. There are no bond angle outliers. There are no chirality outliers.

There are no planarity outliers.

### 5.2 Too-close contacts

In the following table, the Non-H and H(model) columns list the number of non-hydrogen atoms and hydrogen atoms in the chain respectively. The H(added) column lists the number of hydrogen atoms added and optimized by MolProbity. The Clashes column lists the number of clashes within the asymmetric unit, whereas Symm-Clashes lists symmetry-related clashes.

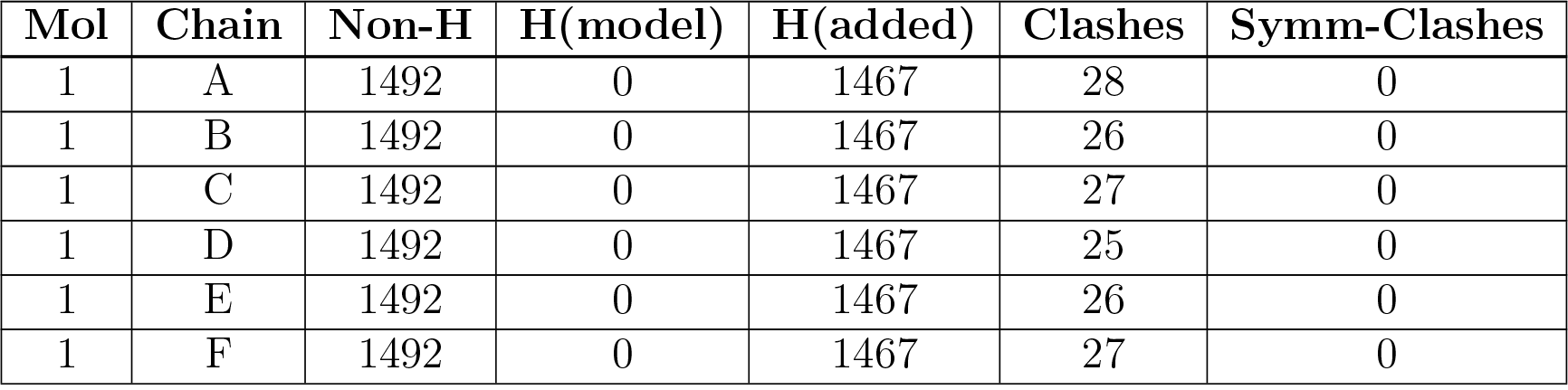
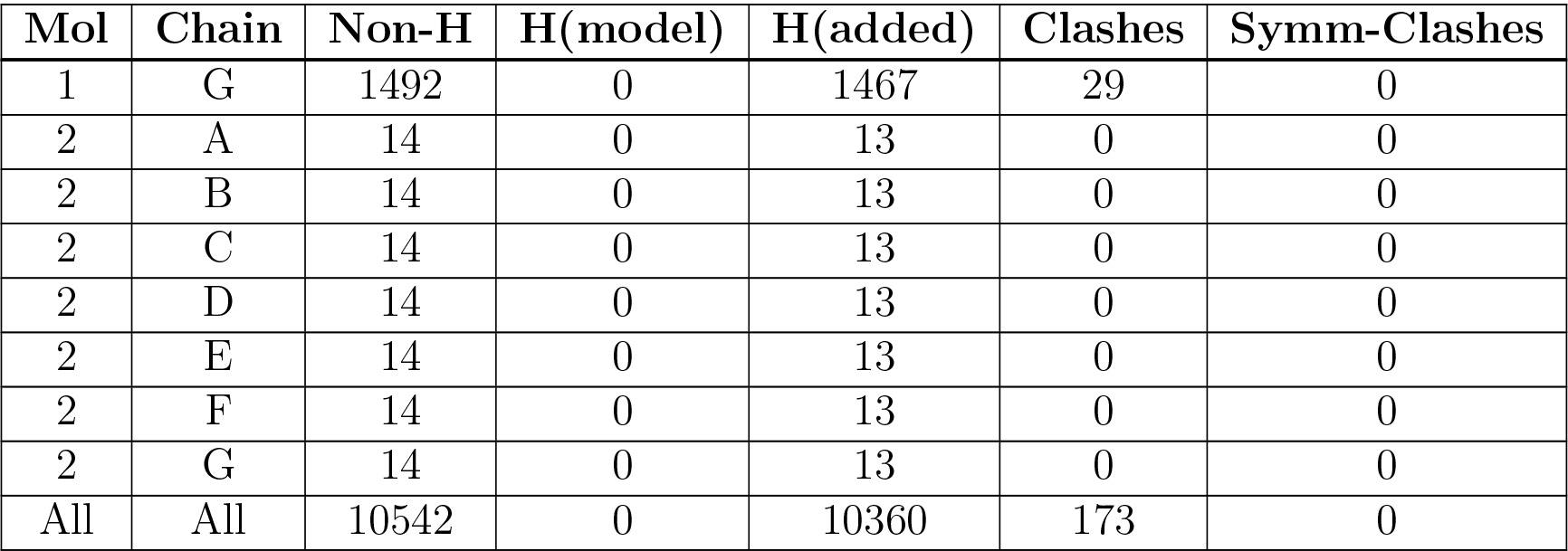

| Mol | Chain | Non-H | H(model) | H(added) | Clashes | Symm-Clashes |
| --- | --- | --- | --- | --- | --- | --- |
| 1 | A | 1492 | 0 | 1467 | 28 | 0 |
| 1 | B | 1492 | 0 | 1467 | 26 | 0 |
| 1 | C | 1492 | 0 | 1467 | 27 | 0 |
| 1 | D | 1492 | 0 | 1467 | 25 | 0 |
| 1 | E | 1492 | 0 | 1467 | 26 | 0 |
| 1 | F | 1492 | 0 | 1467 | 27 | 0 |
| 1 | G | 1492 | 0 | 1467 | 29 | 0 |
| 2 | A | 14 | 0 | 13 | 0 | 0 |
| 2 | B | 14 | 0 | 13 | 0 | 0 |
| 2 | C | 14 | 0 | 13 | 0 | 0 |
| 2 | D | 14 | 0 | 13 | 0 | 0 |
| 2 | E | 14 | 0 | 13 | 0 | 0 |
| 2 | F | 14 | 0 | 13 | 0 | 0 |
| 2 | G | 14 | 0 | 13 | 0 | 0 |
| All | All | 10542 | 0 | 10360 | 173 | 0 |

The all-atom clashscore is defined as the number of clashes found per 1000 atoms (including hydrogen atoms). The all-atom clashscore for this structure is 8.

All (173) close contacts within the same asymmetric unit are listed below, sorted by their clash magnitude.

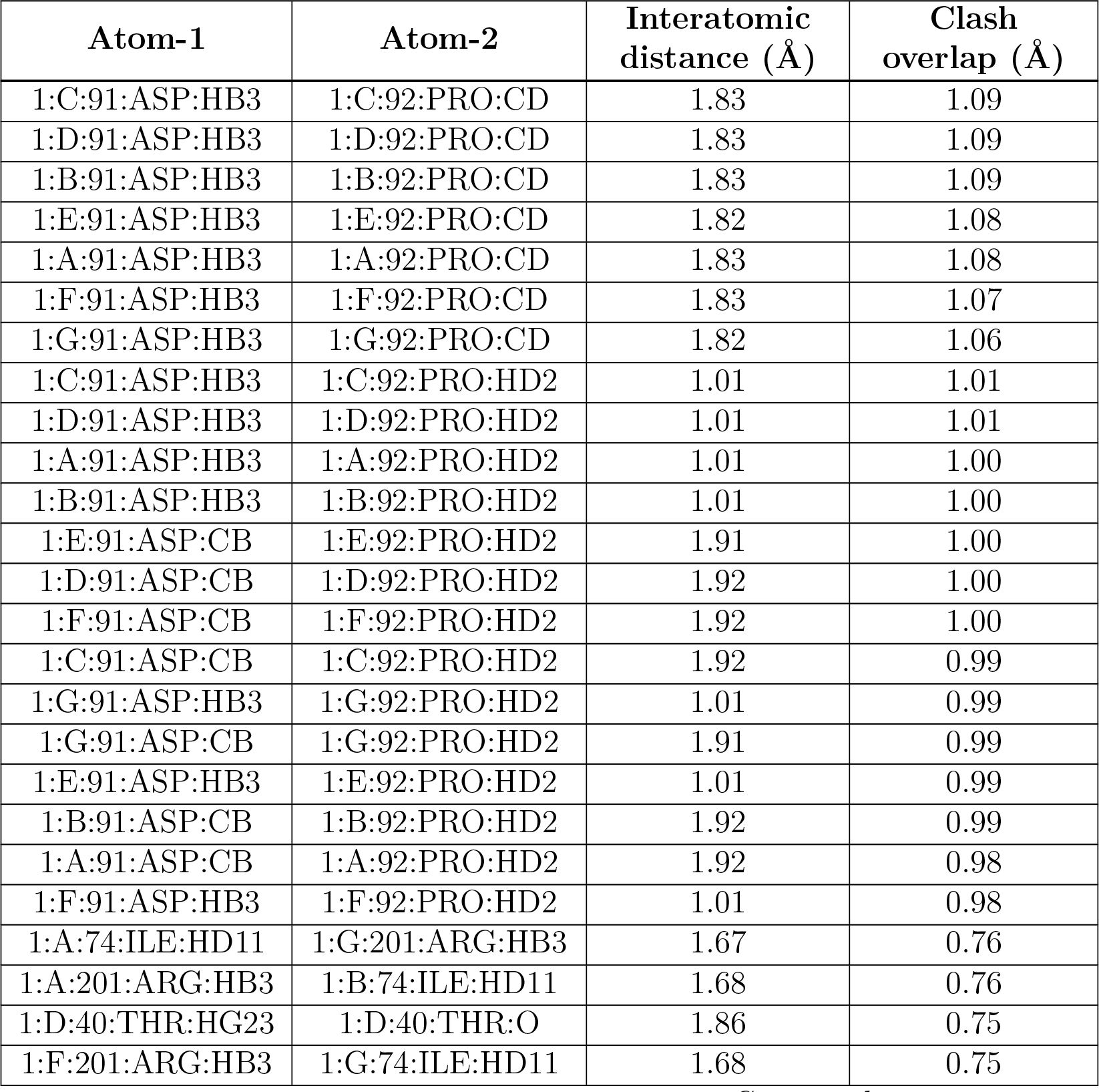
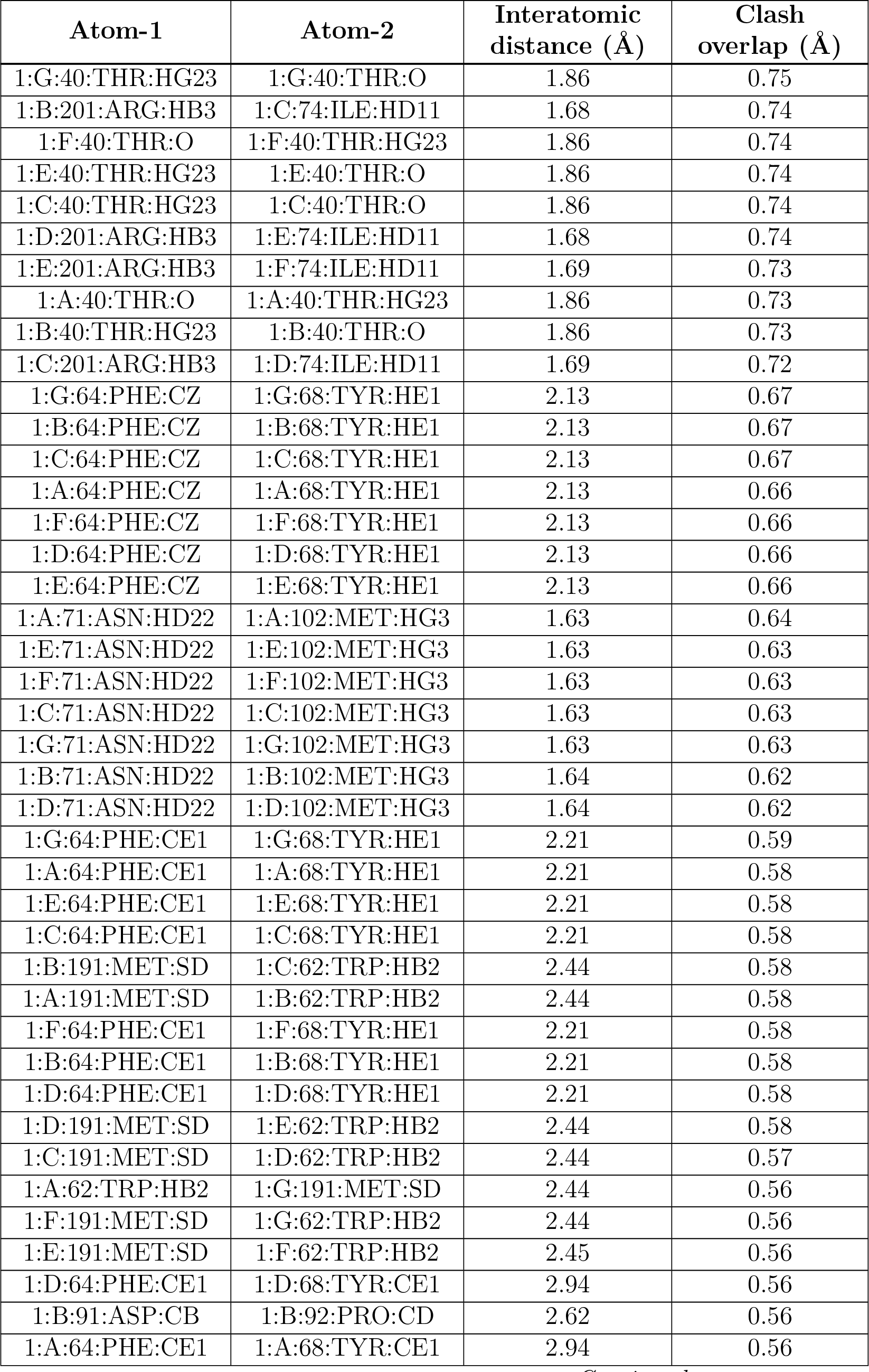
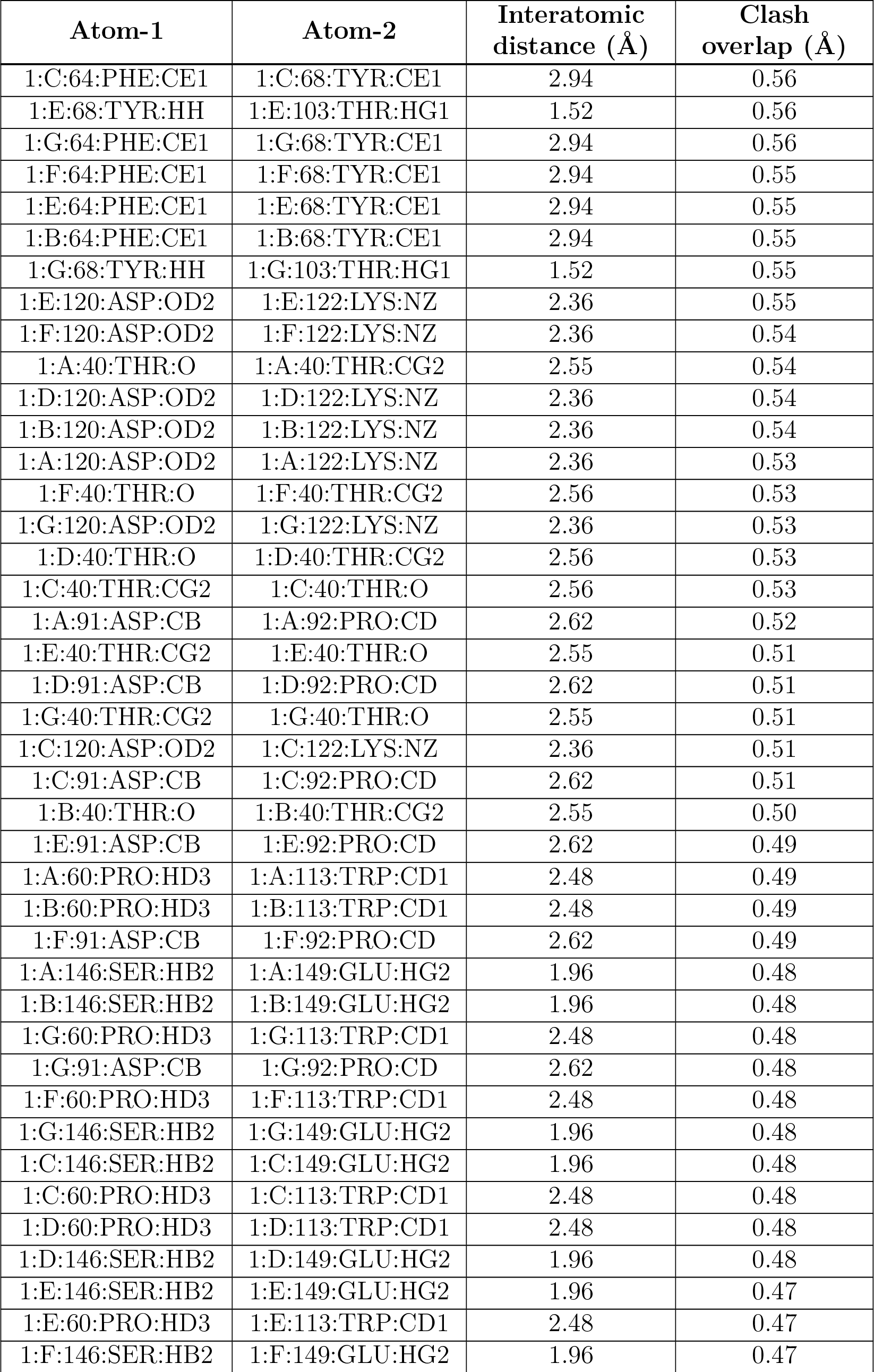
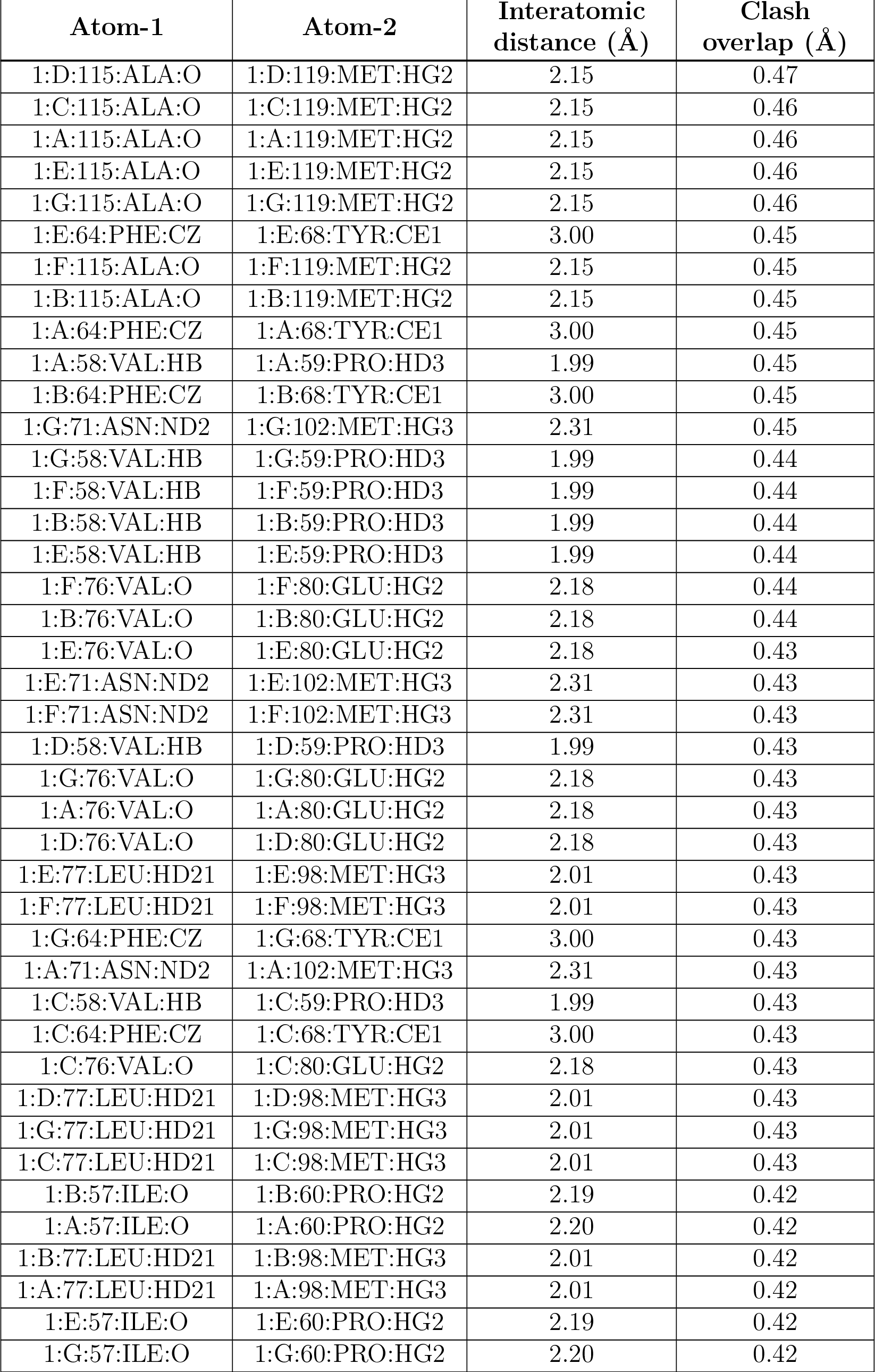
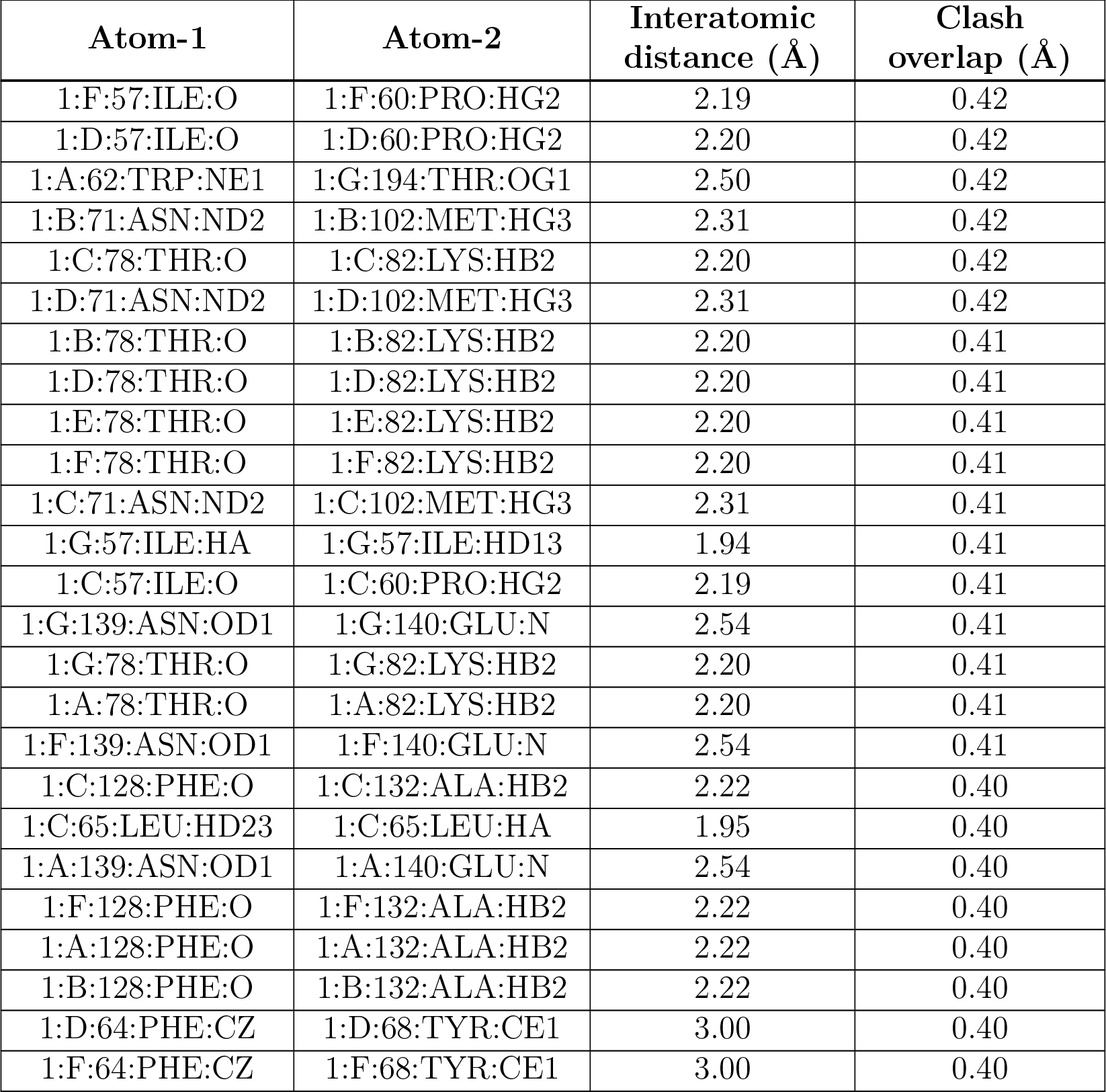

| Atom-1 | Atom-2 | Interatomic distance (Å) | Clash overlap (Å) |
| --- | --- | --- | --- |
| 1:C:91:ASP:HB3 | 1:C:92:PRO:CD | 1.83 | 1.09 |
| 1:D:91:ASP:HB3 | 1:D:92:PRO:CD | 1.83 | 1.09 |
| 1:B:91:ASP:HB3 | 1:B:92:PRO:CD | 1.83 | 1.09 |
| 1:E:91:ASP:HB3 | 1:E:92:PRO:CD | 1.82 | 1.08 |
| 1:A:91:ASP:HB3 | 1:A:92:PRO:CD | 1.83 | 1.08 |
| 1:F:91:ASP:HB3 | 1:F:92:PRO:CD | 1.83 | 1.07 |
| 1:G:91:ASP:HB3 | 1:G:92:PRO:CD | 1.82 | 1.06 |
| 1:C:91:ASP:HB3 | 1:C:92:PRO:HD2 | 1.01 | 1.01 |
| 1:D:91:ASP:HB3 | 1:D:92:PRO:HD2 | 1.01 | 1.01 |
| 1:A:91:ASP:HB3 | 1:A:92:PRO:HD2 | 1.01 | 1.00 |
| 1:B:91:ASP:HB3 | 1:B:92:PRO:HD2 | 1.01 | 1.00 |
| 1:E:91:ASP:CB | 1:E:92:PRO:HD2 | 1.91 | 1.00 |
| 1:D:91:ASP:CB | 1:D:92:PRO:HD2 | 1.92 | 1.00 |
| 1:F:91:ASP:CB | 1:F:92:PRO:HD2 | 1.92 | 1.00 |
| 1:C:91:ASP:CB | 1:C:92:PRO:HD2 | 1.92 | 0.99 |
| 1:G:91:ASP:HB3 | 1:G:92:PRO:HD2 | 1.01 | 0.99 |
| 1:G:91:ASP:CB | 1:G:92:PRO:HD2 | 1.91 | 0.99 |
| 1:E:91:ASP:HB3 | 1:E:92:PRO:HD2 | 1.01 | 0.99 |
| 1:B:91:ASP:CB | 1:B:92:PRO:HD2 | 1.92 | 0.99 |
| 1:A:91:ASP:CB | 1:A:92:PRO:HD2 | 1.92 | 0.98 |
| 1:F:91:ASP:HB3 | 1:F:92:PRO:HD2 | 1.01 | 0.98 |
| 1:A:74:ILE:HD11 | 1:G:201:ARG:HB3 | 1.67 | 0.76 |
| 1:A:201:ARG:HB3 | 1:B:74:ILE:HD11 | 1.68 | 0.76 |
| 1:D:40:THR:HG23 | 1:D:40:THR:O | 1.86 | 0.75 |
| 1:F:201:ARG:HB3 | 1:G:74:ILE:HD11 | 1.68 | 0.75 |
| 1:G:40:THR:HG23 | 1:G:40:THR:O | 1.86 | 0.75 |
| 1:B:201:ARG:HB3 | 1:C:74:ILE:HD11 | 1.68 | 0.74 |
| 1:F:40:THR:O | 1:F:40:THR:HG23 | 1.86 | 0.74 |
| 1:E:40:THR:HG23 | 1:E:40:THR:O | 1.86 | 0.74 |
| 1:C:40:THR:HG23 | 1:C:40:THR:O | 1.86 | 0.74 |
| 1:D:201:ARG:HB3 | 1:E:74:ILE:HD11 | 1.68 | 0.74 |
| 1:E:201:ARG:HB3 | 1:F:74:ILE:HD11 | 1.69 | 0.73 |
| 1:A:40:THR:O | 1:A:40:THR:HG23 | 1.86 | 0.73 |
| 1:B:40:THR:HG23 | 1:B:40:THR:O | 1.86 | 0.73 |
| 1:C:201:ARG:HB3 | 1:D:74:ILE:HD11 | 1.69 | 0.72 |
| 1:G:64:PHE:CZ | 1:G:68:TYR:HE1 | 2.13 | 0.67 |
| 1:B:64:PHE:CZ | 1:B:68:TYR:HE1 | 2.13 | 0.67 |
| 1:C:64:PHE:CZ | 1:C:68:TYR:HE1 | 2.13 | 0.67 |
| 1:A:64:PHE:CZ | 1:A:68:TYR:HE1 | 2.13 | 0.66 |
| 1:F:64:PHE:CZ | 1:F:68:TYR:HE1 | 2.13 | 0.66 |
| 1:D:64:PHE:CZ | 1:D:68:TYR:HE1 | 2.13 | 0.66 |
| 1:E:64:PHE:CZ | 1:E:68:TYR:HE1 | 2.13 | 0.66 |
| 1:A:71:ASN:HD22 | 1:A:102:MET:HG3 | 1.63 | 0.64 |
| 1:E:71:ASN:HD22 | 1:E:102:MET:HG3 | 1.63 | 0.63 |
| 1:F:71:ASN:HD22 | 1:F:102:MET:HG3 | 1.63 | 0.63 |
| 1:C:71:ASN:HD22 | 1:C:102:MET:HG3 | 1.63 | 0.63 |
| 1:G:71:ASN:HD22 | 1:G:102:MET:HG3 | 1.63 | 0.63 |
| 1:B:71:ASN:HD22 | 1:B:102:MET:HG3 | 1.64 | 0.62 |
| 1:D:71:ASN:HD22 | 1:D:102:MET:HG3 | 1.64 | 0.62 |
| 1:G:64:PHE:CE1 | 1:G:68:TYR:HE1 | 2.21 | 0.59 |
| 1:A:64:PHE:CE1 | 1:A:68:TYR:HE1 | 2.21 | 0.58 |
| 1:E:64:PHE:CE1 | 1:E:68:TYR:HE1 | 2.21 | 0.58 |
| 1:C:64:PHE:CE1 | 1:C:68:TYR:HE1 | 2.21 | 0.58 |
| 1:B:191:MET:SD | 1:C:62:TRP:HB2 | 2.44 | 0.58 |
| 1:A:191:MET:SD | 1:B:62:TRP:HB2 | 2.44 | 0.58 |
| 1:F:64:PHE:CE1 | 1:F:68:TYR:HE1 | 2.21 | 0.58 |
| 1:B:64:PHE:CE1 | 1:B:68:TYR:HE1 | 2.21 | 0.58 |
| 1:D:64:PHE:CE1 | 1:D:68:TYR:HE1 | 2.21 | 0.58 |
| 1:D:191:MET:SD | 1:E:62:TRP:HB2 | 2.44 | 0.58 |
| 1:C:191:MET:SD | 1:D:62:TRP:HB2 | 2.44 | 0.57 |
| 1:A:62:TRP:HB2 | 1:G:191:MET:SD | 2.44 | 0.56 |
| 1:F:191:MET:SD | 1:G:62:TRP:HB2 | 2.44 | 0.56 |
| 1:E:191:MET:SD | 1:F:62:TRP:HB2 | 2.45 | 0.56 |
| 1:D:64:PHE:CE1 | 1:D:68:TYR:CE1 | 2.94 | 0.56 |
| 1:B:91:ASP:CB | 1:B:92:PRO:CD | 2.62 | 0.56 |
| 1:A:64:PHE:CE1 | 1:A:68:TYR:CE1 | 2.94 | 0.56 |
| 1:C:64:PHE:CE1 | 1:C:68:TYR:CE1 | 2.94 | 0.56 |
| 1:E:68:TYR:HH | 1:E:103:THR:HG1 | 1.52 | 0.56 |
| 1:G:64:PHE:CE1 | 1:G:68:TYR:CE1 | 2.94 | 0.56 |
| 1:F:64:PHE:CE1 | 1:F:68:TYR:CE1 | 2.94 | 0.55 |
| 1:E:64:PHE:CE1 | 1:E:68:TYR:CE1 | 2.94 | 0.55 |
| 1:B:64:PHE:CE1 | 1:B:68:TYR:CE1 | 2.94 | 0.55 |
| 1:G:68:TYR:HH | 1:G:103:THR:HG1 | 1.52 | 0.55 |
| 1:E:120:ASP:OD2 | 1:E:122:LYS:NZ | 2.36 | 0.55 |
| 1:F:120:ASP:OD2 | 1:F:122:LYS:NZ | 2.36 | 0.54 |
| 1:A:40:THR:O | 1:A:40:THR:CG2 | 2.55 | 0.54 |
| 1:D:120:ASP:OD2 | 1:D:122:LYS:NZ | 2.36 | 0.54 |
| 1:B:120:ASP:OD2 | 1:B:122:LYS:NZ | 2.36 | 0.54 |
| 1:A:120:ASP:OD2 | 1:A:122:LYS:NZ | 2.36 | 0.53 |
| 1:F:40:THR:O | 1:F:40:THR:CG2 | 2.56 | 0.53 |
| 1:G:120:ASP:OD2 | 1:G:122:LYS:NZ | 2.36 | 0.53 |
| 1:D:40:THR:O | 1:D:40:THR:CG2 | 2.56 | 0.53 |
| 1:C:40:THR:CG2 | 1:C:40:THR:O | 2.56 | 0.53 |
| 1:A:91:ASP:CB | 1:A:92:PRO:CD | 2.62 | 0.52 |
| 1:E:40:THR:CG2 | 1:E:40:THR:O | 2.55 | 0.51 |
| 1:D:91:ASP:CB | 1:D:92:PRO:CD | 2.62 | 0.51 |
| 1:G:40:THR:CG2 | 1:G:40:THR:O | 2.55 | 0.51 |
| 1:C:120:ASP:OD2 | 1:C:122:LYS:NZ | 2.36 | 0.51 |
| 1:C:91:ASP:CB | 1:C:92:PRO:CD | 2.62 | 0.51 |
| 1:B:40:THR:O | 1:B:40:THR:CG2 | 2.55 | 0.50 |
| 1:E:91:ASP:CB | 1:E:92:PRO:CD | 2.62 | 0.49 |
| 1:A:60:PRO:HD3 | 1:A:113:TRP:CD1 | 2.48 | 0.49 |
| 1:B:60:PRO:HD3 | 1:B:113:TRP:CD1 | 2.48 | 0.49 |
| 1:F:91:ASP:CB | 1:F:92:PRO:CD | 2.62 | 0.49 |
| 1:A:146:SER:HB2 | 1:A:149:GLU:HG2 | 1.96 | 0.48 |
| 1:B:146:SER:HB2 | 1:B:149:GLU:HG2 | 1.96 | 0.48 |
| 1:G:60:PRO:HD3 | 1:G:113:TRP:CD1 | 2.48 | 0.48 |
| 1:G:91:ASP:CB | 1:G:92:PRO:CD | 2.62 | 0.48 |
| 1:F:60:PRO:HD3 | 1:F:113:TRP:CD1 | 2.48 | 0.48 |
| 1:G:146:SER:HB2 | 1:G:149:GLU:HG2 | 1.96 | 0.48 |
| 1:C:146:SER:HB2 | 1:C:149:GLU:HG2 | 1.96 | 0.48 |
| 1:C:60:PRO:HD3 | 1:C:113:TRP:CD1 | 2.48 | 0.48 |
| 1:D:60:PRO:HD3 | 1:D:113:TRP:CD1 | 2.48 | 0.48 |
| 1:D:146:SER:HB2 | 1:D:149:GLU:HG2 | 1.96 | 0.48 |
| 1:E:146:SER:HB2 | 1:E:149:GLU:HG2 | 1.96 | 0.47 |
| 1:E:60:PRO:HD3 | 1:E:113:TRP:CD1 | 2.48 | 0.47 |
| 1:F:146:SER:HB2 | 1:F:149:GLU:HG2 | 1.96 | 0.47 |
| 1:D:115:ALA:O | 1:D:119:MET:HG2 | 2.15 | 0.47 |
| 1:C:115:ALA:O | 1:C:119:MET:HG2 | 2.15 | 0.46 |
| 1:A:115:ALA:O | 1:A:119:MET:HG2 | 2.15 | 0.46 |
| 1:E:115:ALA:O | 1:E:119:MET:HG2 | 2.15 | 0.46 |
| 1:G:115:ALA:O | 1:G:119:MET:HG2 | 2.15 | 0.46 |
| 1:E:64:PHE:CZ | 1:E:68:TYR:CE1 | 3.00 | 0.45 |
| 1:F:115:ALA:O | 1:F:119:MET:HG2 | 2.15 | 0.45 |
| 1:B:115:ALA:O | 1:B:119:MET:HG2 | 2.15 | 0.45 |
| 1:A:64:PHE:CZ | 1:A:68:TYR:CE1 | 3.00 | 0.45 |
| 1:A:58:VAL:HB | 1:A:59:PRO:HD3 | 1.99 | 0.45 |
| 1:B:64:PHE:CZ | 1:B:68:TYR:CE1 | 3.00 | 0.45 |
| 1:G:71:ASN:ND2 | 1:G:102:MET:HG3 | 2.31 | 0.45 |
| 1:G:58:VAL:HB | 1:G:59:PRO:HD3 | 1.99 | 0.44 |
| 1:F:58:VAL:HB | 1:F:59:PRO:HD3 | 1.99 | 0.44 |
| 1:B:58:VAL:HB | 1:B:59:PRO:HD3 | 1.99 | 0.44 |
| 1:E:58:VAL:HB | 1:E:59:PRO:HD3 | 1.99 | 0.44 |
| 1:F:76:VAL:O | 1:F:80:GLU:HG2 | 2.18 | 0.44 |
| 1:B:76:VAL:O | 1:B:80:GLU:HG2 | 2.18 | 0.44 |
| 1:E:76:VAL:O | 1:E:80:GLU:HG2 | 2.18 | 0.43 |
| 1:E:71:ASN:ND2 | 1:E:102:MET:HG3 | 2.31 | 0.43 |
| 1:F:71:ASN:ND2 | 1:F:102:MET:HG3 | 2.31 | 0.43 |
| 1:D:58:VAL:HB | 1:D:59:PRO:HD3 | 1.99 | 0.43 |
| 1:G:76:VAL:O | 1:G:80:GLU:HG2 | 2.18 | 0.43 |
| 1:A:76:VAL:O | 1:A:80:GLU:HG2 | 2.18 | 0.43 |
| 1:D:76:VAL:O | 1:D:80:GLU:HG2 | 2.18 | 0.43 |
| 1:E:77:LEU:HD21 | 1:E:98:MET:HG3 | 2.01 | 0.43 |
| 1:F:77:LEU:HD21 | 1:F:98:MET:HG3 | 2.01 | 0.43 |
| 1:G:64:PHE:CZ | 1:G:68:TYR:CE1 | 3.00 | 0.43 |
| 1:A:71:ASN:ND2 | 1:A:102:MET:HG3 | 2.31 | 0.43 |
| 1:C:58:VAL:HB | 1:C:59:PRO:HD3 | 1.99 | 0.43 |
| 1:C:64:PHE:CZ | 1:C:68:TYR:CE1 | 3.00 | 0.43 |
| 1:C:76:VAL:O | 1:C:80:GLU:HG2 | 2.18 | 0.43 |
| 1:D:77:LEU:HD21 | 1:D:98:MET:HG3 | 2.01 | 0.43 |
| 1:G:77:LEU:HD21 | 1:G:98:MET:HG3 | 2.01 | 0.43 |
| 1:C:77:LEU:HD21 | 1:C:98:MET:HG3 | 2.01 | 0.43 |
| 1:B:57:ILE:O | 1:B:60:PRO:HG2 | 2.19 | 0.42 |
| 1:A:57:ILE:O | 1:A:60:PRO:HG2 | 2.20 | 0.42 |
| 1:B:77:LEU:HD21 | 1:B:98:MET:HG3 | 2.01 | 0.42 |
| 1:A:77:LEU:HD21 | 1:A:98:MET:HG3 | 2.01 | 0.42 |
| 1:E:57:ILE:O | 1:E:60:PRO:HG2 | 2.19 | 0.42 |
| 1:G:57:ILE:O | 1:G:60:PRO:HG2 | 2.20 | 0.42 |
| 1:F:57:ILE:O | 1:F:60:PRO:HG2 | 2.19 | 0.42 |
| 1:D:57:ILE:O | 1:D:60:PRO:HG2 | 2.20 | 0.42 |
| 1:A:62:TRP:NE1 | 1:G:194:THR:OG1 | 2.50 | 0.42 |
| 1:B:71:ASN:ND2 | 1:B:102:MET:HG3 | 2.31 | 0.42 |
| 1:C:78:THR:O | 1:C:82:LYS:HB2 | 2.20 | 0.42 |
| 1:D:71:ASN:ND2 | 1:D:102:MET:HG3 | 2.31 | 0.42 |
| 1:B:78:THR:O | 1:B:82:LYS:HB2 | 2.20 | 0.41 |
| 1:D:78:THR:O | 1:D:82:LYS:HB2 | 2.20 | 0.41 |
| 1:E:78:THR:O | 1:E:82:LYS:HB2 | 2.20 | 0.41 |
| 1:F:78:THR:O | 1:F:82:LYS:HB2 | 2.20 | 0.41 |
| 1:C:71:ASN:ND2 | 1:C:102:MET:HG3 | 2.31 | 0.41 |
| 1:G:57:ILE:HA | 1:G:57:ILE:HD13 | 1.94 | 0.41 |
| 1:C:57:ILE:O | 1:C:60:PRO:HG2 | 2.19 | 0.41 |
| 1:G:139:ASN:OD1 | 1:G:140:GLU:N | 2.54 | 0.41 |
| 1:G:78:THR:O | 1:G:82:LYS:HB2 | 2.20 | 0.41 |
| 1:A:78:THR:O | 1:A:82:LYS:HB2 | 2.20 | 0.41 |
| 1:F:139:ASN:OD1 | 1:F:140:GLU:N | 2.54 | 0.41 |
| 1:C:128:PHE:O | 1:C:132:ALA:HB2 | 2.22 | 0.40 |
| 1:C:65:LEU:HD23 | 1:C:65:LEU:HA | 1.95 | 0.40 |
| 1:A:139:ASN:OD1 | 1:A:140:GLU:N | 2.54 | 0.40 |
| 1:F:128:PHE:O | 1:F:132:ALA:HB2 | 2.22 | 0.40 |
| 1:A:128:PHE:O | 1:A:132:ALA:HB2 | 2.22 | 0.40 |
| 1:B:128:PHE:O | 1:B:132:ALA:HB2 | 2.22 | 0.40 |
| 1:D:64:PHE:CZ | 1:D:68:TYR:CE1 | 3.00 | 0.40 |
| 1:F:64:PHE:CZ | 1:F:68:TYR:CE1 | 3.00 | 0.40 |

There are no symmetry-related clashes.

### 5.3 Torsion angles

#### 5.3.1 Protein backbone

In the following table, the Percentiles column shows the percent Ramachandran outliers of the chain as a percentile score with respect to all PDB entries followed by that with respect to all EM entries.

The Analysed column shows the number of residues for which the backbone conformation was analysed, and the total number of residues.

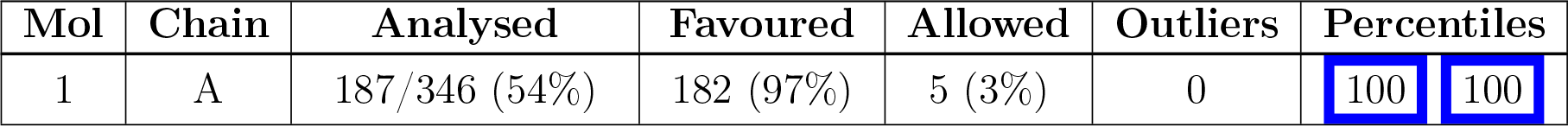
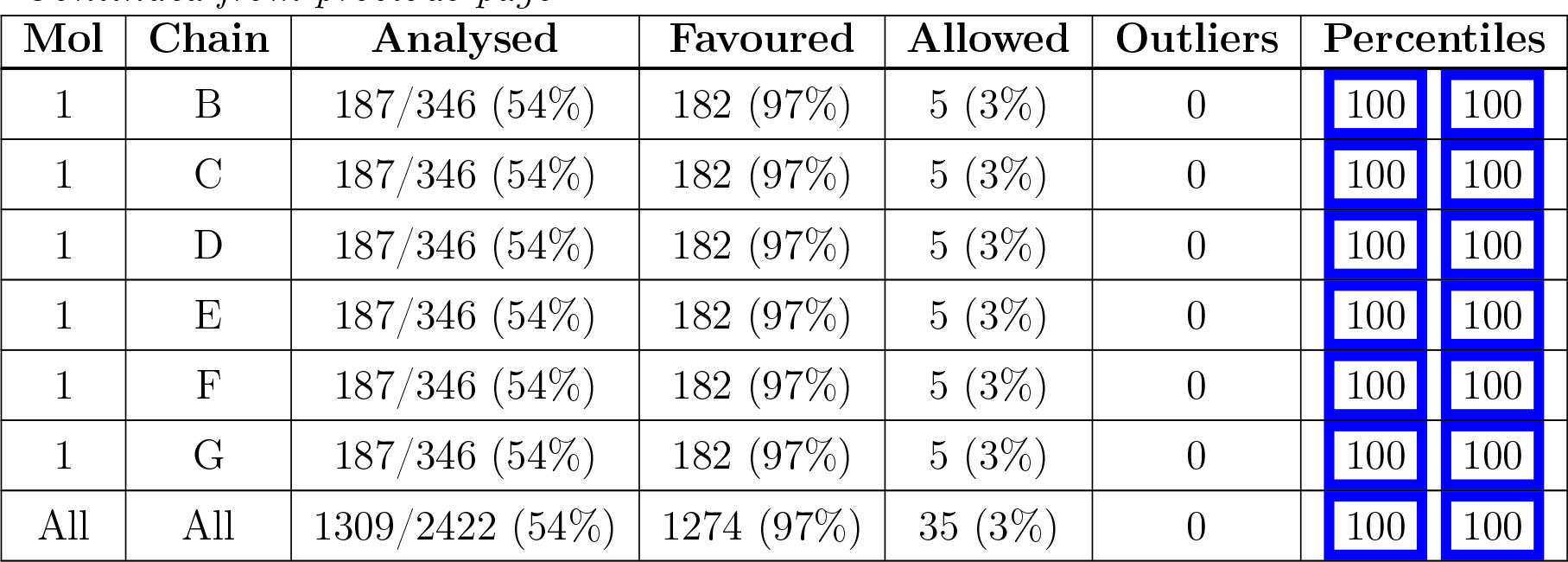

| Mol | Chain | Analysed | Favoured | Allowed | Outliers | Percentiles |  |
| --- | --- | --- | --- | --- | --- | --- | --- |
| 1 | A | 187/346 (54%) | 182 (97%) | 5 (3%) | 0 | 100 | 100 |
| 1 | B | 187/346 (54%) | 182 (97%) | 5 (3%) | 0 | 100 | 100 |
| 1 | C | 187/346 (54%) | 182 (97%) | 5 (3%) | 0 | 100 | 100 |
| 1 | D | 187/346 (54%) | 182 (97%) | 5 (3%) | 0 | 100 | 100 |
| 1 | E | 187/346 (54%) | 182 (97%) | 5 (3%) | 0 | 100 | 100 |
| 1 | F | 187/346 (54%) | 182 (97%) | 5 (3%) | 0 | 100 | 100 |
| 1 | G | 187/346 (54%) | 182 (97%) | 5 (3%) | 0 | 100 | 100 |
| All | All | 1309/2422 (54%) | 1274 (97%) | 35 (3%) | 0 | 100 | 100 |

There are no Ramachandran outliers to report.

#### 5.3.2 Protein sidechains

In the following table, the Percentiles column shows the percent sidechain outliers of the chain as a percentile score with respect to all PDB entries followed by that with respect to all EM entries.

The Analysed column shows the number of residues for which the sidechain conformation was analysed, and the total number of residues.

| Mol | Chain | Analysed | Rotameric | Outliers | Percentiles |  |
| --- | --- | --- | --- | --- | --- | --- |
| 1 | A | 155/306 (51%) | 153 (99%) | 2 (1%) | 69 | 90 |
| 1 | B | 155/306 (51%) | 153 (99%) | 2 (1%) | 69 | 90 |
| 1 | C | 155/306 (51%) | 153 (99%) | 2 (1%) | 69 | 90 |
| 1 | D | 155/306 (51%) | 153 (99%) | 2 (1%) | 69 | 90 |
| 1 | E | 155/306 (51%) | 153 (99%) | 2 (1%) | 69 | 90 |
| 1 | F | 155/306 (51%) | 153 (99%) | 2 (1%) | 69 | 90 |
| 1 | G | 155/306 (51%) | 153 (99%) | 2 (1%) | 69 | 90 |
| All | All | 1085/2142 (51%) | 1071 (99%) | 14 (1%) | 70 | 90 |

All (14) residues with a non-rotameric sidechain are listed below:

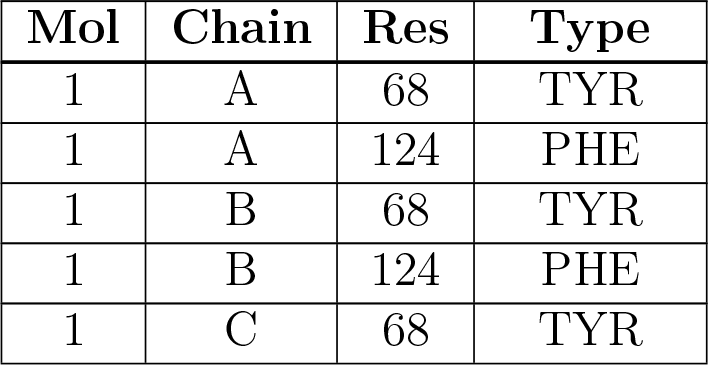
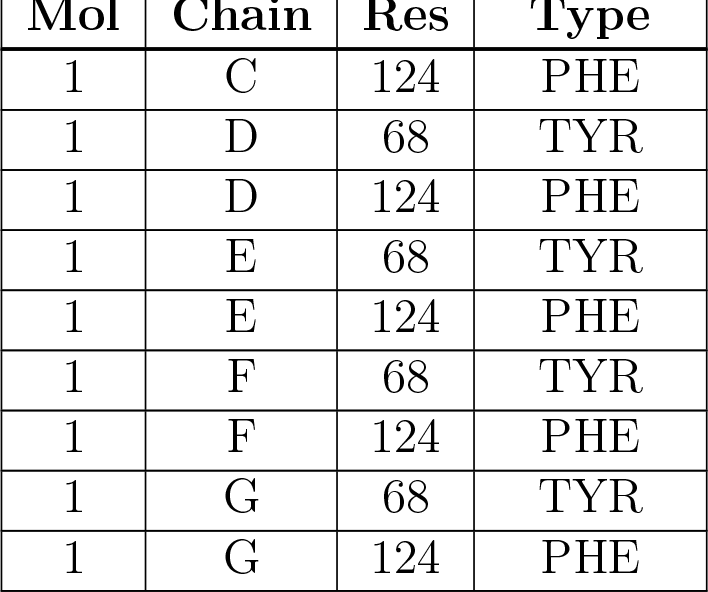

| Mol | Chain | Res | Type |
| --- | --- | --- | --- |
| 1 | A | 68 | TYR |
| 1 | A | 124 | PHE |
| 1 | B | 68 | TYR |
| 1 | B | 124 | PHE |
| 1 | C | 68 | TYR |
| 1 | C | 124 | PHE |
| 1 | D | 68 | TYR |
| 1 | D | 124 | PHE |
| 1 | E | 68 | TYR |
| 1 | E | 124 | PHE |
| 1 | F | 68 | TYR |
| 1 | F | 124 | PHE |
| 1 | G | 68 | TYR |
| 1 | G | 124 | PHE |

Sometimes sidechains can be flipped to improve hydrogen bonding and reduce clashes. All (14) such sidechains are listed below:

| Mol | Chain | Res | Type |
| --- | --- | --- | --- |
| 1 | A | 71 | ASN |
| 1 | A | 154 | GLN |
| 1 | B | 71 | ASN |
| 1 | B | 154 | GLN |
| 1 | C | 71 | ASN |
| 1 | C | 154 | GLN |
| 1 | D | 71 | ASN |
| 1 | D | 154 | GLN |
| 1 | E | 71 | ASN |
| 1 | E | 154 | GLN |
| 1 | F | 71 | ASN |
| 1 | F | 154 | GLN |
| 1 | G | 71 | ASN |
| 1 | G | 154 | GLN |

#### 5.3.3 RNA

There are no RNA molecules in this entry.

### 5.4 Non-standard residues in protein, DNA, RNA chains

There are no non-standard protein/DNA/RNA residues in this entry.

### 5.5 Carbohydrates

There are no monosaccharides in this entry.

### 5.6 Ligand geometry

7 ligands are modelled in this entry.

In the following table, the Counts columns list the number of bonds (or angles) for which Mogul statistics could be retrieved, the number of bonds (or angles) that are observed in the model and the number of bonds (or angles) that are defined in the Chemical Component Dictionary. The Link column lists molecule types, if any, to which the group is linked. The Z score for a bond length (or angle) is the number of standard deviations the observed value is removed from the expected value. A bond length (or angle) with *|Z| >* 2 is considered an outlier worth inspection. RMSZ is the root-mean-square of all Z scores of the bond lengths (or angles).

| Mol | Type | Chain | Res | Link | Bond lengths |  |  | Bond angles |  |  |
| --- | --- | --- | --- | --- | --- | --- | --- | --- | --- | --- |
|  |  |  |  |  | Counts | RMSZ | #\|Z\| > 2 | Counts | RMSZ | #\|Z\| > 2 |
| 2 | NAG | A | 1001 | 1 | 14,14,15 | 0.27 | 0 | 17,19,21 | 0.50 | 0 |
| 2 | NAG | G | 1001 | 1 | 14,14,15 | 0.27 | 0 | 17,19,21 | 0.50 | 0 |
| 2 | NAG | C | 1001 | 1 | 14,14,15 | 0.28 | 0 | 17,19,21 | 0.50 | 0 |
| 2 | NAG | E | 1001 | 1 | 14,14,15 | 0.28 | 0 | 17,19,21 | 0.51 | 0 |
| 2 | NAG | F | 1001 | 1 | 14,14,15 | 0.29 | 0 | 17,19,21 | 0.51 | 0 |
| 2 | NAG | D | 1001 | 1 | 14,14,15 | 0.27 | 0 | 17,19,21 | 0.52 | 0 |
| 2 | NAG | B | 1001 | 1 | 14,14,15 | 0.28 | 0 | 17,19,21 | 0.51 | 0 |

In the following table, the Chirals column lists the number of chiral outliers, the number of chiral centers analysed, the number of these observed in the model and the number defined in the Chemical Component Dictionary. Similar counts are reported in the Torsion and Rings columns. ’-’ means no outliers of that kind were identified.

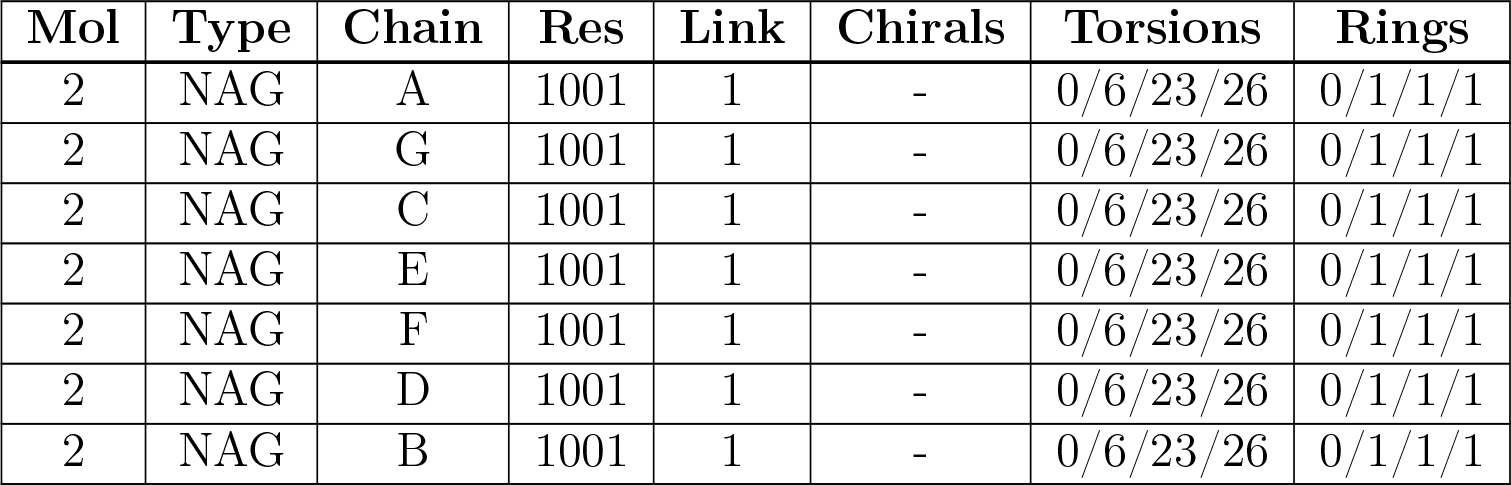

There are no bond length outliers.

There are no bond angle outliers.

There are no chirality outliers.

There are no torsion outliers.

There are no ring outliers.

No monomer is involved in short contacts.

### 5.7 Other polymers

There are no such residues in this entry.

### 5.8 Polymer linkage issues

There are no chain breaks in this entry.

## 6 Map visualisation

This section contains visualisations of the EMDB entry EMD-30831. These allow visual inspection of the internal detail of the map and identification of artifacts.

No raw map or half-maps were deposited for this entry and therefore no images, graphs, etc. pertaining to the raw map can be shown.

### 6.1 Orthogonal projections

#### 6.1.1 Primary map

The images above show the map projected in three orthogonal directions.

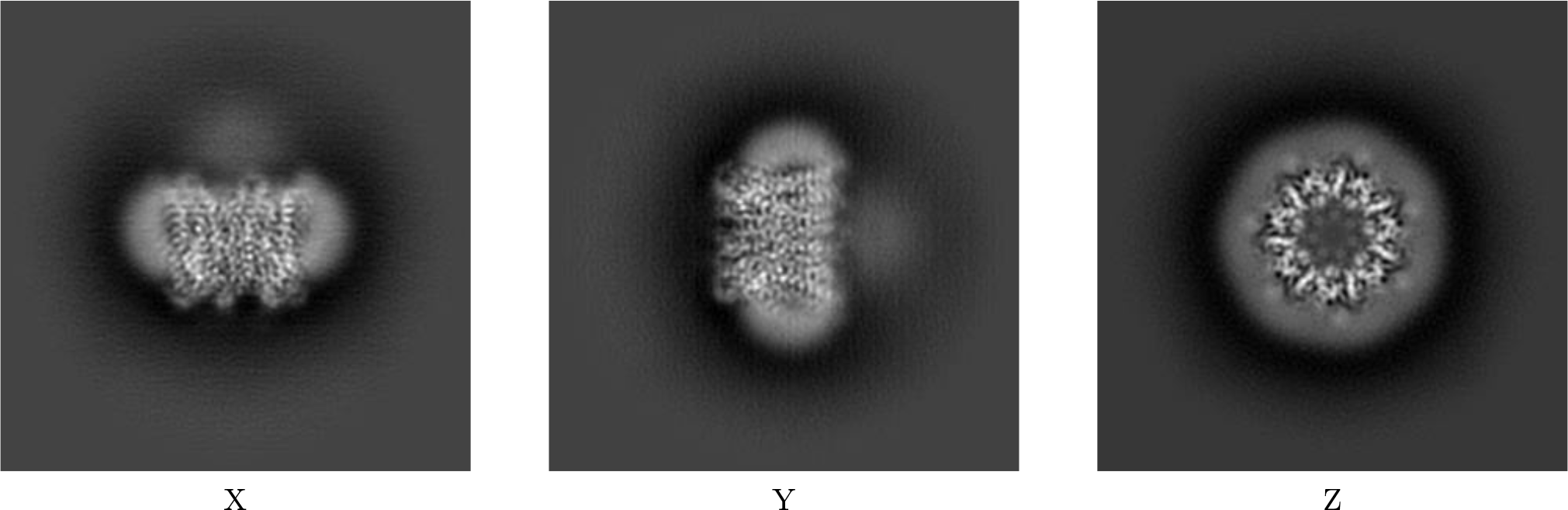

### 6.2 Central slices

#### 6.2.1 Primary map

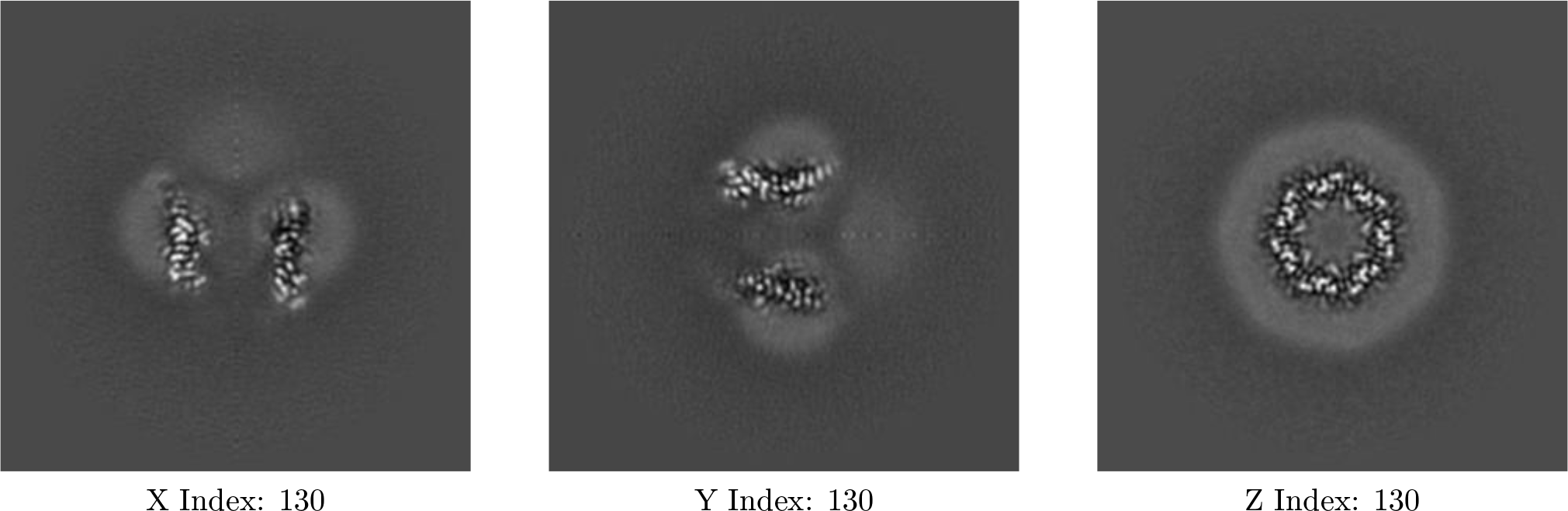

The images above show central slices of the map in three orthogonal directions.

### 6.3 Largest variance slices

#### 6.3.1 Primary map

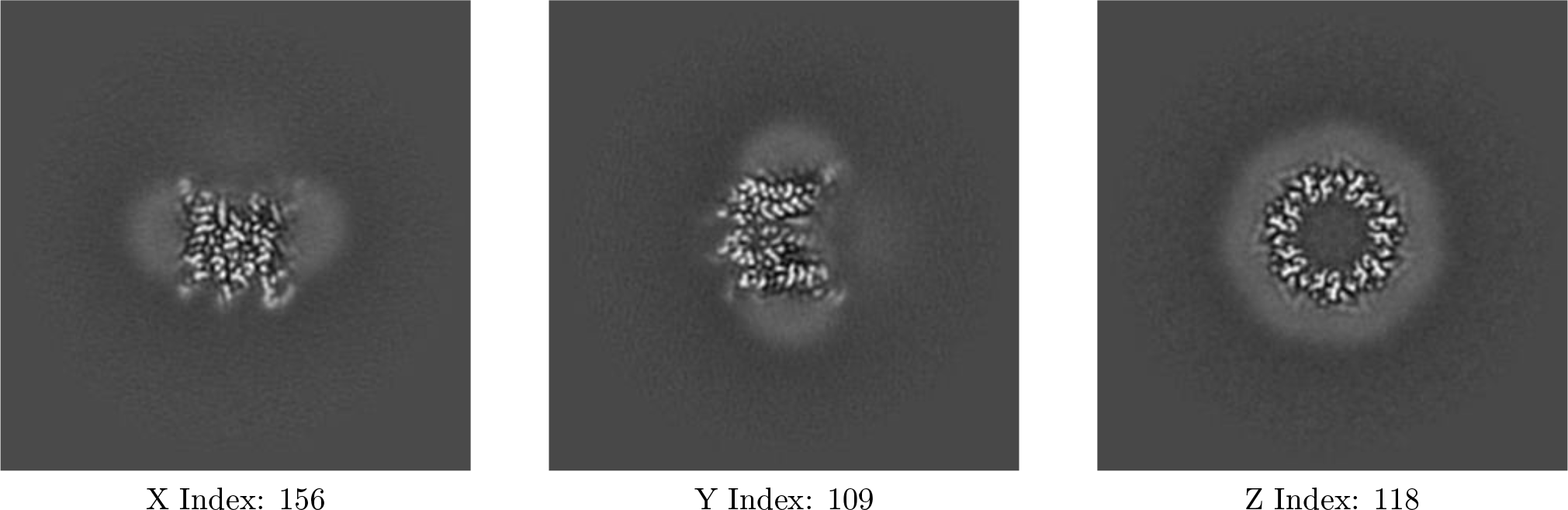

The images above show the largest variance slices of the map in three orthogonal directions.

### 6.4 Orthogonal surface views

#### 6.4.1 Primary map

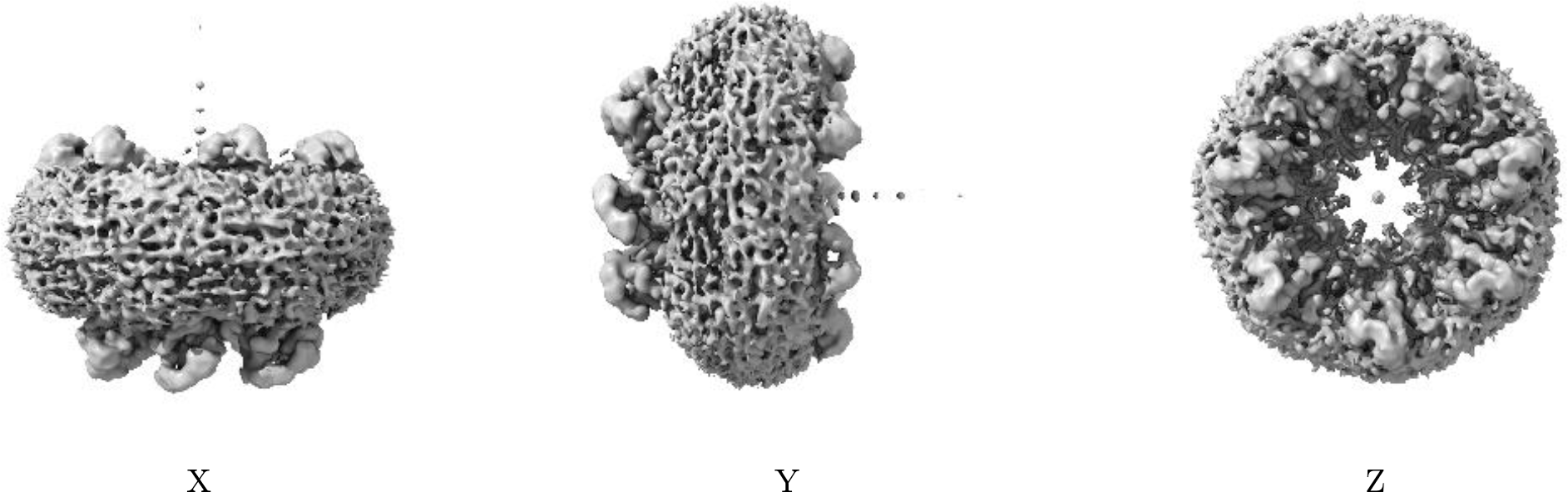

The images above show the 3D surface view of the map at the recommended contour level 0.154. These images, in conjunction with the slice images, may facilitate assessment of whether an appropriate contour level has been provided.

### 6.5 Mask visualisation

This section was not generated. No masks/segmentation were deposited.

## 7 Map analysis

This section contains the results of statistical analysis of the map.

### 7.1 Map-value distribution

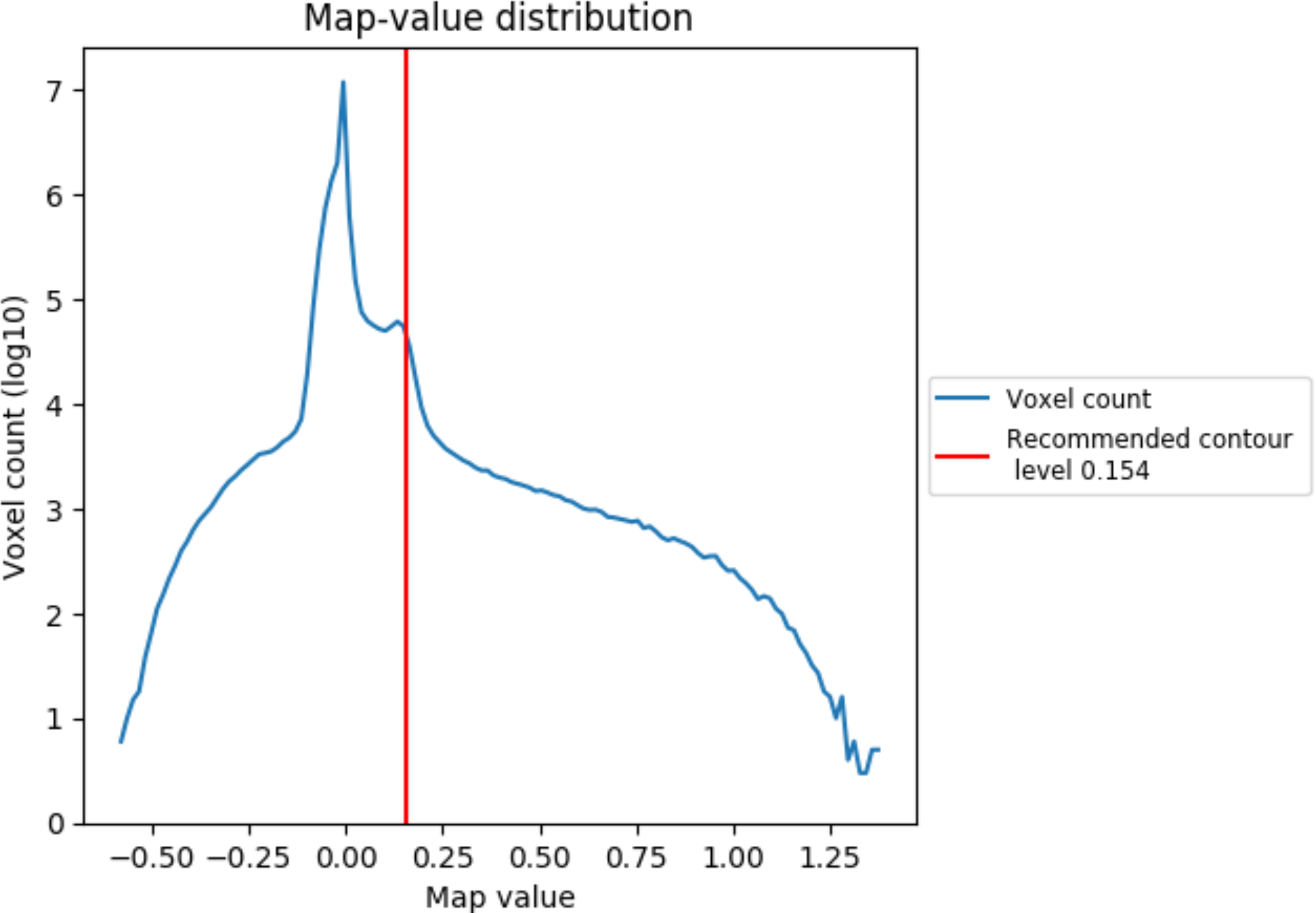

The map-value distribution is plotted in 128 intervals along the x-axis. The y-axis is logarithmic. A spike in this graph at zero usually indicates that the volume has been masked.

### 7.2 Volume estimate

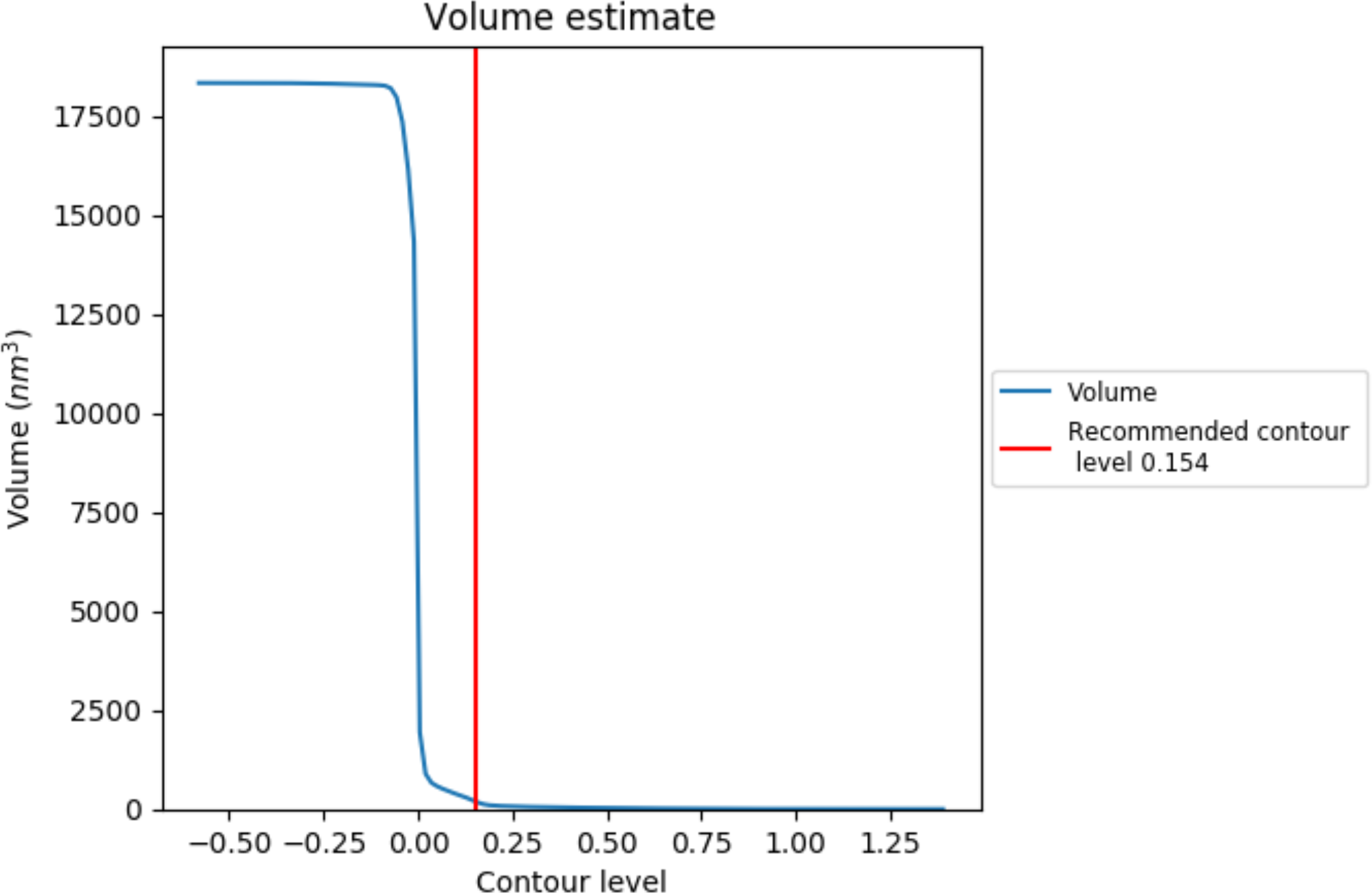

The volume at the recommended contour level is 185 nm^3^; this corresponds to an approximate mass of 167 kDa.

The volume estimate graph shows how the enclosed volume varies with the contour level. The recommended contour level is shown as a vertical line and the intersection between the line and the curve gives the volume of the enclosed surface at the given level.

### 7.3 Rotationally averaged power spectrum

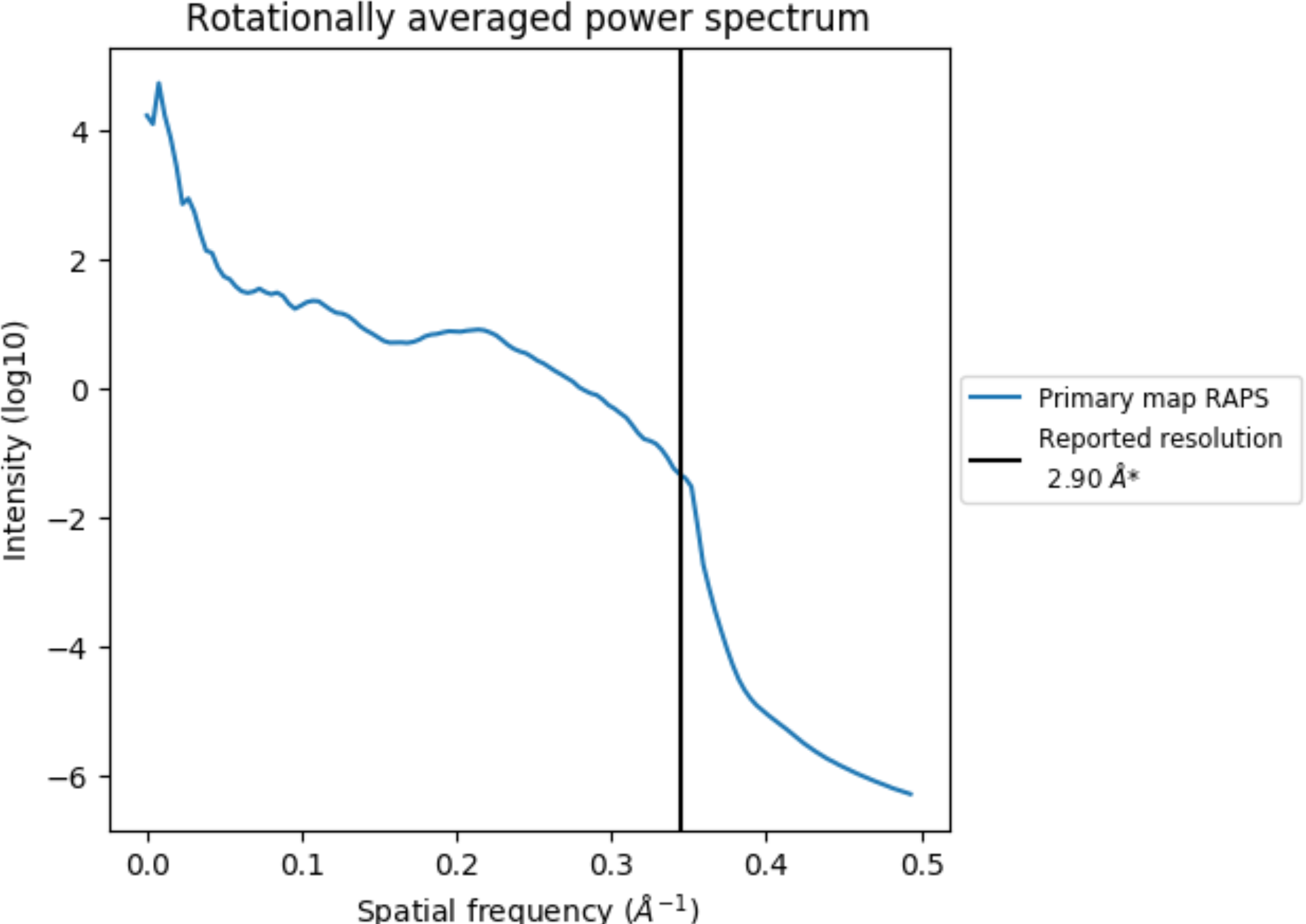

## 8 Fourier-Shell correlation

This section was not generated. No FSC curve or half-maps provided.

## 9 Map-model fit

This section contains information regarding the fit between EMDB map EMD-30831 and PDB model 7DSD. Per-residue inclusion information can be found in section 3 on page 5.

### 9.1 Map-model overlay

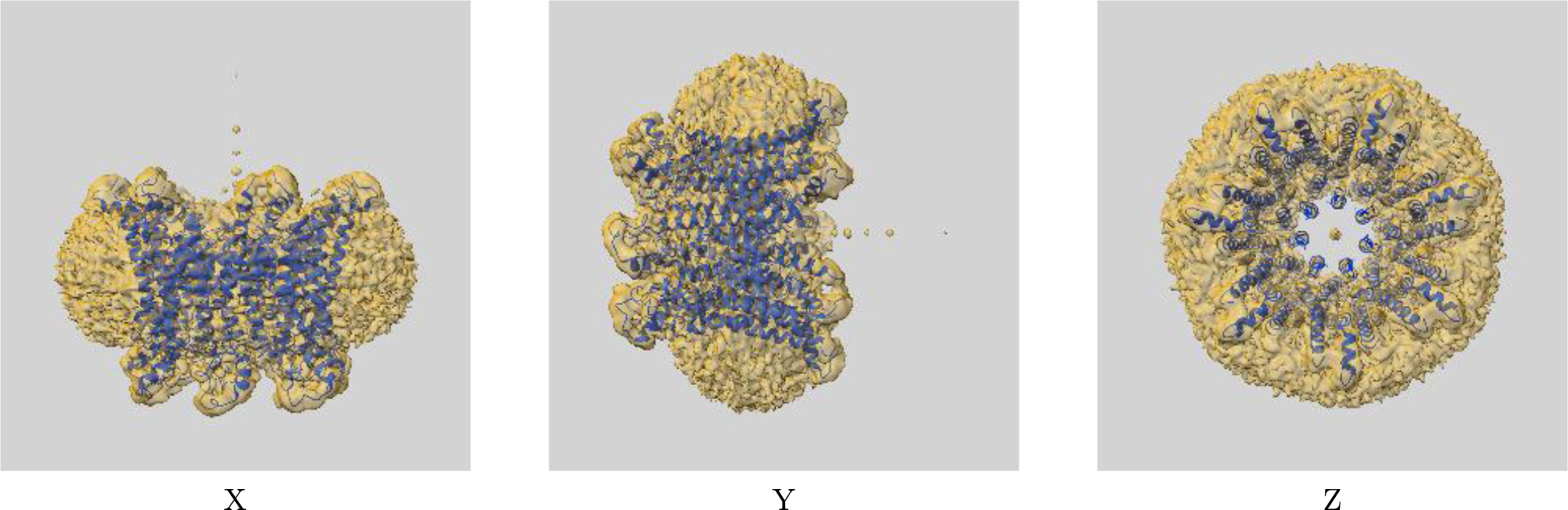

The images above show the 3D surface view of the map at the recommended contour level 0.154 at 50% transparency in yellow overlaid with a ribbon representation of the model coloured in blue. These images allow for the visual assessment of the quality of fit between the atomic model and the map.

### 9.2 Atom inclusion

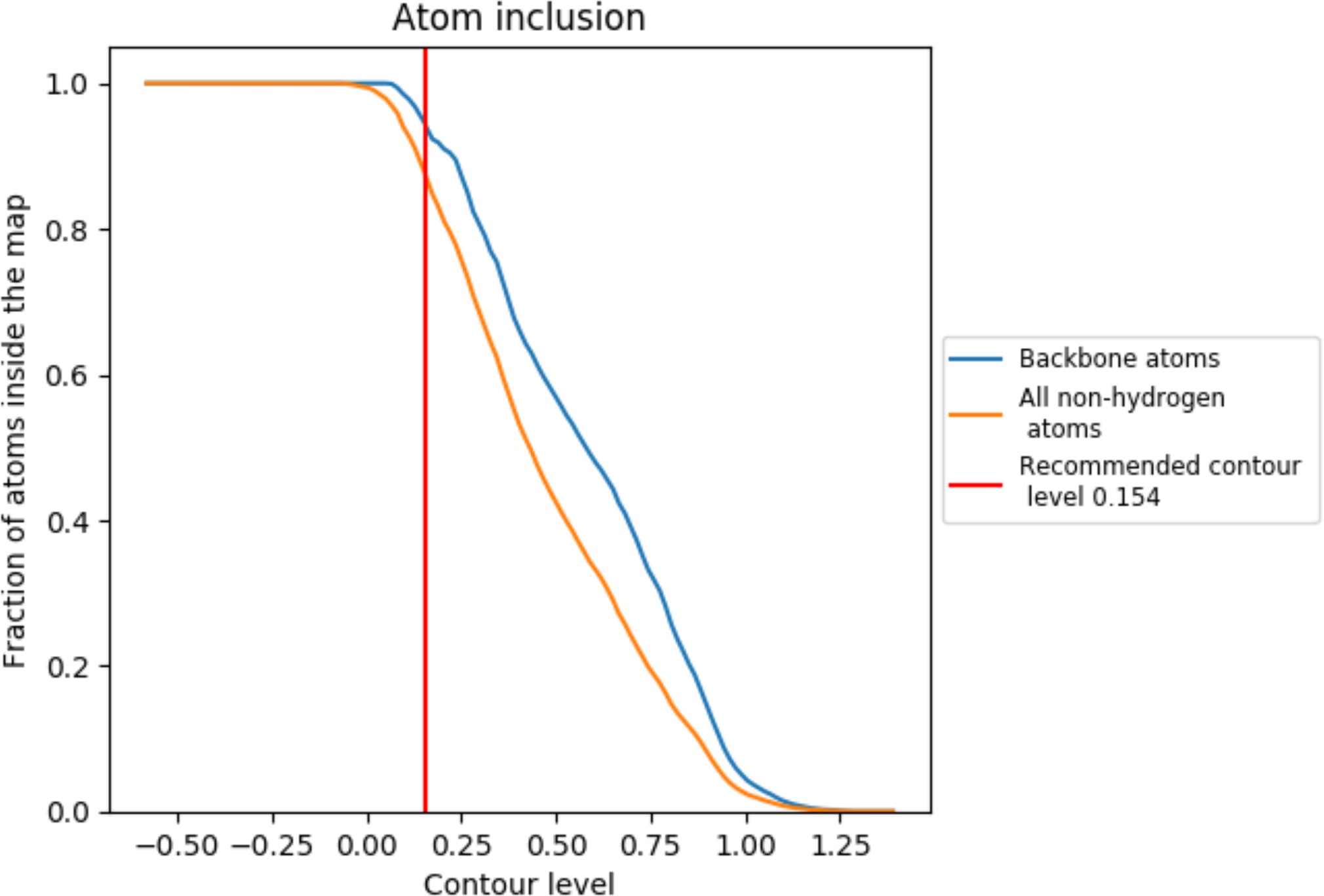

At the recommended contour level, 95% of all backbone atoms, 88% of all non-hydrogen atoms, are inside the map.

## 1 Overall quality at a glance

The following experimental techniques were used to determine the structure: *ELECTRON MICROSCOPY*

The reported resolution of this entry is 3.50 Å.

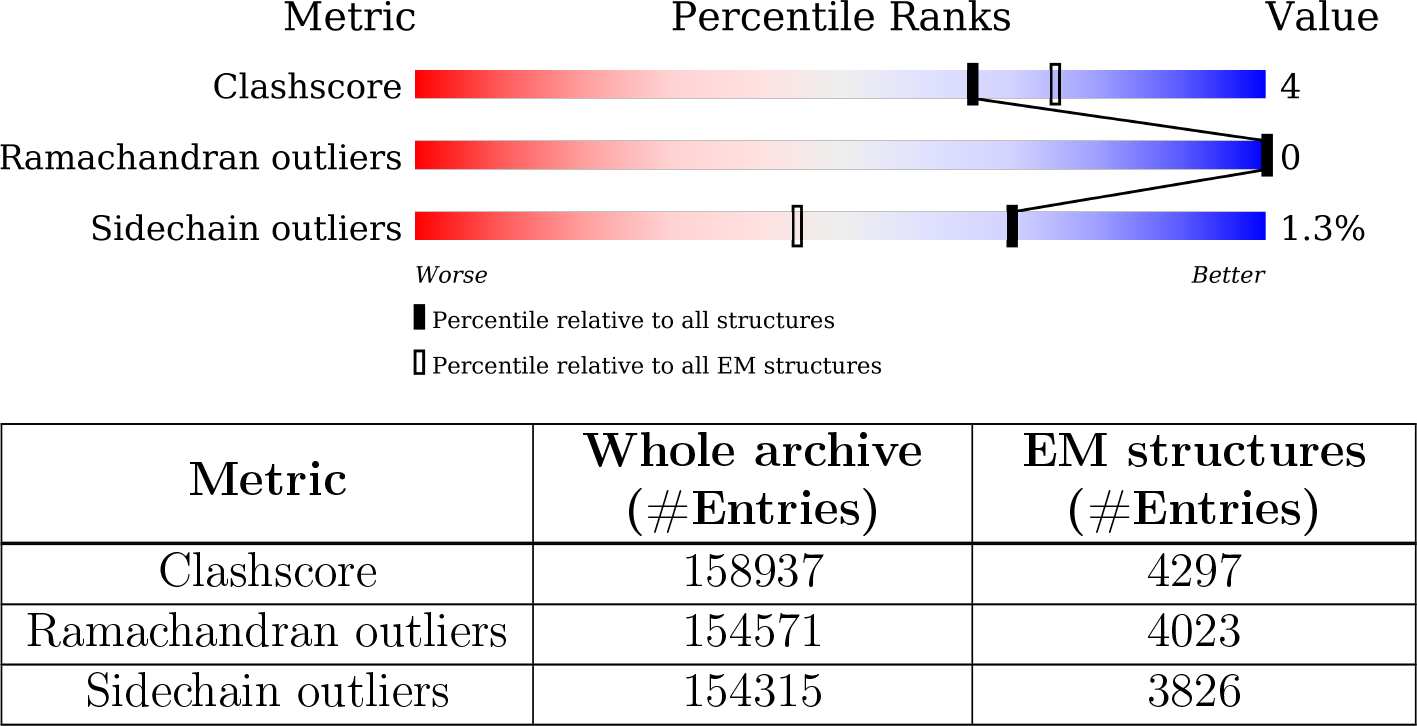

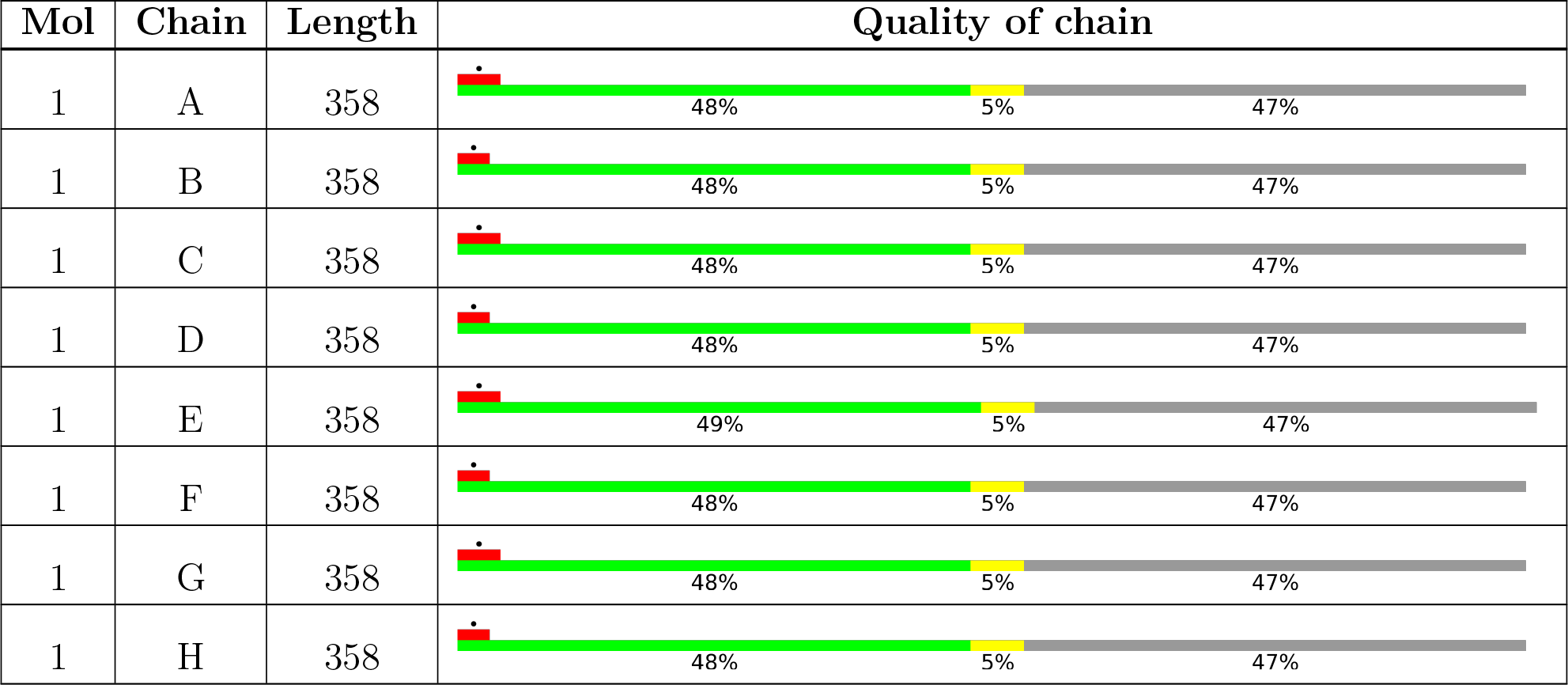

## 2 Entry composition

There are 2 unique types of molecules in this entry. The entry contains 11848 atoms, of which 0 are hydrogens and 0 are deuteriums.

• Molecule 1 is a protein called Calcium homeostasis modulator 1.

| Mol | Chain | Residues | Atoms |  |  |  |  | AltConf | Trace |
| --- | --- | --- | --- | --- | --- | --- | --- | --- | --- |
| 1 | A | 191 | Total | C | N | O | S | 0 | 0 |
|  |  |  | 1467 | 959 | 235 | 260 | 13 |  |  |
| 1 | B | 191 | Total | C | N | O | S | 0 | 0 |
|  |  |  | 1467 | 959 | 235 | 260 | 13 |  |  |
| 1 | C | 191 | Total | C | N | O | S | 0 | 0 |
|  |  |  | 1467 | 959 | 235 | 260 | 13 |  |  |
| 1 | D | 191 | Total | C | N | O | S | 0 | 0 |
|  |  |  | 1467 | 959 | 235 | 260 | 13 |  |  |
| 1 | E | 191 | Total | C | N | O | S | 0 | 0 |
|  |  |  | 1467 | 959 | 235 | 260 | 13 |  |  |
| 1 | F | 191 | Total | C | N | O | S | 0 | 0 |
|  |  |  | 1467 | 959 | 235 | 260 | 13 |  |  |
| 1 | G | 191 | Total | C | N | O | S | 0 | 0 |
|  |  |  | 1467 | 959 | 235 | 260 | 13 |  |  |
| 1 | H | 191 | Total | C | N | O | S | 0 | 0 |
|  |  |  | 1467 | 959 | 235 | 260 | 13 |  |  |

There are 104 discrepancies between the modelled and reference sequences:

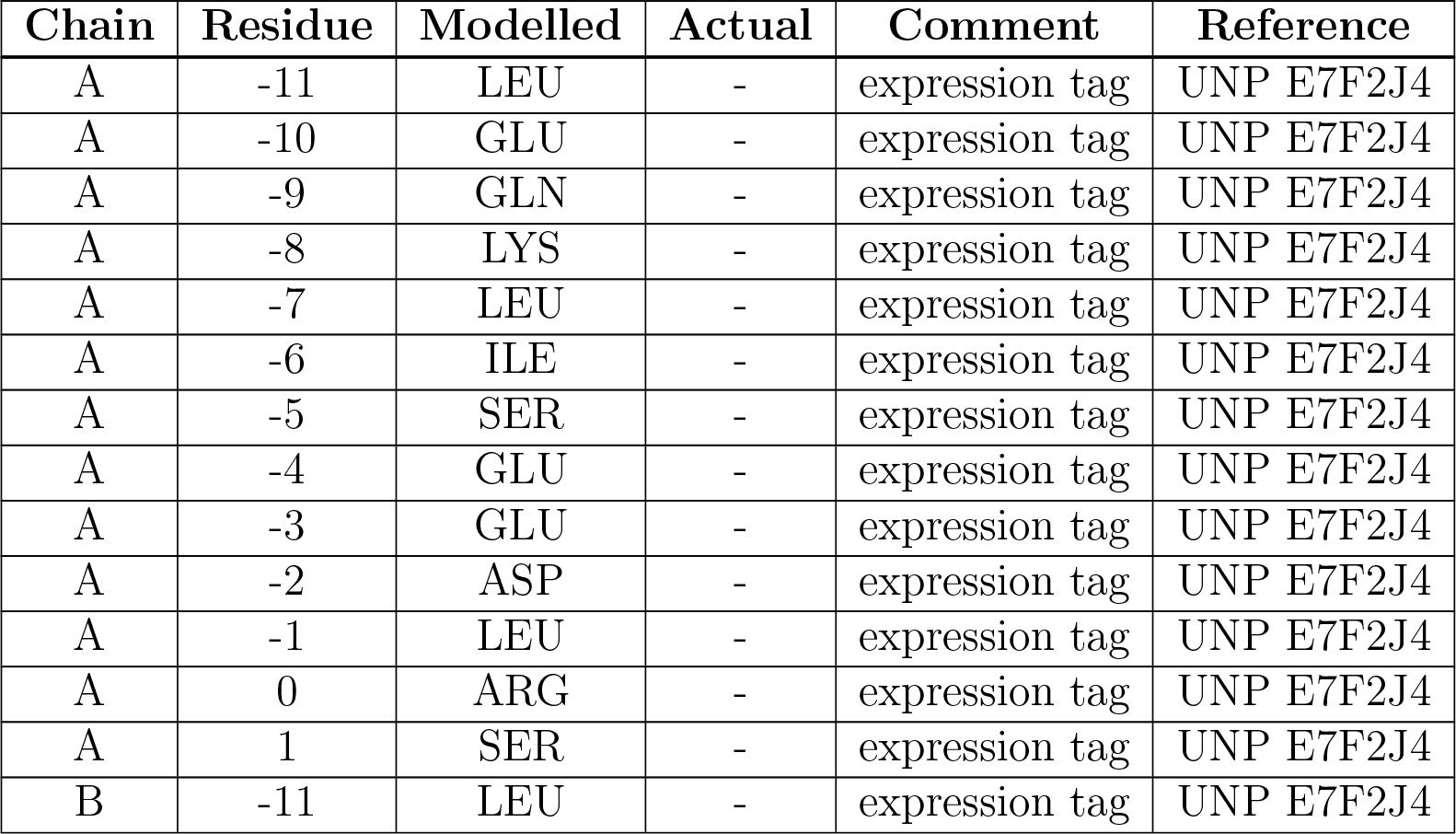
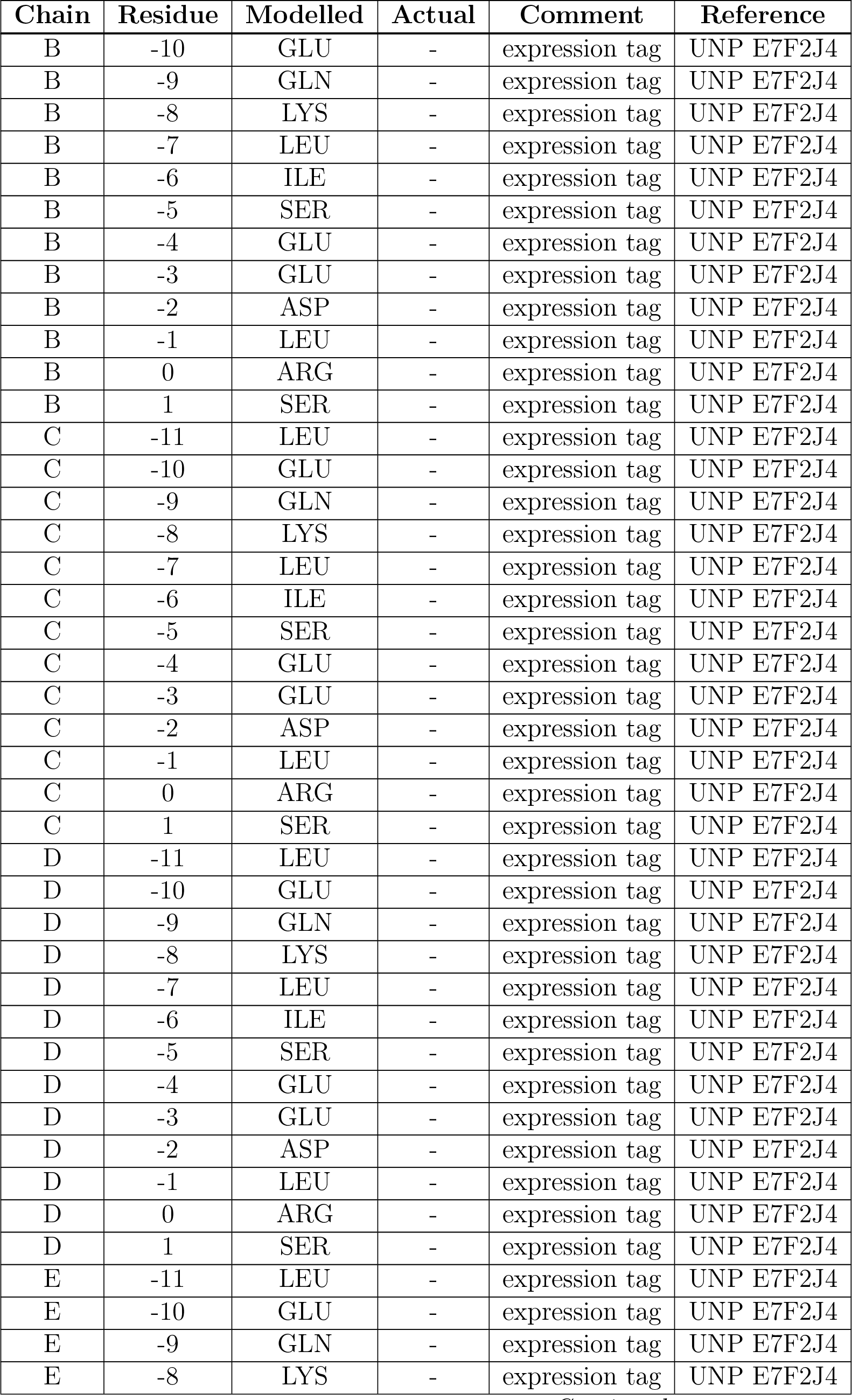
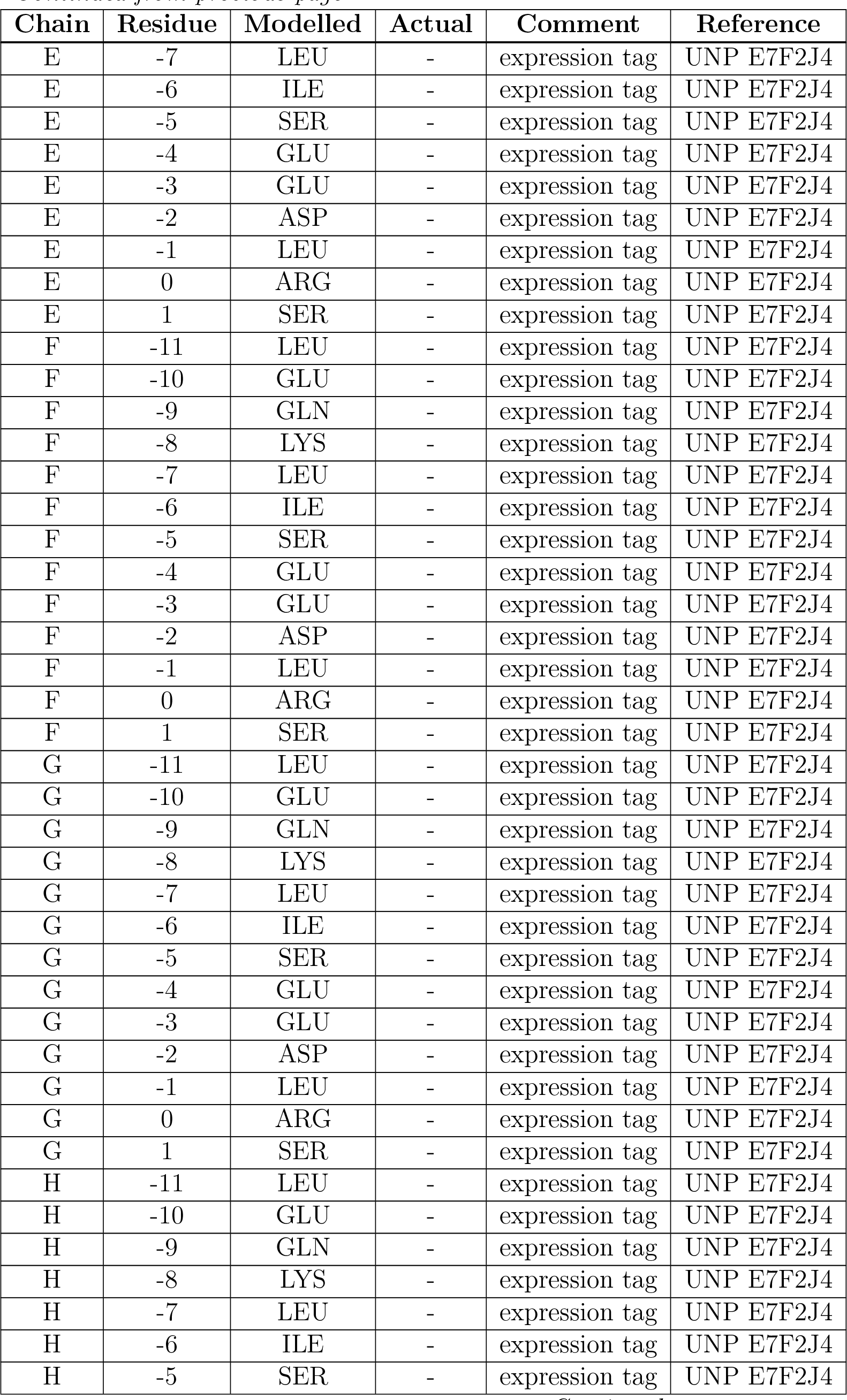
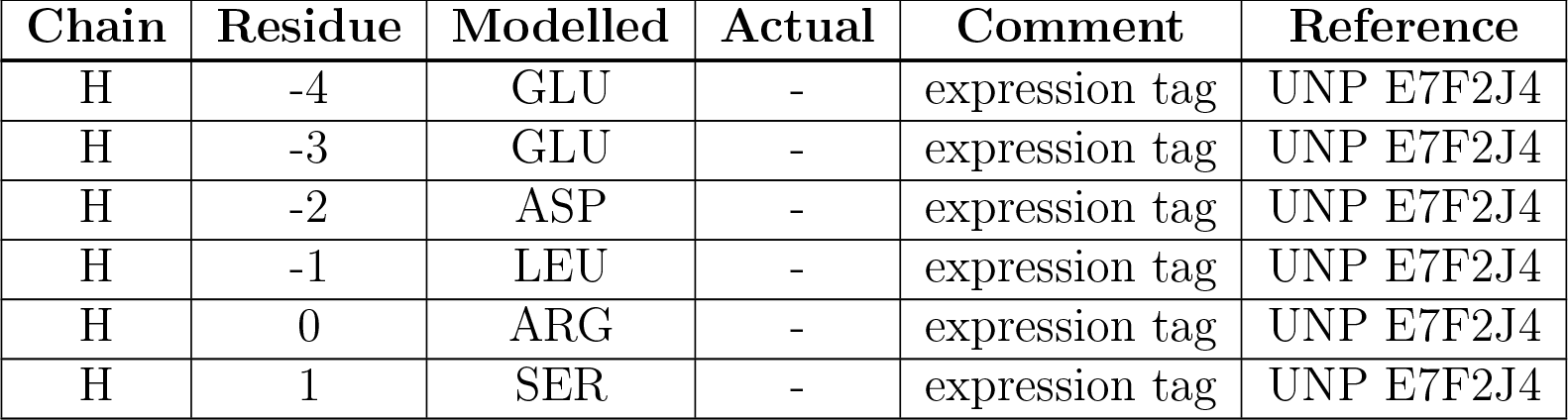

| Chain | Residue | Modelled | Actual | Comment | Reference |
| --- | --- | --- | --- | --- | --- |
| A | -11 | LEU | - | expression tag | UNP E7F2J4 |
| A | -10 | GLU | - | expression tag | UNP E7F2J4 |
| A | -9 | GLN | - | expression tag | UNP E7F2J4 |
| A | -8 | LYS | - | expression tag | UNP E7F2J4 |
| A | -7 | LEU | - | expression tag | UNP E7F2J4 |
| A | -6 | ILE | - | expression tag | UNP E7F2J4 |
| A | -5 | SER | - | expression tag | UNP E7F2J4 |
| A | -4 | GLU | - | expression tag | UNP E7F2J4 |
| A | -3 | GLU | - | expression tag | UNP E7F2J4 |
| A | -2 | ASP | - | expression tag | UNP E7F2J4 |
| A | -1 | LEU | - | expression tag | UNP E7F2J4 |
| A | 0 | ARG | - | expression tag | UNP E7F2J4 |
| A | 1 | SER | - | expression tag | UNP E7F2J4 |
| B | -11 | LEU | - | expression tag | UNP E7F2J4 |
| B | -10 | GLU | - | expression tag | UNP E7F2J4 |
| B | -9 | GLN | - | expression tag | UNP E7F2J4 |
| B | -8 | LYS | - | expression tag | UNP E7F2J4 |
| B | -7 | LEU | - | expression tag | UNP E7F2J4 |
| B | -6 | ILE | - | expression tag | UNP E7F2J4 |
| B | -5 | SER | - | expression tag | UNP E7F2J4 |
| B | -4 | GLU | - | expression tag | UNP E7F2J4 |
| B | -3 | GLU | - | expression tag | UNP E7F2J4 |
| B | -2 | ASP | - | expression tag | UNP E7F2J4 |
| B | -1 | LEU | - | expression tag | UNP E7F2J4 |
| B | 0 | ARG | - | expression tag | UNP E7F2J4 |
| B | 1 | SER | - | expression tag | UNP E7F2J4 |
| C | -11 | LEU | - | expression tag | UNP E7F2J4 |
| C | -10 | GLU | - | expression tag | UNP E7F2J4 |
| C | -9 | GLN | - | expression tag | UNP E7F2J4 |
| C | -8 | LYS | - | expression tag | UNP E7F2J4 |
| C | -7 | LEU | - | expression tag | UNP E7F2J4 |
| C | -6 | ILE | - | expression tag | UNP E7F2J4 |
| C | -5 | SER | - | expression tag | UNP E7F2J4 |
| C | -4 | GLU | - | expression tag | UNP E7F2J4 |
| C | -3 | GLU | - | expression tag | UNP E7F2J4 |
| C | -2 | ASP | - | expression tag | UNP E7F2J4 |
| C | -1 | LEU | - | expression tag | UNP E7F2J4 |
| C | 0 | ARG | - | expression tag | UNP E7F2J4 |
| C | 1 | SER | - | expression tag | UNP E7F2J4 |
| D | -11 | LEU | - | expression tag | UNP E7F2J4 |
| D | -10 | GLU | - | expression tag | UNP E7F2J4 |
| D | -9 | GLN | - | expression tag | UNP E7F2J4 |
| D | -8 | LYS | - | expression tag | UNP E7F2J4 |
| D | -7 | LEU | - | expression tag | UNP E7F2J4 |
| D | -6 | ILE | - | expression tag | UNP E7F2J4 |
| D | -5 | SER | - | expression tag | UNP E7F2J4 |
| D | -4 | GLU | - | expression tag | UNP E7F2J4 |
| D | -3 | GLU | - | expression tag | UNP E7F2J4 |
| D | -2 | ASP | - | expression tag | UNP E7F2J4 |
| D | -1 | LEU | - | expression tag | UNP E7F2J4 |
| D | 0 | ARG | - | expression tag | UNP E7F2J4 |
| D | 1 | SER | - | expression tag | UNP E7F2J4 |
| E | -11 | LEU | - | expression tag | UNP E7F2J4 |
| E | -10 | GLU | - | expression tag | UNP E7F2J4 |
| E | -9 | GLN | - | expression tag | UNP E7F2J4 |
| E | -8 | LYS | - | expression tag | UNP E7F2J4 |
| E | -7 | LEU | - | expression tag | UNP E7F2J4 |
| E | -6 | ILE | - | expression tag | UNP E7F2J4 |
| E | -5 | SER | - | expression tag | UNP E7F2J4 |
| E | -4 | GLU | - | expression tag | UNP E7F2J4 |
| E | -3 | GLU | - | expression tag | UNP E7F2J4 |
| E | -2 | ASP | - | expression tag | UNP E7F2J4 |
| E | -1 | LEU | - | expression tag | UNP E7F2J4 |
| E | 0 | ARG | - | expression tag | UNP E7F2J4 |
| E | 1 | SER | - | expression tag | UNP E7F2J4 |
| F | -11 | LEU | - | expression tag | UNP E7F2J4 |
| F | -10 | GLU | - | expression tag | UNP E7F2J4 |
| F | -9 | GLN | - | expression tag | UNP E7F2J4 |
| F | -8 | LYS | - | expression tag | UNP E7F2J4 |
| F | -7 | LEU | - | expression tag | UNP E7F2J4 |
| F | -6 | ILE | - | expression tag | UNP E7F2J4 |
| F | -5 | SER | - | expression tag | UNP E7F2J4 |
| F | -4 | GLU | - | expression tag | UNP E7F2J4 |
| F | -3 | GLU | - | expression tag | UNP E7F2J4 |
| F | -2 | ASP | - | expression tag | UNP E7F2J4 |
| F | -1 | LEU | - | expression tag | UNP E7F2J4 |
| F | 0 | ARG | - | expression tag | UNP E7F2J4 |
| F | 1 | SER | - | expression tag | UNP E7F2J4 |
| G | -11 | LEU | - | expression tag | UNP E7F2J4 |
| G | -10 | GLU | - | expression tag | UNP E7F2J4 |
| G | -9 | GLN | - | expression tag | UNP E7F2J4 |
| G | -8 | LYS | - | expression tag | UNP E7F2J4 |
| G | -7 | LEU | - | expression tag | UNP E7F2J4 |
| G | -6 | ILE | - | expression tag | UNP E7F2J4 |
| G | -5 | SER | - | expression tag | UNP E7F2J4 |
| G | -4 | GLU | - | expression tag | UNP E7F2J4 |
| G | -3 | GLU | - | expression tag | UNP E7F2J4 |
| G | -2 | ASP | - | expression tag | UNP E7F2J4 |
| G | -1 | LEU | - | expression tag | UNP E7F2J4 |
| G | 0 | ARG | - | expression tag | UNP E7F2J4 |
| G | 1 | SER | - | expression tag | UNP E7F2J4 |
| H | -11 | LEU | - | expression tag | UNP E7F2J4 |
| H | -10 | GLU | - | expression tag | UNP E7F2J4 |
| H | -9 | GLN | - | expression tag | UNP E7F2J4 |
| H | -8 | LYS | - | expression tag | UNP E7F2J4 |
| H | -7 | LEU | - | expression tag | UNP E7F2J4 |
| H | -6 | ILE | - | expression tag | UNP E7F2J4 |
| H | -5 | SER | - | expression tag | UNP E7F2J4 |
| H | -4 | GLU | - | expression tag | UNP E7F2J4 |
| H | -3 | GLU | - | expression tag | UNP E7F2J4 |
| H | -2 | ASP | - | expression tag | UNP E7F2J4 |
| H | -1 | LEU | - | expression tag | UNP E7F2J4 |
| H | 0 | ARG | - | expression tag | UNP E7F2J4 |
| H | 1 | SER | - | expression tag | UNP E7F2J4 |

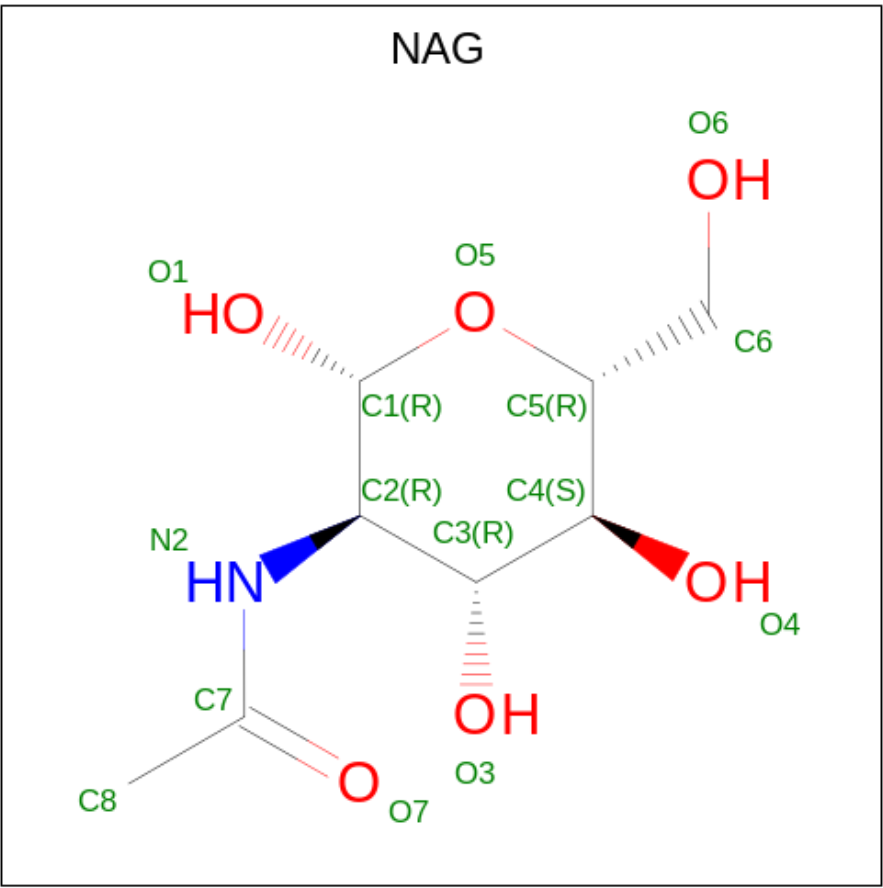

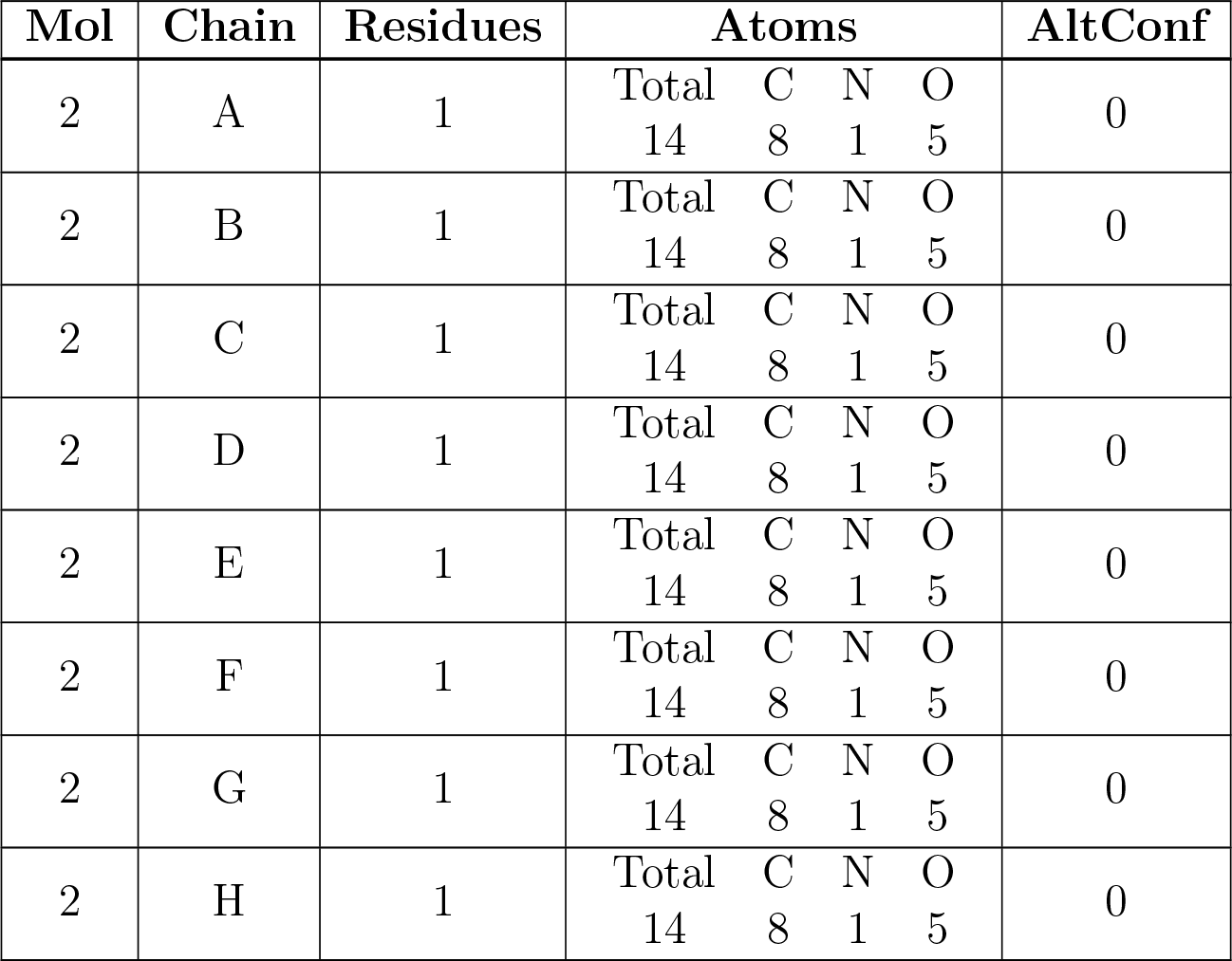

## 3 Residue-property plots

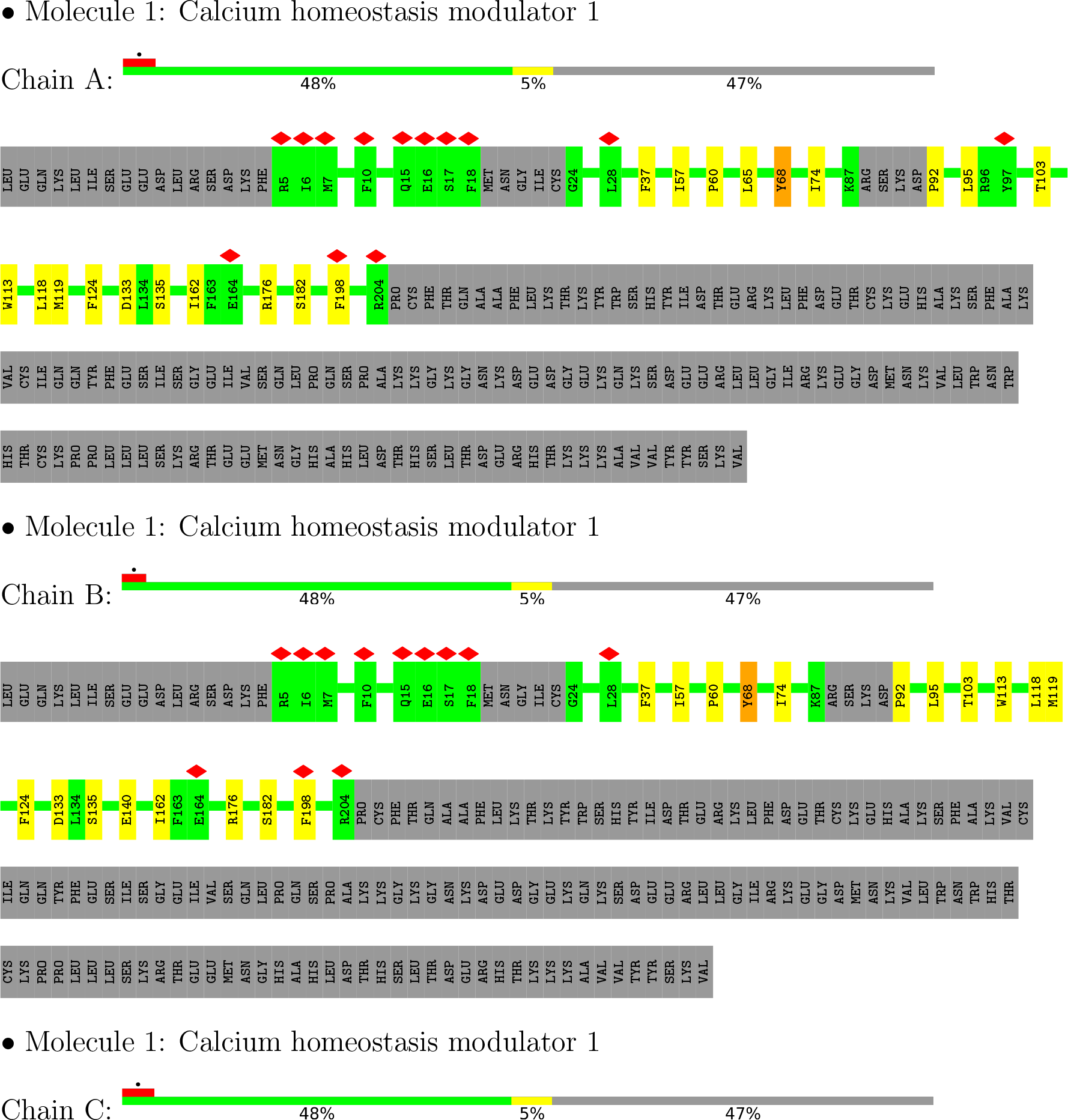
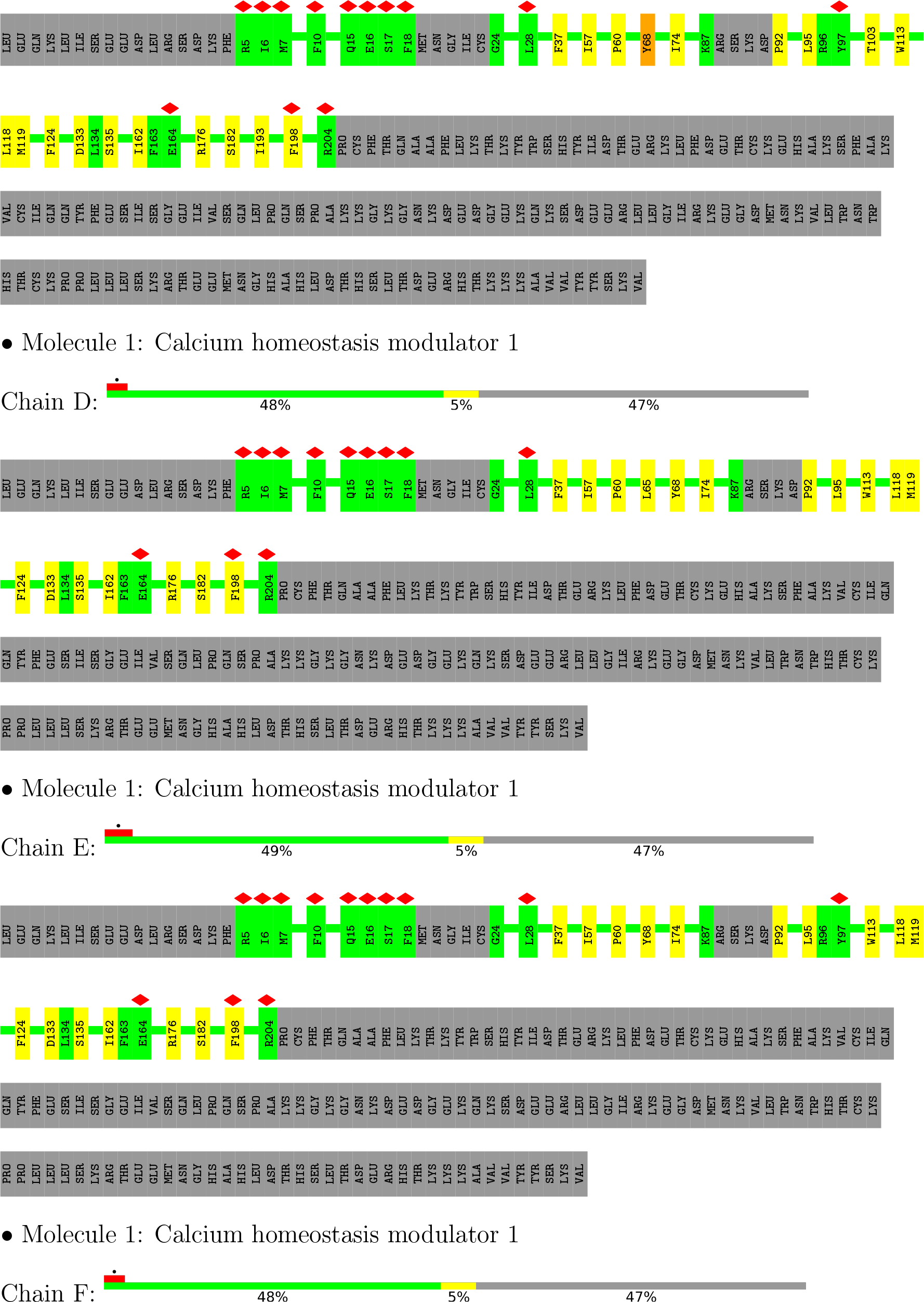
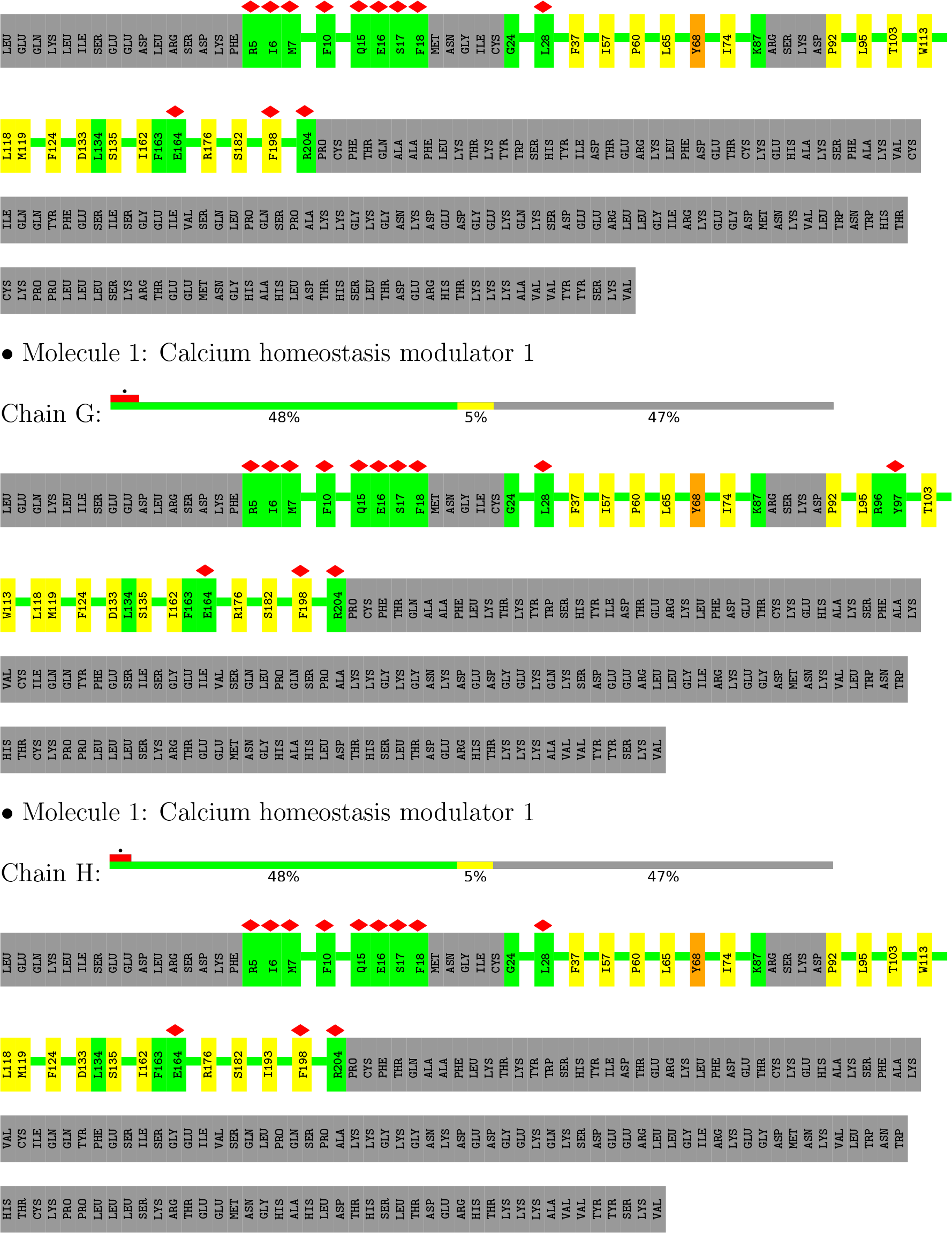

## 4 Experimental information

| Property | Value | Source |
| --- | --- | --- |
| EM reconstruction method | SINGLE PARTICLE | Depositor |
| Imposed symmetry | POINT, Not provided |  |
| Number of particles used | 44894 | Depositor |
| Resolution determination method | FSC 0.143 CUT-OFF | Depositor |
| CTF correction method | NONE | Depositor |
| Microscope | FEI TITAN KRIOS | Depositor |
| Voltage (kV) | 300 | Depositor |
| Electron dose ($e^-/\text{\AA}^2$) | 50 | Depositor |
| Minimum defocus (nm) | Not provided |  |
| Maximum defocus (nm) | Not provided |  |
| Magnification | Not provided |  |
| Image detector | GATAN K2 SUMMIT (4k x 4k) | Depositor |
| Maximum map value | 0.069 | Depositor |
| Minimum map value | -0.040 | Depositor |
| Average map value | 0.000 | Depositor |
| Map value standard deviation | 0.002 | Depositor |
| Recommended contour level | 0.0075 | Depositor |
| Map size (Å) | 263.64, 263.64, 263.64 | wwPDB |
| Map dimensions | 260, 260, 260 | wwPDB |
| Map angles (°) | 90.0, 90.0, 90.0 | wwPDB |
| Pixel spacing (Å) | 1.014, 1.014, 1.014 | Depositor |

## 5 Model quality

### 5.1 Standard geometry

Bond lengths and bond angles in the following residue types are not validated in this section: NAG

| Mol | Chain | Bond lengths |  | Bond angles |  |
| --- | --- | --- | --- | --- | --- |
|  |  | RMSZ | #\|Z\| >5 | RMSZ | #\|Z\| >5 |
| 1 | A | 0.27 | 0/1501 | 0.40 | 0/2040 |
| 1 | B | 0.27 | 0/1501 | 0.40 | 0/2040 |
| 1 | C | 0.27 | 0/1501 | 0.40 | 0/2040 |
| 1 | D | 0.27 | 0/1501 | 0.40 | 0/2040 |
| 1 | E | 0.27 | 0/1501 | 0.40 | 0/2040 |
| 1 | F | 0.27 | 0/1501 | 0.40 | 0/2040 |
| 1 | G | 0.27 | 0/1501 | 0.40 | 0/2040 |
| 1 | H | 0.27 | 0/1501 | 0.40 | 0/2040 |
| All | All | 0.27 | 0/12008 | 0.40 | 0/16320 |

There are no bond length outliers.

There are no bond angle outliers.

There are no chirality outliers.

There are no planarity outliers.

### 5.2 Too-close contacts

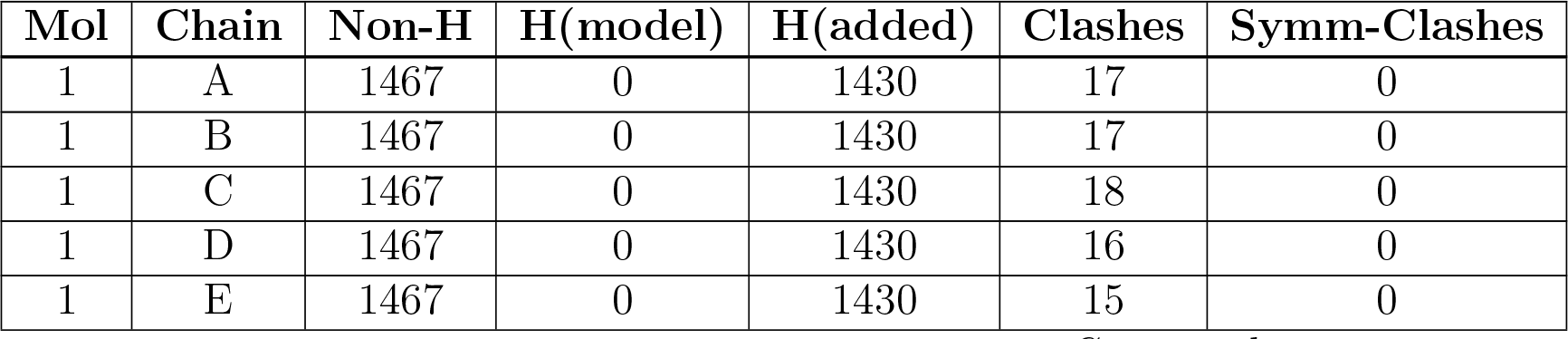
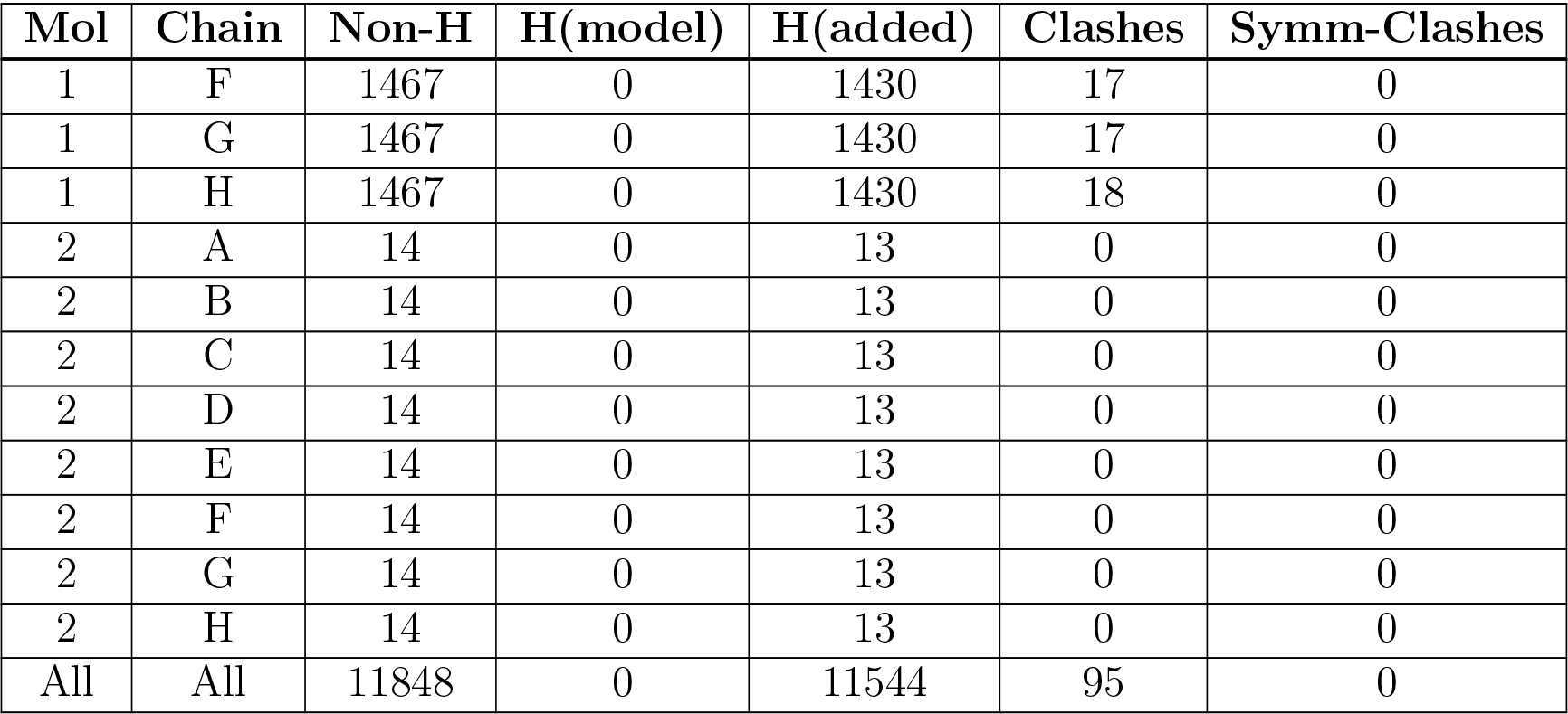

| Mol | Chain | Non-H | H(model) | H(added) | Clashes | Symm-Clashes |
| --- | --- | --- | --- | --- | --- | --- |
| 1 | A | 1467 | 0 | 1430 | 17 | 0 |
| 1 | B | 1467 | 0 | 1430 | 17 | 0 |
| 1 | C | 1467 | 0 | 1430 | 18 | 0 |
| 1 | D | 1467 | 0 | 1430 | 16 | 0 |
| 1 | E | 1467 | 0 | 1430 | 15 | 0 |
| 1 | F | 1467 | 0 | 1430 | 17 | 0 |
| 1 | G | 1467 | 0 | 1430 | 17 | 0 |
| 1 | H | 1467 | 0 | 1430 | 18 | 0 |
| 2 | A | 14 | 0 | 13 | 0 | 0 |
| 2 | B | 14 | 0 | 13 | 0 | 0 |
| 2 | C | 14 | 0 | 13 | 0 | 0 |
| 2 | D | 14 | 0 | 13 | 0 | 0 |
| 2 | E | 14 | 0 | 13 | 0 | 0 |
| 2 | F | 14 | 0 | 13 | 0 | 0 |
| 2 | G | 14 | 0 | 13 | 0 | 0 |
| 2 | H | 14 | 0 | 13 | 0 | 0 |
| All | All | 11848 | 0 | 11544 | 95 | 0 |

The all-atom clashscore is defined as the number of clashes found per 1000 atoms (including hydrogen atoms). The all-atom clashscore for this structure is 4.

All (95) close contacts within the same asymmetric unit are listed below, sorted by their clash magnitude.

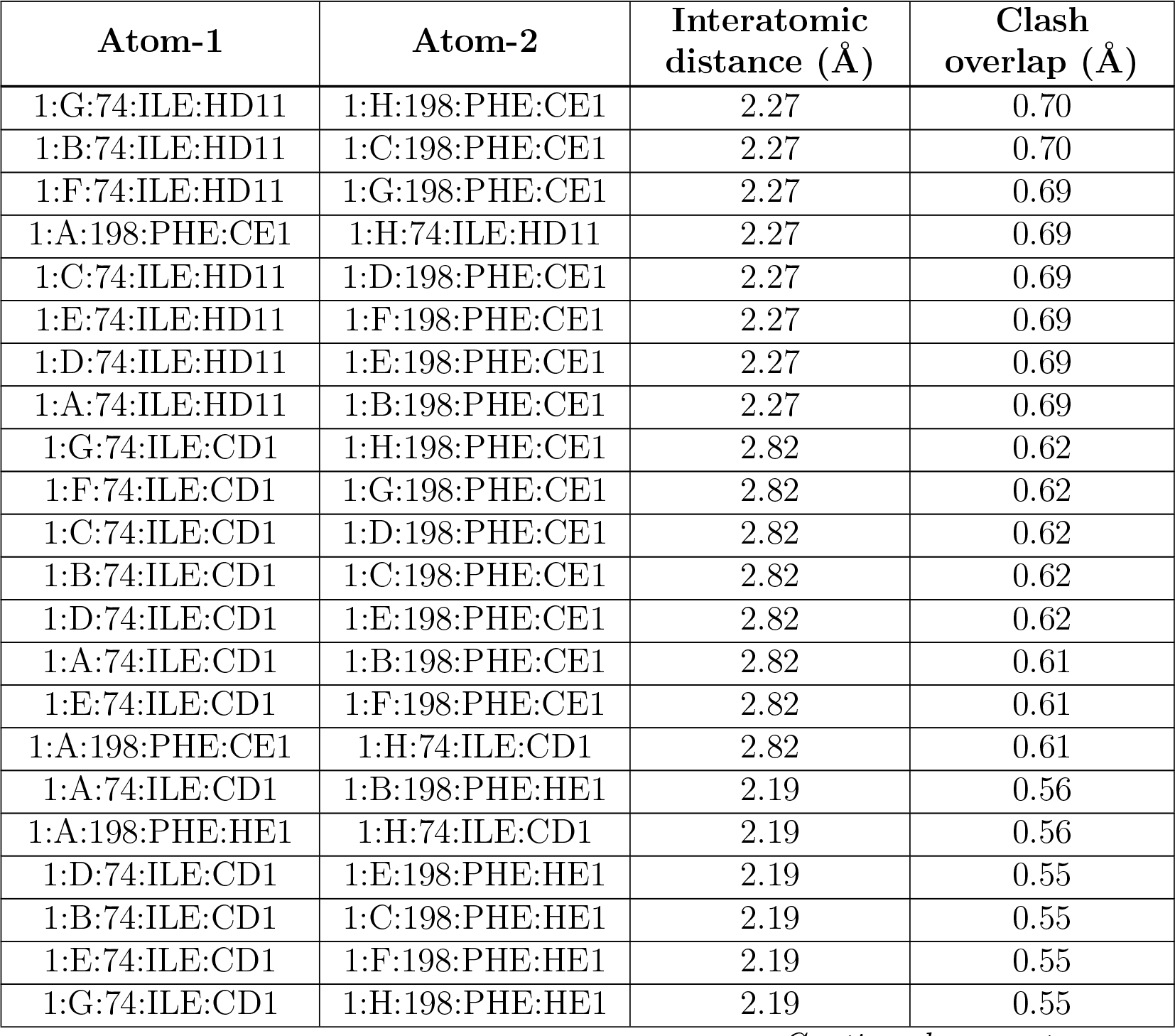
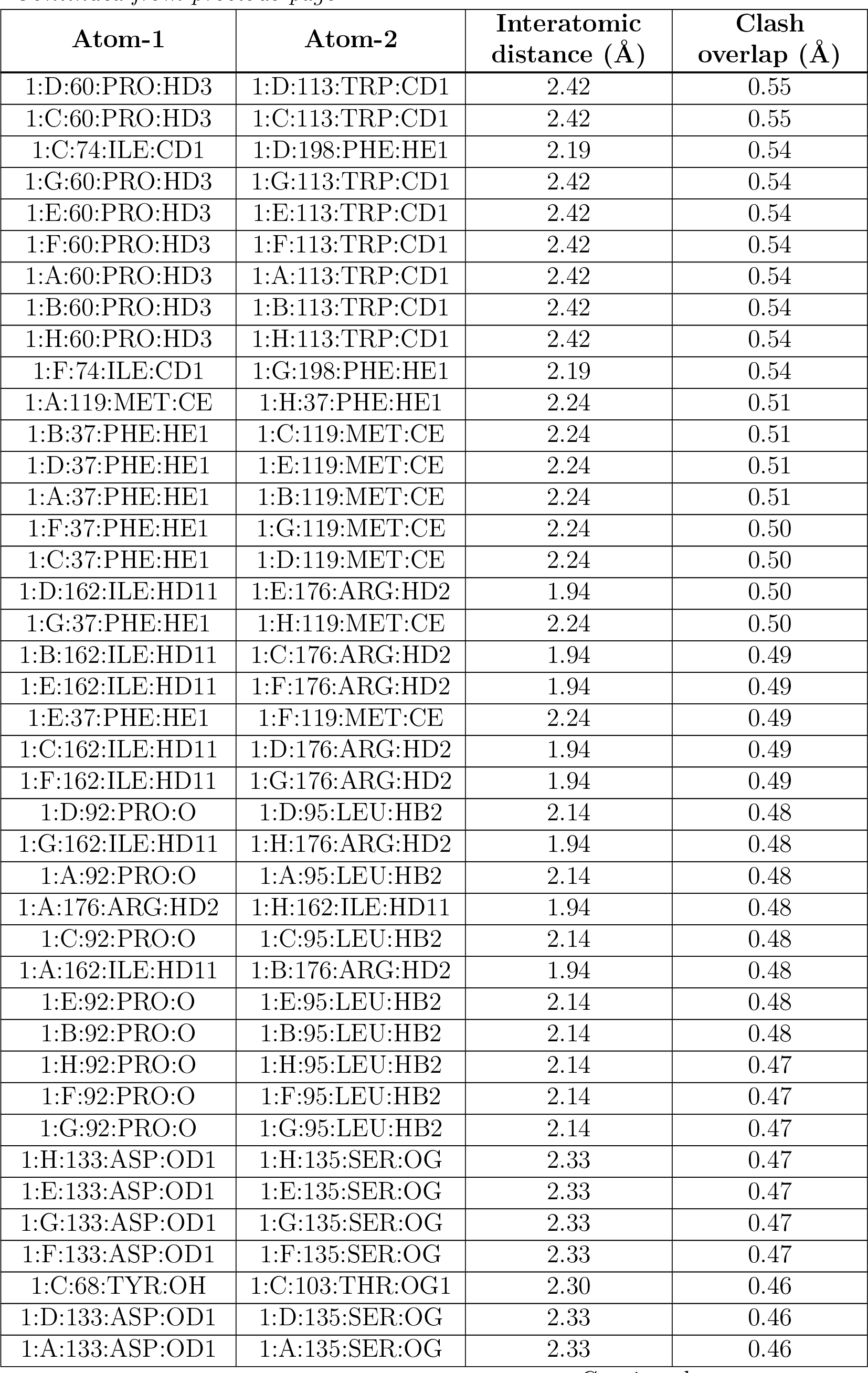
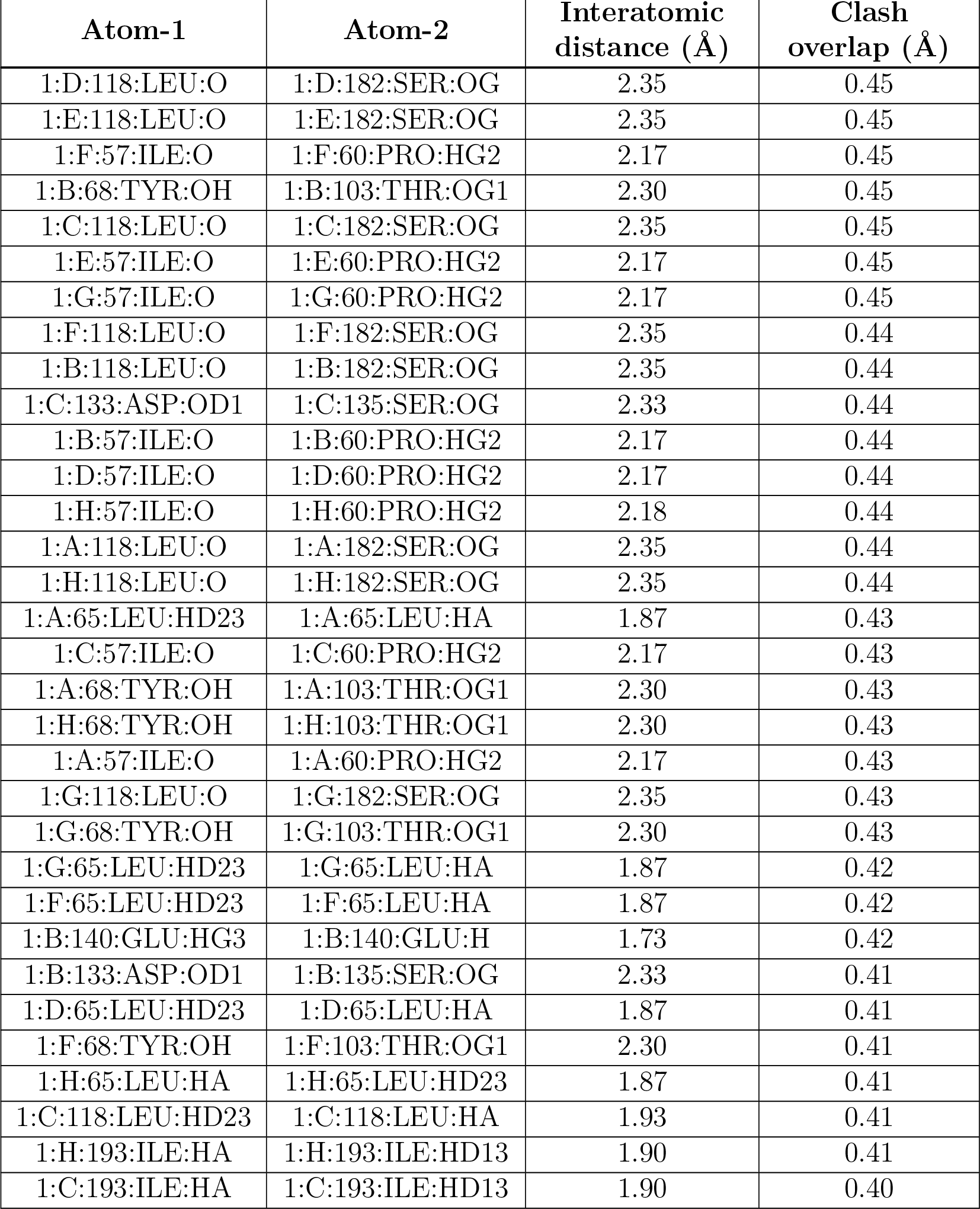

| Atom-1 | Atom-2 | Interatomic distance (Å) | Clash overlap (Å) |
| --- | --- | --- | --- |
| 1:G:74:ILE:HD11 | 1:H:198:PHE:CE1 | 2.27 | 0.70 |
| 1:B:74:ILE:HD11 | 1:C:198:PHE:CE1 | 2.27 | 0.70 |
| 1:F:74:ILE:HD11 | 1:G:198:PHE:CE1 | 2.27 | 0.69 |
| 1:A:198:PHE:CE1 | 1:H:74:ILE:HD11 | 2.27 | 0.69 |
| 1:C:74:ILE:HD11 | 1:D:198:PHE:CE1 | 2.27 | 0.69 |
| 1:E:74:ILE:HD11 | 1:F:198:PHE:CE1 | 2.27 | 0.69 |
| 1:D:74:ILE:HD11 | 1:E:198:PHE:CE1 | 2.27 | 0.69 |
| 1:A:74:ILE:HD11 | 1:B:198:PHE:CE1 | 2.27 | 0.69 |
| 1:G:74:ILE:CD1 | 1:H:198:PHE:CE1 | 2.82 | 0.62 |
| 1:F:74:ILE:CD1 | 1:G:198:PHE:CE1 | 2.82 | 0.62 |
| 1:C:74:ILE:CD1 | 1:D:198:PHE:CE1 | 2.82 | 0.62 |
| 1:B:74:ILE:CD1 | 1:C:198:PHE:CE1 | 2.82 | 0.62 |
| 1:D:74:ILE:CD1 | 1:E:198:PHE:CE1 | 2.82 | 0.62 |
| 1:A:74:ILE:CD1 | 1:B:198:PHE:CE1 | 2.82 | 0.61 |
| 1:E:74:ILE:CD1 | 1:F:198:PHE:CE1 | 2.82 | 0.61 |
| 1:A:198:PHE:CE1 | 1:H:74:ILE:CD1 | 2.82 | 0.61 |
| 1:A:74:ILE:CD1 | 1:B:198:PHE:HE1 | 2.19 | 0.56 |
| 1:A:198:PHE:HE1 | 1:H:74:ILE:CD1 | 2.19 | 0.56 |
| 1:D:74:ILE:CD1 | 1:E:198:PHE:HE1 | 2.19 | 0.55 |
| 1:B:74:ILE:CD1 | 1:C:198:PHE:HE1 | 2.19 | 0.55 |
| 1:E:74:ILE:CD1 | 1:F:198:PHE:HE1 | 2.19 | 0.55 |
| 1:G:74:ILE:CD1 | 1:H:198:PHE:HE1 | 2.19 | 0.55 |
| 1:D:60:PRO:HD3 | 1:D:113:TRP:CD1 | 2.42 | 0.55 |
| 1:C:60:PRO:HD3 | 1:C:113:TRP:CD1 | 2.42 | 0.55 |
| 1:C:74:ILE:CD1 | 1:D:198:PHE:HE1 | 2.19 | 0.54 |
| 1:G:60:PRO:HD3 | 1:G:113:TRP:CD1 | 2.42 | 0.54 |
| 1:E:60:PRO:HD3 | 1:E:113:TRP:CD1 | 2.42 | 0.54 |
| 1:F:60:PRO:HD3 | 1:F:113:TRP:CD1 | 2.42 | 0.54 |
| 1:A:60:PRO:HD3 | 1:A:113:TRP:CD1 | 2.42 | 0.54 |
| 1:B:60:PRO:HD3 | 1:B:113:TRP:CD1 | 2.42 | 0.54 |
| 1:H:60:PRO:HD3 | 1:H:113:TRP:CD1 | 2.42 | 0.54 |
| 1:F:74:ILE:CD1 | 1:G:198:PHE:HE1 | 2.19 | 0.54 |
| 1:A:119:MET:CE | 1:H:37:PHE:HE1 | 2.24 | 0.51 |
| 1:B:37:PHE:HE1 | 1:C:119:MET:CE | 2.24 | 0.51 |
| 1:D:37:PHE:HE1 | 1:E:119:MET:CE | 2.24 | 0.51 |
| 1:A:37:PHE:HE1 | 1:B:119:MET:CE | 2.24 | 0.51 |
| 1:F:37:PHE:HE1 | 1:G:119:MET:CE | 2.24 | 0.50 |
| 1:C:37:PHE:HE1 | 1:D:119:MET:CE | 2.24 | 0.50 |
| 1:D:162:ILE:HD11 | 1:E:176:ARG:HD2 | 1.94 | 0.50 |
| 1:G:37:PHE:HE1 | 1:H:119:MET:CE | 2.24 | 0.50 |
| 1:B:162:ILE:HD11 | 1:C:176:ARG:HD2 | 1.94 | 0.49 |
| 1:E:162:ILE:HD11 | 1:F:176:ARG:HD2 | 1.94 | 0.49 |
| 1:E:37:PHE:HE1 | 1:F:119:MET:CE | 2.24 | 0.49 |
| 1:C:162:ILE:HD11 | 1:D:176:ARG:HD2 | 1.94 | 0.49 |
| 1:F:162:ILE:HD11 | 1:G:176:ARG:HD2 | 1.94 | 0.49 |
| 1:D:92:PRO:O | 1:D:95:LEU:HB2 | 2.14 | 0.48 |
| 1:G:162:ILE:HD11 | 1:H:176:ARG:HD2 | 1.94 | 0.48 |
| 1:A:92:PRO:O | 1:A:95:LEU:HB2 | 2.14 | 0.48 |
| 1:A:176:ARG:HD2 | 1:H:162:ILE:HD11 | 1.94 | 0.48 |
| 1:C:92:PRO:O | 1:C:95:LEU:HB2 | 2.14 | 0.48 |
| 1:A:162:ILE:HD11 | 1:B:176:ARG:HD2 | 1.94 | 0.48 |
| 1:E:92:PRO:O | 1:E:95:LEU:HB2 | 2.14 | 0.48 |
| 1:B:92:PRO:O | 1:B:95:LEU:HB2 | 2.14 | 0.48 |
| 1:H:92:PRO:O | 1:H:95:LEU:HB2 | 2.14 | 0.47 |
| 1:F:92:PRO:O | 1:F:95:LEU:HB2 | 2.14 | 0.47 |
| 1:G:92:PRO:O | 1:G:95:LEU:HB2 | 2.14 | 0.47 |
| 1:H:133:ASP:OD1 | 1:H:135:SER:OG | 2.33 | 0.47 |
| 1:E:133:ASP:OD1 | 1:E:135:SER:OG | 2.33 | 0.47 |
| 1:G:133:ASP:OD1 | 1:G:135:SER:OG | 2.33 | 0.47 |
| 1:F:133:ASP:OD1 | 1:F:135:SER:OG | 2.33 | 0.47 |
| 1:C:68:TYR:OH | 1:C:103:THR:OG1 | 2.30 | 0.46 |
| 1:D:133:ASP:OD1 | 1:D:135:SER:OG | 2.33 | 0.46 |
| 1:A:133:ASP:OD1 | 1:A:135:SER:OG | 2.33 | 0.46 |
| 1:D:118:LEU:O | 1:D:182:SER:OG | 2.35 | 0.45 |
| 1:E:118:LEU:O | 1:E:182:SER:OG | 2.35 | 0.45 |
| 1:F:57:ILE:O | 1:F:60:PRO:HG2 | 2.17 | 0.45 |
| 1:B:68:TYR:OH | 1:B:103:THR:OG1 | 2.30 | 0.45 |
| 1:C:118:LEU:O | 1:C:182:SER:OG | 2.35 | 0.45 |
| 1:E:57:ILE:O | 1:E:60:PRO:HG2 | 2.17 | 0.45 |
| 1:G:57:ILE:O | 1:G:60:PRO:HG2 | 2.17 | 0.45 |
| 1:F:118:LEU:O | 1:F:182:SER:OG | 2.35 | 0.44 |
| 1:B:118:LEU:O | 1:B:182:SER:OG | 2.35 | 0.44 |
| 1:C:133:ASP:OD1 | 1:C:135:SER:OG | 2.33 | 0.44 |
| 1:B:57:ILE:O | 1:B:60:PRO:HG2 | 2.17 | 0.44 |
| 1:D:57:ILE:O | 1:D:60:PRO:HG2 | 2.17 | 0.44 |
| 1:H:57:ILE:O | 1:H:60:PRO:HG2 | 2.18 | 0.44 |
| 1:A:118:LEU:O | 1:A:182:SER:OG | 2.35 | 0.44 |
| 1:H:118:LEU:O | 1:H:182:SER:OG | 2.35 | 0.44 |
| 1:A:65:LEU:HD23 | 1:A:65:LEU:HA | 1.87 | 0.43 |
| 1:C:57:ILE:O | 1:C:60:PRO:HG2 | 2.17 | 0.43 |
| 1:A:68:TYR:OH | 1:A:103:THR:OG1 | 2.30 | 0.43 |
| 1:H:68:TYR:OH | 1:H:103:THR:OG1 | 2.30 | 0.43 |
| 1:A:57:ILE:O | 1:A:60:PRO:HG2 | 2.17 | 0.43 |
| 1:G:118:LEU:O | 1:G:182:SER:OG | 2.35 | 0.43 |
| 1:G:68:TYR:OH | 1:G:103:THR:OG1 | 2.30 | 0.43 |
| 1:G:65:LEU:HD23 | 1:G:65:LEU:HA | 1.87 | 0.42 |
| 1:F:65:LEU:HD23 | 1:F:65:LEU:HA | 1.87 | 0.42 |
| 1:B:140:GLU:HG3 | 1:B:140:GLU:H | 1.73 | 0.42 |
| 1:B:133:ASP:OD1 | 1:B:135:SER:OG | 2.33 | 0.41 |
| 1:D:65:LEU:HD23 | 1:D:65:LEU:HA | 1.87 | 0.41 |
| 1:F:68:TYR:OH | 1:F:103:THR:OG1 | 2.30 | 0.41 |
| 1:H:65:LEU:HA | 1:H:65:LEU:HD23 | 1.87 | 0.41 |
| 1:C:118:LEU:HD23 | 1:C:118:LEU:HA | 1.93 | 0.41 |
| 1:H:193:ILE:HA | 1:H:193:ILE:HD13 | 1.90 | 0.41 |
| 1:C:193:ILE:HA | 1:C:193:ILE:HD13 | 1.90 | 0.40 |

There are no symmetry-related clashes.

### 5.3 Torsion angles

#### 5.3.1 Protein backbone

| Mol | Chain | Analysed | Favoured | Allowed | Outliers | Percentiles |  |
| --- | --- | --- | --- | --- | --- | --- | --- |
| 1 | A | 185/358 (52%) | 182 (98%) | 3 (2%) | 0 | 100 | 100 |
| 1 | B | 185/358 (52%) | 182 (98%) | 3 (2%) | 0 | 100 | 100 |
| 1 | C | 185/358 (52%) | 182 (98%) | 3 (2%) | 0 | 100 | 100 |
| 1 | D | 185/358 (52%) | 182 (98%) | 3 (2%) | 0 | 100 | 100 |
| 1 | E | 185/358 (52%) | 182 (98%) | 3 (2%) | 0 | 100 | 100 |
| 1 | F | 185/358 (52%) | 182 (98%) | 3 (2%) | 0 | 100 | 100 |
| 1 | G | 185/358 (52%) | 182 (98%) | 3 (2%) | 0 | 100 | 100 |
| 1 | H | 185/358 (52%) | 182 (98%) | 3 (2%) | 0 | 100 | 100 |
| All | All | 1480/2864 (52%) | 1456 (98%) | 24 (2%) | 0 | 100 | 100 |

There are no Ramachandran outliers to report.

#### 5.3.2 Protein sidechains

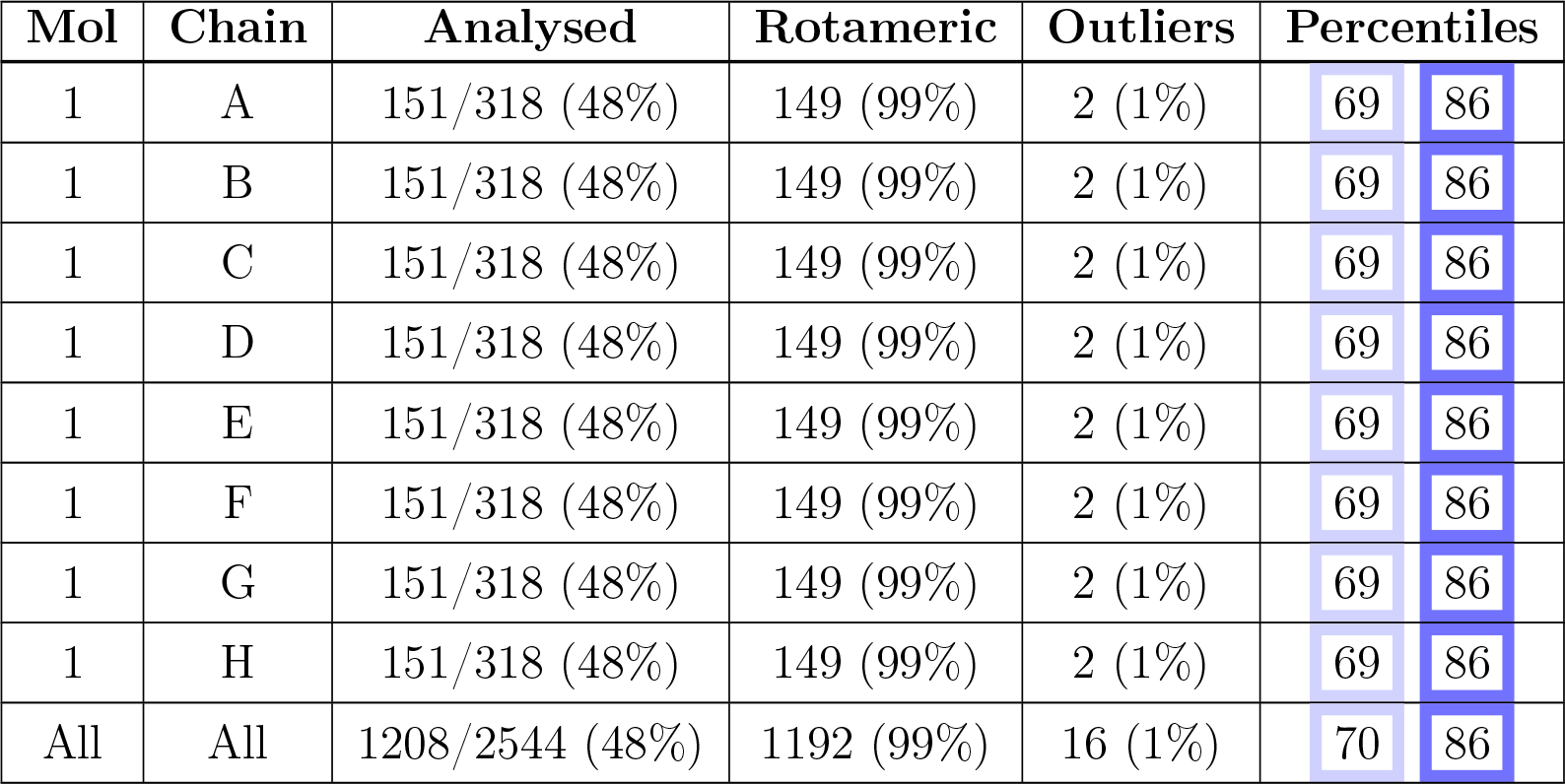

| Mol | Chain | Analysed | Rotameric | Outliers | Percentiles |  |
| --- | --- | --- | --- | --- | --- | --- |
| 1 | A | 151/318 (48%) | 149 (99%) | 2 (1%) | 69 | 86 |
| 1 | B | 151/318 (48%) | 149 (99%) | 2 (1%) | 69 | 86 |
| 1 | C | 151/318 (48%) | 149 (99%) | 2 (1%) | 69 | 86 |
| 1 | D | 151/318 (48%) | 149 (99%) | 2 (1%) | 69 | 86 |
| 1 | E | 151/318 (48%) | 149 (99%) | 2 (1%) | 69 | 86 |
| 1 | F | 151/318 (48%) | 149 (99%) | 2 (1%) | 69 | 86 |
| 1 | G | 151/318 (48%) | 149 (99%) | 2 (1%) | 69 | 86 |
| 1 | H | 151/318 (48%) | 149 (99%) | 2 (1%) | 69 | 86 |
| All | All | 1208/2544 (48%) | 1192 (99%) | 16 (1%) | 70 | 86 |

All (16) residues with a non-rotameric sidechain are listed below:

| Mol | Chain | Res | Type |
| --- | --- | --- | --- |
| 1 | A | 68 | TYR |
| 1 | A | 124 | PHE |
| 1 | B | 68 | TYR |
| 1 | B | 124 | PHE |
| 1 | C | 68 | TYR |
| 1 | C | 124 | PHE |
| 1 | D | 68 | TYR |
| 1 | D | 124 | PHE |
| 1 | E | 68 | TYR |
| 1 | E | 124 | PHE |
| 1 | F | 68 | TYR |
| 1 | F | 124 | PHE |
| 1 | G | 68 | TYR |
| 1 | G | 124 | PHE |
| 1 | H | 68 | TYR |
| 1 | H | 124 | PHE |

Sometimes sidechains can be flipped to improve hydrogen bonding and reduce clashes. There are no such sidechains identified.

#### 5.3.3 RNA

There are no RNA molecules in this entry.

### 5.4 Non-standard residues in protein, DNA, RNA chains

There are no non-standard protein/DNA/RNA residues in this entry.

### 5.5 Carbohydrates

There are no monosaccharides in this entry.

### 5.6 Ligand geometry

8 ligands are modelled in this entry.

| Mol | Type | Chain | Res | Link | Bond lengths |  |  | Bond angles |  |  |
| --- | --- | --- | --- | --- | --- | --- | --- | --- | --- | --- |
| | | | | | Counts | RMSZ | $\#\vert Z\vert > 2$ | Counts | RMSZ | $\#\vert Z\vert > 2$ |
| 2 | NAG | F | 1001 | 1 | 14,14,15 | 0.27 | 0 | 17,19,21 | 0.52 | 0 |
| 2 | NAG | D | 1001 | 1 | 14,14,15 | 0.26 | 0 | 17,19,21 | 0.52 | 0 |
| 2 | NAG | E | 1001 | 1 | 14,14,15 | 0.26 | 0 | 17,19,21 | 0.52 | 0 |
| 2 | NAG | C | 1001 | 1 | 14,14,15 | 0.26 | 0 | 17,19,21 | 0.52 | 0 |
| 2 | NAG | G | 1001 | 1 | 14,14,15 | 0.26 | 0 | 17,19,21 | 0.52 | 0 |
| 2 | NAG | H | 1001 | 1 | 14,14,15 | 0.26 | 0 | 17,19,21 | 0.52 | 0 |
| 2 | NAG | A | 1001 | 1 | 14,14,15 | 0.26 | 0 | 17,19,21 | 0.52 | 0 |
| 2 | NAG | B | 1001 | 1 | 14,14,15 | 0.27 | 0 | 17,19,21 | 0.52 | 0 |

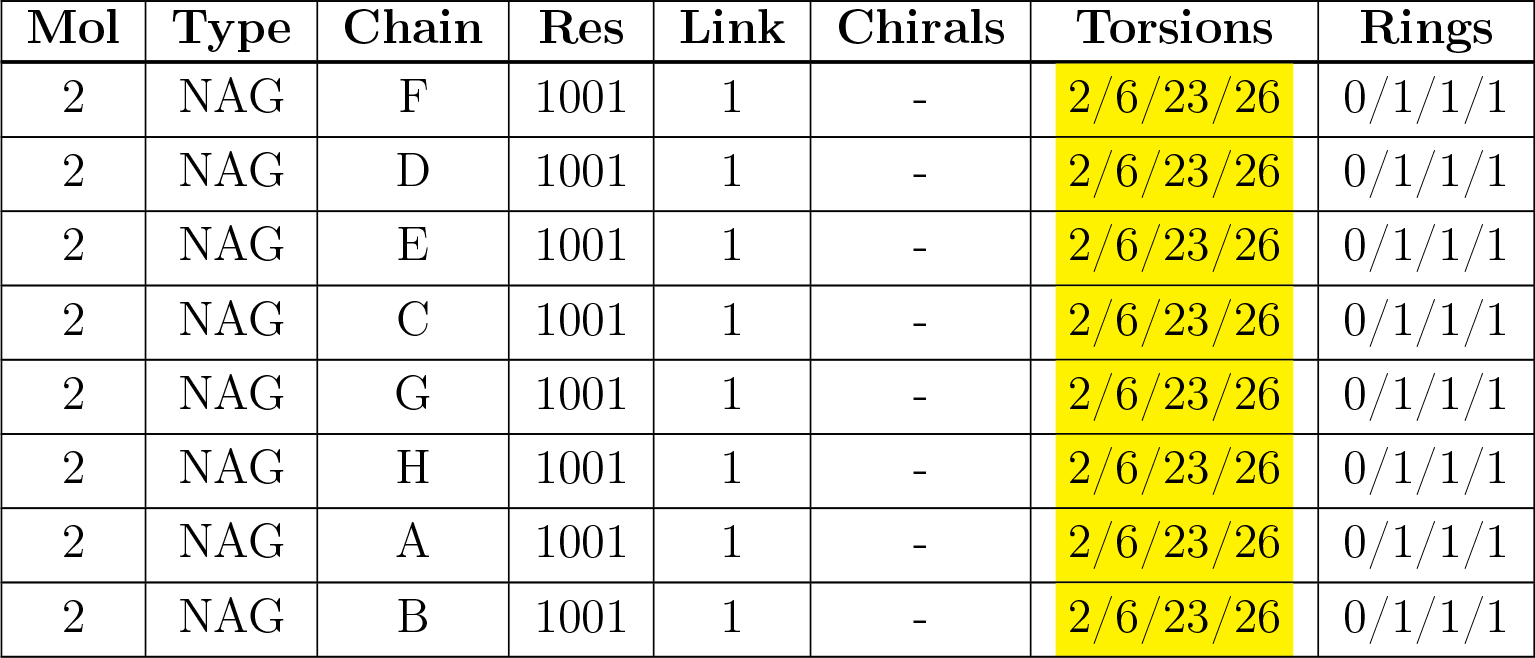

| Mol | Type | Chain | Res | Link | Chirals | Torsions | Rings |
| --- | --- | --- | --- | --- | --- | --- | --- |
| 2 | NAG | F | 1001 | 1 | - | 2/6/23/26 | 0/1/1/1 |
| 2 | NAG | D | 1001 | 1 | - | 2/6/23/26 | 0/1/1/1 |
| 2 | NAG | E | 1001 | 1 | - | 2/6/23/26 | 0/1/1/1 |
| 2 | NAG | C | 1001 | 1 | - | 2/6/23/26 | 0/1/1/1 |
| 2 | NAG | G | 1001 | 1 | - | 2/6/23/26 | 0/1/1/1 |
| 2 | NAG | H | 1001 | 1 | - | 2/6/23/26 | 0/1/1/1 |
| 2 | NAG | A | 1001 | 1 | - | 2/6/23/26 | 0/1/1/1 |
| 2 | NAG | B | 1001 | 1 | - | 2/6/23/26 | 0/1/1/1 |

There are no bond length outliers.

There are no bond angle outliers.

There are no chirality outliers.

All (16) torsion outliers are listed below:

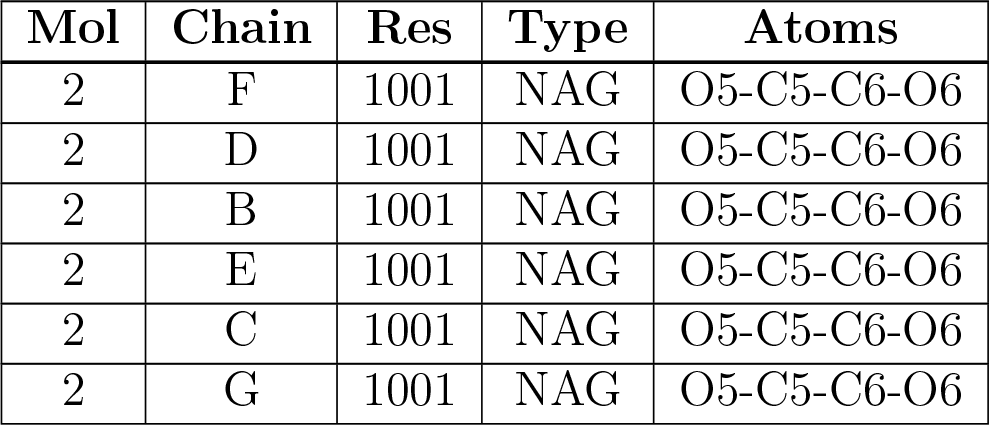
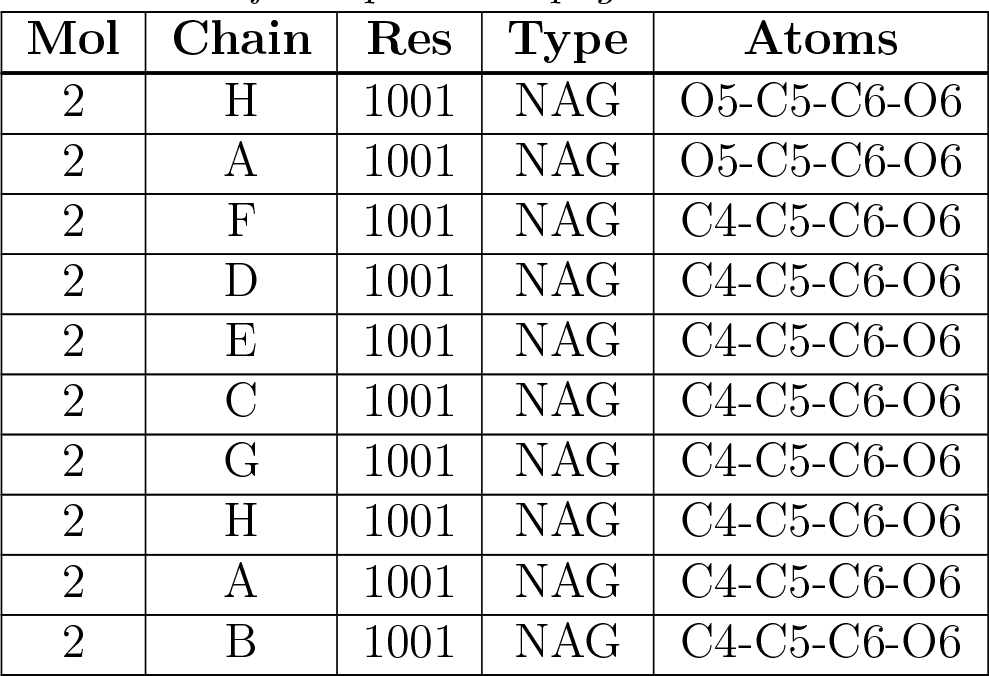

| Mol | Chain | Res | Type | Atoms |
| --- | --- | --- | --- | --- |
| 2 | F | 1001 | NAG | O5-C5-C6-O6 |
| 2 | D | 1001 | NAG | O5-C5-C6-O6 |
| 2 | B | 1001 | NAG | O5-C5-C6-O6 |
| 2 | E | 1001 | NAG | O5-C5-C6-O6 |
| 2 | C | 1001 | NAG | O5-C5-C6-O6 |
| 2 | G | 1001 | NAG | O5-C5-C6-O6 |
| 2 | H | 1001 | NAG | O5-C5-C6-O6 |
| 2 | A | 1001 | NAG | O5-C5-C6-O6 |
| 2 | F | 1001 | NAG | C4-C5-C6-O6 |
| 2 | D | 1001 | NAG | C4-C5-C6-O6 |
| 2 | E | 1001 | NAG | C4-C5-C6-O6 |
| 2 | C | 1001 | NAG | C4-C5-C6-O6 |
| 2 | G | 1001 | NAG | C4-C5-C6-O6 |
| 2 | H | 1001 | NAG | C4-C5-C6-O6 |
| 2 | A | 1001 | NAG | C4-C5-C6-O6 |
| 2 | B | 1001 | NAG | C4-C5-C6-O6 |

There are no ring outliers.

No monomer is involved in short contacts.

### 5.7 Other polymers

There are no such residues in this entry.

### 5.8 Polymer linkage issues

There are no chain breaks in this entry.

## 6 Map visualisation

This section contains visualisations of the EMDB entry EMD-30830. These allow visual inspection of the internal detail of the map and identification of artifacts.

### 6.1 Orthogonal projections

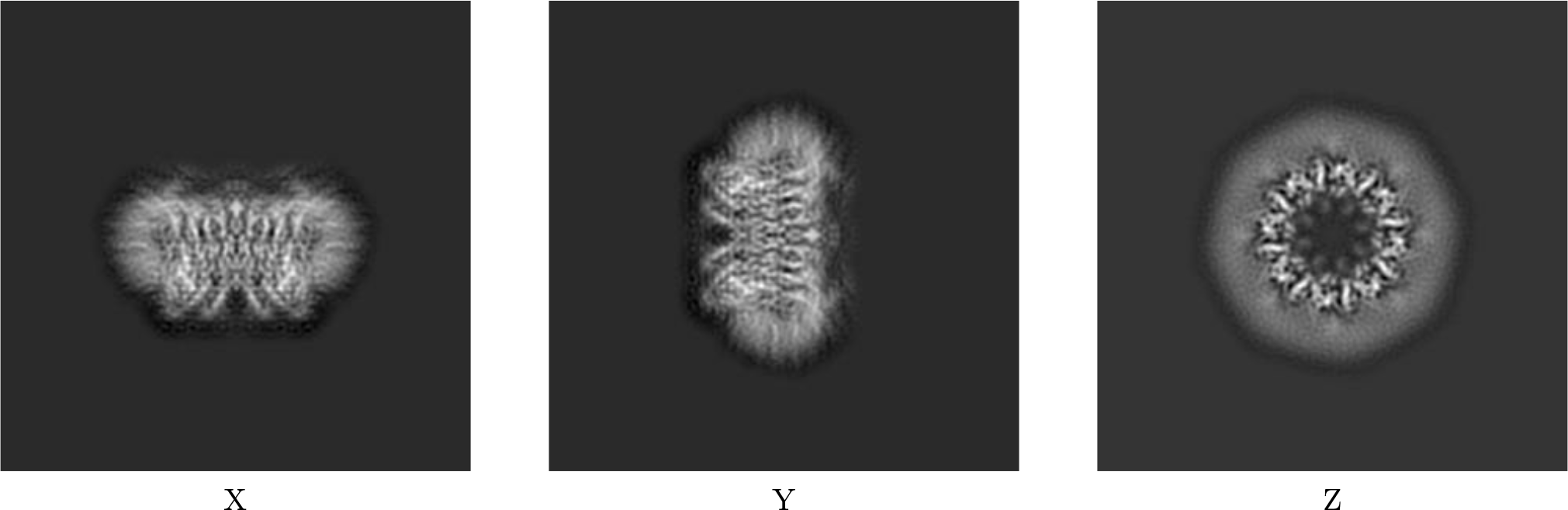

#### 6.1.1 Primary map

The images above show the map projected in three orthogonal directions.

### 6.2 Central slices

#### 6.2.1 Primary map

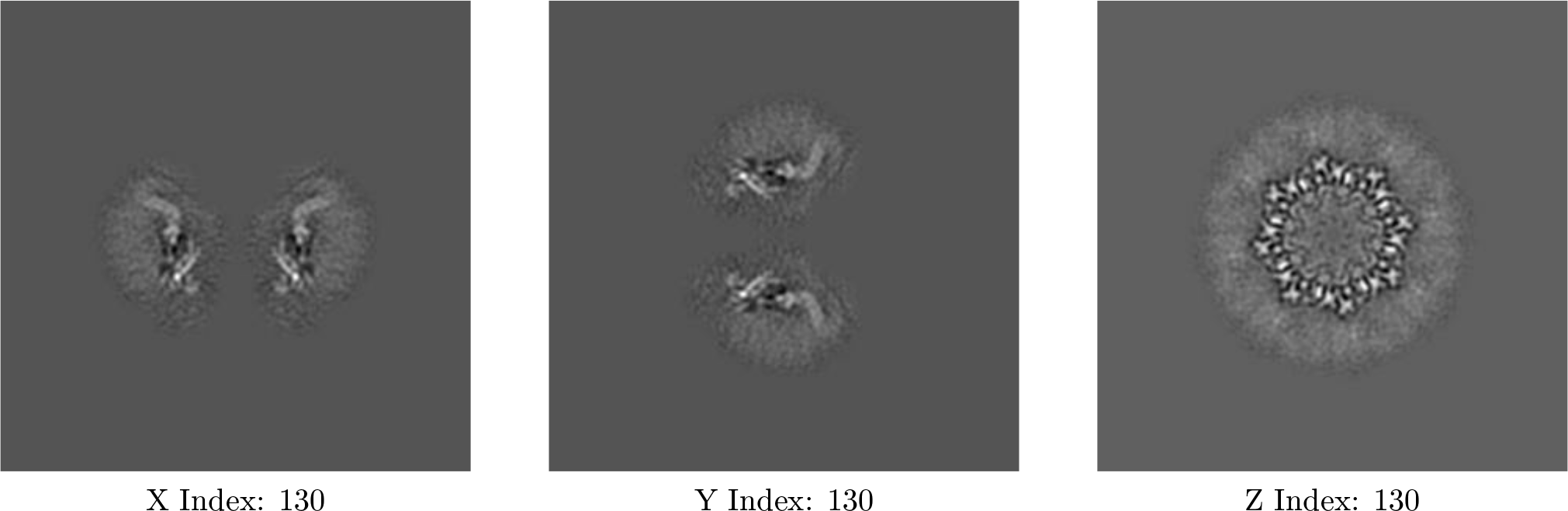

The images above show central slices of the map in three orthogonal directions.

### 6.3 Largest variance slices

#### 6.3.1 Primary map

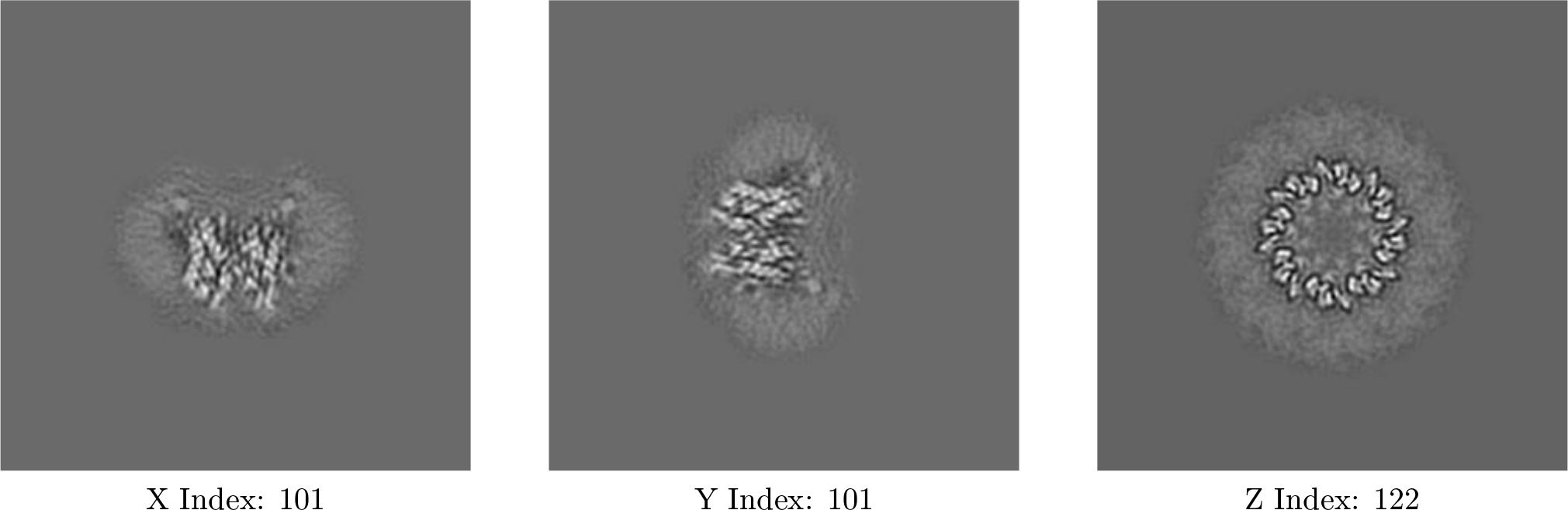

The images above show the largest variance slices of the map in three orthogonal directions.

### 6.4 Orthogonal surface views

#### 6.4.1 Primary map

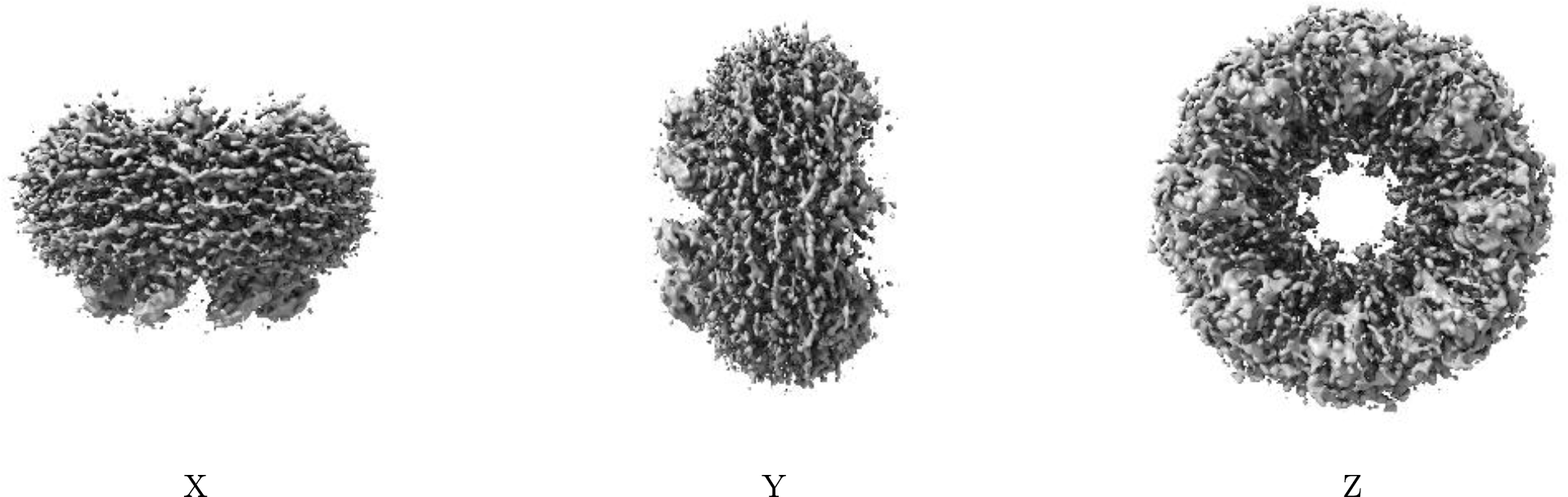

The images above show the 3D surface view of the map at the recommended contour level 0.0075. These images, in conjunction with the slice images, may facilitate assessment of whether an ap- propriate contour level has been provided.

### 6.5 Mask visualisation

This section was not generated. No masks/segmentation were deposited.

## 7 Map analysis

This section contains the results of statistical analysis of the map.

### 7.1 Map-value distribution

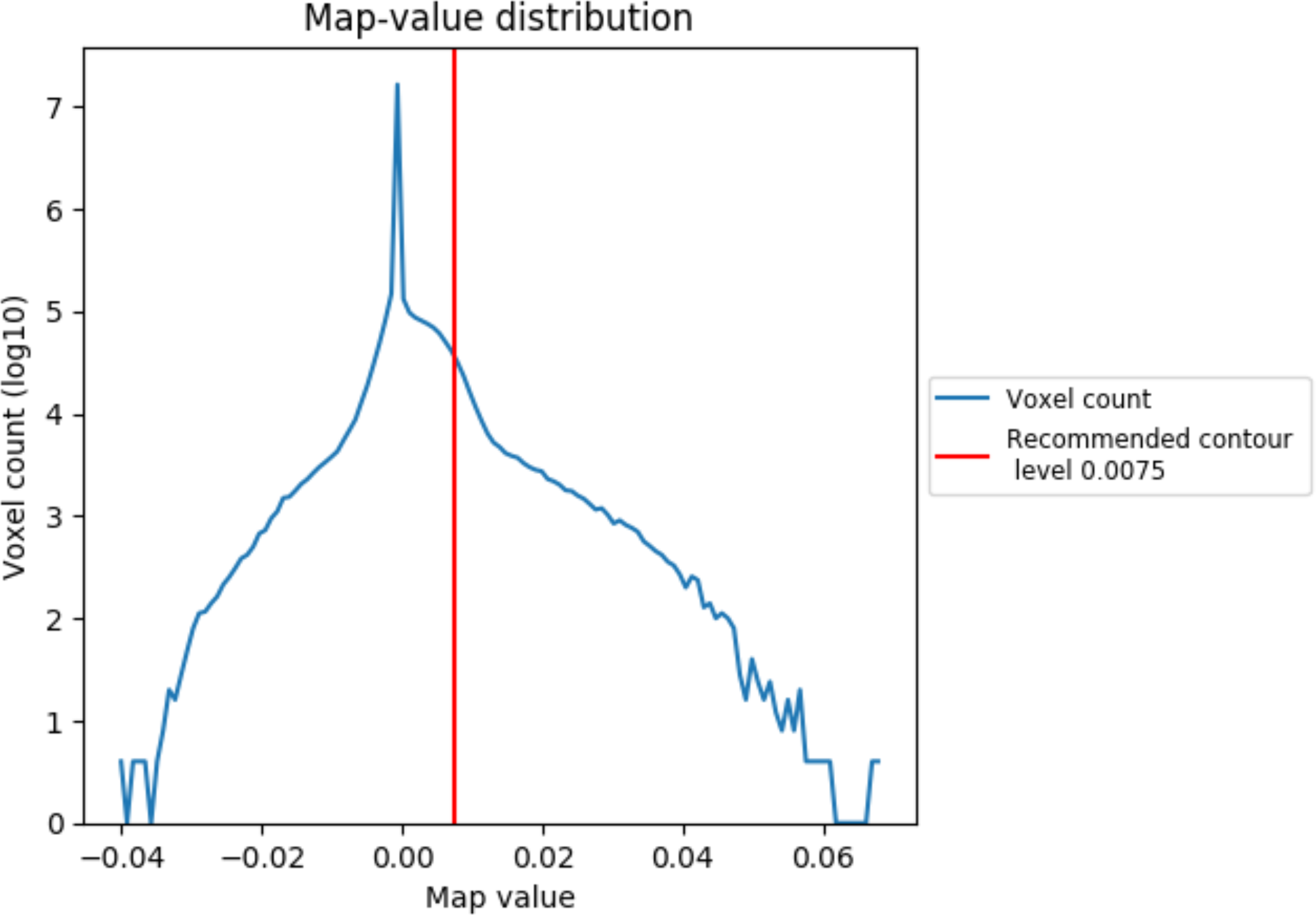

### 7.2 Volume estimate

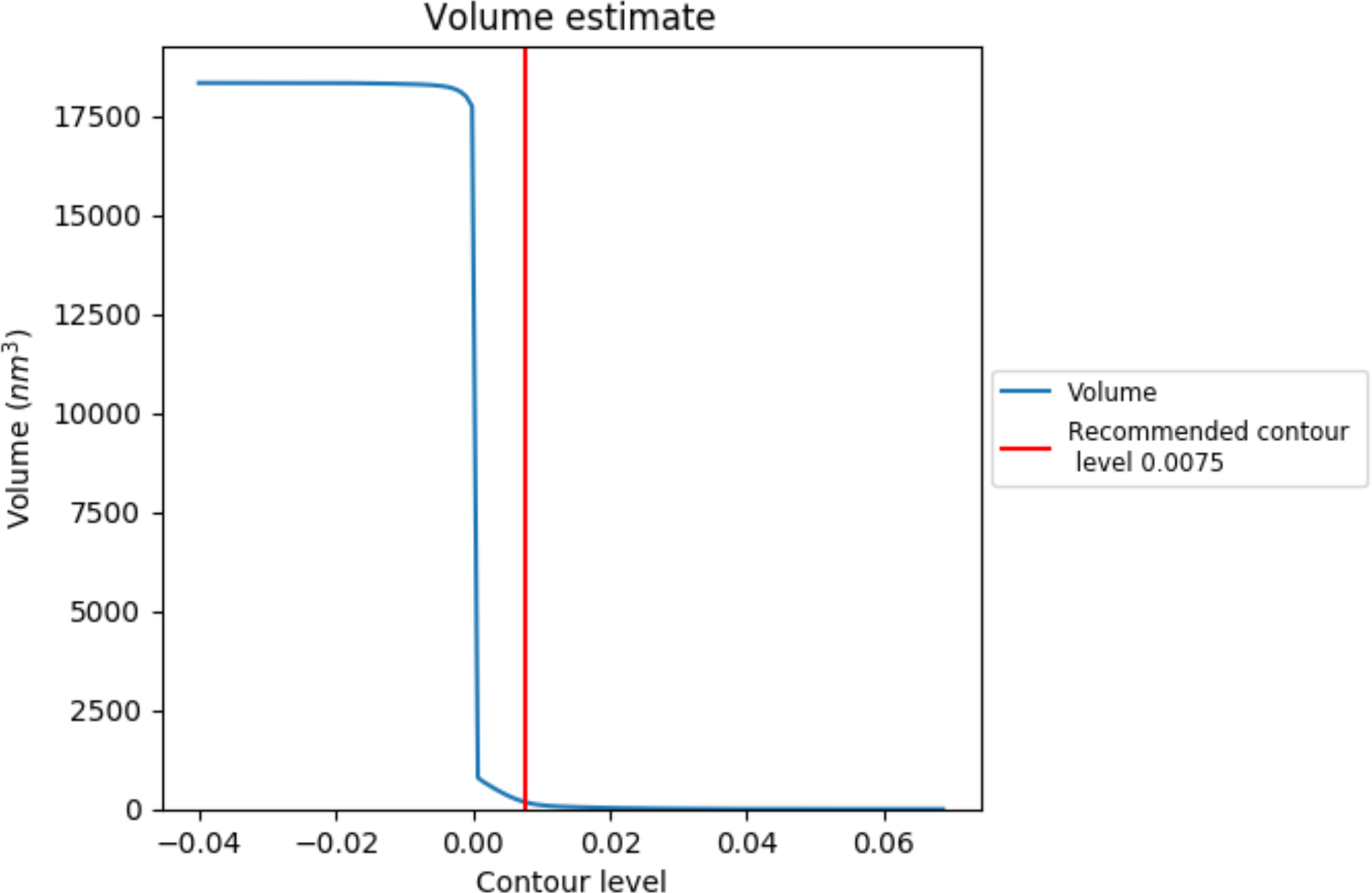

The volume at the recommended contour level is 182 nm^3^; this corresponds to an approximate mass of 164 kDa.

### 7.3 Rotationally averaged power spectrum

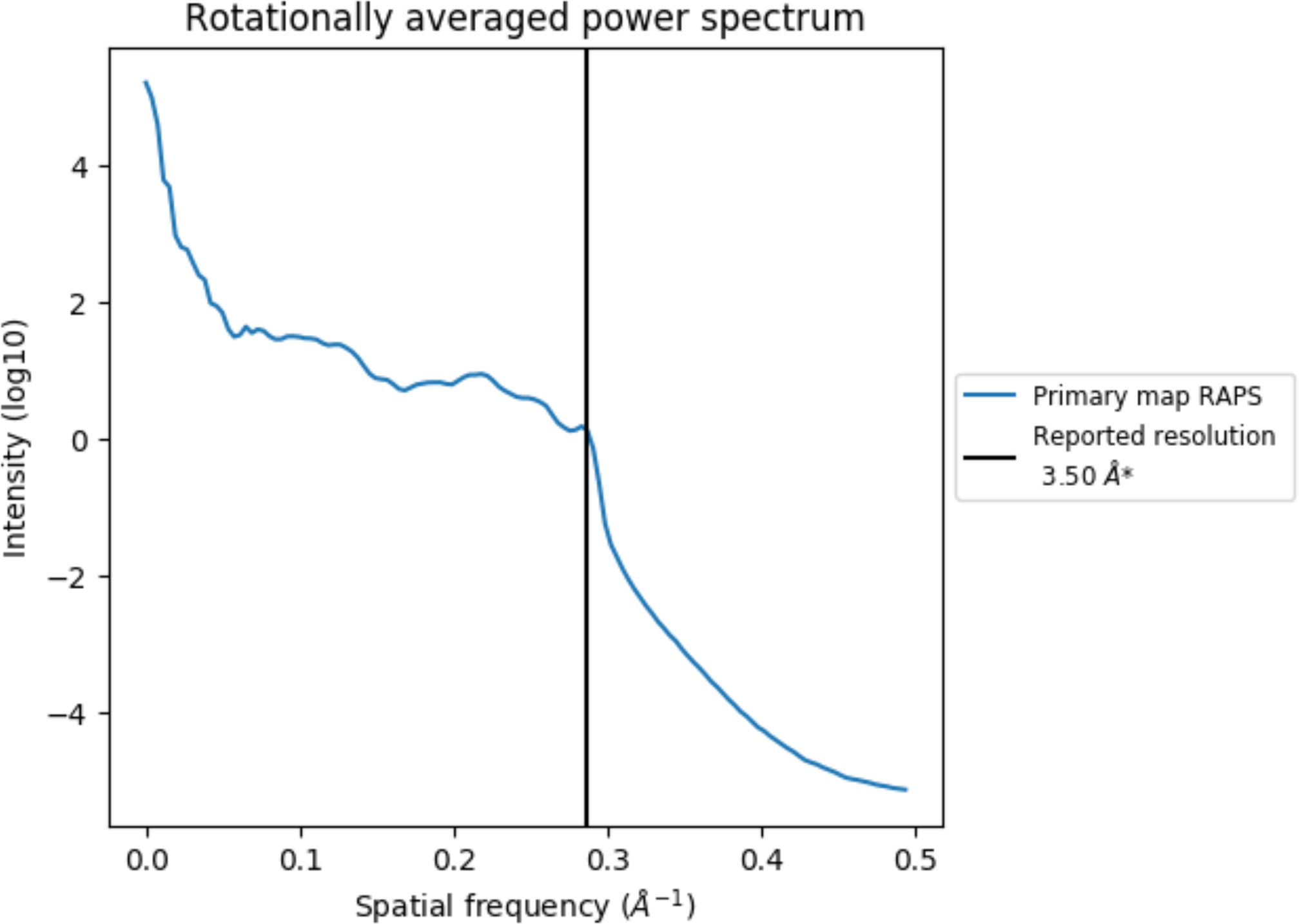

## 8 Fourier-Shell correlation

This section was not generated. No FSC curve or half-maps provided.

## 9 Map-model fit

This section contains information regarding the fit between EMDB map EMD-30830 and PDB model 7DSC. Per-residue inclusion information can be found in section 3 on page 7.

### 9.1 Map-model overlay

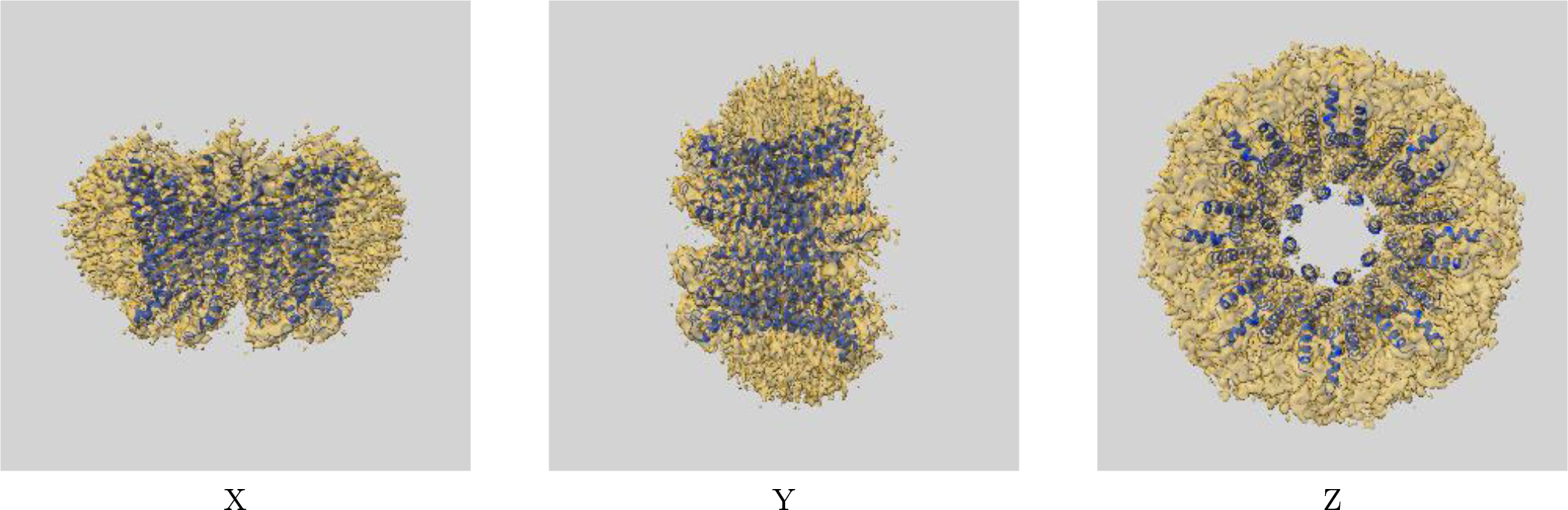

The images above show the 3D surface view of the map at the recommended contour level 0.0075 at 50% transparency in yellow overlaid with a ribbon representation of the model coloured in blue. These images allow for the visual assessment of the quality of fit between the atomic model and the map.

### 9.2 Atom inclusion

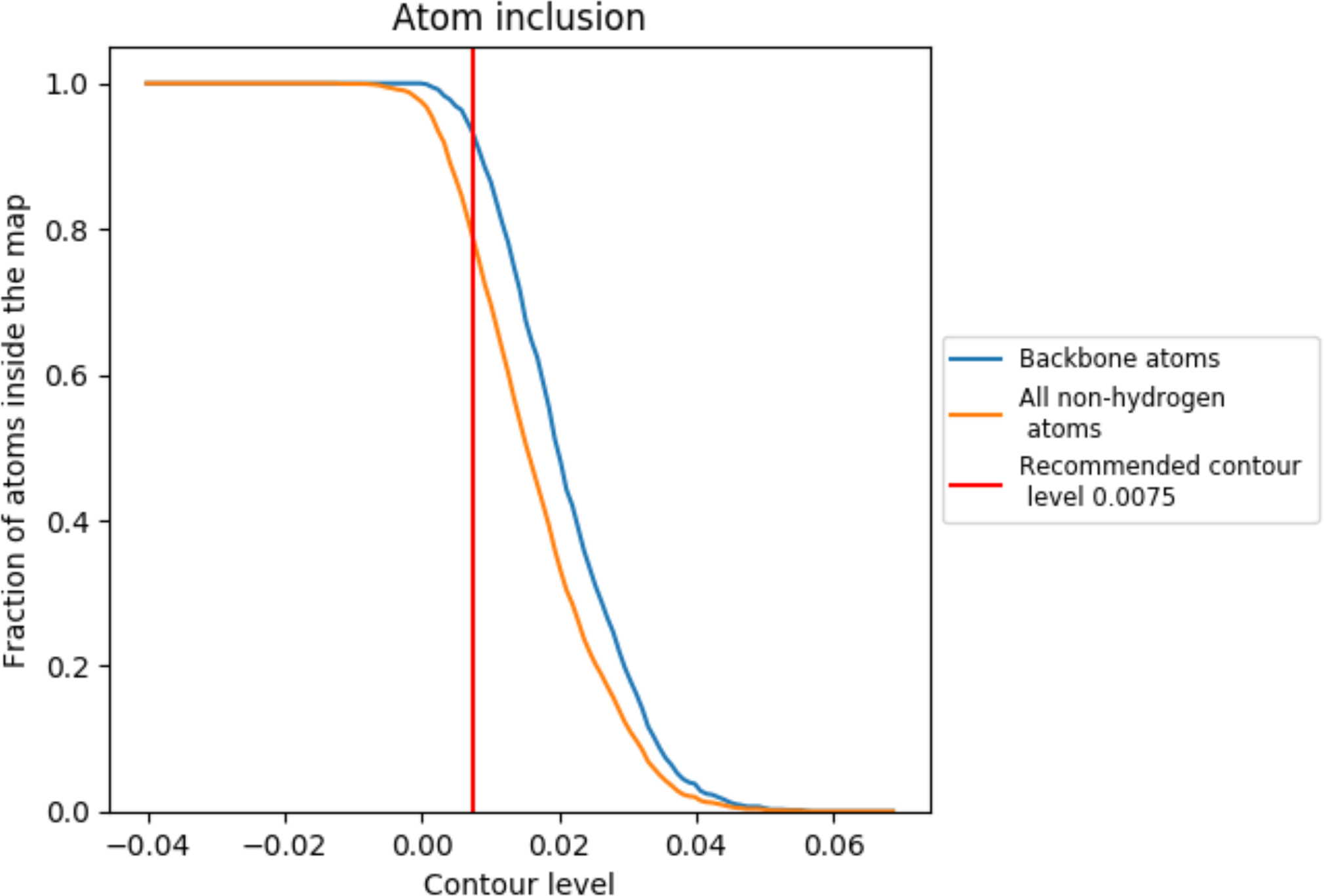

At the recommended contour level, 93% of all backbone atoms, 79% of all non-hydrogen atoms, are inside the map.

## 1 Overall quality at a glance

The following experimental techniques were used to determine the structure: *ELECTRON MICROSCOPY*

The reported resolution of this entry is 3.20 Å.

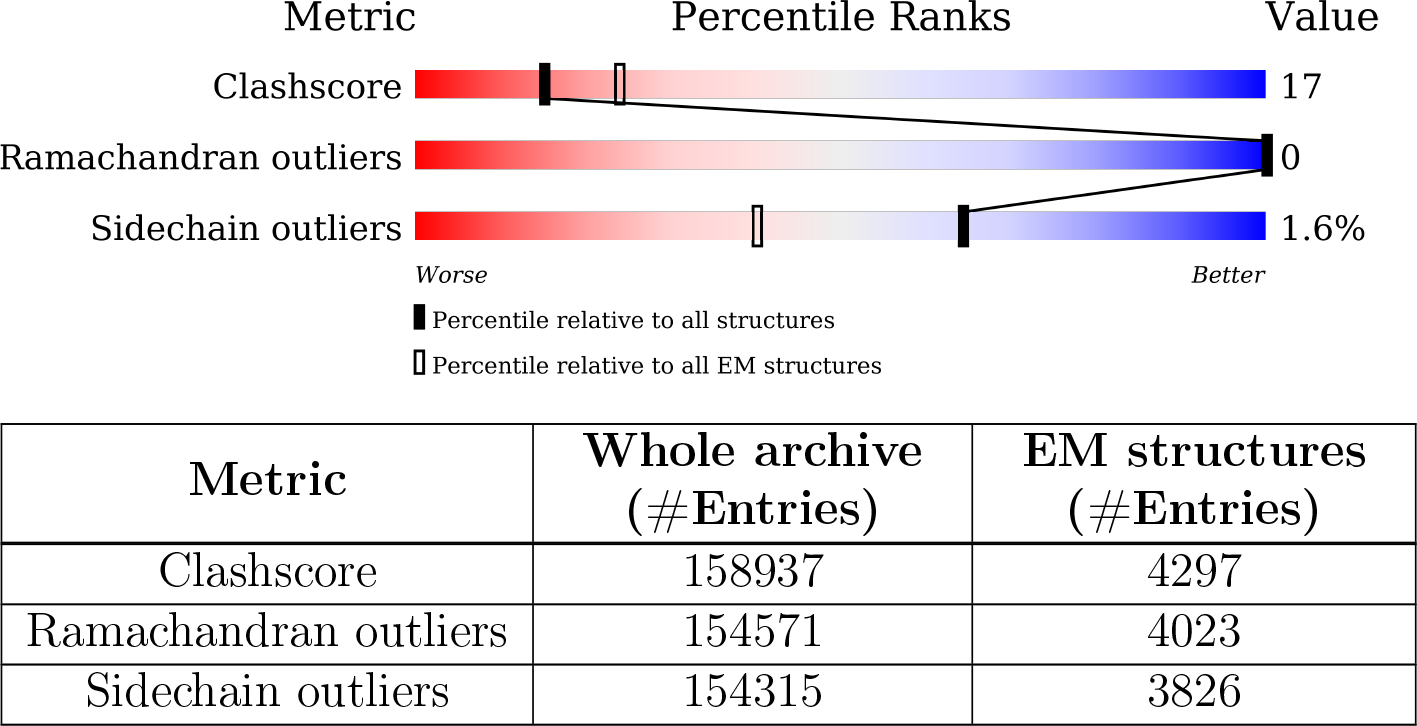

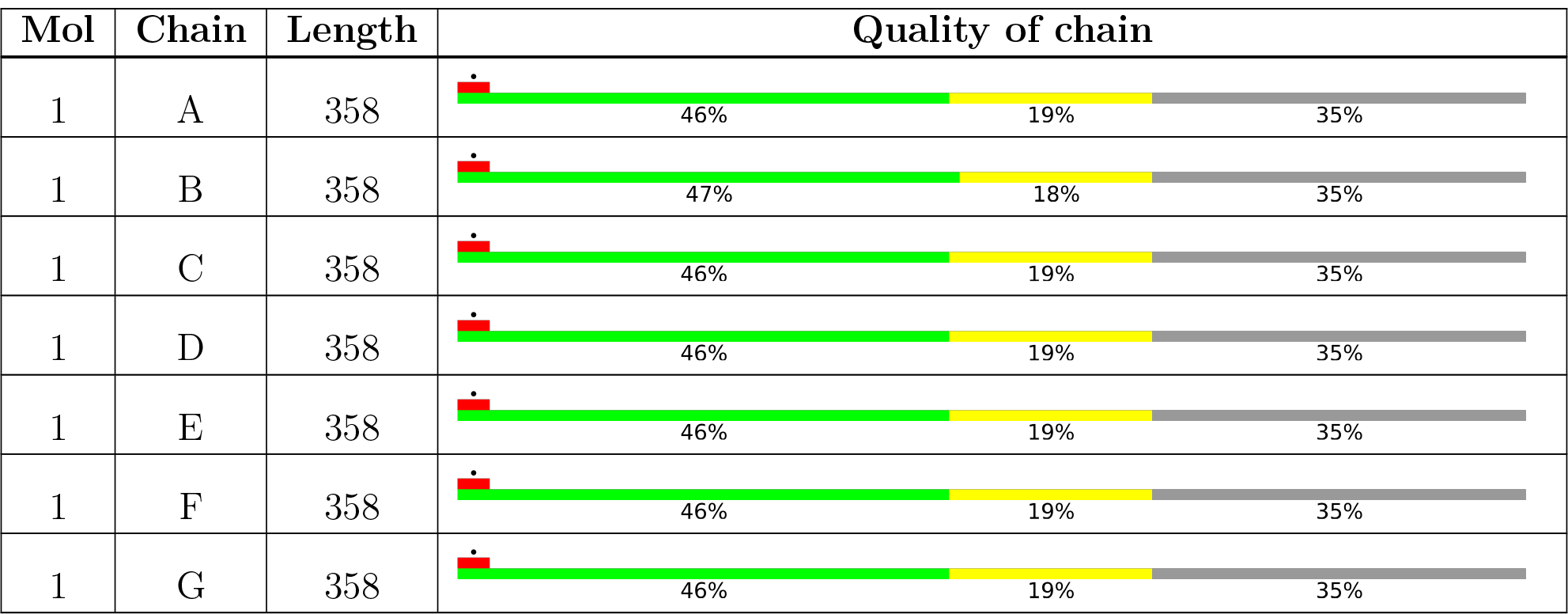

## 2 Entry composition

There are 2 unique types of molecules in this entry. The entry contains 13006 atoms, of which 0 are hydrogens and 0 are deuteriums.

• Molecule 1 is a protein called Calcium homeostasis modulator 1.

| Mol | Chain | Residues | Atoms |  |  |  |  | AltConf | Trace |
| --- | --- | --- | --- | --- | --- | --- | --- | --- | --- |
| 1 | A | 233 | Total | C | N | O | S | 0 | 0 |
|  |  |  | 1844 | 1203 | 298 | 328 | 15 |  |  |
| 1 | B | 233 | Total | C | N | O | S | 0 | 0 |
|  |  |  | 1844 | 1203 | 298 | 328 | 15 |  |  |
| 1 | C | 233 | Total | C | N | O | S | 0 | 0 |
|  |  |  | 1844 | 1203 | 298 | 328 | 15 |  |  |
| 1 | D | 233 | Total | C | N | O | S | 0 | 0 |
|  |  |  | 1844 | 1203 | 298 | 328 | 15 |  |  |
| 1 | E | 233 | Total | C | N | O | S | 0 | 0 |
|  |  |  | 1844 | 1203 | 298 | 328 | 15 |  |  |
| 1 | F | 233 | Total | C | N | O | S | 0 | 0 |
|  |  |  | 1844 | 1203 | 298 | 328 | 15 |  |  |
| 1 | G | 233 | Total | C | N | O | S | 0 | 0 |
|  |  |  | 1844 | 1203 | 298 | 328 | 15 |  |  |

There are 91 discrepancies between the modelled and reference sequences:

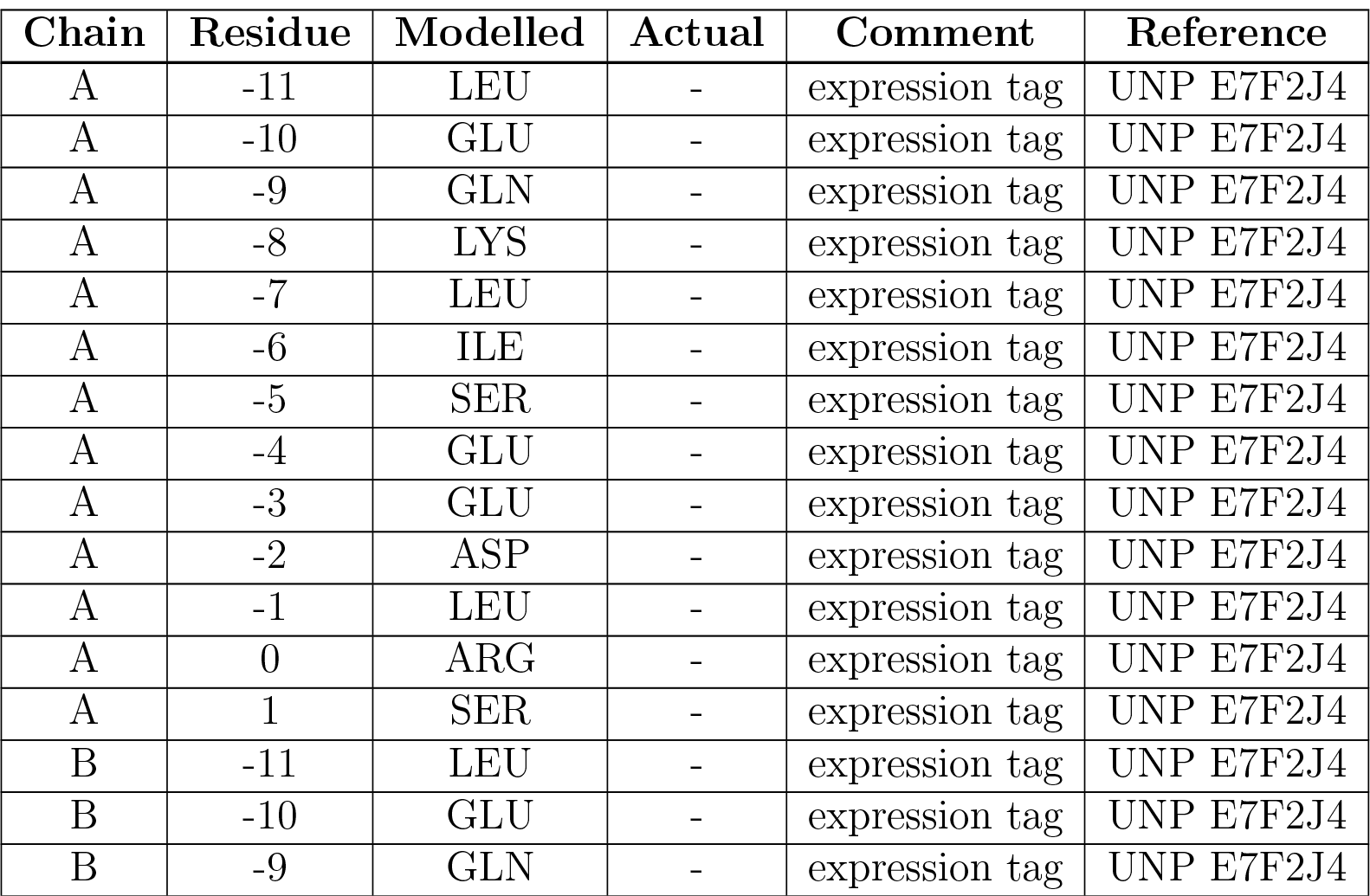
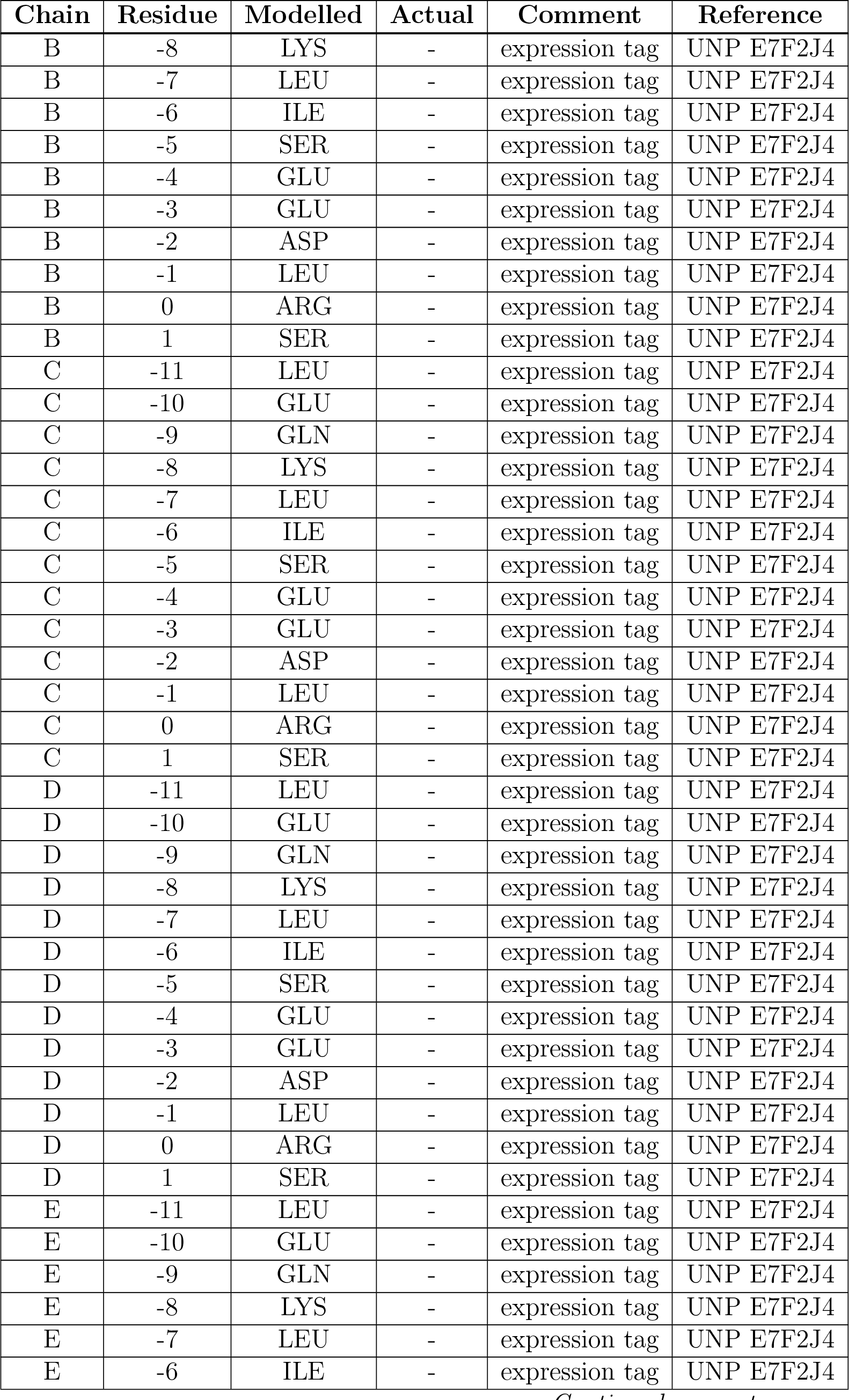
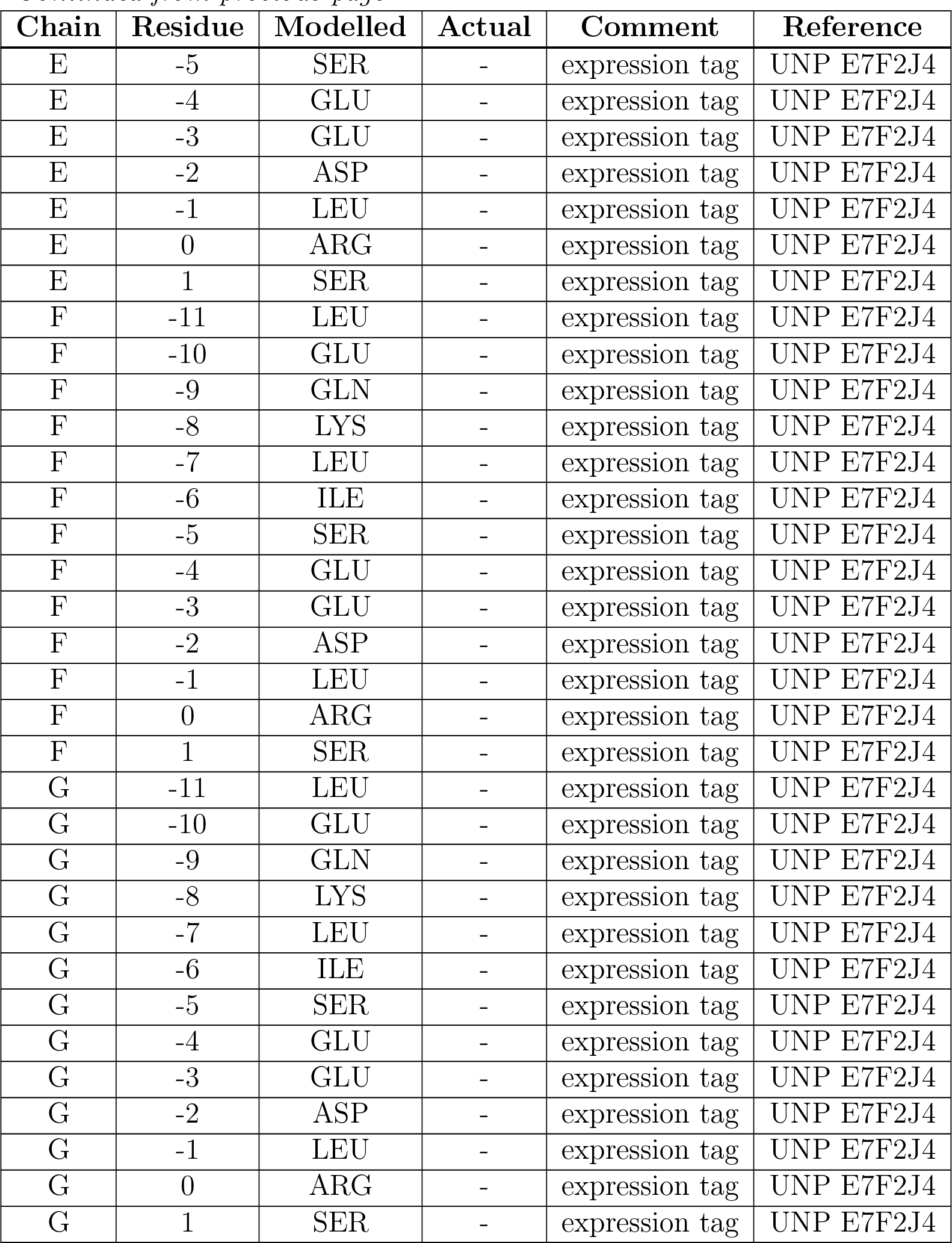

| Chain | Residue | Modelled | Actual | Comment | Reference |
| --- | --- | --- | --- | --- | --- |
| A | -11 | LEU | - | expression tag | UNP E7F2J4 |
| A | -10 | GLU | - | expression tag | UNP E7F2J4 |
| A | -9 | GLN | - | expression tag | UNP E7F2J4 |
| A | -8 | LYS | - | expression tag | UNP E7F2J4 |
| A | -7 | LEU | - | expression tag | UNP E7F2J4 |
| A | -6 | ILE | - | expression tag | UNP E7F2J4 |
| A | -5 | SER | - | expression tag | UNP E7F2J4 |
| A | -4 | GLU | - | expression tag | UNP E7F2J4 |
| A | -3 | GLU | - | expression tag | UNP E7F2J4 |
| A | -2 | ASP | - | expression tag | UNP E7F2J4 |
| A | -1 | LEU | - | expression tag | UNP E7F2J4 |
| A | 0 | ARG | - | expression tag | UNP E7F2J4 |
| A | 1 | SER | - | expression tag | UNP E7F2J4 |
| B | -11 | LEU | - | expression tag | UNP E7F2J4 |
| B | -10 | GLU | - | expression tag | UNP E7F2J4 |
| B | -9 | GLN | - | expression tag | UNP E7F2J4 |
| B | -8 | LYS | - | expression tag | UNP E7F2J4 |
| B | -7 | LEU | - | expression tag | UNP E7F2J4 |
| B | -6 | ILE | - | expression tag | UNP E7F2J4 |
| B | -5 | SER | - | expression tag | UNP E7F2J4 |
| B | -4 | GLU | - | expression tag | UNP E7F2J4 |
| B | -3 | GLU | - | expression tag | UNP E7F2J4 |
| B | -2 | ASP | - | expression tag | UNP E7F2J4 |
| B | -1 | LEU | - | expression tag | UNP E7F2J4 |
| B | 0 | ARG | - | expression tag | UNP E7F2J4 |
| B | 1 | SER | - | expression tag | UNP E7F2J4 |
| C | -11 | LEU | - | expression tag | UNP E7F2J4 |
| C | -10 | GLU | - | expression tag | UNP E7F2J4 |
| C | -9 | GLN | - | expression tag | UNP E7F2J4 |
| C | -8 | LYS | - | expression tag | UNP E7F2J4 |
| C | -7 | LEU | - | expression tag | UNP E7F2J4 |
| C | -6 | ILE | - | expression tag | UNP E7F2J4 |
| C | -5 | SER | - | expression tag | UNP E7F2J4 |
| C | -4 | GLU | - | expression tag | UNP E7F2J4 |
| C | -3 | GLU | - | expression tag | UNP E7F2J4 |
| C | -2 | ASP | - | expression tag | UNP E7F2J4 |
| C | -1 | LEU | - | expression tag | UNP E7F2J4 |
| C | 0 | ARG | - | expression tag | UNP E7F2J4 |
| C | 1 | SER | - | expression tag | UNP E7F2J4 |
| D | -11 | LEU | - | expression tag | UNP E7F2J4 |
| D | -10 | GLU | - | expression tag | UNP E7F2J4 |
| D | -9 | GLN | - | expression tag | UNP E7F2J4 |
| D | -8 | LYS | - | expression tag | UNP E7F2J4 |
| D | -7 | LEU | - | expression tag | UNP E7F2J4 |
| D | -6 | ILE | - | expression tag | UNP E7F2J4 |
| D | -5 | SER | - | expression tag | UNP E7F2J4 |
| D | -4 | GLU | - | expression tag | UNP E7F2J4 |
| D | -3 | GLU | - | expression tag | UNP E7F2J4 |
| D | -2 | ASP | - | expression tag | UNP E7F2J4 |
| D | -1 | LEU | - | expression tag | UNP E7F2J4 |
| D | 0 | ARG | - | expression tag | UNP E7F2J4 |
| D | 1 | SER | - | expression tag | UNP E7F2J4 |
| E | -11 | LEU | - | expression tag | UNP E7F2J4 |
| E | -10 | GLU | - | expression tag | UNP E7F2J4 |
| E | -9 | GLN | - | expression tag | UNP E7F2J4 |
| E | -8 | LYS | - | expression tag | UNP E7F2J4 |
| E | -7 | LEU | - | expression tag | UNP E7F2J4 |
| E | -6 | ILE | - | expression tag | UNP E7F2J4 |
| E | -5 | SER | - | expression tag | UNP E7F2J4 |
| E | -4 | GLU | - | expression tag | UNP E7F2J4 |
| E | -3 | GLU | - | expression tag | UNP E7F2J4 |
| E | -2 | ASP | - | expression tag | UNP E7F2J4 |
| E | -1 | LEU | - | expression tag | UNP E7F2J4 |
| E | 0 | ARG | - | expression tag | UNP E7F2J4 |
| E | 1 | SER | - | expression tag | UNP E7F2J4 |
| F | -11 | LEU | - | expression tag | UNP E7F2J4 |
| F | -10 | GLU | - | expression tag | UNP E7F2J4 |
| F | -9 | GLN | - | expression tag | UNP E7F2J4 |
| F | -8 | LYS | - | expression tag | UNP E7F2J4 |
| F | -7 | LEU | - | expression tag | UNP E7F2J4 |
| F | -6 | ILE | - | expression tag | UNP E7F2J4 |
| F | -5 | SER | - | expression tag | UNP E7F2J4 |
| F | -4 | GLU | - | expression tag | UNP E7F2J4 |
| F | -3 | GLU | - | expression tag | UNP E7F2J4 |
| F | -2 | ASP | - | expression tag | UNP E7F2J4 |
| F | -1 | LEU | - | expression tag | UNP E7F2J4 |
| F | 0 | ARG | - | expression tag | UNP E7F2J4 |
| F | 1 | SER | - | expression tag | UNP E7F2J4 |
| G | -11 | LEU | - | expression tag | UNP E7F2J4 |
| G | -10 | GLU | - | expression tag | UNP E7F2J4 |
| G | -9 | GLN | - | expression tag | UNP E7F2J4 |
| G | -8 | LYS | - | expression tag | UNP E7F2J4 |
| G | -7 | LEU | - | expression tag | UNP E7F2J4 |
| G | -6 | ILE | - | expression tag | UNP E7F2J4 |
| G | -5 | SER | - | expression tag | UNP E7F2J4 |
| G | -4 | GLU | - | expression tag | UNP E7F2J4 |
| G | -3 | GLU | - | expression tag | UNP E7F2J4 |
| G | -2 | ASP | - | expression tag | UNP E7F2J4 |
| G | -1 | LEU | - | expression tag | UNP E7F2J4 |
| G | 0 | ARG | - | expression tag | UNP E7F2J4 |
| G | 1 | SER | - | expression tag | UNP E7F2J4 |

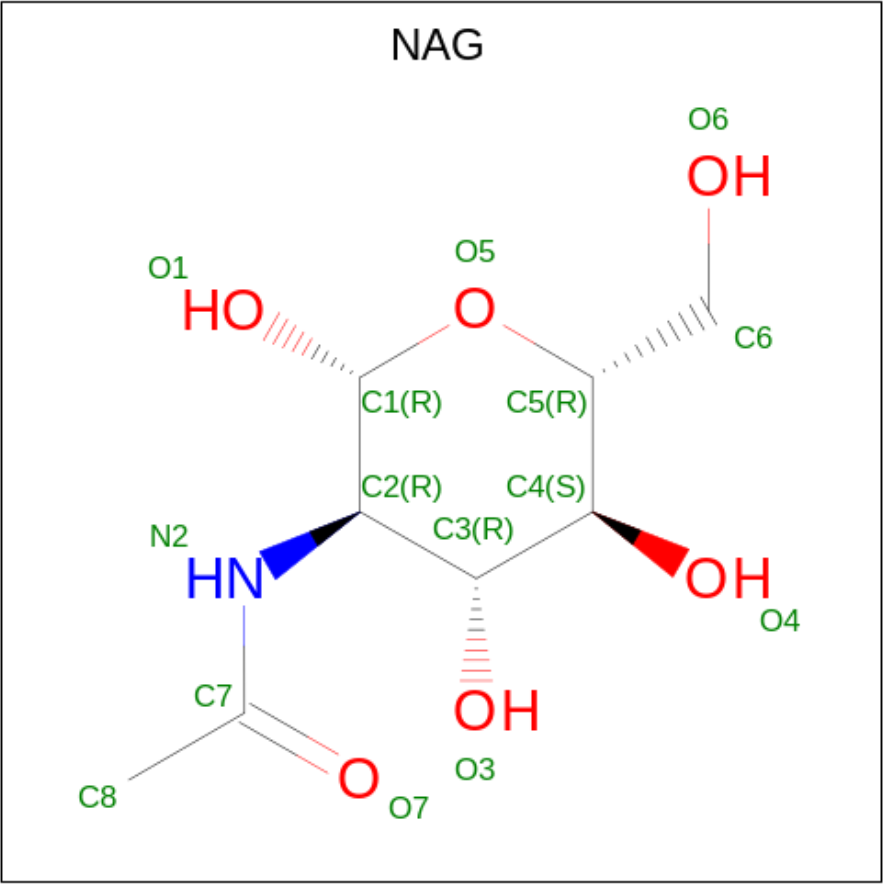

| Mol | Chain | Residues | Atoms |  |  |  | AltConf |
| --- | --- | --- | --- | --- | --- | --- | --- |
|  |  |  | Total | C | N | O |  |
| 2 | A | 1 | Total<br>14 | 8 | 1 | 5 | 0 |
| 2 | B | 1 | Total<br>14 | 8 | 1 | 5 | 0 |
| 2 | C | 1 | Total<br>14 | 8 | 1 | 5 | 0 |
| 2 | D | 1 | Total<br>14 | 8 | 1 | 5 | 0 |
| 2 | E | 1 | Total<br>14 | 8 | 1 | 5 | 0 |
| 2 | F | 1 | Total<br>14 | 8 | 1 | 5 | 0 |
| 2 | G | 1 | Total<br>14 | 8 | 1 | 5 | 0 |

## 3 Residue-property plots

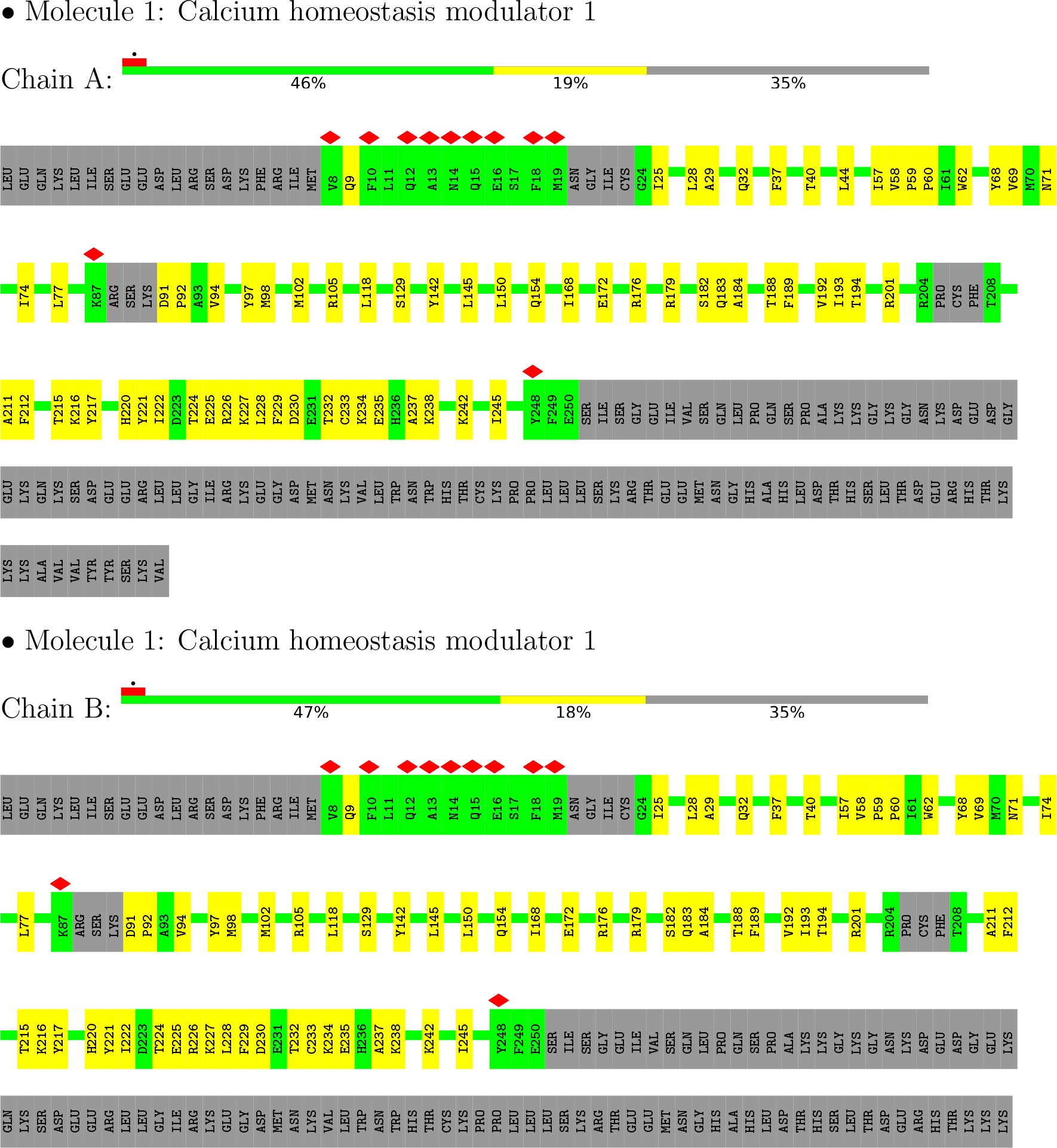
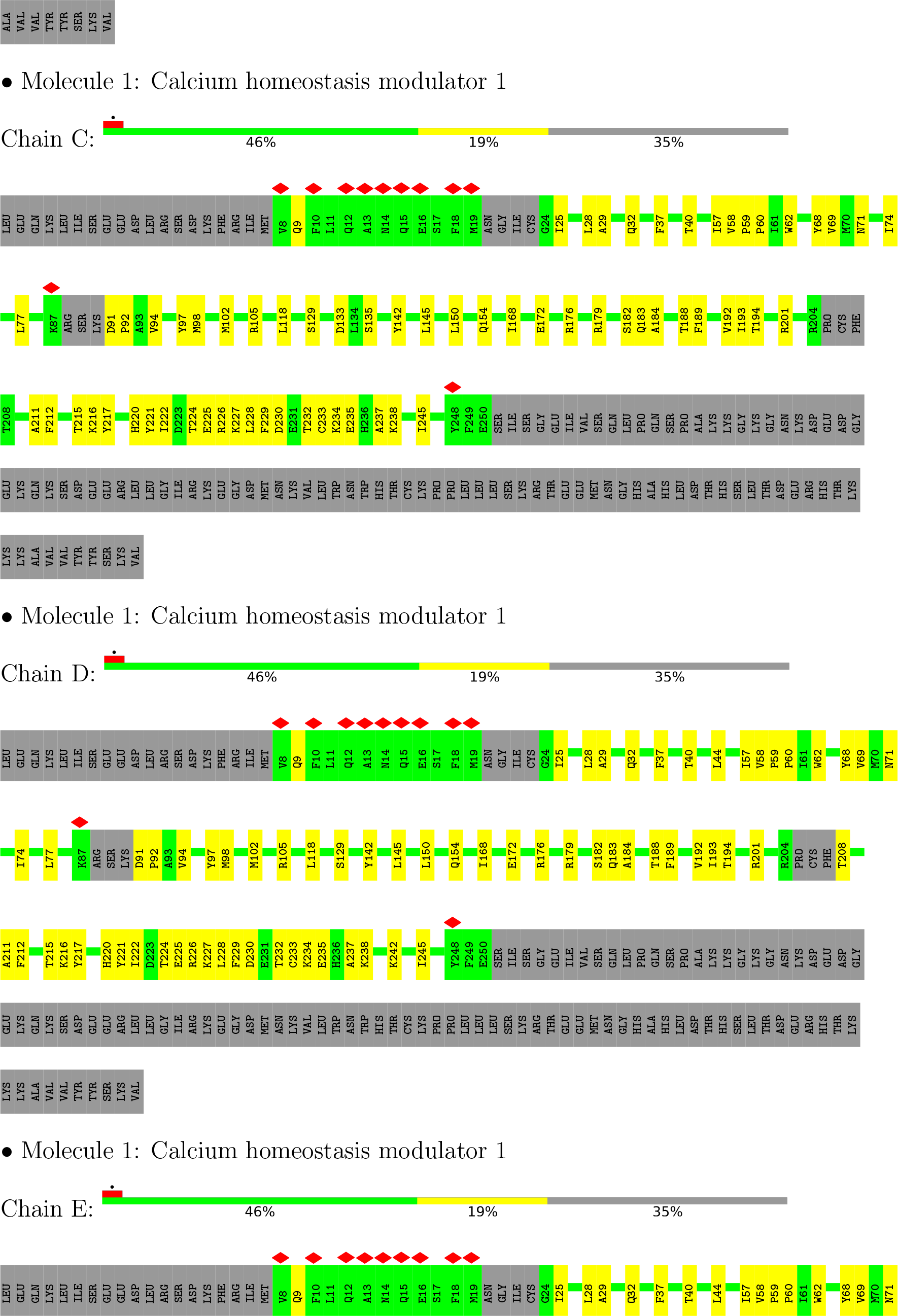
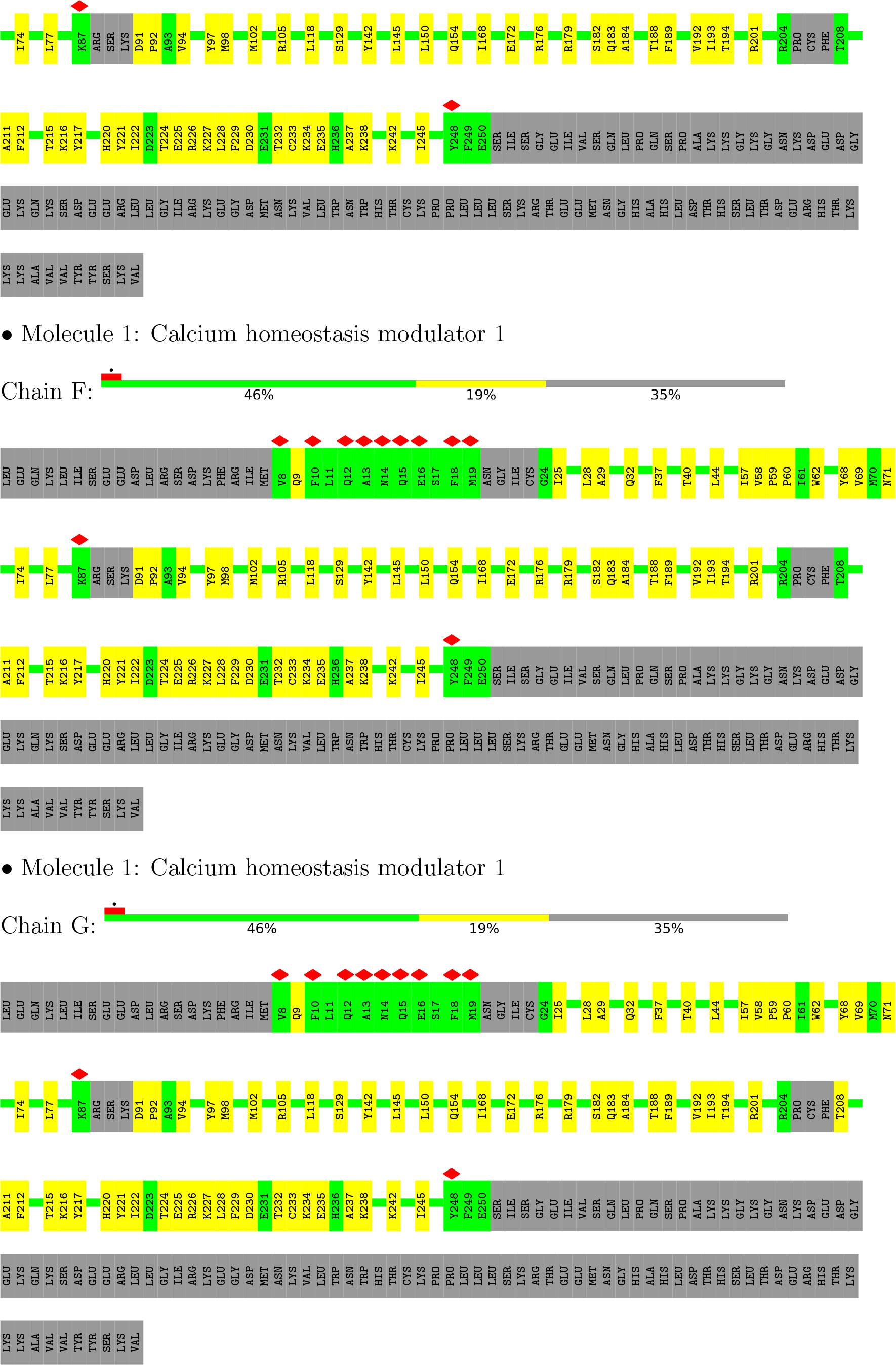

## 4 Experimental information

| Property | Value | Source |
| --- | --- | --- |
| EM reconstruction method | SINGLE PARTICLE | Depositor |
| Imposed symmetry | POINT, Not provided |  |
| Number of particles used | 12671 | Depositor |
| Resolution determination method | FSC 0.143 CUT-OFF | Depositor |
| CTF correction method | NONE | Depositor |
| Microscope | FEI TITAN KRIOS | Depositor |
| Voltage (kV) | 300 | Depositor |
| Electron dose ($e^-/\text{\AA}^2$) | 50 | Depositor |
| Minimum defocus (nm) | Not provided |  |
| Maximum defocus (nm) | Not provided |  |
| Magnification | Not provided |  |
| Image detector | GATAN K2 SUMMIT (4k x 4k) | Depositor |
| Maximum map value | 1.569 | Depositor |
| Minimum map value | -0.651 | Depositor |
| Average map value | -0.002 | Depositor |
| Map value standard deviation | 0.057 | Depositor |
| Recommended contour level | 0.209 | Depositor |
| Map size (Å) | 263.64, 263.64, 263.64 | wwPDB |
| Map dimensions | 260, 260, 260 | wwPDB |
| Map angles (°) | 90.0, 90.0, 90.0 | wwPDB |
| Pixel spacing (Å) | 1.014, 1.014, 1.014 | Depositor |

## 5 Model quality

### 5.1 Standard geometry

Bond lengths and bond angles in the following residue types are not validated in this section: NAG

| Mol | Chain | Bond lengths |  | Bond angles |  |
| --- | --- | --- | --- | --- | --- |
|  |  | RMSZ | #\|Z\| >5 | RMSZ | #\|Z\| >5 |
| 1 | A | 0.31 | 0/1888 | 0.41 | 0/2557 |
| 1 | B | 0.31 | 0/1888 | 0.41 | 0/2557 |
| 1 | C | 0.31 | 0/1888 | 0.41 | 0/2557 |
| 1 | D | 0.31 | 0/1888 | 0.41 | 0/2557 |
| 1 | E | 0.31 | 0/1888 | 0.41 | 0/2557 |
| 1 | F | 0.31 | 0/1888 | 0.41 | 0/2557 |
| 1 | G | 0.31 | 0/1888 | 0.41 | 0/2557 |
| All | All | 0.31 | 0/13216 | 0.41 | 0/17899 |

There are no bond length outliers.

There are no bond angle outliers.

There are no chirality outliers.

There are no planarity outliers.

### 5.2 Too-close contacts

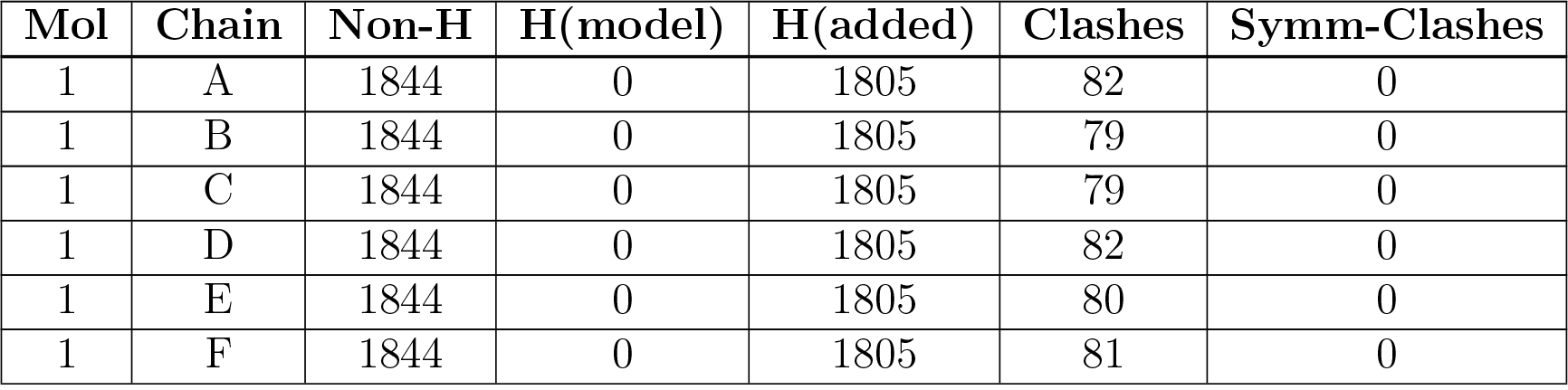
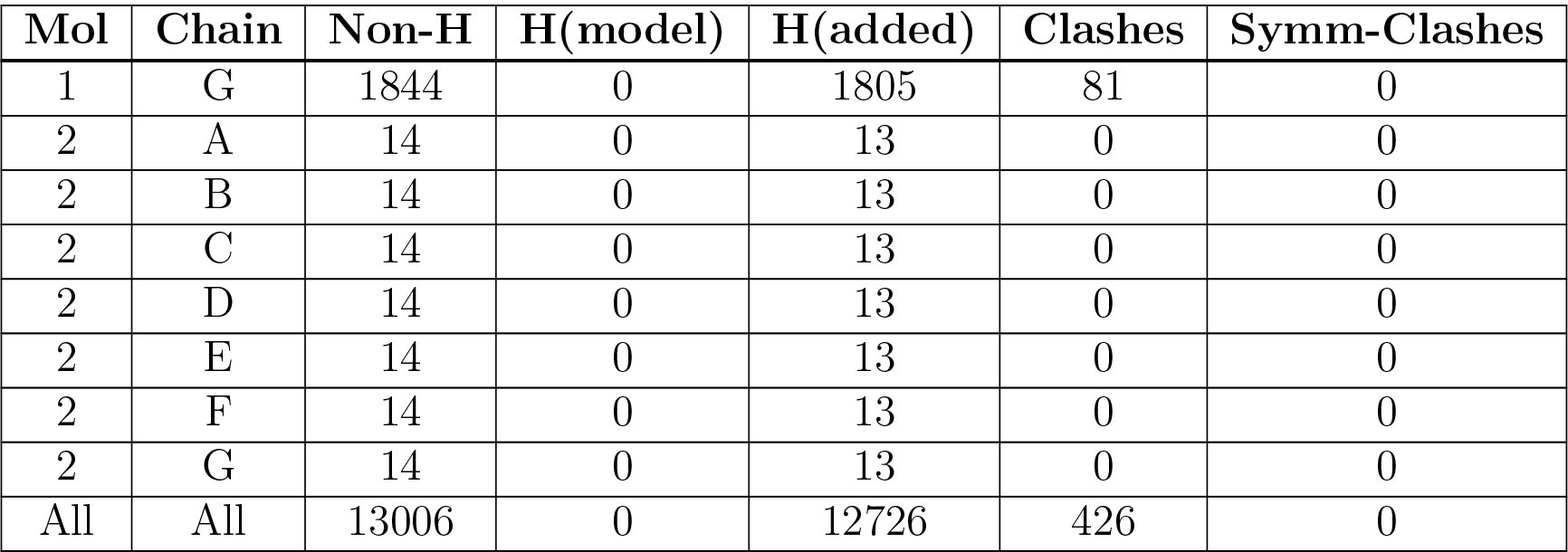

| Mol | Chain | Non-H | H(model) | H(added) | Clashes | Symm-Clashes |
| --- | --- | --- | --- | --- | --- | --- |
| 1 | A | 1844 | 0 | 1805 | 82 | 0 |
| 1 | B | 1844 | 0 | 1805 | 79 | 0 |
| 1 | C | 1844 | 0 | 1805 | 79 | 0 |
| 1 | D | 1844 | 0 | 1805 | 82 | 0 |
| 1 | E | 1844 | 0 | 1805 | 80 | 0 |
| 1 | F | 1844 | 0 | 1805 | 81 | 0 |
| 1 | G | 1844 | 0 | 1805 | 81 | 0 |
| 2 | A | 14 | 0 | 13 | 0 | 0 |
| 2 | B | 14 | 0 | 13 | 0 | 0 |
| 2 | C | 14 | 0 | 13 | 0 | 0 |
| 2 | D | 14 | 0 | 13 | 0 | 0 |
| 2 | E | 14 | 0 | 13 | 0 | 0 |
| 2 | F | 14 | 0 | 13 | 0 | 0 |
| 2 | G | 14 | 0 | 13 | 0 | 0 |
| All | All | 13006 | 0 | 12726 | 426 | 0 |

The all-atom clashscore is defined as the number of clashes found per 1000 atoms (including hydrogen atoms). The all-atom clashscore for this structure is 17.

All (426) close contacts within the same asymmetric unit are listed below, sorted by their clash magnitude.

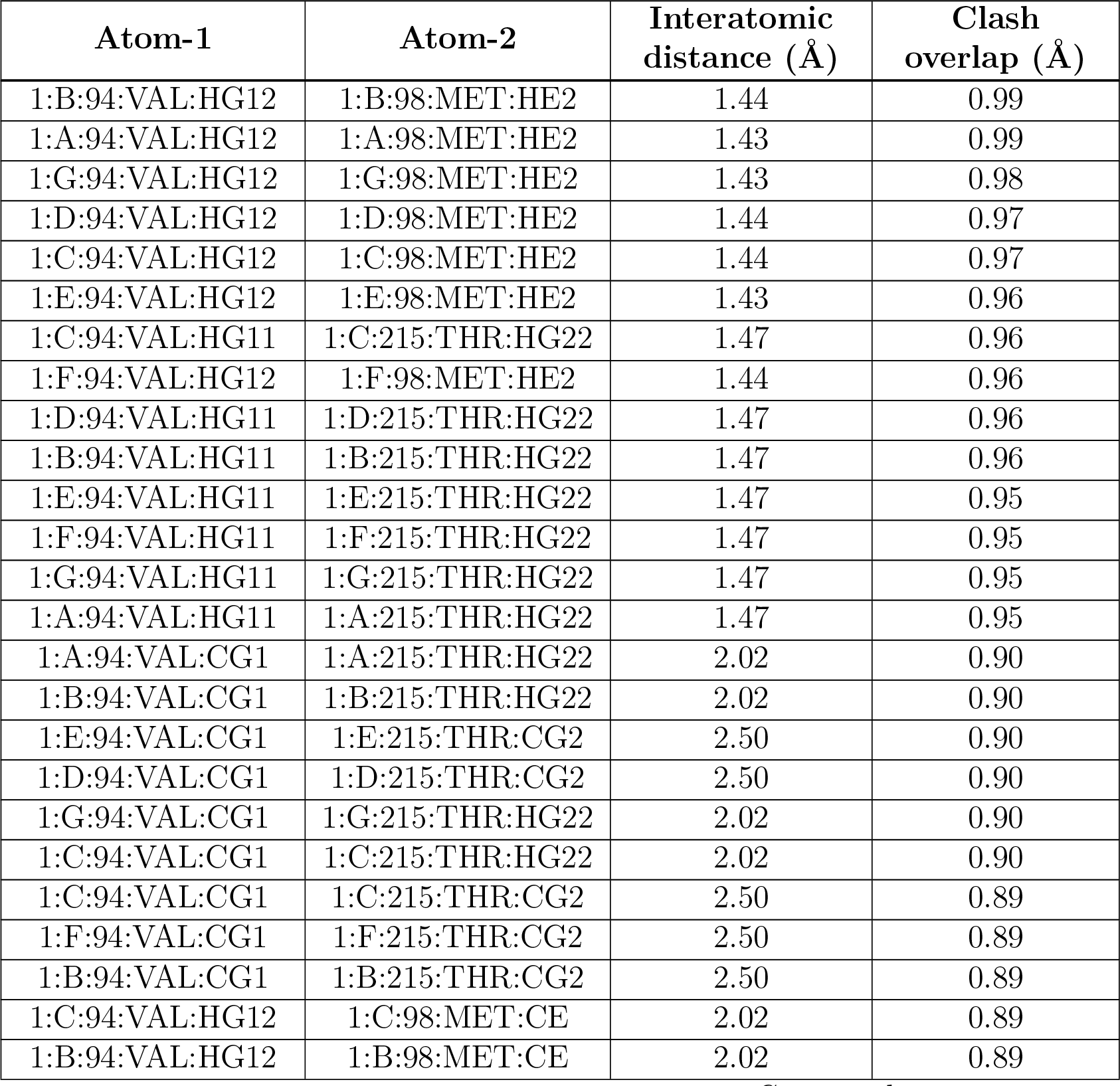
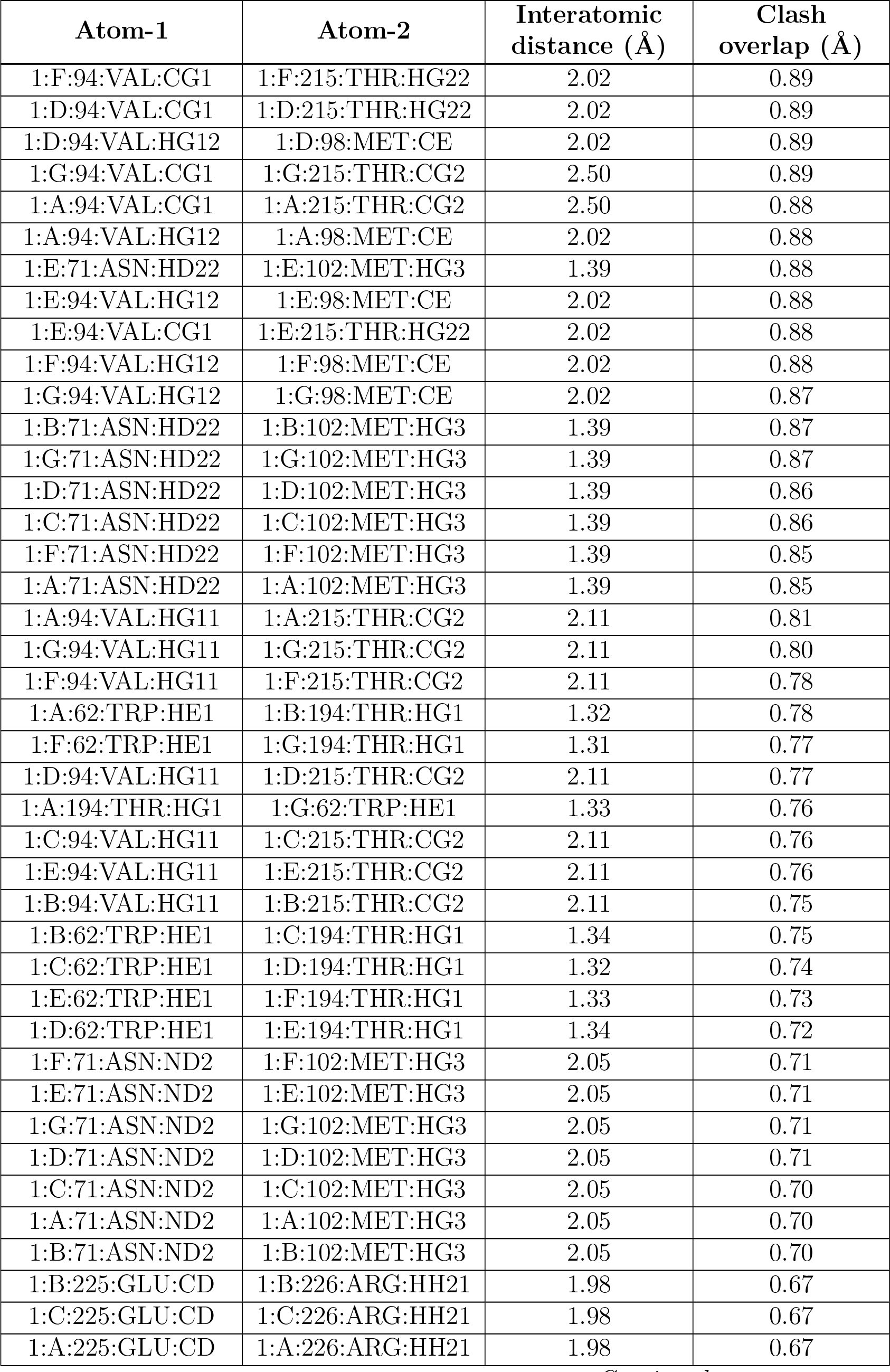
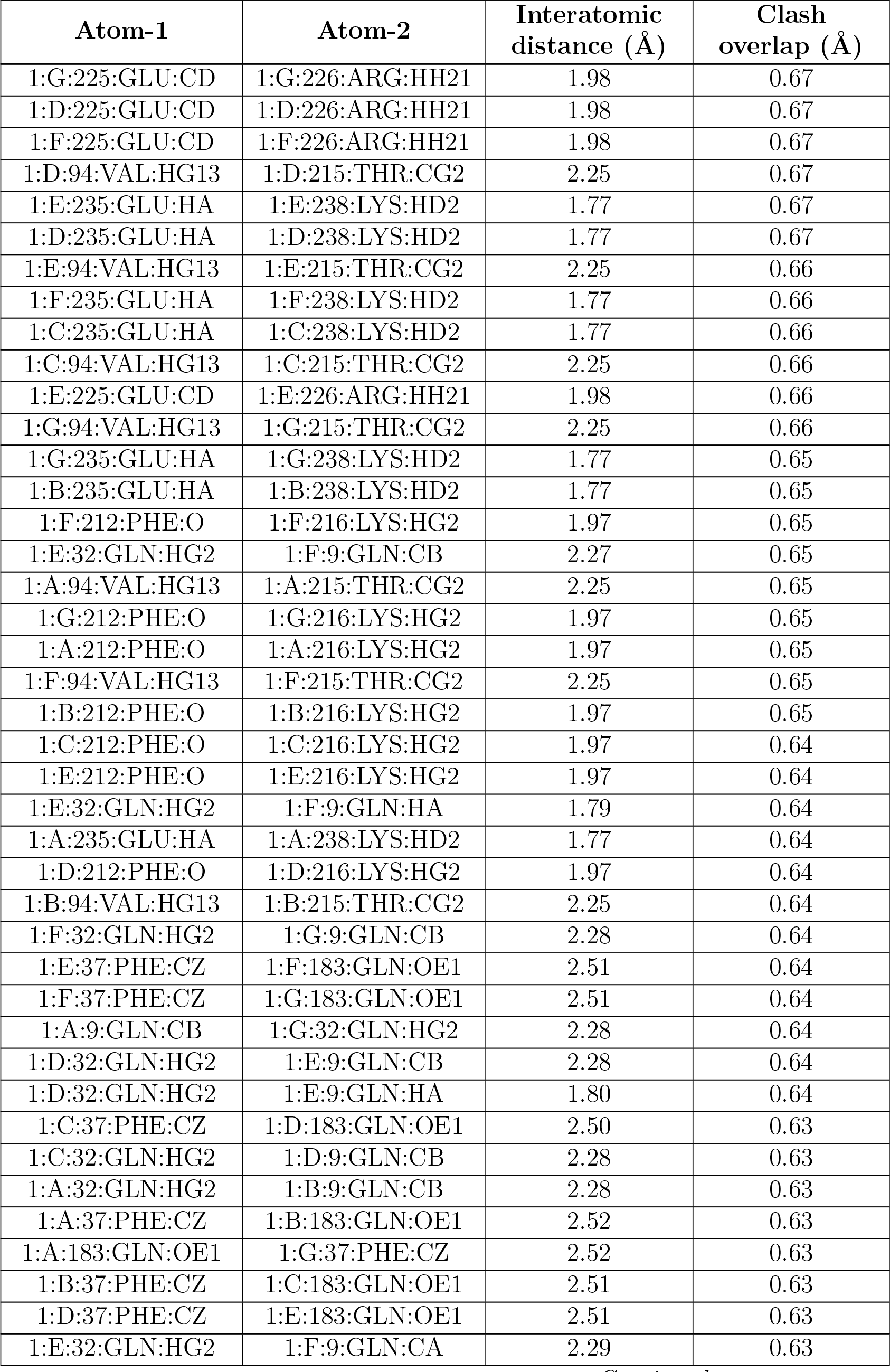
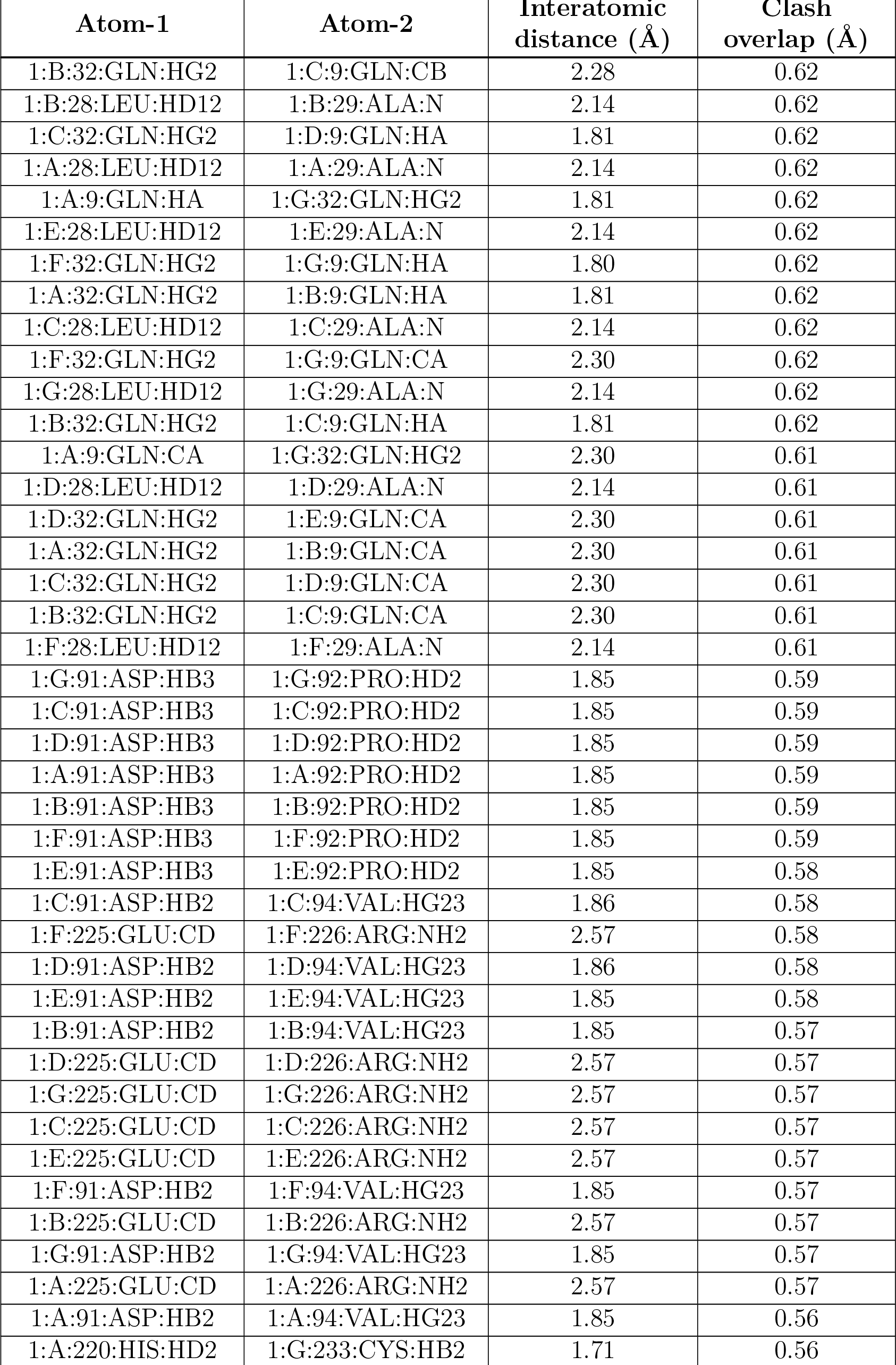
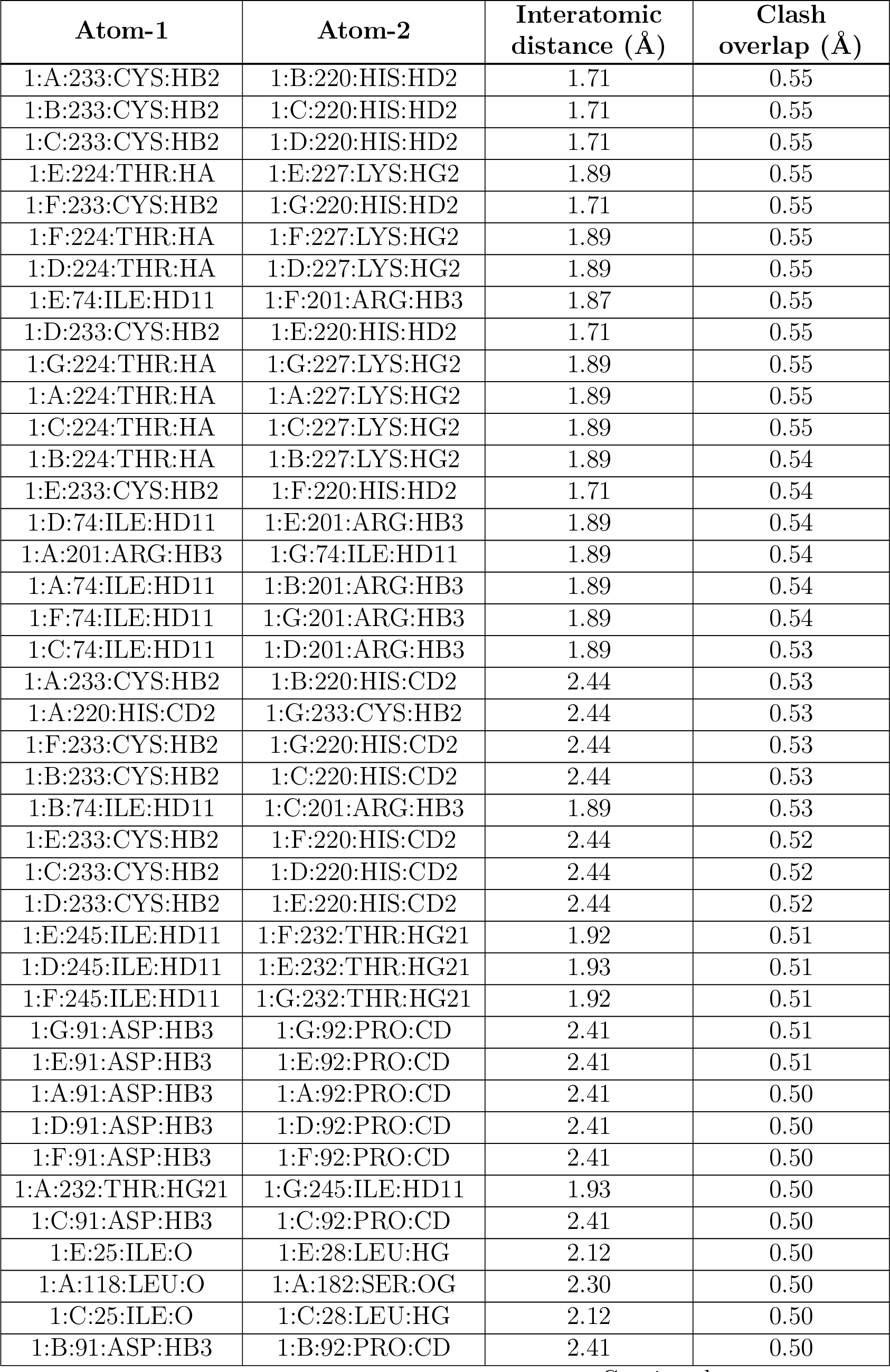
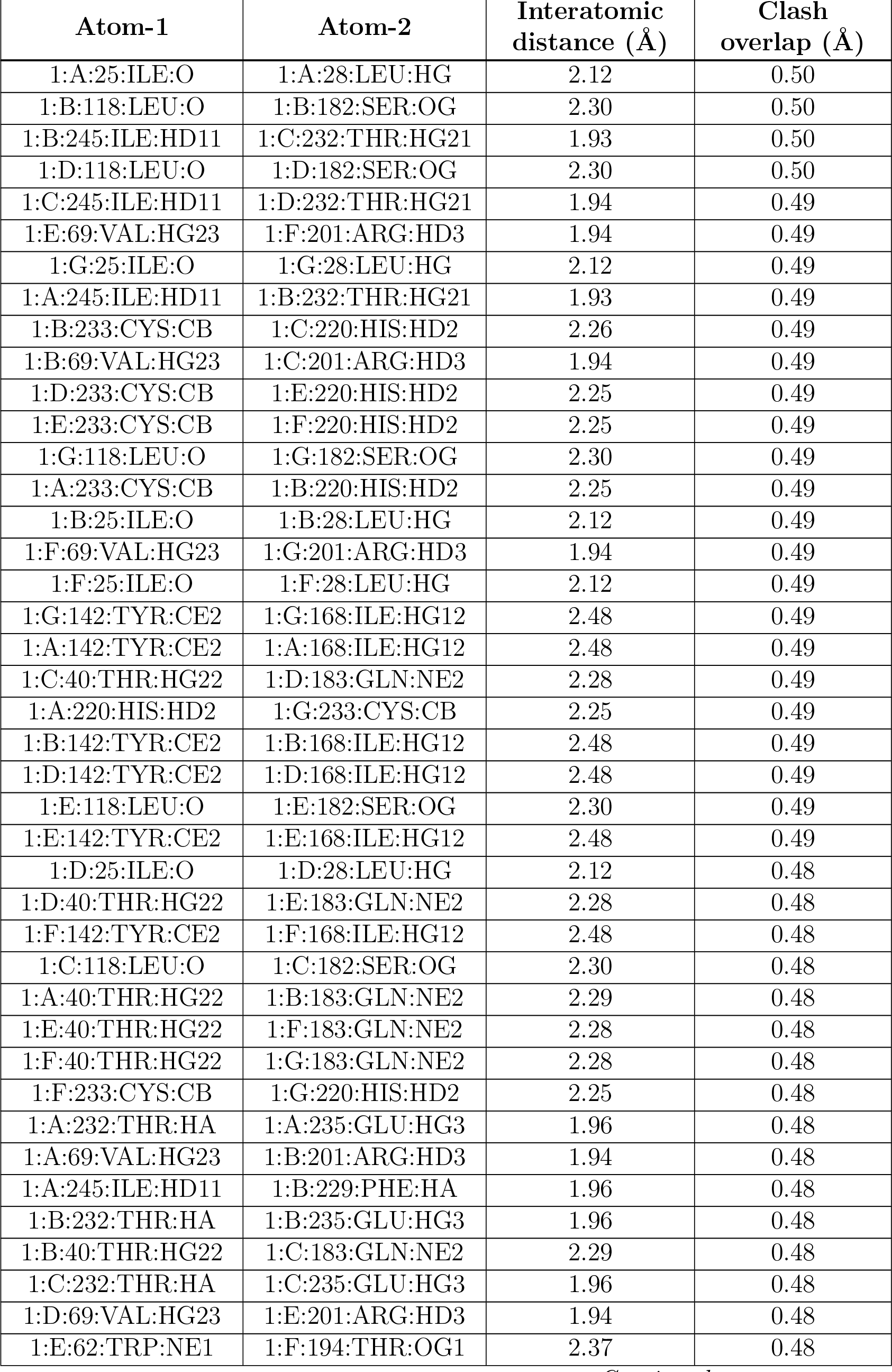
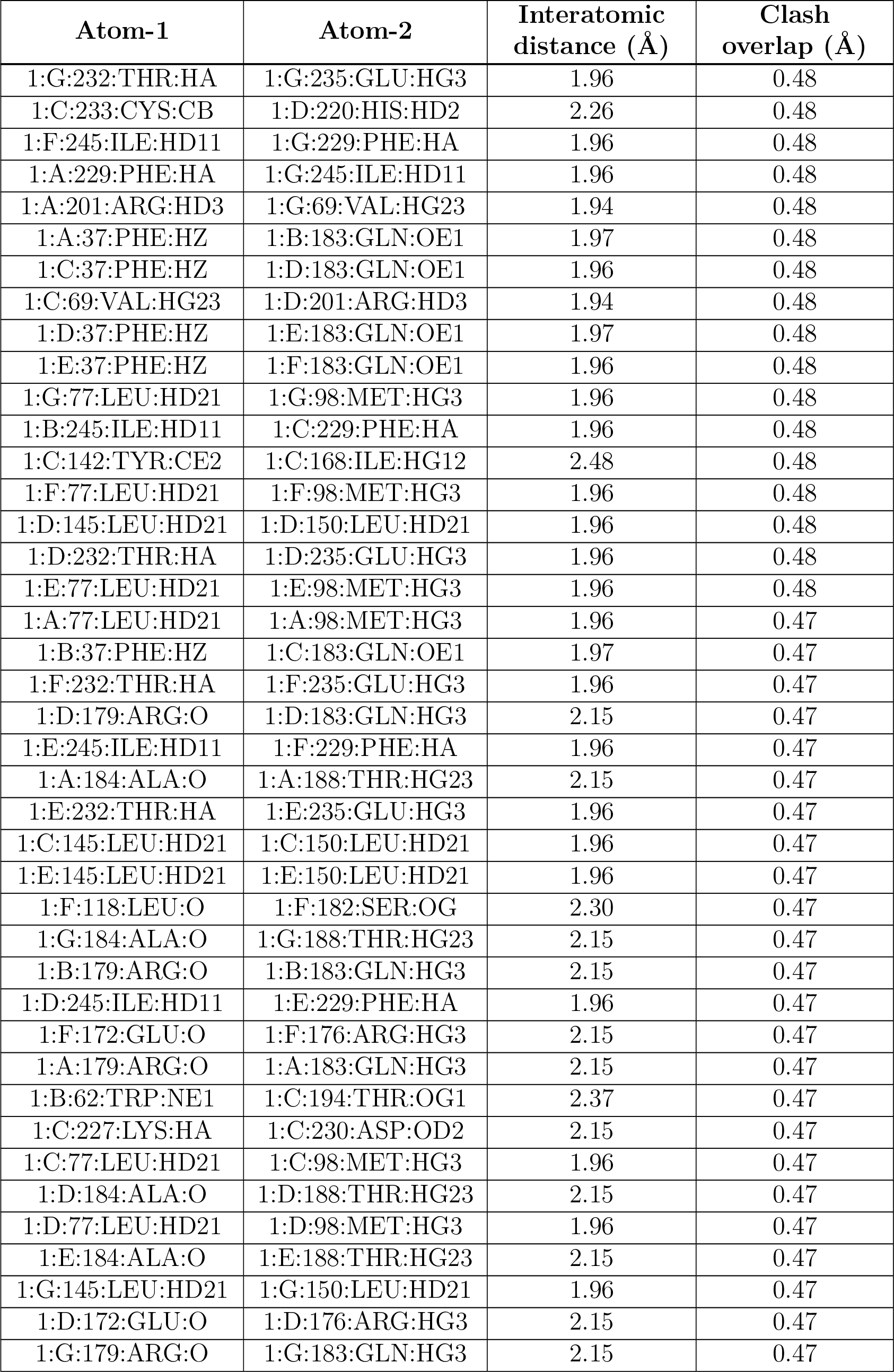
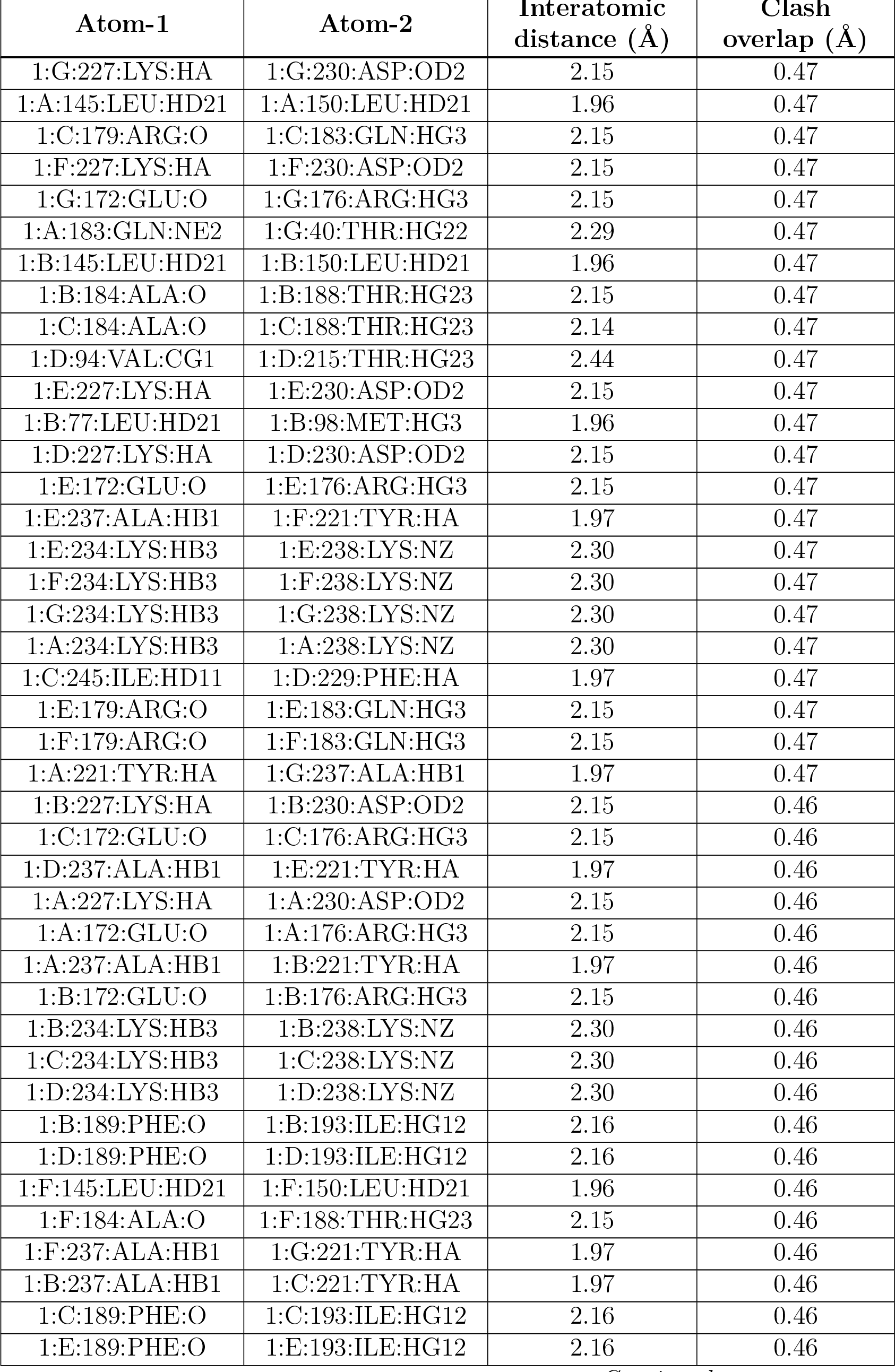
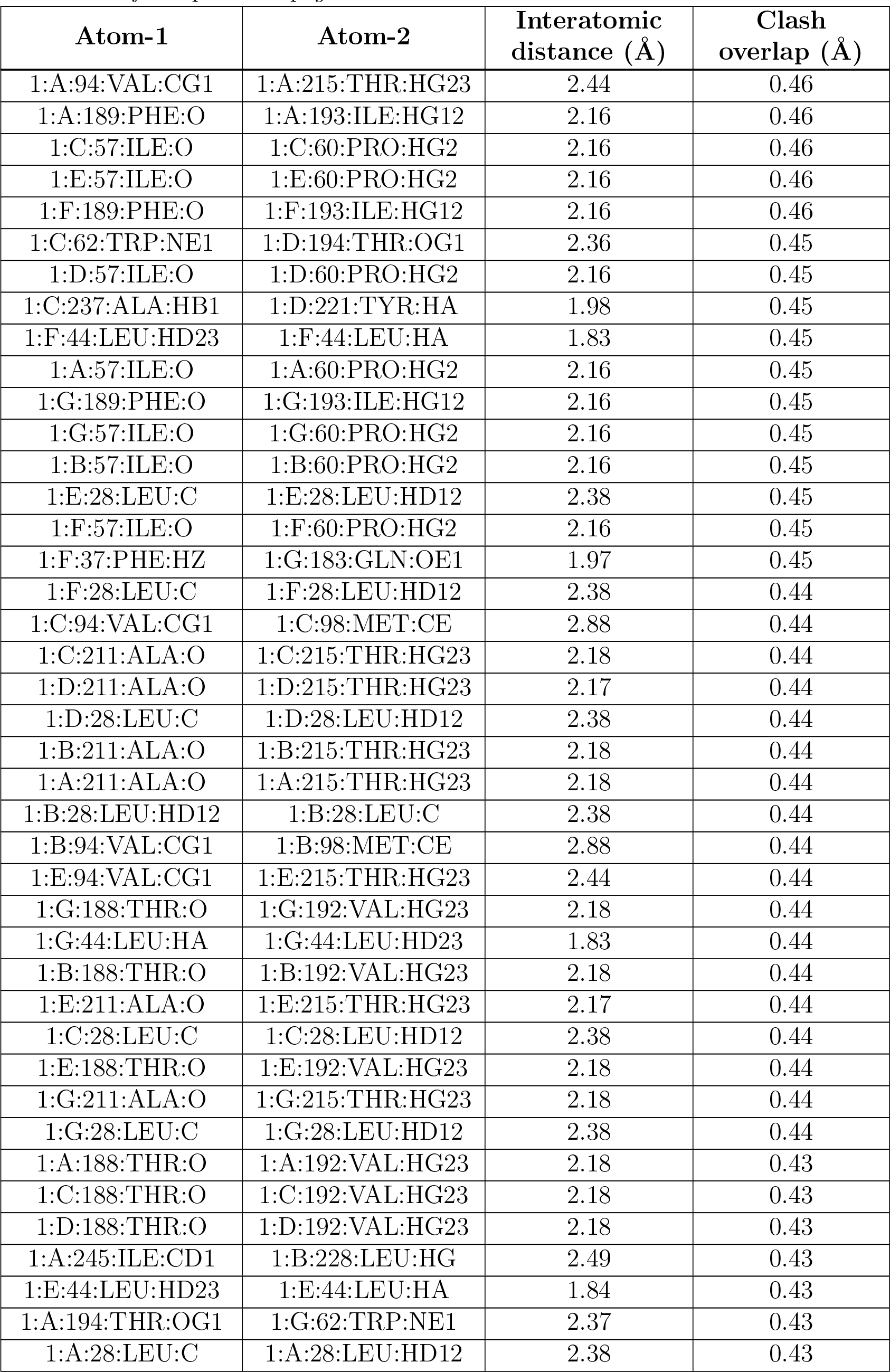
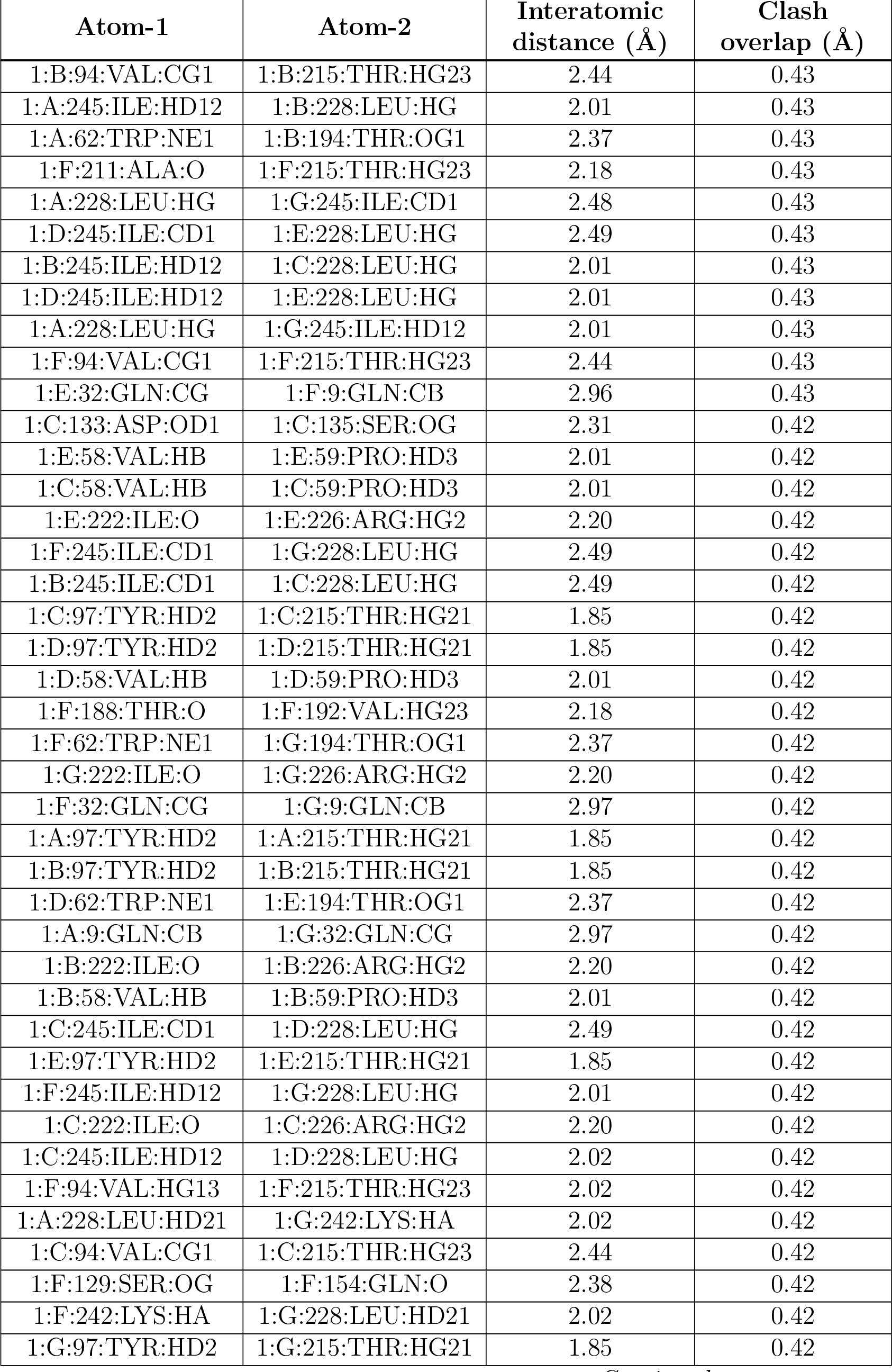
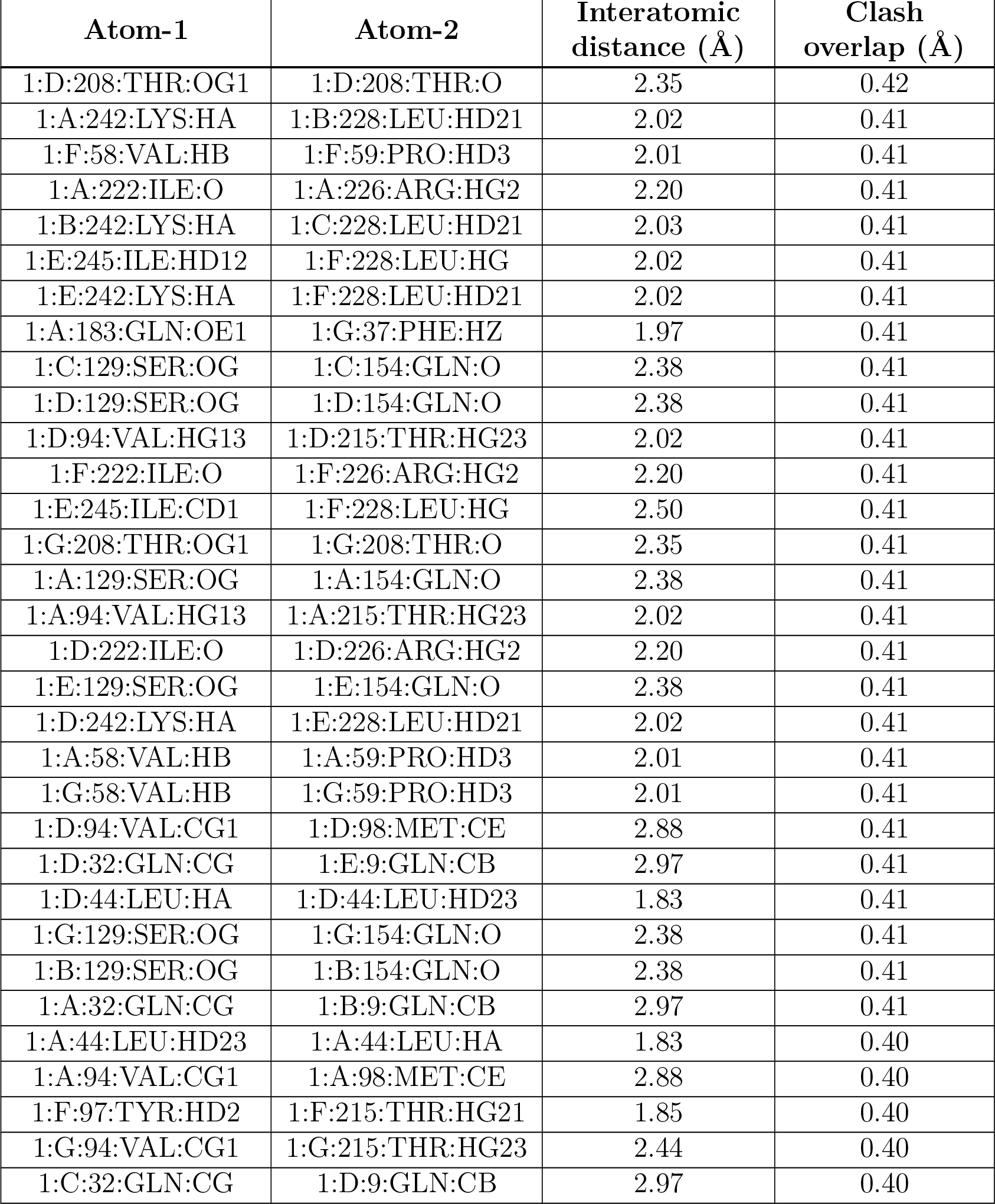

| Atom-1 | Atom-2 | Interatomic distance (Å) | Clash overlap (Å) |
| --- | --- | --- | --- |
| 1:B:94:VAL:HG12 | 1:B:98:MET:HE2 | 1.44 | 0.99 |
| 1:A:94:VAL:HG12 | 1:A:98:MET:HE2 | 1.43 | 0.99 |
| 1:G:94:VAL:HG12 | 1:G:98:MET:HE2 | 1.43 | 0.98 |
| 1:D:94:VAL:HG12 | 1:D:98:MET:HE2 | 1.44 | 0.97 |
| 1:C:94:VAL:HG12 | 1:C:98:MET:HE2 | 1.44 | 0.97 |
| 1:E:94:VAL:HG12 | 1:E:98:MET:HE2 | 1.43 | 0.96 |
| 1:C:94:VAL:HG11 | 1:C:215:THR:HG22 | 1.47 | 0.96 |
| 1:F:94:VAL:HG12 | 1:F:98:MET:HE2 | 1.44 | 0.96 |
| 1:D:94:VAL:HG11 | 1:D:215:THR:HG22 | 1.47 | 0.96 |
| 1:B:94:VAL:HG11 | 1:B:215:THR:HG22 | 1.47 | 0.96 |
| 1:E:94:VAL:HG11 | 1:E:215:THR:HG22 | 1.47 | 0.95 |
| 1:F:94:VAL:HG11 | 1:F:215:THR:HG22 | 1.47 | 0.95 |
| 1:G:94:VAL:HG11 | 1:G:215:THR:HG22 | 1.47 | 0.95 |
| 1:A:94:VAL:HG11 | 1:A:215:THR:HG22 | 1.47 | 0.95 |
| 1:A:94:VAL:CG1 | 1:A:215:THR:HG22 | 2.02 | 0.90 |
| 1:B:94:VAL:CG1 | 1:B:215:THR:HG22 | 2.02 | 0.90 |
| 1:E:94:VAL:CG1 | 1:E:215:THR:CG2 | 2.50 | 0.90 |
| 1:D:94:VAL:CG1 | 1:D:215:THR:CG2 | 2.50 | 0.90 |
| 1:G:94:VAL:CG1 | 1:G:215:THR:HG22 | 2.02 | 0.90 |
| 1:C:94:VAL:CG1 | 1:C:215:THR:HG22 | 2.02 | 0.90 |
| 1:C:94:VAL:CG1 | 1:C:215:THR:CG2 | 2.50 | 0.89 |
| 1:F:94:VAL:CG1 | 1:F:215:THR:CG2 | 2.50 | 0.89 |
| 1:B:94:VAL:CG1 | 1:B:215:THR:CG2 | 2.50 | 0.89 |
| 1:C:94:VAL:HG12 | 1:C:98:MET:CE | 2.02 | 0.89 |
| 1:B:94:VAL:HG12 | 1:B:98:MET:CE | 2.02 | 0.89 |
| 1:F:94:VAL:CG1 | 1:F:215:THR:HG22 | 2.02 | 0.89 |
| 1:D:94:VAL:CG1 | 1:D:215:THR:HG22 | 2.02 | 0.89 |
| 1:D:94:VAL:HG12 | 1:D:98:MET:CE | 2.02 | 0.89 |
| 1:G:94:VAL:CG1 | 1:G:215:THR:CG2 | 2.50 | 0.89 |
| 1:A:94:VAL:CG1 | 1:A:215:THR:CG2 | 2.50 | 0.88 |
| 1:A:94:VAL:HG12 | 1:A:98:MET:CE | 2.02 | 0.88 |
| 1:E:71:ASN:HD22 | 1:E:102:MET:HG3 | 1.39 | 0.88 |
| 1:E:94:VAL:HG12 | 1:E:98:MET:CE | 2.02 | 0.88 |
| 1:E:94:VAL:CG1 | 1:E:215:THR:HG22 | 2.02 | 0.88 |
| 1:F:94:VAL:HG12 | 1:F:98:MET:CE | 2.02 | 0.88 |
| 1:G:94:VAL:HG12 | 1:G:98:MET:CE | 2.02 | 0.87 |
| 1:B:71:ASN:HD22 | 1:B:102:MET:HG3 | 1.39 | 0.87 |
| 1:G:71:ASN:HD22 | 1:G:102:MET:HG3 | 1.39 | 0.87 |
| 1:D:71:ASN:HD22 | 1:D:102:MET:HG3 | 1.39 | 0.86 |
| 1:C:71:ASN:HD22 | 1:C:102:MET:HG3 | 1.39 | 0.86 |
| 1:F:71:ASN:HD22 | 1:F:102:MET:HG3 | 1.39 | 0.85 |
| 1:A:71:ASN:HD22 | 1:A:102:MET:HG3 | 1.39 | 0.85 |
| 1:A:94:VAL:HG11 | 1:A:215:THR:CG2 | 2.11 | 0.81 |
| 1:G:94:VAL:HG11 | 1:G:215:THR:CG2 | 2.11 | 0.80 |
| 1:F:94:VAL:HG11 | 1:F:215:THR:CG2 | 2.11 | 0.78 |
| 1:A:62:TRP:HE1 | 1:B:194:THR:HG1 | 1.32 | 0.78 |
| 1:F:62:TRP:HE1 | 1:G:194:THR:HG1 | 1.31 | 0.77 |
| 1:D:94:VAL:HG11 | 1:D:215:THR:CG2 | 2.11 | 0.77 |
| 1:A:194:THR:HG1 | 1:G:62:TRP:HE1 | 1.33 | 0.76 |
| 1:C:94:VAL:HG11 | 1:C:215:THR:CG2 | 2.11 | 0.76 |
| 1:E:94:VAL:HG11 | 1:E:215:THR:CG2 | 2.11 | 0.76 |
| 1:B:94:VAL:HG11 | 1:B:215:THR:CG2 | 2.11 | 0.75 |
| 1:B:62:TRP:HE1 | 1:C:194:THR:HG1 | 1.34 | 0.75 |
| 1:C:62:TRP:HE1 | 1:D:194:THR:HG1 | 1.32 | 0.74 |
| 1:E:62:TRP:HE1 | 1:F:194:THR:HG1 | 1.33 | 0.73 |
| 1:D:62:TRP:HE1 | 1:E:194:THR:HG1 | 1.34 | 0.72 |
| 1:F:71:ASN:ND2 | 1:F:102:MET:HG3 | 2.05 | 0.71 |
| 1:E:71:ASN:ND2 | 1:E:102:MET:HG3 | 2.05 | 0.71 |
| 1:G:71:ASN:ND2 | 1:G:102:MET:HG3 | 2.05 | 0.71 |
| 1:D:71:ASN:ND2 | 1:D:102:MET:HG3 | 2.05 | 0.71 |
| 1:C:71:ASN:ND2 | 1:C:102:MET:HG3 | 2.05 | 0.70 |
| 1:A:71:ASN:ND2 | 1:A:102:MET:HG3 | 2.05 | 0.70 |
| 1:B:71:ASN:ND2 | 1:B:102:MET:HG3 | 2.05 | 0.70 |
| 1:B:225:GLU:CD | 1:B:226:ARG:HH21 | 1.98 | 0.67 |
| 1:C:225:GLU:CD | 1:C:226:ARG:HH21 | 1.98 | 0.67 |
| 1:A:225:GLU:CD | 1:A:226:ARG:HH21 | 1.98 | 0.67 |
| 1:G:225:GLU:CD | 1:G:226:ARG:HH21 | 1.98 | 0.67 |
| 1:D:225:GLU:CD | 1:D:226:ARG:HH21 | 1.98 | 0.67 |
| 1:F:225:GLU:CD | 1:F:226:ARG:HH21 | 1.98 | 0.67 |
| 1:D:94:VAL:HG13 | 1:D:215:THR:CG2 | 2.25 | 0.67 |
| 1:E:235:GLU:HA | 1:E:238:LYS:HD2 | 1.77 | 0.67 |
| 1:D:235:GLU:HA | 1:D:238:LYS:HD2 | 1.77 | 0.67 |
| 1:E:94:VAL:HG13 | 1:E:215:THR:CG2 | 2.25 | 0.66 |
| 1:F:235:GLU:HA | 1:F:238:LYS:HD2 | 1.77 | 0.66 |
| 1:C:235:GLU:HA | 1:C:238:LYS:HD2 | 1.77 | 0.66 |
| 1:C:94:VAL:HG13 | 1:C:215:THR:CG2 | 2.25 | 0.66 |
| 1:E:225:GLU:CD | 1:E:226:ARG:HH21 | 1.98 | 0.66 |
| 1:G:94:VAL:HG13 | 1:G:215:THR:CG2 | 2.25 | 0.66 |
| 1:G:235:GLU:HA | 1:G:238:LYS:HD2 | 1.77 | 0.65 |
| 1:B:235:GLU:HA | 1:B:238:LYS:HD2 | 1.77 | 0.65 |
| 1:F:212:PHE:O | 1:F:216:LYS:HG2 | 1.97 | 0.65 |
| 1:E:32:GLN:HG2 | 1:F:9:GLN:CB | 2.27 | 0.65 |
| 1:A:94:VAL:HG13 | 1:A:215:THR:CG2 | 2.25 | 0.65 |
| 1:G:212:PHE:O | 1:G:216:LYS:HG2 | 1.97 | 0.65 |
| 1:A:212:PHE:O | 1:A:216:LYS:HG2 | 1.97 | 0.65 |
| 1:F:94:VAL:HG13 | 1:F:215:THR:CG2 | 2.25 | 0.65 |
| 1:B:212:PHE:O | 1:B:216:LYS:HG2 | 1.97 | 0.65 |
| 1:C:212:PHE:O | 1:C:216:LYS:HG2 | 1.97 | 0.64 |
| 1:E:212:PHE:O | 1:E:216:LYS:HG2 | 1.97 | 0.64 |
| 1:E:32:GLN:HG2 | 1:F:9:GLN:HA | 1.79 | 0.64 |
| 1:A:235:GLU:HA | 1:A:238:LYS:HD2 | 1.77 | 0.64 |
| 1:D:212:PHE:O | 1:D:216:LYS:HG2 | 1.97 | 0.64 |
| 1:B:94:VAL:HG13 | 1:B:215:THR:CG2 | 2.25 | 0.64 |
| 1:F:32:GLN:HG2 | 1:G:9:GLN:CB | 2.28 | 0.64 |
| 1:E:37:PHE:CZ | 1:F:183:GLN:OE1 | 2.51 | 0.64 |
| 1:F:37:PHE:CZ | 1:G:183:GLN:OE1 | 2.51 | 0.64 |
| 1:A:9:GLN:CB | 1:G:32:GLN:HG2 | 2.28 | 0.64 |
| 1:D:32:GLN:HG2 | 1:E:9:GLN:CB | 2.28 | 0.64 |
| 1:D:32:GLN:HG2 | 1:E:9:GLN:HA | 1.80 | 0.64 |
| 1:C:37:PHE:CZ | 1:D:183:GLN:OE1 | 2.50 | 0.63 |
| 1:C:32:GLN:HG2 | 1:D:9:GLN:CB | 2.28 | 0.63 |
| 1:A:32:GLN:HG2 | 1:B:9:GLN:CB | 2.28 | 0.63 |
| 1:A:37:PHE:CZ | 1:B:183:GLN:OE1 | 2.52 | 0.63 |
| 1:A:183:GLN:OE1 | 1:G:37:PHE:CZ | 2.52 | 0.63 |
| 1:B:37:PHE:CZ | 1:C:183:GLN:OE1 | 2.51 | 0.63 |
| 1:D:37:PHE:CZ | 1:E:183:GLN:OE1 | 2.51 | 0.63 |
| 1:E:32:GLN:HG2 | 1:F:9:GLN:CA | 2.29 | 0.63 |
| 1:B:32:GLN:HG2 | 1:C:9:GLN:CB | 2.28 | 0.62 |
| 1:B:28:LEU:HD12 | 1:B:29:ALA:N | 2.14 | 0.62 |
| 1:C:32:GLN:HG2 | 1:D:9:GLN:HA | 1.81 | 0.62 |
| 1:A:28:LEU:HD12 | 1:A:29:ALA:N | 2.14 | 0.62 |
| 1:A:9:GLN:HA | 1:G:32:GLN:HG2 | 1.81 | 0.62 |
| 1:E:28:LEU:HD12 | 1:E:29:ALA:N | 2.14 | 0.62 |
| 1:F:32:GLN:HG2 | 1:G:9:GLN:HA | 1.80 | 0.62 |
| 1:A:32:GLN:HG2 | 1:B:9:GLN:HA | 1.81 | 0.62 |
| 1:C:28:LEU:HD12 | 1:C:29:ALA:N | 2.14 | 0.62 |
| 1:F:32:GLN:HG2 | 1:G:9:GLN:CA | 2.30 | 0.62 |
| 1:G:28:LEU:HD12 | 1:G:29:ALA:N | 2.14 | 0.62 |
| 1:B:32:GLN:HG2 | 1:C:9:GLN:HA | 1.81 | 0.62 |
| 1:A:9:GLN:CA | 1:G:32:GLN:HG2 | 2.30 | 0.61 |
| 1:D:28:LEU:HD12 | 1:D:29:ALA:N | 2.14 | 0.61 |
| 1:D:32:GLN:HG2 | 1:E:9:GLN:CA | 2.30 | 0.61 |
| 1:A:32:GLN:HG2 | 1:B:9:GLN:CA | 2.30 | 0.61 |
| 1:C:32:GLN:HG2 | 1:D:9:GLN:CA | 2.30 | 0.61 |
| 1:B:32:GLN:HG2 | 1:C:9:GLN:CA | 2.30 | 0.61 |
| 1:F:28:LEU:HD12 | 1:F:29:ALA:N | 2.14 | 0.61 |
| 1:G:91:ASP:HB3 | 1:G:92:PRO:HD2 | 1.85 | 0.59 |
| 1:C:91:ASP:HB3 | 1:C:92:PRO:HD2 | 1.85 | 0.59 |
| 1:D:91:ASP:HB3 | 1:D:92:PRO:HD2 | 1.85 | 0.59 |
| 1:A:91:ASP:HB3 | 1:A:92:PRO:HD2 | 1.85 | 0.59 |
| 1:B:91:ASP:HB3 | 1:B:92:PRO:HD2 | 1.85 | 0.59 |
| 1:F:91:ASP:HB3 | 1:F:92:PRO:HD2 | 1.85 | 0.59 |
| 1:E:91:ASP:HB3 | 1:E:92:PRO:HD2 | 1.85 | 0.58 |
| 1:C:91:ASP:HB2 | 1:C:94:VAL:HG23 | 1.86 | 0.58 |
| 1:F:225:GLU:CD | 1:F:226:ARG:NH2 | 2.57 | 0.58 |
| 1:D:91:ASP:HB2 | 1:D:94:VAL:HG23 | 1.86 | 0.58 |
| 1:E:91:ASP:HB2 | 1:E:94:VAL:HG23 | 1.85 | 0.58 |
| 1:B:91:ASP:HB2 | 1:B:94:VAL:HG23 | 1.85 | 0.57 |
| 1:D:225:GLU:CD | 1:D:226:ARG:NH2 | 2.57 | 0.57 |
| 1:G:225:GLU:CD | 1:G:226:ARG:NH2 | 2.57 | 0.57 |
| 1:C:225:GLU:CD | 1:C:226:ARG:NH2 | 2.57 | 0.57 |
| 1:E:225:GLU:CD | 1:E:226:ARG:NH2 | 2.57 | 0.57 |
| 1:F:91:ASP:HB2 | 1:F:94:VAL:HG23 | 1.85 | 0.57 |
| 1:B:225:GLU:CD | 1:B:226:ARG:NH2 | 2.57 | 0.57 |
| 1:G:91:ASP:HB2 | 1:G:94:VAL:HG23 | 1.85 | 0.57 |
| 1:A:225:GLU:CD | 1:A:226:ARG:NH2 | 2.57 | 0.57 |
| 1:A:91:ASP:HB2 | 1:A:94:VAL:HG23 | 1.85 | 0.56 |
| 1:A:220:HIS:HD2 | 1:G:233:CYS:HB2 | 1.71 | 0.56 |
| 1:A:233:CYS:HB2 | 1:B:220:HIS:HD2 | 1.71 | 0.55 |
| 1:B:233:CYS:HB2 | 1:C:220:HIS:HD2 | 1.71 | 0.55 |
| 1:C:233:CYS:HB2 | 1:D:220:HIS:HD2 | 1.71 | 0.55 |
| 1:E:224:THR:HA | 1:E:227:LYS:HG2 | 1.89 | 0.55 |
| 1:F:233:CYS:HB2 | 1:G:220:HIS:HD2 | 1.71 | 0.55 |
| 1:F:224:THR:HA | 1:F:227:LYS:HG2 | 1.89 | 0.55 |
| 1:D:224:THR:HA | 1:D:227:LYS:HG2 | 1.89 | 0.55 |
| 1:E:74:ILE:HD11 | 1:F:201:ARG:HB3 | 1.87 | 0.55 |
| 1:D:233:CYS:HB2 | 1:E:220:HIS:HD2 | 1.71 | 0.55 |
| 1:G:224:THR:HA | 1:G:227:LYS:HG2 | 1.89 | 0.55 |
| 1:A:224:THR:HA | 1:A:227:LYS:HG2 | 1.89 | 0.55 |
| 1:C:224:THR:HA | 1:C:227:LYS:HG2 | 1.89 | 0.55 |
| 1:B:224:THR:HA | 1:B:227:LYS:HG2 | 1.89 | 0.54 |
| 1:E:233:CYS:HB2 | 1:F:220:HIS:HD2 | 1.71 | 0.54 |
| 1:D:74:ILE:HD11 | 1:E:201:ARG:HB3 | 1.89 | 0.54 |
| 1:A:201:ARG:HB3 | 1:G:74:ILE:HD11 | 1.89 | 0.54 |
| 1:A:74:ILE:HD11 | 1:B:201:ARG:HB3 | 1.89 | 0.54 |
| 1:F:74:ILE:HD11 | 1:G:201:ARG:HB3 | 1.89 | 0.54 |
| 1:C:74:ILE:HD11 | 1:D:201:ARG:HB3 | 1.89 | 0.53 |
| 1:A:233:CYS:HB2 | 1:B:220:HIS:CD2 | 2.44 | 0.53 |
| 1:A:220:HIS:CD2 | 1:G:233:CYS:HB2 | 2.44 | 0.53 |
| 1:F:233:CYS:HB2 | 1:G:220:HIS:CD2 | 2.44 | 0.53 |
| 1:B:233:CYS:HB2 | 1:C:220:HIS:CD2 | 2.44 | 0.53 |
| 1:B:74:ILE:HD11 | 1:C:201:ARG:HB3 | 1.89 | 0.53 |
| 1:E:233:CYS:HB2 | 1:F:220:HIS:CD2 | 2.44 | 0.52 |
| 1:C:233:CYS:HB2 | 1:D:220:HIS:CD2 | 2.44 | 0.52 |
| 1:D:233:CYS:HB2 | 1:E:220:HIS:CD2 | 2.44 | 0.52 |
| 1:E:245:ILE:HD11 | 1:F:232:THR:HG21 | 1.92 | 0.51 |
| 1:D:245:ILE:HD11 | 1:E:232:THR:HG21 | 1.93 | 0.51 |
| 1:F:245:ILE:HD11 | 1:G:232:THR:HG21 | 1.92 | 0.51 |
| 1:G:91:ASP:HB3 | 1:G:92:PRO:CD | 2.41 | 0.51 |
| 1:E:91:ASP:HB3 | 1:E:92:PRO:CD | 2.41 | 0.51 |
| 1:A:91:ASP:HB3 | 1:A:92:PRO:CD | 2.41 | 0.50 |
| 1:D:91:ASP:HB3 | 1:D:92:PRO:CD | 2.41 | 0.50 |
| 1:F:91:ASP:HB3 | 1:F:92:PRO:CD | 2.41 | 0.50 |
| 1:A:232:THR:HG21 | 1:G:245:ILE:HD11 | 1.93 | 0.50 |
| 1:C:91:ASP:HB3 | 1:C:92:PRO:CD | 2.41 | 0.50 |
| 1:E:25:ILE:O | 1:E:28:LEU:HG | 2.12 | 0.50 |
| 1:A:118:LEU:O | 1:A:182:SER:OG | 2.30 | 0.50 |
| 1:C:25:ILE:O | 1:C:28:LEU:HG | 2.12 | 0.50 |
| 1:B:91:ASP:HB3 | 1:B:92:PRO:CD | 2.41 | 0.50 |
| 1:A:25:ILE:O | 1:A:28:LEU:HG | 2.12 | 0.50 |
| 1:B:118:LEU:O | 1:B:182:SER:OG | 2.30 | 0.50 |
| 1:B:245:ILE:HD11 | 1:C:232:THR:HG21 | 1.93 | 0.50 |
| 1:D:118:LEU:O | 1:D:182:SER:OG | 2.30 | 0.50 |
| 1:C:245:ILE:HD11 | 1:D:232:THR:HG21 | 1.94 | 0.49 |
| 1:E:69:VAL:HG23 | 1:F:201:ARG:HD3 | 1.94 | 0.49 |
| 1:G:25:ILE:O | 1:G:28:LEU:HG | 2.12 | 0.49 |
| 1:A:245:ILE:HD11 | 1:B:232:THR:HG21 | 1.93 | 0.49 |
| 1:B:233:CYS:CB | 1:C:220:HIS:HD2 | 2.26 | 0.49 |
| 1:B:69:VAL:HG23 | 1:C:201:ARG:HD3 | 1.94 | 0.49 |
| 1:D:233:CYS:CB | 1:E:220:HIS:HD2 | 2.25 | 0.49 |
| 1:E:233:CYS:CB | 1:F:220:HIS:HD2 | 2.25 | 0.49 |
| 1:G:118:LEU:O | 1:G:182:SER:OG | 2.30 | 0.49 |
| 1:A:233:CYS:CB | 1:B:220:HIS:HD2 | 2.25 | 0.49 |
| 1:B:25:ILE:O | 1:B:28:LEU:HG | 2.12 | 0.49 |
| 1:F:69:VAL:HG23 | 1:G:201:ARG:HD3 | 1.94 | 0.49 |
| 1:F:25:ILE:O | 1:F:28:LEU:HG | 2.12 | 0.49 |
| 1:G:142:TYR:CE2 | 1:G:168:ILE:HG12 | 2.48 | 0.49 |
| 1:A:142:TYR:CE2 | 1:A:168:ILE:HG12 | 2.48 | 0.49 |
| 1:C:40:THR:HG22 | 1:D:183:GLN:NE2 | 2.28 | 0.49 |
| 1:A:220:HIS:HD2 | 1:G:233:CYS:CB | 2.25 | 0.49 |
| 1:B:142:TYR:CE2 | 1:B:168:ILE:HG12 | 2.48 | 0.49 |
| 1:D:142:TYR:CE2 | 1:D:168:ILE:HG12 | 2.48 | 0.49 |
| 1:E:118:LEU:O | 1:E:182:SER:OG | 2.30 | 0.49 |
| 1:E:142:TYR:CE2 | 1:E:168:ILE:HG12 | 2.48 | 0.49 |
| 1:D:25:ILE:O | 1:D:28:LEU:HG | 2.12 | 0.48 |
| 1:D:40:THR:HG22 | 1:E:183:GLN:NE2 | 2.28 | 0.48 |
| 1:F:142:TYR:CE2 | 1:F:168:ILE:HG12 | 2.48 | 0.48 |
| 1:C:118:LEU:O | 1:C:182:SER:OG | 2.30 | 0.48 |
| 1:A:40:THR:HG22 | 1:B:183:GLN:NE2 | 2.29 | 0.48 |
| 1:E:40:THR:HG22 | 1:F:183:GLN:NE2 | 2.28 | 0.48 |
| 1:F:40:THR:HG22 | 1:G:183:GLN:NE2 | 2.28 | 0.48 |
| 1:F:233:CYS:CB | 1:G:220:HIS:HD2 | 2.25 | 0.48 |
| 1:A:232:THR:HA | 1:A:235:GLU:HG3 | 1.96 | 0.48 |
| 1:A:69:VAL:HG23 | 1:B:201:ARG:HD3 | 1.94 | 0.48 |
| 1:A:245:ILE:HD11 | 1:B:229:PHE:HA | 1.96 | 0.48 |
| 1:B:232:THR:HA | 1:B:235:GLU:HG3 | 1.96 | 0.48 |
| 1:B:40:THR:HG22 | 1:C:183:GLN:NE2 | 2.29 | 0.48 |
| 1:C:232:THR:HA | 1:C:235:GLU:HG3 | 1.96 | 0.48 |
| 1:D:69:VAL:HG23 | 1:E:201:ARG:HD3 | 1.94 | 0.48 |
| 1:E:62:TRP:NE1 | 1:F:194:THR:OG1 | 2.37 | 0.48 |
| 1:G:232:THR:HA | 1:G:235:GLU:HG3 | 1.96 | 0.48 |
| 1:C:233:CYS:CB | 1:D:220:HIS:HD2 | 2.26 | 0.48 |
| 1:F:245:ILE:HD11 | 1:G:229:PHE:HA | 1.96 | 0.48 |
| 1:A:229:PHE:HA | 1:G:245:ILE:HD11 | 1.96 | 0.48 |
| 1:A:201:ARG:HD3 | 1:G:69:VAL:HG23 | 1.94 | 0.48 |
| 1:A:37:PHE:HZ | 1:B:183:GLN:OE1 | 1.97 | 0.48 |
| 1:C:37:PHE:HZ | 1:D:183:GLN:OE1 | 1.96 | 0.48 |
| 1:C:69:VAL:HG23 | 1:D:201:ARG:HD3 | 1.94 | 0.48 |
| 1:D:37:PHE:HZ | 1:E:183:GLN:OE1 | 1.97 | 0.48 |
| 1:E:37:PHE:HZ | 1:F:183:GLN:OE1 | 1.96 | 0.48 |
| 1:G:77:LEU:HD21 | 1:G:98:MET:HG3 | 1.96 | 0.48 |
| 1:B:245:ILE:HD11 | 1:C:229:PHE:HA | 1.96 | 0.48 |
| 1:C:142:TYR:CE2 | 1:C:168:ILE:HG12 | 2.48 | 0.48 |
| 1:F:77:LEU:HD21 | 1:F:98:MET:HG3 | 1.96 | 0.48 |
| 1:D:145:LEU:HD21 | 1:D:150:LEU:HD21 | 1.96 | 0.48 |
| 1:D:232:THR:HA | 1:D:235:GLU:HG3 | 1.96 | 0.48 |
| 1:E:77:LEU:HD21 | 1:E:98:MET:HG3 | 1.96 | 0.48 |
| 1:A:77:LEU:HD21 | 1:A:98:MET:HG3 | 1.96 | 0.47 |
| 1:B:37:PHE:HZ | 1:C:183:GLN:OE1 | 1.97 | 0.47 |
| 1:F:232:THR:HA | 1:F:235:GLU:HG3 | 1.96 | 0.47 |
| 1:D:179:ARG:O | 1:D:183:GLN:HG3 | 2.15 | 0.47 |
| 1:E:245:ILE:HD11 | 1:F:229:PHE:HA | 1.96 | 0.47 |
| 1:A:184:ALA:O | 1:A:188:THR:HG23 | 2.15 | 0.47 |
| 1:E:232:THR:HA | 1:E:235:GLU:HG3 | 1.96 | 0.47 |
| 1:C:145:LEU:HD21 | 1:C:150:LEU:HD21 | 1.96 | 0.47 |
| 1:E:145:LEU:HD21 | 1:E:150:LEU:HD21 | 1.96 | 0.47 |
| 1:F:118:LEU:O | 1:F:182:SER:OG | 2.30 | 0.47 |
| 1:G:184:ALA:O | 1:G:188:THR:HG23 | 2.15 | 0.47 |
| 1:B:179:ARG:O | 1:B:183:GLN:HG3 | 2.15 | 0.47 |
| 1:D:245:ILE:HD11 | 1:E:229:PHE:HA | 1.96 | 0.47 |
| 1:F:172:GLU:O | 1:F:176:ARG:HG3 | 2.15 | 0.47 |
| 1:A:179:ARG:O | 1:A:183:GLN:HG3 | 2.15 | 0.47 |
| 1:B:62:TRP:NE1 | 1:C:194:THR:OG1 | 2.37 | 0.47 |
| 1:C:227:LYS:HA | 1:C:230:ASP:OD2 | 2.15 | 0.47 |
| 1:C:77:LEU:HD21 | 1:C:98:MET:HG3 | 1.96 | 0.47 |
| 1:D:184:ALA:O | 1:D:188:THR:HG23 | 2.15 | 0.47 |
| 1:D:77:LEU:HD21 | 1:D:98:MET:HG3 | 1.96 | 0.47 |
| 1:E:184:ALA:O | 1:E:188:THR:HG23 | 2.15 | 0.47 |
| 1:G:145:LEU:HD21 | 1:G:150:LEU:HD21 | 1.96 | 0.47 |
| 1:D:172:GLU:O | 1:D:176:ARG:HG3 | 2.15 | 0.47 |
| 1:G:179:ARG:O | 1:G:183:GLN:HG3 | 2.15 | 0.47 |
| 1:G:227:LYS:HA | 1:G:230:ASP:OD2 | 2.15 | 0.47 |
| 1:A:145:LEU:HD21 | 1:A:150:LEU:HD21 | 1.96 | 0.47 |
| 1:C:179:ARG:O | 1:C:183:GLN:HG3 | 2.15 | 0.47 |
| 1:F:227:LYS:HA | 1:F:230:ASP:OD2 | 2.15 | 0.47 |
| 1:G:172:GLU:O | 1:G:176:ARG:HG3 | 2.15 | 0.47 |
| 1:A:183:GLN:NE2 | 1:G:40:THR:HG22 | 2.29 | 0.47 |
| 1:B:145:LEU:HD21 | 1:B:150:LEU:HD21 | 1.96 | 0.47 |
| 1:B:184:ALA:O | 1:B:188:THR:HG23 | 2.15 | 0.47 |
| 1:C:184:ALA:O | 1:C:188:THR:HG23 | 2.14 | 0.47 |
| 1:D:94:VAL:CG1 | 1:D:215:THR:HG23 | 2.44 | 0.47 |
| 1:E:227:LYS:HA | 1:E:230:ASP:OD2 | 2.15 | 0.47 |
| 1:B:77:LEU:HD21 | 1:B:98:MET:HG3 | 1.96 | 0.47 |
| 1:D:227:LYS:HA | 1:D:230:ASP:OD2 | 2.15 | 0.47 |
| 1:E:172:GLU:O | 1:E:176:ARG:HG3 | 2.15 | 0.47 |
| 1:E:237:ALA:HB1 | 1:F:221:TYR:HA | 1.97 | 0.47 |
| 1:E:234:LYS:HB3 | 1:E:238:LYS:NZ | 2.30 | 0.47 |
| 1:F:234:LYS:HB3 | 1:F:238:LYS:NZ | 2.30 | 0.47 |
| 1:G:234:LYS:HB3 | 1:G:238:LYS:NZ | 2.30 | 0.47 |
| 1:A:234:LYS:HB3 | 1:A:238:LYS:NZ | 2.30 | 0.47 |
| 1:C:245:ILE:HD11 | 1:D:229:PHE:HA | 1.97 | 0.47 |
| 1:E:179:ARG:O | 1:E:183:GLN:HG3 | 2.15 | 0.47 |
| 1:F:179:ARG:O | 1:F:183:GLN:HG3 | 2.15 | 0.47 |
| 1:A:221:TYR:HA | 1:G:237:ALA:HB1 | 1.97 | 0.47 |
| 1:B:227:LYS:HA | 1:B:230:ASP:OD2 | 2.15 | 0.46 |
| 1:C:172:GLU:O | 1:C:176:ARG:HG3 | 2.15 | 0.46 |
| 1:D:237:ALA:HB1 | 1:E:221:TYR:HA | 1.97 | 0.46 |
| 1:A:227:LYS:HA | 1:A:230:ASP:OD2 | 2.15 | 0.46 |
| 1:A:172:GLU:O | 1:A:176:ARG:HG3 | 2.15 | 0.46 |
| 1:A:237:ALA:HB1 | 1:B:221:TYR:HA | 1.97 | 0.46 |
| 1:B:172:GLU:O | 1:B:176:ARG:HG3 | 2.15 | 0.46 |
| 1:B:234:LYS:HB3 | 1:B:238:LYS:NZ | 2.30 | 0.46 |
| 1:C:234:LYS:HB3 | 1:C:238:LYS:NZ | 2.30 | 0.46 |
| 1:D:234:LYS:HB3 | 1:D:238:LYS:NZ | 2.30 | 0.46 |
| 1:B:189:PHE:O | 1:B:193:ILE:HG12 | 2.16 | 0.46 |
| 1:D:189:PHE:O | 1:D:193:ILE:HG12 | 2.16 | 0.46 |
| 1:F:145:LEU:HD21 | 1:F:150:LEU:HD21 | 1.96 | 0.46 |
| 1:F:184:ALA:O | 1:F:188:THR:HG23 | 2.15 | 0.46 |
| 1:F:237:ALA:HB1 | 1:G:221:TYR:HA | 1.97 | 0.46 |
| 1:B:237:ALA:HB1 | 1:C:221:TYR:HA | 1.97 | 0.46 |
| 1:C:189:PHE:O | 1:C:193:ILE:HG12 | 2.16 | 0.46 |
| 1:E:189:PHE:O | 1:E:193:ILE:HG12 | 2.16 | 0.46 |
| 1:A:94:VAL:CG1 | 1:A:215:THR:HG23 | 2.44 | 0.46 |
| 1:A:189:PHE:O | 1:A:193:ILE:HG12 | 2.16 | 0.46 |
| 1:C:57:ILE:O | 1:C:60:PRO:HG2 | 2.16 | 0.46 |
| 1:E:57:ILE:O | 1:E:60:PRO:HG2 | 2.16 | 0.46 |
| 1:F:189:PHE:O | 1:F:193:ILE:HG12 | 2.16 | 0.46 |
| 1:C:62:TRP:NE1 | 1:D:194:THR:OG1 | 2.36 | 0.45 |
| 1:D:57:ILE:O | 1:D:60:PRO:HG2 | 2.16 | 0.45 |
| 1:C:237:ALA:HB1 | 1:D:221:TYR:HA | 1.98 | 0.45 |
| 1:F:44:LEU:HD23 | 1:F:44:LEU:HA | 1.83 | 0.45 |
| 1:A:57:ILE:O | 1:A:60:PRO:HG2 | 2.16 | 0.45 |
| 1:G:189:PHE:O | 1:G:193:ILE:HG12 | 2.16 | 0.45 |
| 1:G:57:ILE:O | 1:G:60:PRO:HG2 | 2.16 | 0.45 |
| 1:B:57:ILE:O | 1:B:60:PRO:HG2 | 2.16 | 0.45 |
| 1:E:28:LEU:C | 1:E:28:LEU:HD12 | 2.38 | 0.45 |
| 1:F:57:ILE:O | 1:F:60:PRO:HG2 | 2.16 | 0.45 |
| 1:F:37:PHE:HZ | 1:G:183:GLN:OE1 | 1.97 | 0.45 |
| 1:F:28:LEU:C | 1:F:28:LEU:HD12 | 2.38 | 0.44 |
| 1:C:94:VAL:CG1 | 1:C:98:MET:CE | 2.88 | 0.44 |
| 1:C:211:ALA:O | 1:C:215:THR:HG23 | 2.18 | 0.44 |
| 1:D:211:ALA:O | 1:D:215:THR:HG23 | 2.17 | 0.44 |
| 1:D:28:LEU:C | 1:D:28:LEU:HD12 | 2.38 | 0.44 |
| 1:B:211:ALA:O | 1:B:215:THR:HG23 | 2.18 | 0.44 |
| 1:A:211:ALA:O | 1:A:215:THR:HG23 | 2.18 | 0.44 |
| 1:B:28:LEU:HD12 | 1:B:28:LEU:C | 2.38 | 0.44 |
| 1:B:94:VAL:CG1 | 1:B:98:MET:CE | 2.88 | 0.44 |
| 1:E:94:VAL:CG1 | 1:E:215:THR:HG23 | 2.44 | 0.44 |
| 1:G:188:THR:O | 1:G:192:VAL:HG23 | 2.18 | 0.44 |
| 1:G:44:LEU:HA | 1:G:44:LEU:HD23 | 1.83 | 0.44 |
| 1:B:188:THR:O | 1:B:192:VAL:HG23 | 2.18 | 0.44 |
| 1:E:211:ALA:O | 1:E:215:THR:HG23 | 2.17 | 0.44 |
| 1:C:28:LEU:C | 1:C:28:LEU:HD12 | 2.38 | 0.44 |
| 1:E:188:THR:O | 1:E:192:VAL:HG23 | 2.18 | 0.44 |
| 1:G:211:ALA:O | 1:G:215:THR:HG23 | 2.18 | 0.44 |
| 1:G:28:LEU:C | 1:G:28:LEU:HD12 | 2.38 | 0.44 |
| 1:A:188:THR:O | 1:A:192:VAL:HG23 | 2.18 | 0.43 |
| 1:C:188:THR:O | 1:C:192:VAL:HG23 | 2.18 | 0.43 |
| 1:D:188:THR:O | 1:D:192:VAL:HG23 | 2.18 | 0.43 |
| 1:A:245:ILE:CD1 | 1:B:228:LEU:HG | 2.49 | 0.43 |
| 1:E:44:LEU:HD23 | 1:E:44:LEU:HA | 1.84 | 0.43 |
| 1:A:194:THR:OG1 | 1:G:62:TRP:NE1 | 2.37 | 0.43 |
| 1:A:28:LEU:C | 1:A:28:LEU:HD12 | 2.38 | 0.43 |
| 1:B:94:VAL:CG1 | 1:B:215:THR:HG23 | 2.44 | 0.43 |
| 1:A:245:ILE:HD12 | 1:B:228:LEU:HG | 2.01 | 0.43 |
| 1:A:62:TRP:NE1 | 1:B:194:THR:OG1 | 2.37 | 0.43 |
| 1:F:211:ALA:O | 1:F:215:THR:HG23 | 2.18 | 0.43 |
| 1:A:228:LEU:HG | 1:G:245:ILE:CD1 | 2.48 | 0.43 |
| 1:D:245:ILE:CD1 | 1:E:228:LEU:HG | 2.49 | 0.43 |
| 1:B:245:ILE:HD12 | 1:C:228:LEU:HG | 2.01 | 0.43 |
| 1:D:245:ILE:HD12 | 1:E:228:LEU:HG | 2.01 | 0.43 |
| 1:A:228:LEU:HG | 1:G:245:ILE:HD12 | 2.01 | 0.43 |
| 1:F:94:VAL:CG1 | 1:F:215:THR:HG23 | 2.44 | 0.43 |
| 1:E:32:GLN:CG | 1:F:9:GLN:CB | 2.96 | 0.43 |
| 1:C:133:ASP:OD1 | 1:C:135:SER:OG | 2.31 | 0.42 |
| 1:E:58:VAL:HB | 1:E:59:PRO:HD3 | 2.01 | 0.42 |
| 1:C:58:VAL:HB | 1:C:59:PRO:HD3 | 2.01 | 0.42 |
| 1:E:222:ILE:O | 1:E:226:ARG:HG2 | 2.20 | 0.42 |
| 1:F:245:ILE:CD1 | 1:G:228:LEU:HG | 2.49 | 0.42 |
| 1:B:245:ILE:CD1 | 1:C:228:LEU:HG | 2.49 | 0.42 |
| 1:C:97:TYR:HD2 | 1:C:215:THR:HG21 | 1.85 | 0.42 |
| 1:D:97:TYR:HD2 | 1:D:215:THR:HG21 | 1.85 | 0.42 |
| 1:D:58:VAL:HB | 1:D:59:PRO:HD3 | 2.01 | 0.42 |
| 1:F:188:THR:O | 1:F:192:VAL:HG23 | 2.18 | 0.42 |
| 1:F:62:TRP:NE1 | 1:G:194:THR:OG1 | 2.37 | 0.42 |
| 1:G:222:ILE:O | 1:G:226:ARG:HG2 | 2.20 | 0.42 |
| 1:F:32:GLN:CG | 1:G:9:GLN:CB | 2.97 | 0.42 |
| 1:A:97:TYR:HD2 | 1:A:215:THR:HG21 | 1.85 | 0.42 |
| 1:B:97:TYR:HD2 | 1:B:215:THR:HG21 | 1.85 | 0.42 |
| 1:D:62:TRP:NE1 | 1:E:194:THR:OG1 | 2.37 | 0.42 |
| 1:A:9:GLN:CB | 1:G:32:GLN:CG | 2.97 | 0.42 |
| 1:B:222:ILE:O | 1:B:226:ARG:HG2 | 2.20 | 0.42 |
| 1:B:58:VAL:HB | 1:B:59:PRO:HD3 | 2.01 | 0.42 |
| 1:C:245:ILE:CD1 | 1:D:228:LEU:HG | 2.49 | 0.42 |
| 1:E:97:TYR:HD2 | 1:E:215:THR:HG21 | 1.85 | 0.42 |
| 1:F:245:ILE:HD12 | 1:G:228:LEU:HG | 2.01 | 0.42 |
| 1:C:222:ILE:O | 1:C:226:ARG:HG2 | 2.20 | 0.42 |
| 1:C:245:ILE:HD12 | 1:D:228:LEU:HG | 2.02 | 0.42 |
| 1:F:94:VAL:HG13 | 1:F:215:THR:HG23 | 2.02 | 0.42 |
| 1:A:228:LEU:HD21 | 1:G:242:LYS:HA | 2.02 | 0.42 |
| 1:C:94:VAL:CG1 | 1:C:215:THR:HG23 | 2.44 | 0.42 |
| 1:F:129:SER:OG | 1:F:154:GLN:O | 2.38 | 0.42 |
| 1:F:242:LYS:HA | 1:G:228:LEU:HD21 | 2.02 | 0.42 |
| 1:G:97:TYR:HD2 | 1:G:215:THR:HG21 | 1.85 | 0.42 |
| 1:D:208:THR:OG1 | 1:D:208:THR:O | 2.35 | 0.42 |
| 1:A:242:LYS:HA | 1:B:228:LEU:HD21 | 2.02 | 0.41 |
| 1:F:58:VAL:HB | 1:F:59:PRO:HD3 | 2.01 | 0.41 |
| 1:A:222:ILE:O | 1:A:226:ARG:HG2 | 2.20 | 0.41 |
| 1:B:242:LYS:HA | 1:C:228:LEU:HD21 | 2.03 | 0.41 |
| 1:E:245:ILE:HD12 | 1:F:228:LEU:HG | 2.02 | 0.41 |
| 1:E:242:LYS:HA | 1:F:228:LEU:HD21 | 2.02 | 0.41 |
| 1:A:183:GLN:OE1 | 1:G:37:PHE:HZ | 1.97 | 0.41 |
| 1:C:129:SER:OG | 1:C:154:GLN:O | 2.38 | 0.41 |
| 1:D:129:SER:OG | 1:D:154:GLN:O | 2.38 | 0.41 |
| 1:D:94:VAL:HG13 | 1:D:215:THR:HG23 | 2.02 | 0.41 |
| 1:F:222:ILE:O | 1:F:226:ARG:HG2 | 2.20 | 0.41 |
| 1:E:245:ILE:CD1 | 1:F:228:LEU:HG | 2.50 | 0.41 |
| 1:G:208:THR:OG1 | 1:G:208:THR:O | 2.35 | 0.41 |
| 1:A:129:SER:OG | 1:A:154:GLN:O | 2.38 | 0.41 |
| 1:A:94:VAL:HG13 | 1:A:215:THR:HG23 | 2.02 | 0.41 |
| 1:D:222:ILE:O | 1:D:226:ARG:HG2 | 2.20 | 0.41 |
| 1:E:129:SER:OG | 1:E:154:GLN:O | 2.38 | 0.41 |
| 1:D:242:LYS:HA | 1:E:228:LEU:HD21 | 2.02 | 0.41 |
| 1:A:58:VAL:HB | 1:A:59:PRO:HD3 | 2.01 | 0.41 |
| 1:G:58:VAL:HB | 1:G:59:PRO:HD3 | 2.01 | 0.41 |
| 1:D:94:VAL:CG1 | 1:D:98:MET:CE | 2.88 | 0.41 |
| 1:D:32:GLN:CG | 1:E:9:GLN:CB | 2.97 | 0.41 |
| 1:D:44:LEU:HA | 1:D:44:LEU:HD23 | 1.83 | 0.41 |
| 1:G:129:SER:OG | 1:G:154:GLN:O | 2.38 | 0.41 |
| 1:B:129:SER:OG | 1:B:154:GLN:O | 2.38 | 0.41 |
| 1:A:32:GLN:CG | 1:B:9:GLN:CB | 2.97 | 0.41 |
| 1:A:44:LEU:HD23 | 1:A:44:LEU:HA | 1.83 | 0.40 |
| 1:A:94:VAL:CG1 | 1:A:98:MET:CE | 2.88 | 0.40 |
| 1:F:97:TYR:HD2 | 1:F:215:THR:HG21 | 1.85 | 0.40 |
| 1:G:94:VAL:CG1 | 1:G:215:THR:HG23 | 2.44 | 0.40 |
| 1:C:32:GLN:CG | 1:D:9:GLN:CB | 2.97 | 0.40 |

There are no symmetry-related clashes.

### 5.3 Torsion angles

#### 5.3.1 Protein backbone

| Mol | Chain | Analysed | Favoured | Allowed | Outliers | Percentiles |  |
| --- | --- | --- | --- | --- | --- | --- | --- |
| 1 | A | 225/358 (63%) | 219 (97%) | 6 (3%) | 0 | 100 | 100 |
| 1 | B | 225/358 (63%) | 219 (97%) | 6 (3%) | 0 | 100 | 100 |
| 1 | C | 225/358 (63%) | 219 (97%) | 6 (3%) | 0 | 100 | 100 |
| 1 | D | 225/358 (63%) | 219 (97%) | 6 (3%) | 0 | 100 | 100 |
| 1 | E | 225/358 (63%) | 219 (97%) | 6 (3%) | 0 | 100 | 100 |
| 1 | F | 225/358 (63%) | 219 (97%) | 6 (3%) | 0 | 100 | 100 |
| 1 | G | 225/358 (63%) | 219 (97%) | 6 (3%) | 0 | 100 | 100 |
| All | All | 1575/2506 (63%) | 1533 (97%) | 42 (3%) | 0 | 100 | 100 |

There are no Ramachandran outliers to report.

#### 5.3.2 Protein sidechains

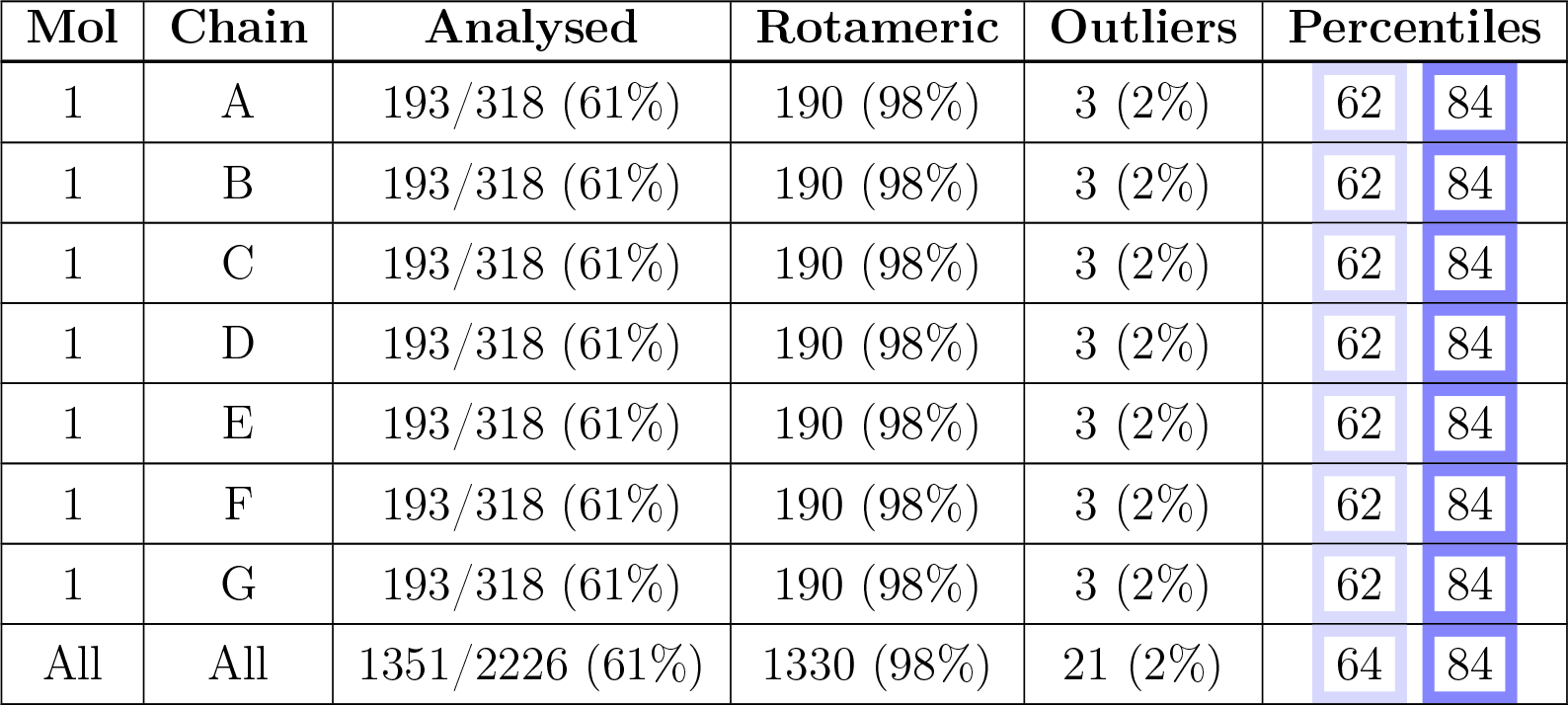

| Mol | Chain | Analysed | Rotameric | Outliers | Percentiles |  |
| --- | --- | --- | --- | --- | --- | --- |
| 1 | A | 193/318 (61%) | 190 (98%) | 3 (2%) | 62 | 84 |
| 1 | B | 193/318 (61%) | 190 (98%) | 3 (2%) | 62 | 84 |
| 1 | C | 193/318 (61%) | 190 (98%) | 3 (2%) | 62 | 84 |
| 1 | D | 193/318 (61%) | 190 (98%) | 3 (2%) | 62 | 84 |
| 1 | E | 193/318 (61%) | 190 (98%) | 3 (2%) | 62 | 84 |
| 1 | F | 193/318 (61%) | 190 (98%) | 3 (2%) | 62 | 84 |
| 1 | G | 193/318 (61%) | 190 (98%) | 3 (2%) | 62 | 84 |
| All | All | 1351/2226 (61%) | 1330 (98%) | 21 (2%) | 64 | 84 |

All (21) residues with a non-rotameric sidechain are listed below:

| Mol | Chain | Res | Type |
| --- | --- | --- | --- |
| 1 | A | 68 | TYR |
| 1 | A | 105 | ARG |
| 1 | A | 217 | TYR |
| 1 | B | 68 | TYR |
| 1 | B | 105 | ARG |
| 1 | B | 217 | TYR |
| 1 | C | 68 | TYR |
| 1 | C | 105 | ARG |
| 1 | C | 217 | TYR |
| 1 | D | 68 | TYR |
| 1 | D | 105 | ARG |
| 1 | D | 217 | TYR |
| 1 | E | 68 | TYR |
| 1 | E | 105 | ARG |
| 1 | E | 217 | TYR |
| 1 | F | 68 | TYR |
| 1 | F | 105 | ARG |
| 1 | F | 217 | TYR |
| 1 | G | 68 | TYR |
| 1 | G | 105 | ARG |
| 1 | G | 217 | TYR |

Sometimes sidechains can be flipped to improve hydrogen bonding and reduce clashes. All (21) such sidechains are listed below:

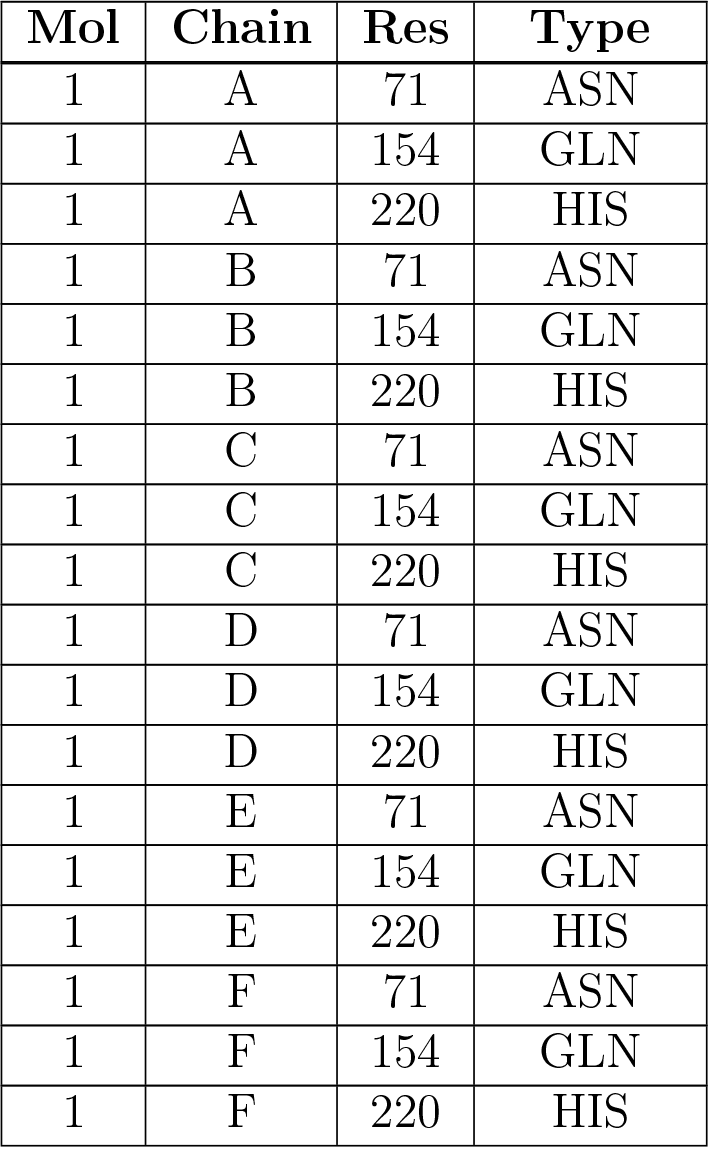
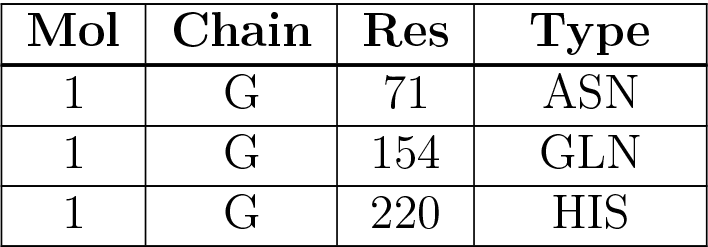

| Mol | Chain | Res | Type |
| --- | --- | --- | --- |
| 1 | A | 71 | ASN |
| 1 | A | 154 | GLN |
| 1 | A | 220 | HIS |
| 1 | B | 71 | ASN |
| 1 | B | 154 | GLN |
| 1 | B | 220 | HIS |
| 1 | C | 71 | ASN |
| 1 | C | 154 | GLN |
| 1 | C | 220 | HIS |
| 1 | D | 71 | ASN |
| 1 | D | 154 | GLN |
| 1 | D | 220 | HIS |
| 1 | E | 71 | ASN |
| 1 | E | 154 | GLN |
| 1 | E | 220 | HIS |
| 1 | F | 71 | ASN |
| 1 | F | 154 | GLN |
| 1 | F | 220 | HIS |
| 1 | G | 71 | ASN |
| 1 | G | 154 | GLN |
| 1 | G | 220 | HIS |

#### 5.3.3 RNA

There are no RNA molecules in this entry.

### 5.4 Non-standard residues in protein, DNA, RNA chains

There are no non-standard protein/DNA/RNA residues in this entry.

### 5.5 Carbohydrates

There are no monosaccharides in this entry.

### 5.6 Ligand geometry

7 ligands are modelled in this entry.

| Mol | Type | Chain | Res | Link | Bond lengths |  |  | Bond angles |  |  |
| --- | --- | --- | --- | --- | --- | --- | --- | --- | --- | --- |
| | | | | | Counts | RMSZ | $\#\vert Z\vert > 2$ | Counts | RMSZ | $\#\vert Z\vert > 2$ |
| 2 | NAG | F | 1001 | 1 | 14,14,15 | 0.25 | 0 | 17,19,21 | 0.61 | 1 (5%) |
| 2 | NAG | E | 1001 | 1 | 14,14,15 | 0.25 | 0 | 17,19,21 | 0.61 | 1 (5%) |
| 2 | NAG | G | 1001 | 1 | 14,14,15 | 0.24 | 0 | 17,19,21 | 0.61 | 1 (5%) |
| 2 | NAG | A | 1001 | 1 | 14,14,15 | 0.25 | 0 | 17,19,21 | 0.61 | 1 (5%) |
| 2 | NAG | C | 1001 | 1 | 14,14,15 | 0.26 | 0 | 17,19,21 | 0.61 | 1 (5%) |
| 2 | NAG | B | 1001 | 1 | 14,14,15 | 0.24 | 0 | 17,19,21 | 0.61 | 1 (5%) |
| 2 | NAG | D | 1001 | 1 | 14,14,15 | 0.25 | 0 | 17,19,21 | 0.61 | 1 (5%) |

| Mol | Type | Chain | Res | Link | Chirals | Torsions | Rings |
| --- | --- | --- | --- | --- | --- | --- | --- |
| 2 | NAG | F | 1001 | 1 | - | 2/6/23/26 | 0/1/1/1 |
| 2 | NAG | E | 1001 | 1 | - | 2/6/23/26 | 0/1/1/1 |
| 2 | NAG | G | 1001 | 1 | - | 2/6/23/26 | 0/1/1/1 |
| 2 | NAG | A | 1001 | 1 | - | 2/6/23/26 | 0/1/1/1 |
| 2 | NAG | C | 1001 | 1 | - | 2/6/23/26 | 0/1/1/1 |
| 2 | NAG | B | 1001 | 1 | - | 2/6/23/26 | 0/1/1/1 |
| 2 | NAG | D | 1001 | 1 | - | 2/6/23/26 | 0/1/1/1 |

There are no bond length outliers.

All (7) bond angle outliers are listed below:

There are no bond length outliers. All (7) bond angle outliers are listed below:
| Mol | Chain | Res | Type | Atoms | Z | Observed(°) | Ideal(°) |
| --- | --- | --- | --- | --- | --- | --- | --- |
| 2 | G | 1001 | NAG | C1-O5-C5 | 2.06 | 114.99 | 112.19 |
| 2 | B | 1001 | NAG | C1-O5-C5 | 2.05 | 114.97 | 112.19 |
| 2 | C | 1001 | NAG | C1-O5-C5 | 2.04 | 114.95 | 112.19 |
| 2 | A | 1001 | NAG | C1-O5-C5 | 2.04 | 114.95 | 112.19 |
| 2 | F | 1001 | NAG | C1-O5-C5 | 2.03 | 114.95 | 112.19 |
| 2 | E | 1001 | NAG | C1-O5-C5 | 2.03 | 114.94 | 112.19 |
| 2 | D | 1001 | NAG | C1-O5-C5 | 2.03 | 114.94 | 112.19 |

There are no chirality outliers.

All (14) torsion outliers are listed below:

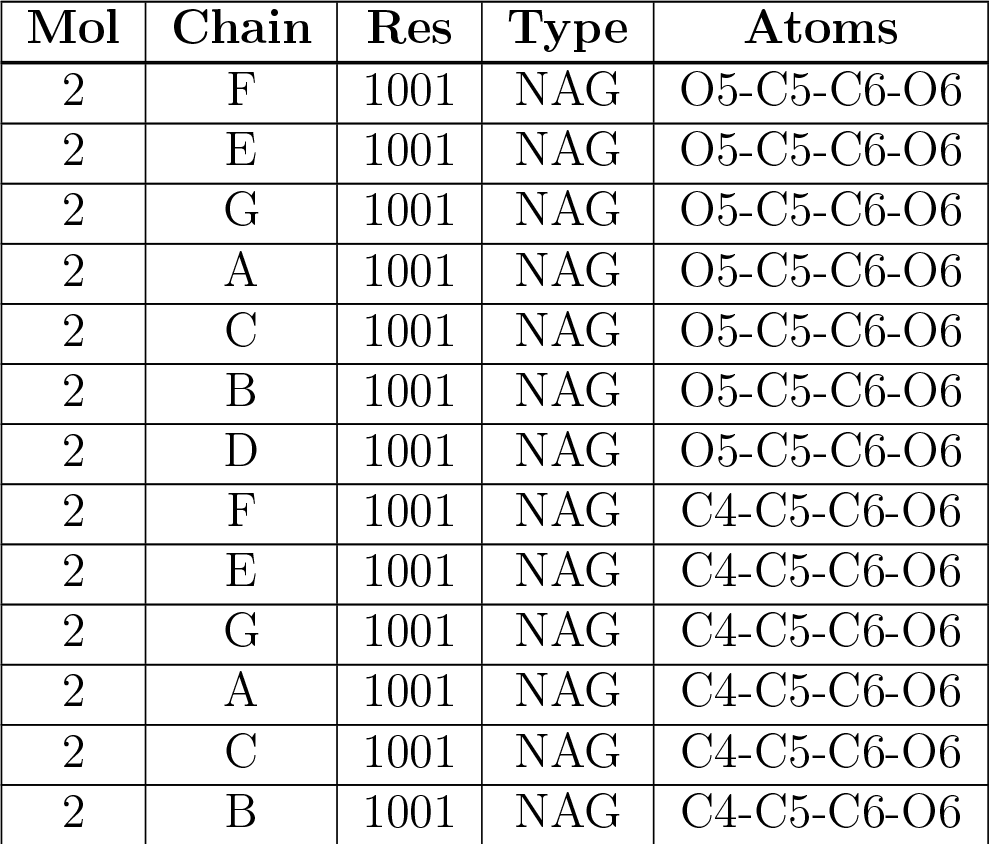
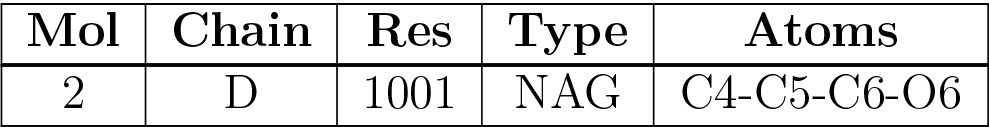

There are no chirality outliers. All (14) torsion outliers are listed below:
| Mol | Chain | Res | Type | Atoms |
| --- | --- | --- | --- | --- |
| 2 | F | 1001 | NAG | O5-C5-C6-O6 |
| 2 | E | 1001 | NAG | O5-C5-C6-O6 |
| 2 | G | 1001 | NAG | O5-C5-C6-O6 |
| 2 | A | 1001 | NAG | O5-C5-C6-O6 |
| 2 | C | 1001 | NAG | O5-C5-C6-O6 |
| 2 | B | 1001 | NAG | O5-C5-C6-O6 |
| 2 | D | 1001 | NAG | O5-C5-C6-O6 |
| 2 | F | 1001 | NAG | C4-C5-C6-O6 |
| 2 | E | 1001 | NAG | C4-C5-C6-O6 |
| 2 | G | 1001 | NAG | C4-C5-C6-O6 |
| 2 | A | 1001 | NAG | C4-C5-C6-O6 |
| 2 | C | 1001 | NAG | C4-C5-C6-O6 |
| 2 | B | 1001 | NAG | C4-C5-C6-O6 |
| 2 | D | 1001 | NAG | C4-C5-C6-O6 |

There are no ring outliers.

No monomer is involved in short contacts.

### 5.7 Other polymers

There are no such residues in this entry.

### 5.8 Polymer linkage issues

There are no chain breaks in this entry.

## 6 Map visualisation

This section contains visualisations of the EMDB entry EMD-30832. These allow visual inspection of the internal detail of the map and identification of artifacts.

### 6.1 Orthogonal projections

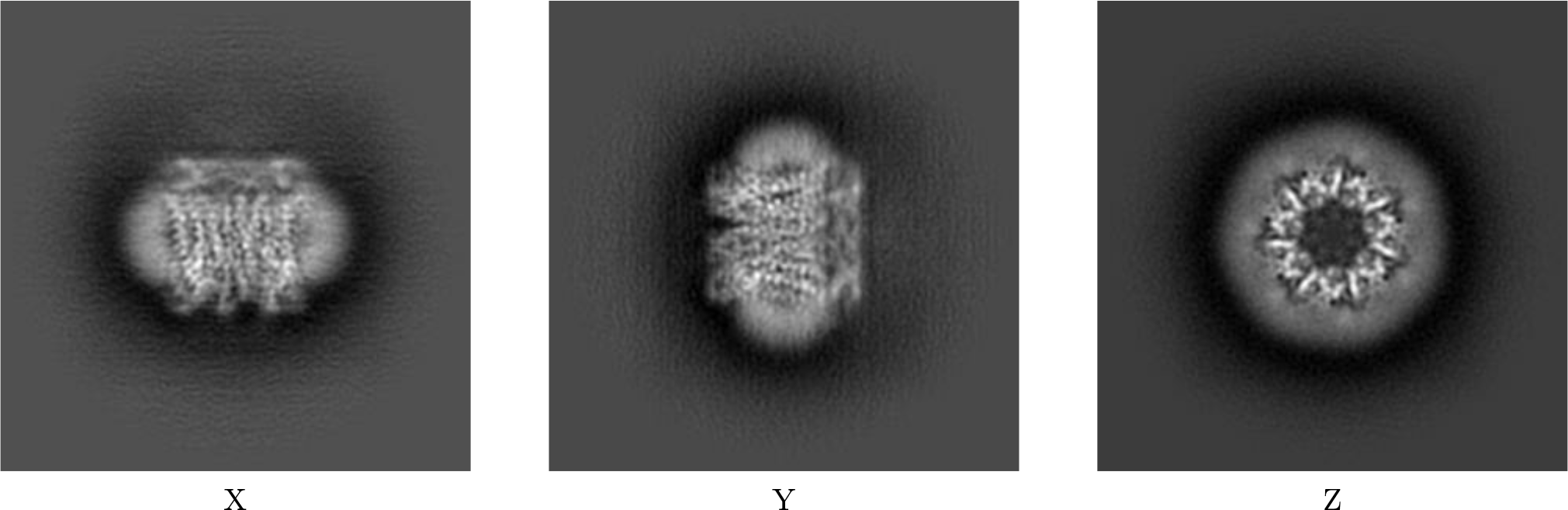

#### 6.1.1 Primary map

The images above show the map projected in three orthogonal directions.

### 6.2 Central slices

#### 6.2.1 Primary map

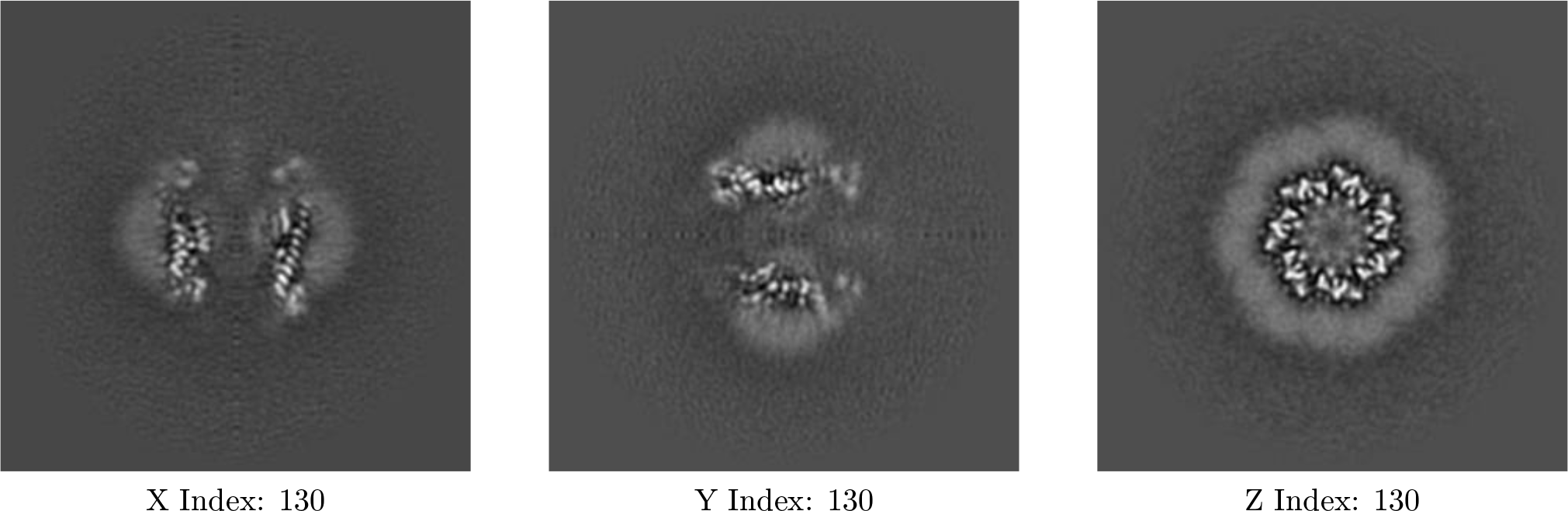

The images above show central slices of the map in three orthogonal directions.

### 6.3 Largest variance slices

#### 6.3.1 Primary map

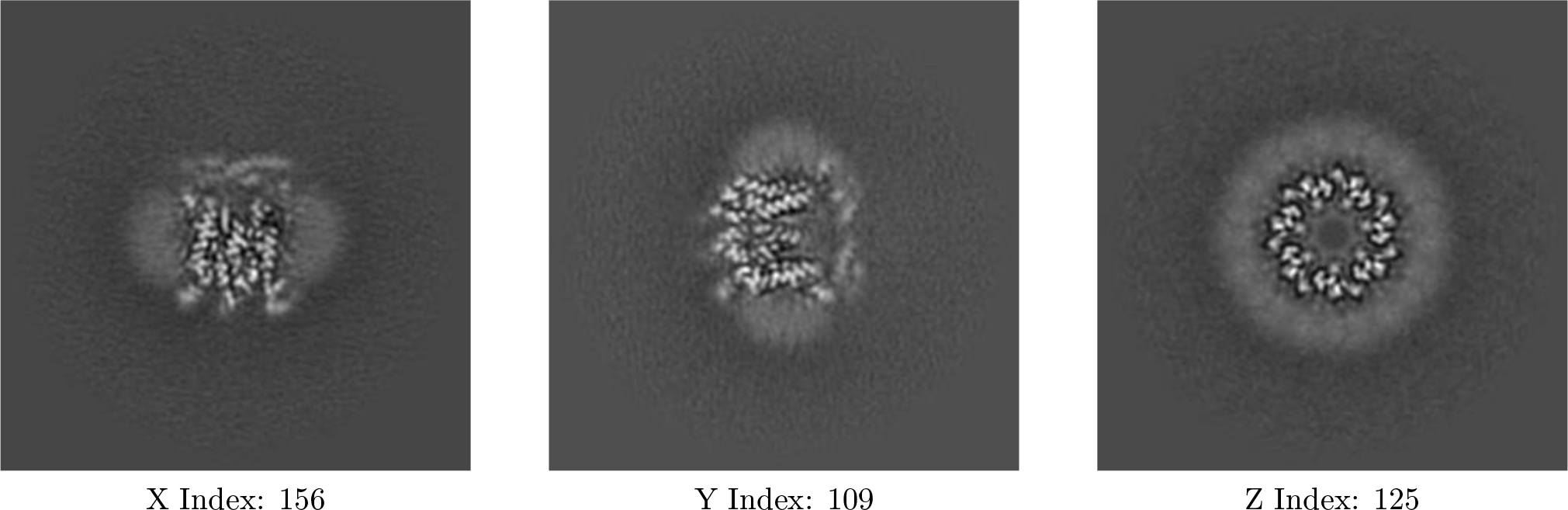

The images above show the largest variance slices of the map in three orthogonal directions.

### 6.4 Orthogonal surface views

#### 6.4.1 Primary map

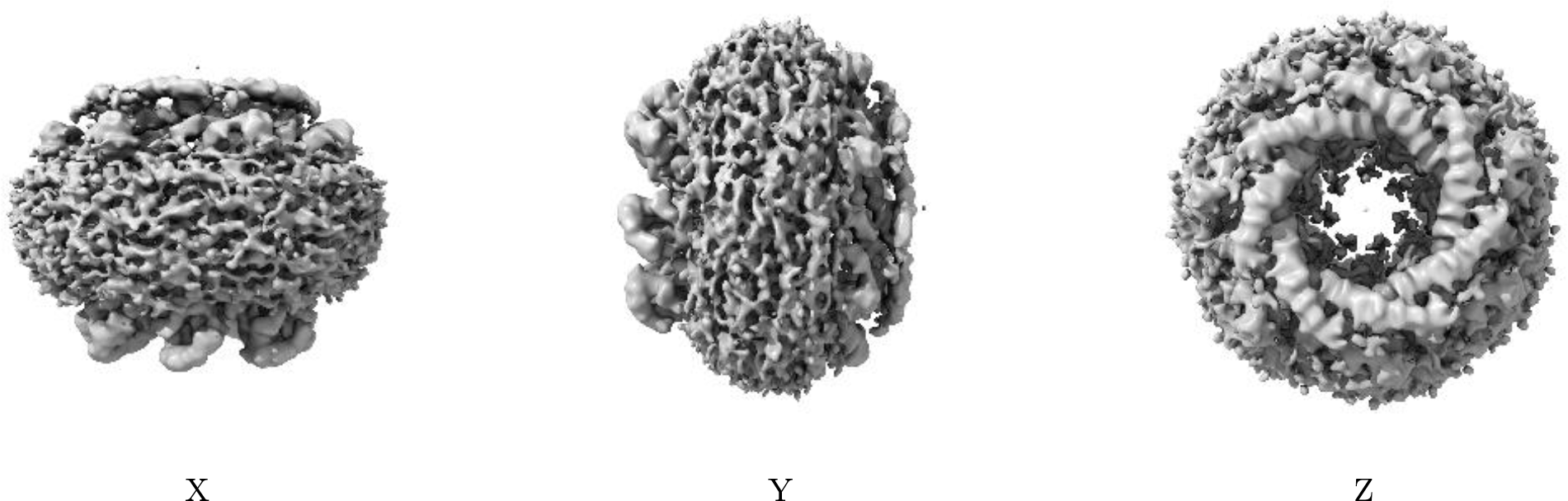

The images above show the 3D surface view of the map at the recommended contour level 0.209. These images, in conjunction with the slice images, may facilitate assessment of whether an appropriate contour level has been provided.

### 6.5 Mask visualisation

This section was not generated. No masks/segmentation were deposited.

## 7 Map analysis

This section contains the results of statistical analysis of the map.

### 7.1 Map-value distribution

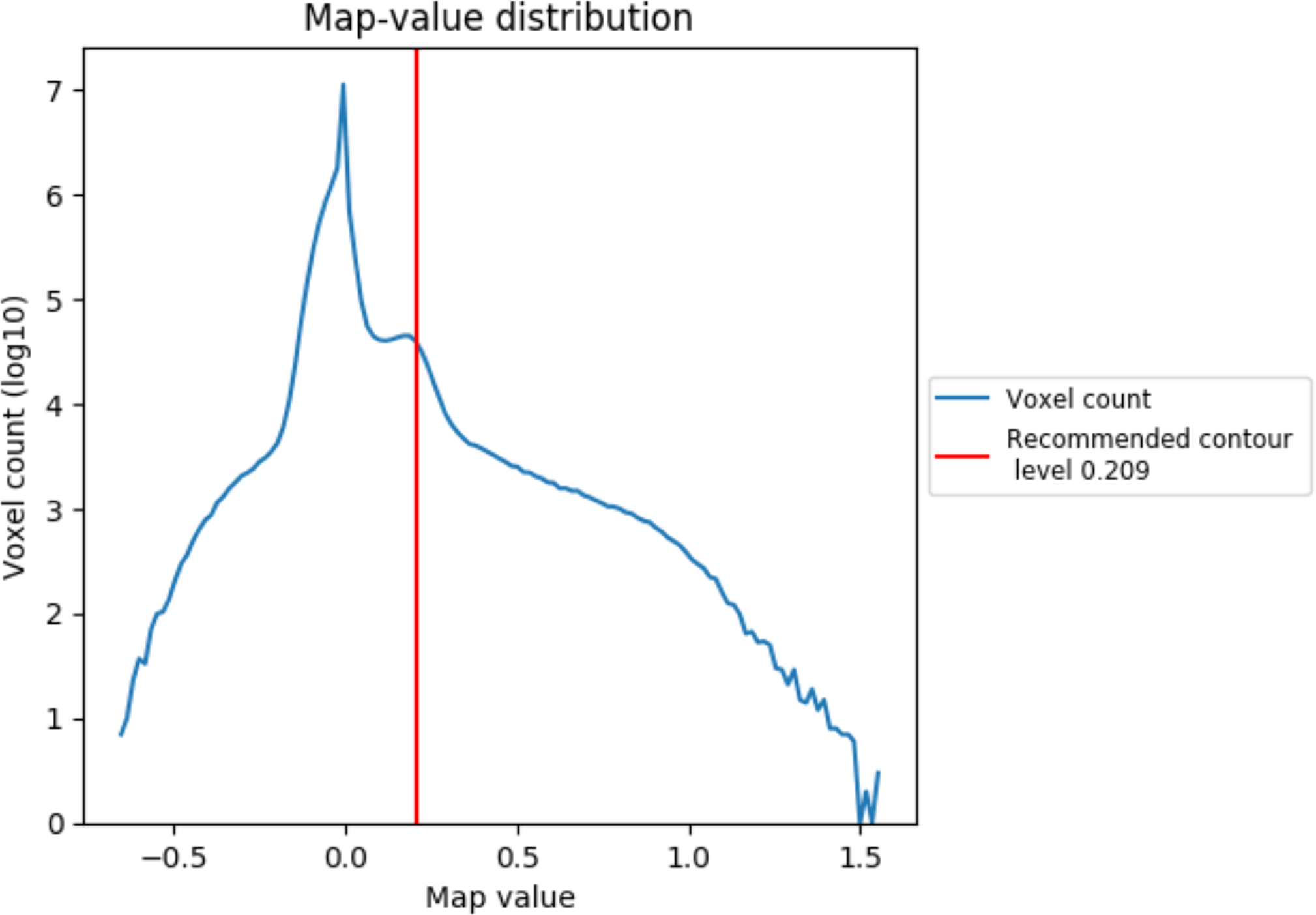

### 7.2 Volume estimate

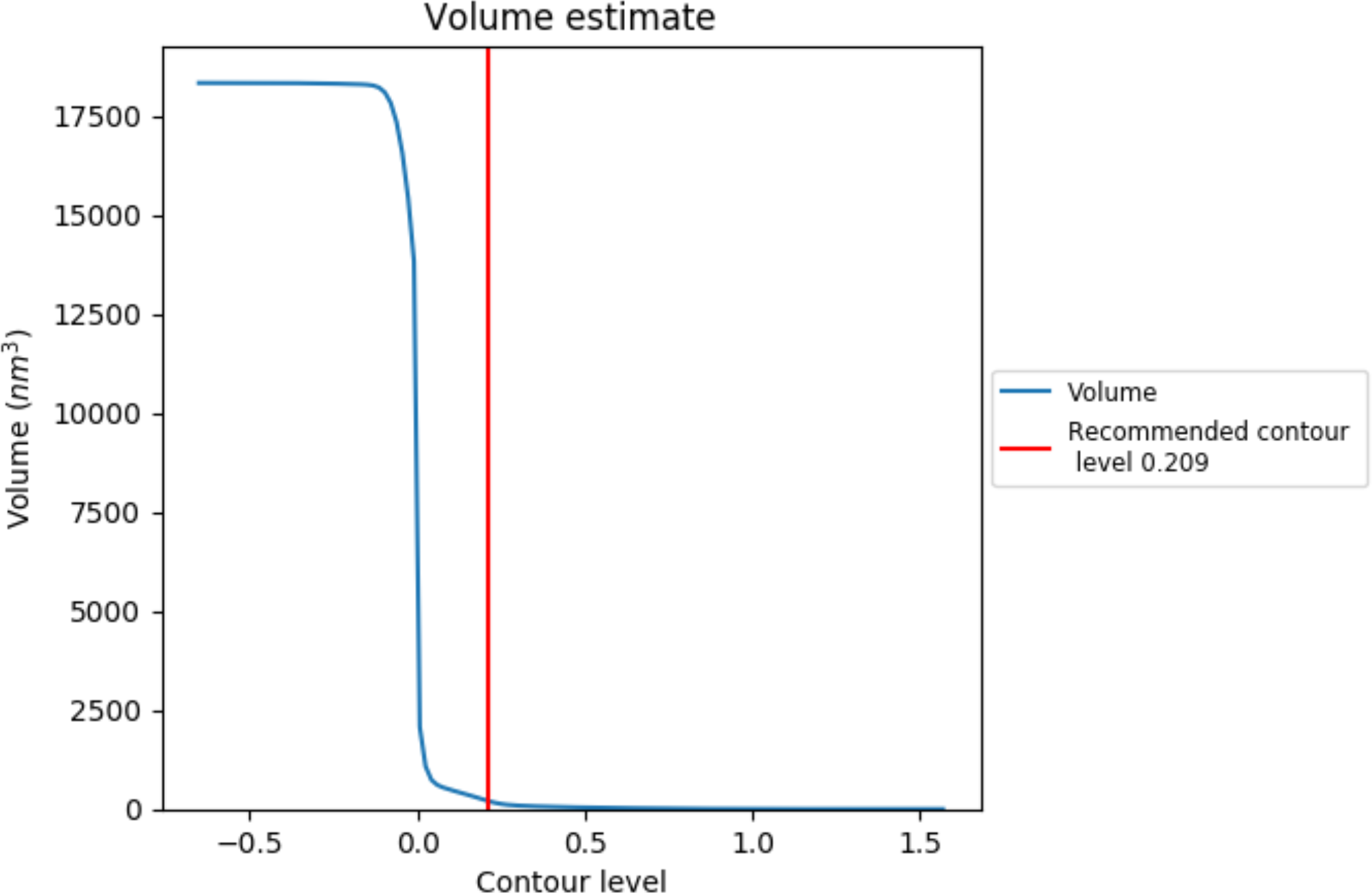

The volume at the recommended contour level is 211 nm^3^; this corresponds to an approximate mass of 191 kDa.

### 7.3 Rotationally averaged power spectrum

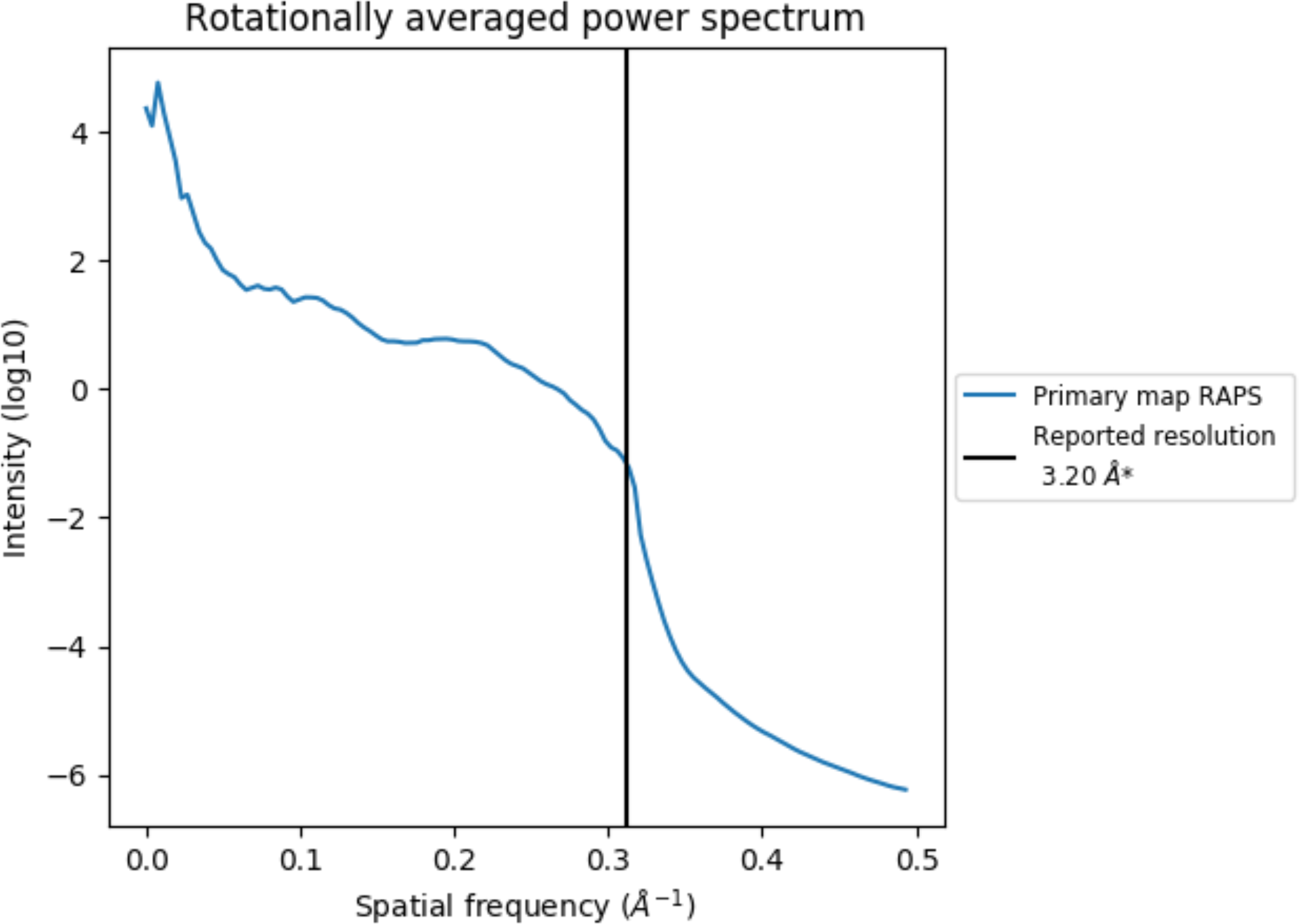

## 8 Fourier-Shell correlation

This section was not generated. No FSC curve or half-maps provided.

## 9 Map-model fit

This section contains information regarding the fit between EMDB map EMD-30832 and PDB model 7DSE. Per-residue inclusion information can be found in section 3 on page 7.

### 9.1 Map-model overlay

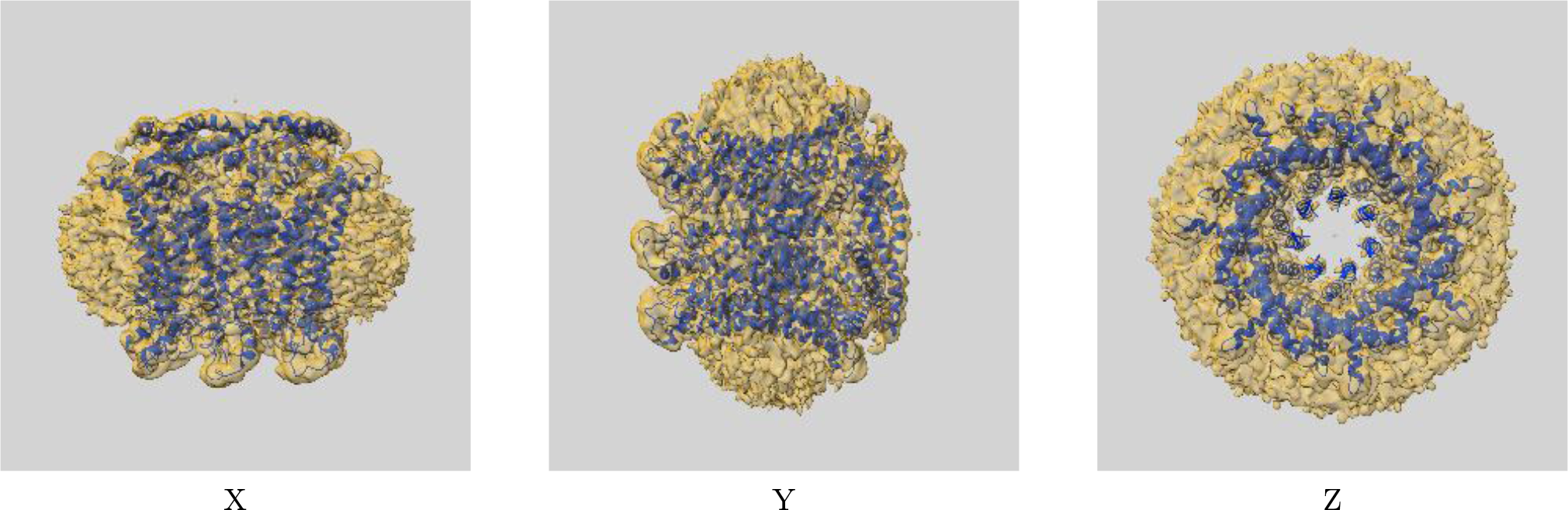

The images above show the 3D surface view of the map at the recommended contour level 0.209 at 50% transparency in yellow overlaid with a ribbon representation of the model coloured in blue. These images allow for the visual assessment of the quality of fit between the atomic model and the map.

### 9.2 Atom inclusion

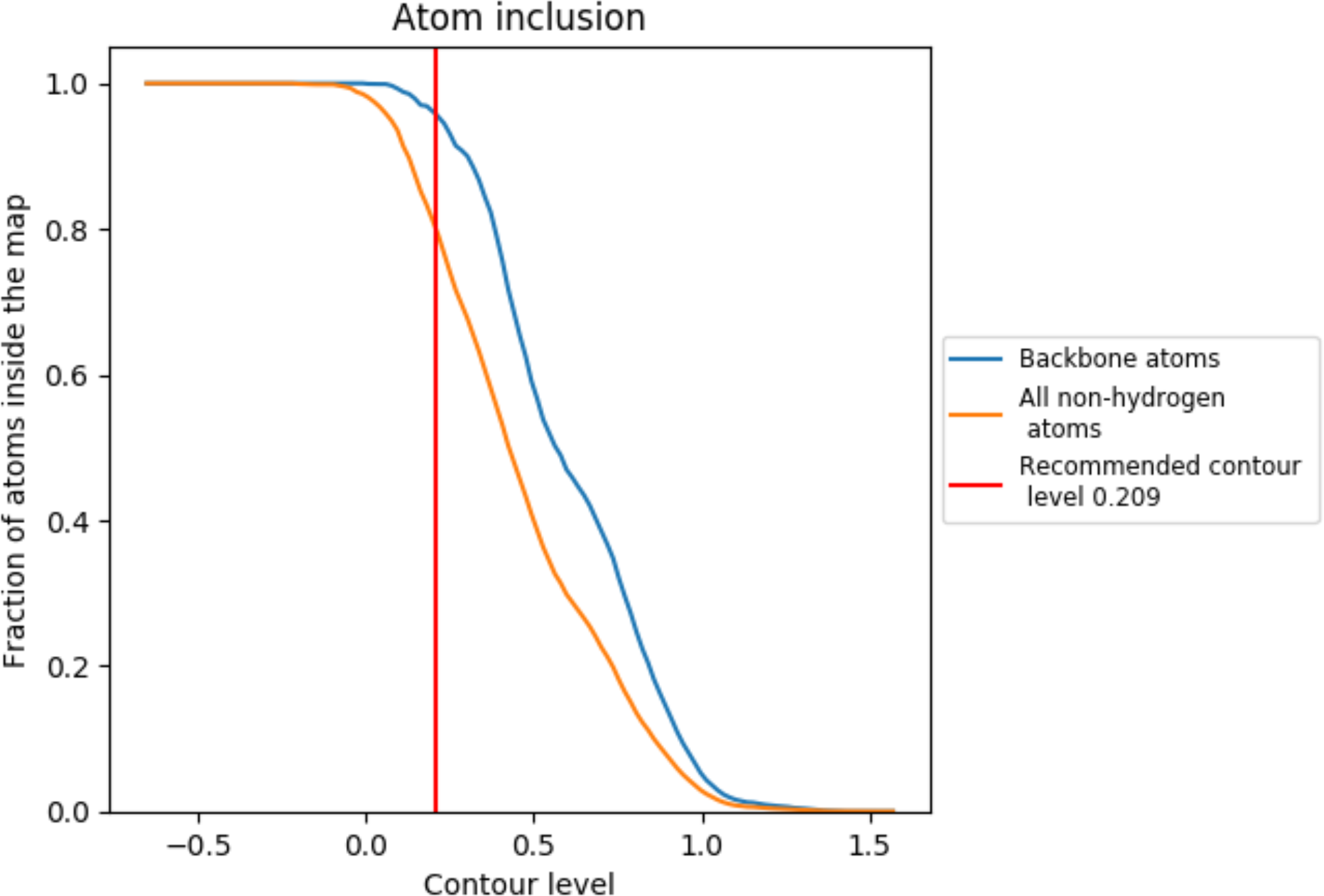

At the recommended contour level, 96% of all backbone atoms, 80% of all non-hydrogen atoms, are inside the map.

---

**Supplemental Figure 1.**
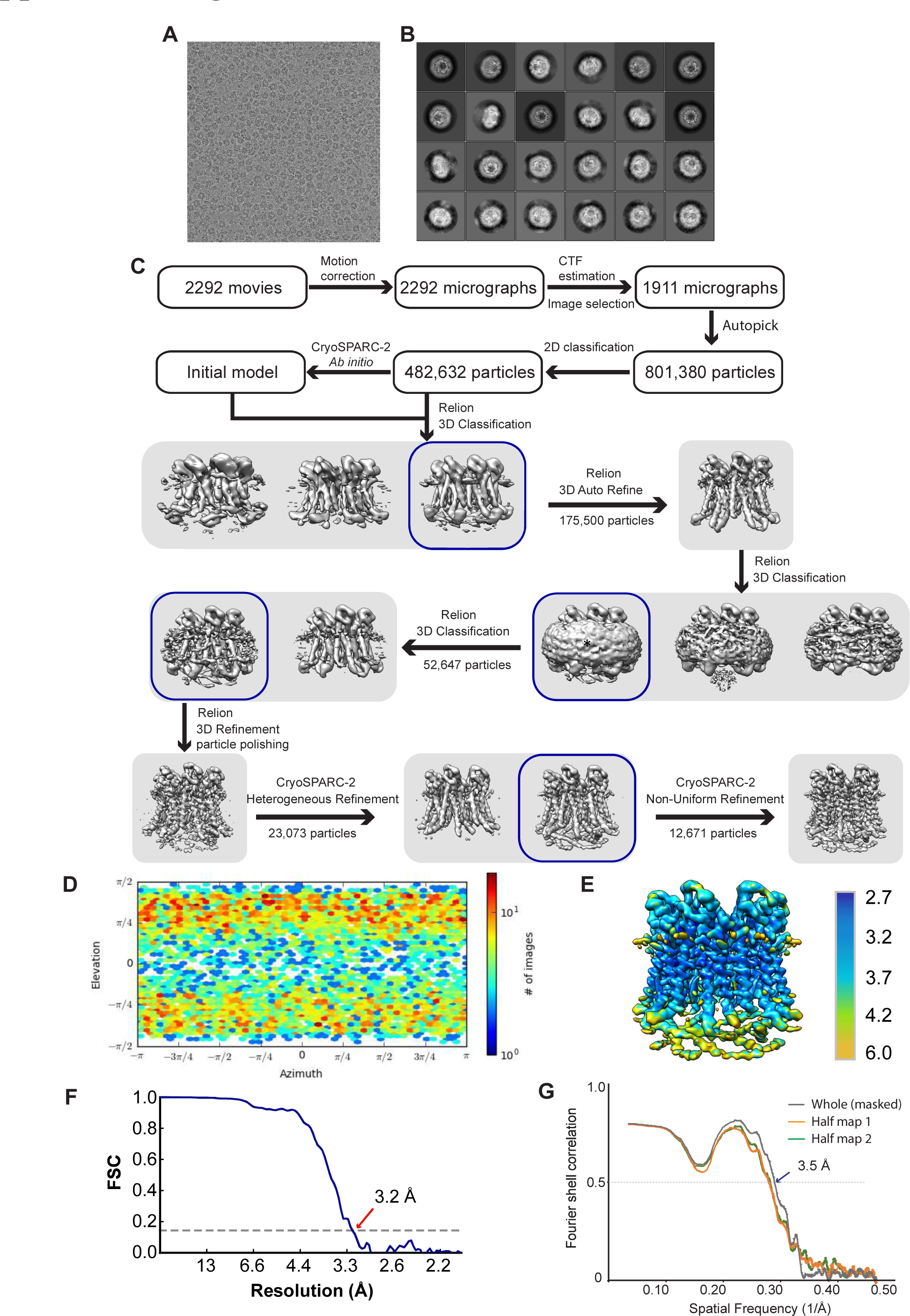
Structure determination of drCALHM1-hepta-LCH channel. (A) A drift-corrected cryoEM micro- graph of the drCALHM1. (B) Representative 2D class averages of the drCALHM1 obtained in Relion-3. (C) Workflow of image processing, 3D reconstruction and structure refinement. (D) Euler angle distribution of drCALHM1 in the final 3D reconstruction in cryoSPARC v2. (E) Local resolution estimation from ResMap. (F) Gold standard FSC curve of drCALHM1 is estimated by cryoSPARC v2. (G) FSC curves for cross-validation between maps and models: model versus summed map (grey), model versus half map 1 (orange) and model versus half map 2 (green).

**Supplemental Figure 2.**
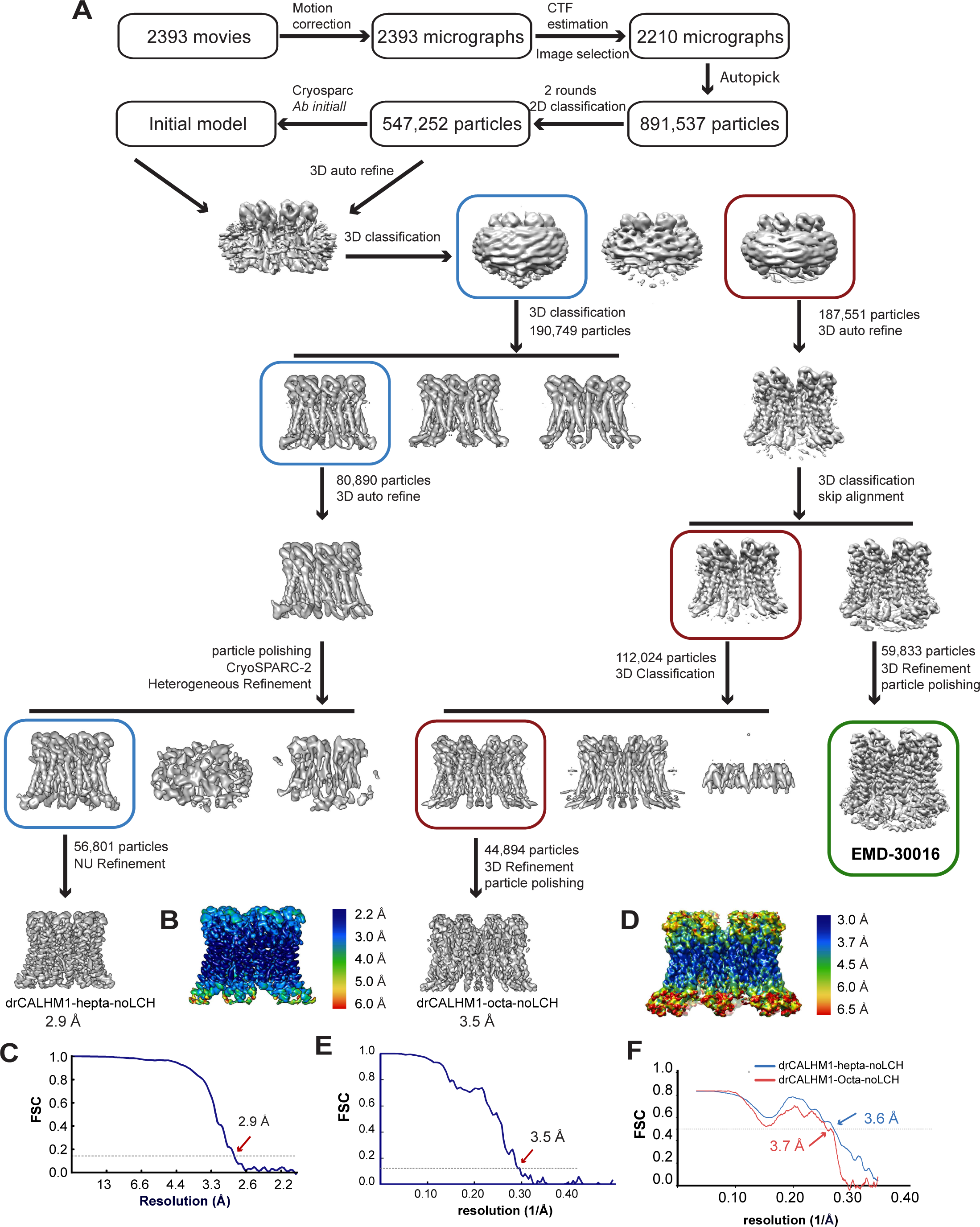
Structure determination of drCALHM1-hepta-noLCH and drCALHM1-octa-noLCH. (A)Workflow if image processing, 2D reconstruction and structure refinement. (B) Local resolution estimation of drCALHM1-hep-ta-noLCH from ResMap. (C) Gold standard FSC curve of drCALHM1-hepta-noLCH. (D) Local resolution estimation of drCALHM1-octa-noLCH from ResMap. (E) Gold standard FSC curve of drCALHM1-octa-noLCH. (F) FSC curves for cross-validation between maps and models of drCALHM1-hepta-noLCH (blue) and drCALHM1-octa-noLCH (red).

**Supplemental Figure 3.**
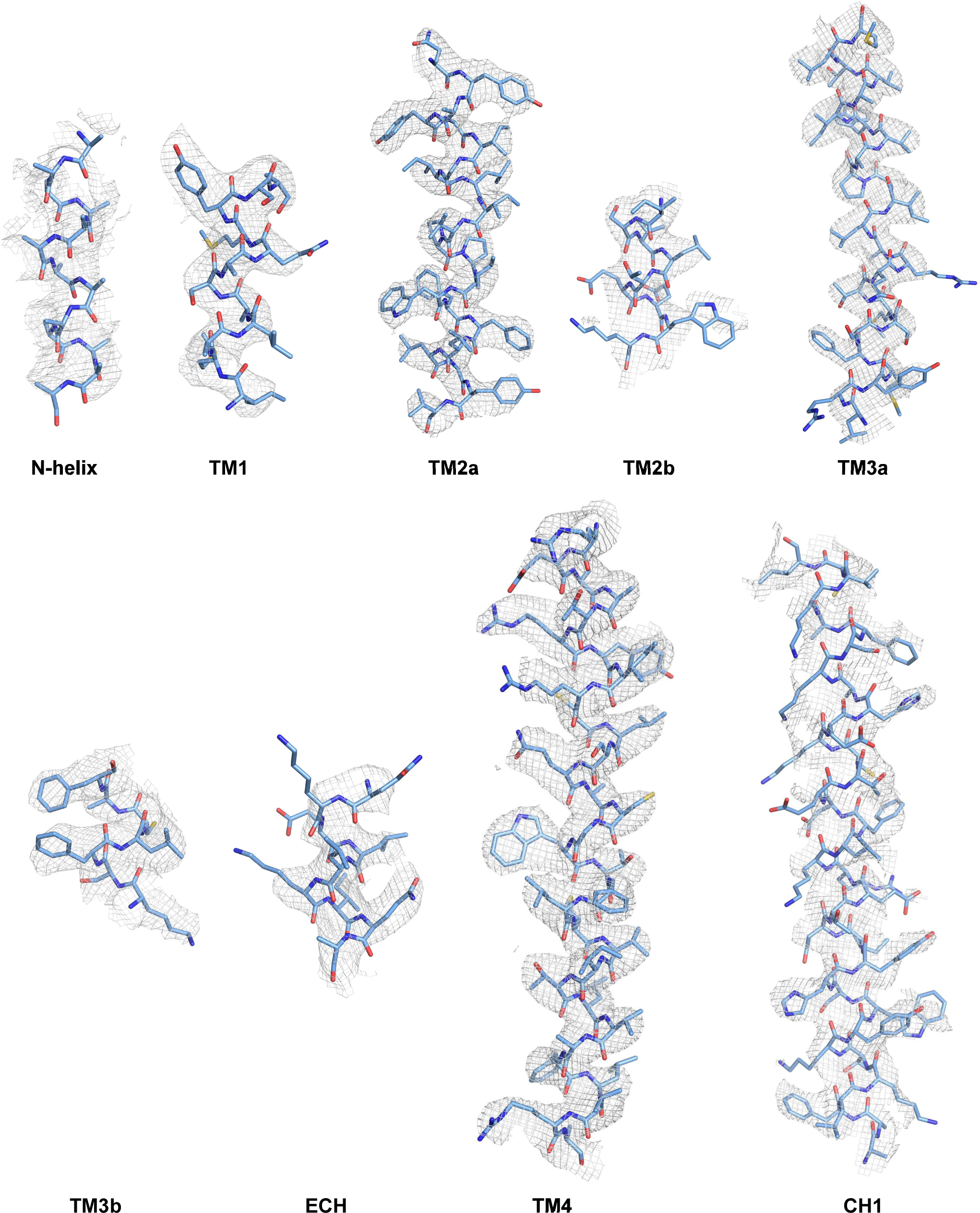
Representative cryoEM density maps (grey mesh) superposed with the atomic model of the drCALHM1-hepta-LCH channel for the helices.

**Supplemental Figure 4.**
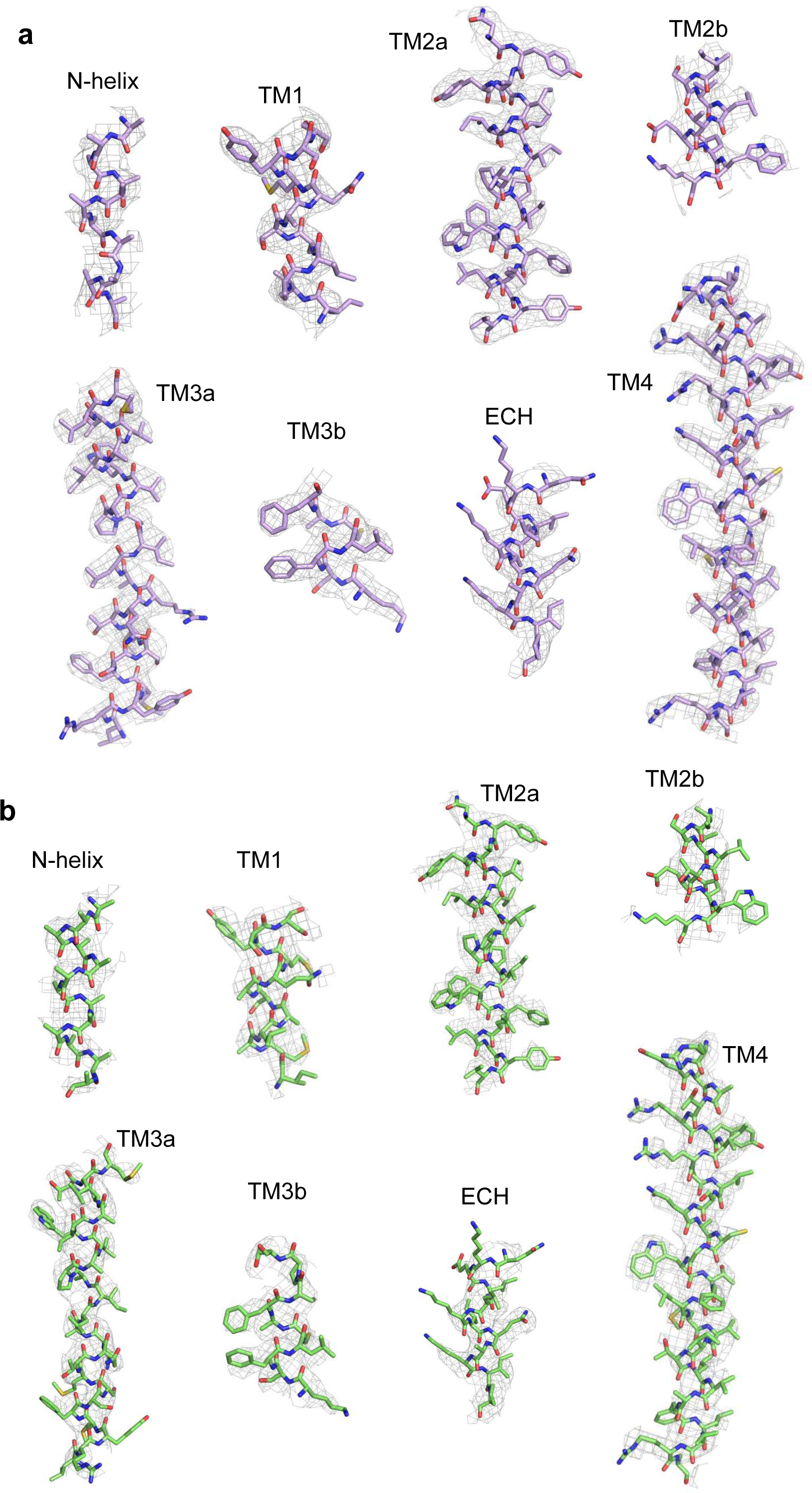
Representative cryo-EM density maps (gray mesh) superposed with the atomic model of the drCALHM1-hepta-noLCH (a) and drCALHM1-octa-noLCH (b) for the helices.

**Supplemental Figure 5.**
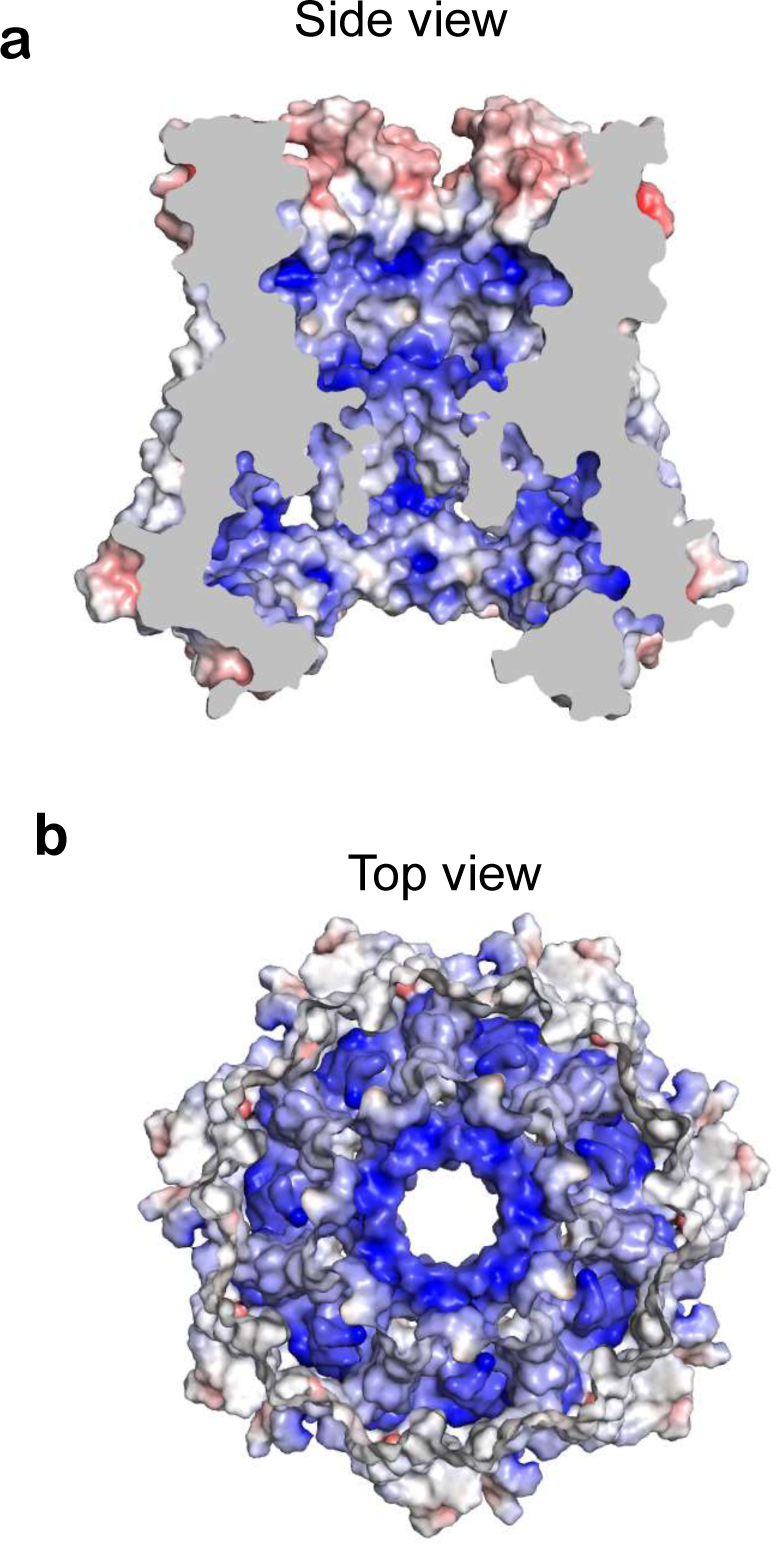
The electrostatic surface potential distribution of the heptameric drCALHM1 channel. (a) Side cut-away view and (b) top view of the electrostatic surface potential distribution of the heptameric drCALHM1 channel. The range of the electrostatic surface potential is shown from -10 kT/e (red) to +10 kT/e (blue).

**Supplemental Figure 6.**
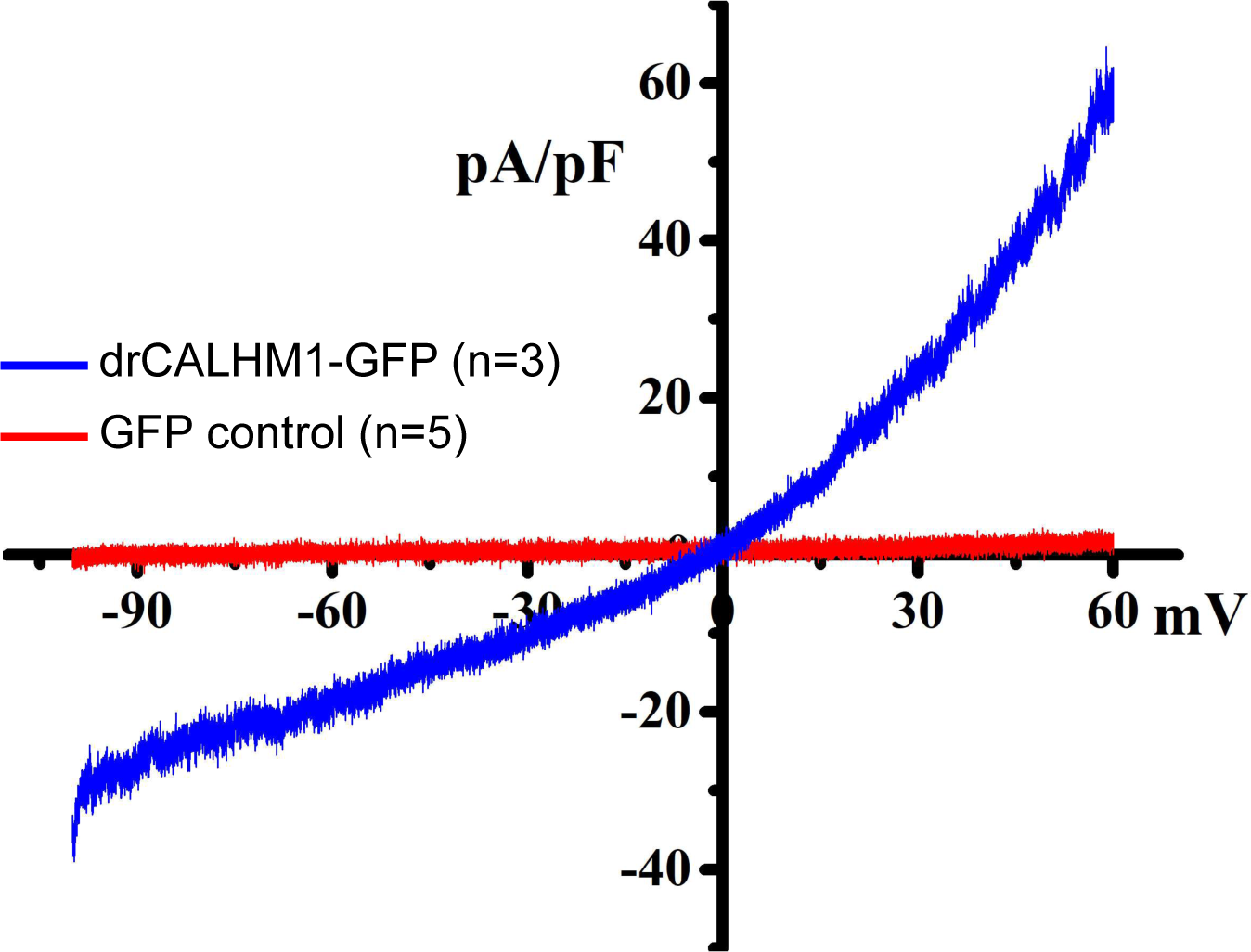
Whole cell patch clamp recordings of drCALHM1. Whole-cell currents measured in control GFP- and drCALHM1-GFP expressing HEK293T cells by a voltage-ramp protocol from -100 to +60 mV over 5 s (-60 mV holding potential) in bath solution containing 1.5 mM Ca^2+^ and 1 mM Mg^2+^.

**Supplemental Figure 7.**
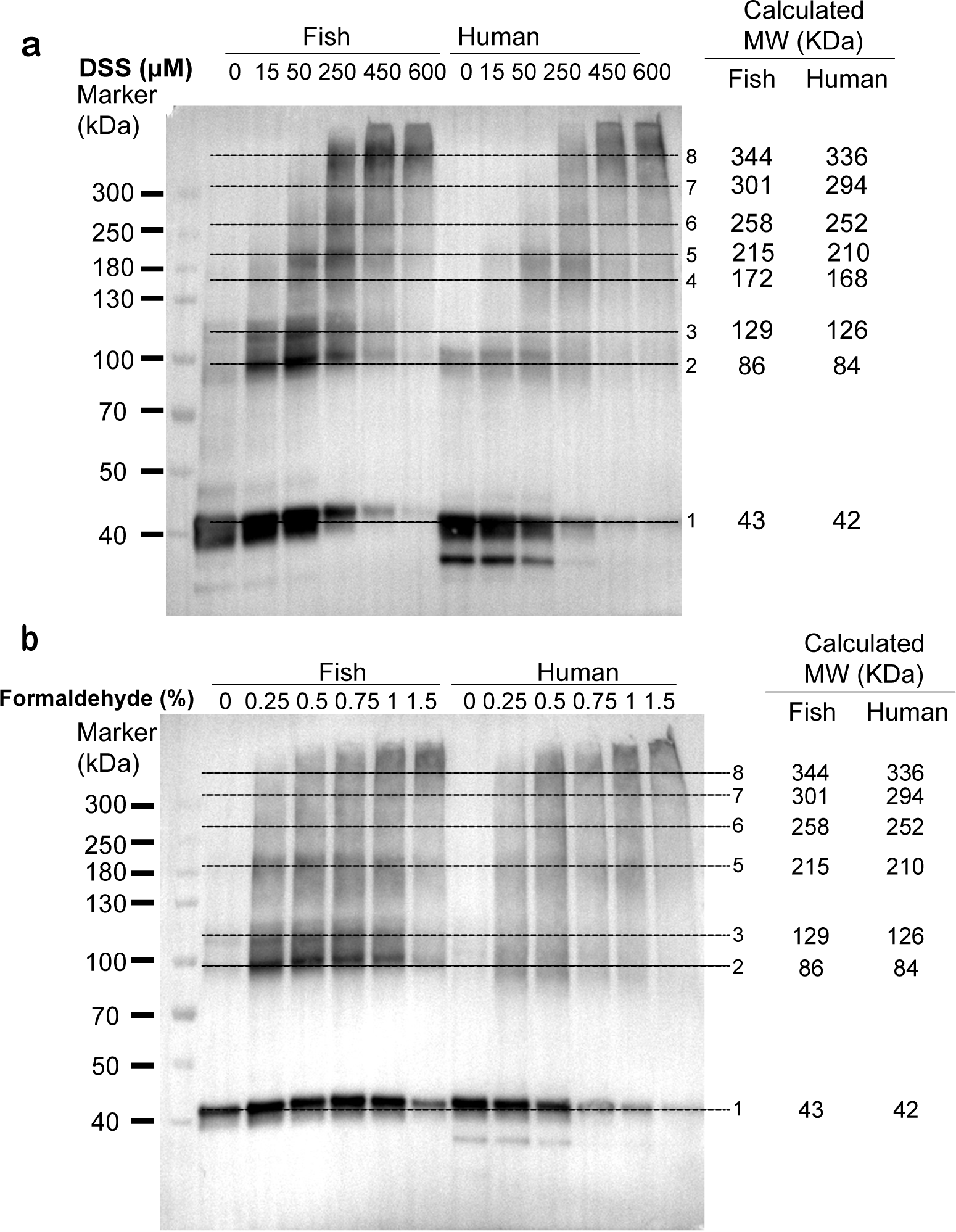
In cell crosslinking of the drCALHM1 channel. Western blot shows the results of cross-linking drCALHM1 in mammalian cells (HEK293T) using different concentrations of (a) DSS and (b) formaldehyde. As the concentration of the cross-linking agent increases, all oligomers on the cross-linking gradually appear until the octamer is formed. The black dashed line marks the position of the bands in different oligomerization states, and the theoretically calculated molecular weight is marked on the right.

**Supplemental Figure 8.**
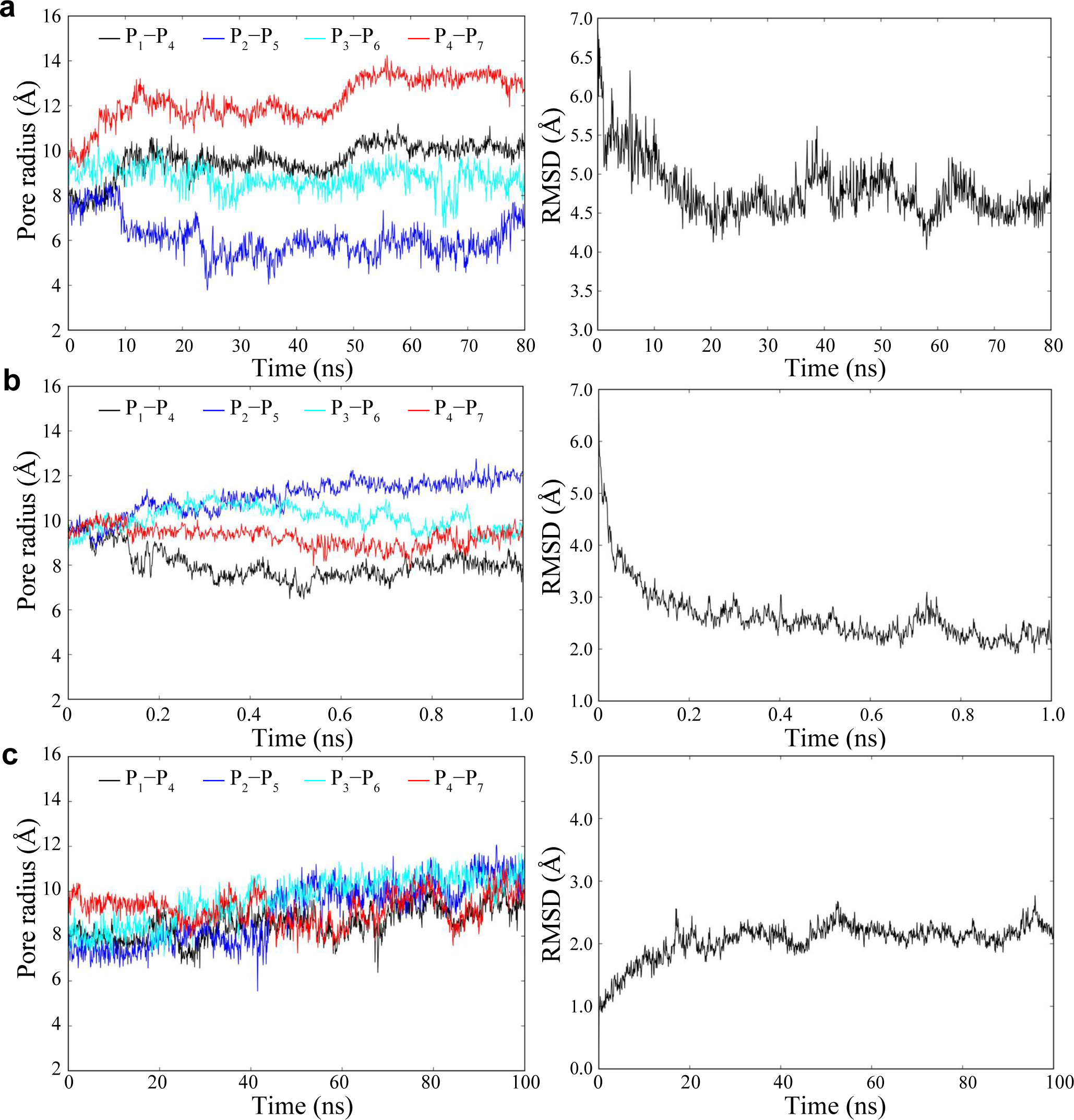
Time dependences of pore dynamics using different MD strategies. Time dependences of RMSD of Cα atoms in N-helix of P compared to that of P in octamer, and the minimum distance between the Cα atoms in N-helix of P_1_ and P_4_ (P_1_ –P_4_ distance), P_2_ −P_3_, P_3_ −P_6_ , and P_4_ –P_7_ distances calculated from (a) the supervised MD, (b) the targeted MD, and (c) the conventional MD trajectories.

**Supplemental Figure 9.**
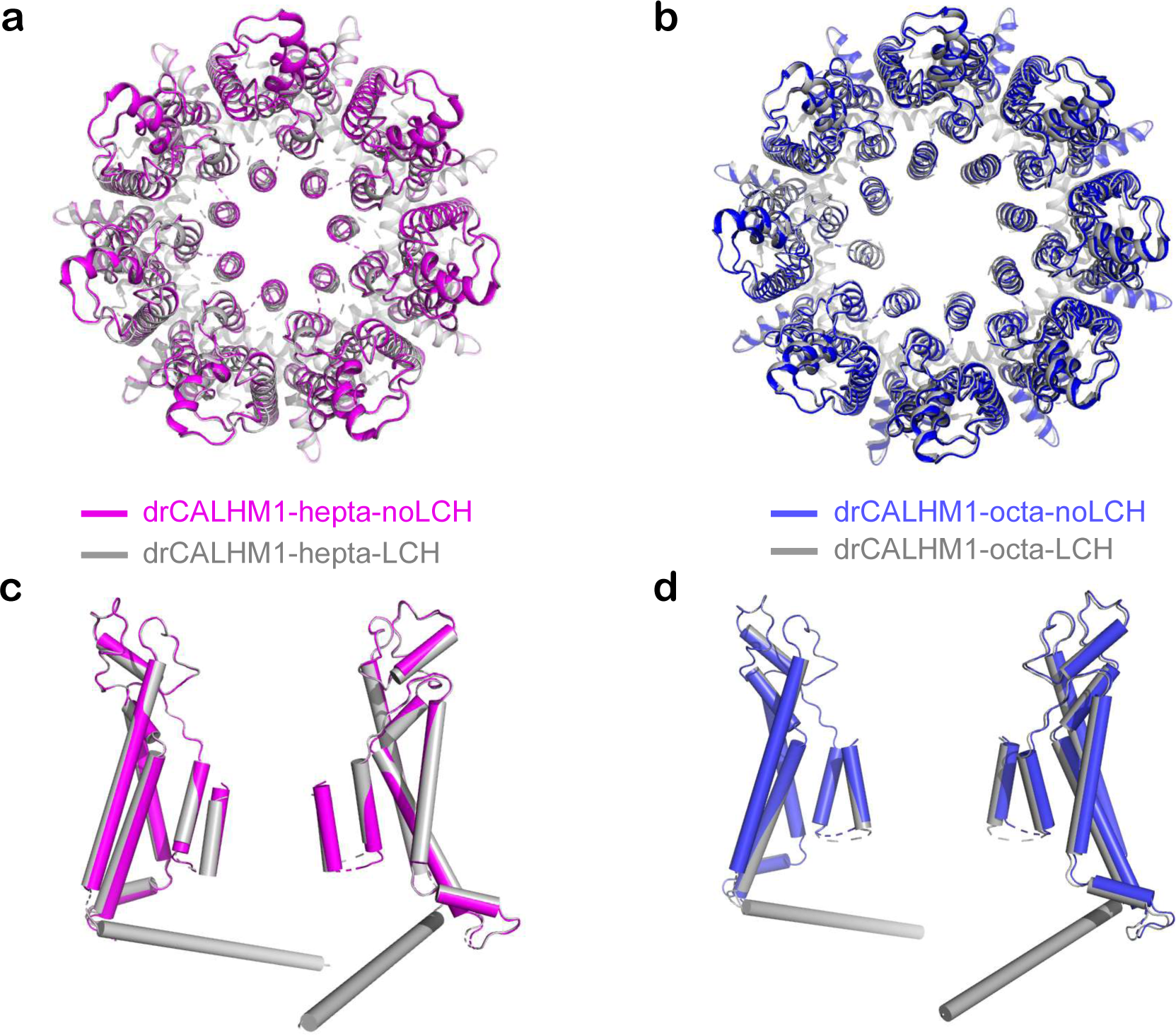
Structural comparison of the two oligomeric states with or without LCH. The two-state structures of the heptamer (a) and octamer (b) are superimposed and viewed from the top of the membrane. The comparison of the two monomer structures on both sides of the pore after the heptamer (c) and the octamer (d) overlap. The octamer and heptamer with LCH in gray. drCALHM1-hepta-noLCH and drCALHM1-octa-noLCH are shown in magenta and blue, respectively.

**Supplemental Figure 10.**
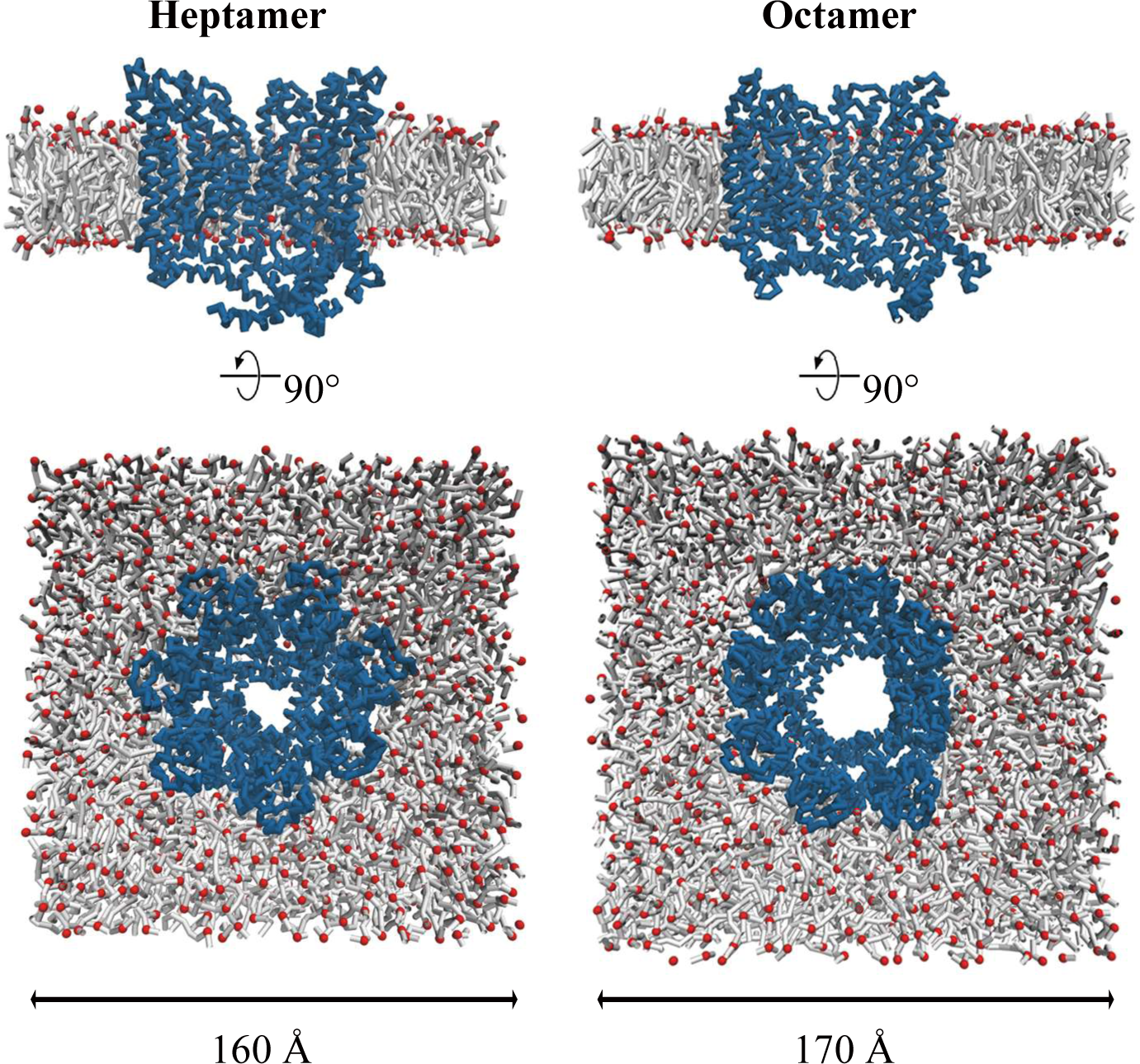
Coarse-grained MD simulations of heptamers (left) and octamers (right) embedded in POPC bilayer membranes. Side (cutaway) and top views of the final frame of one 10-μs replicate are shown in each case. The protein backbone particles are shown as blue ribbons, and phospholipid headgroups and acyl tails are shown as red balls and white sticks, respectively. Water and ions present in the simulation systems are omitted for clarity.

**Table S1.** CryoEM data collection and refinement statistics.

|  | **drCALHM1-hepta-**<br>**LCH** | **drCALHM1-hepta-**<br>**noLCH** | **drCALHM1-octa-**<br>**noLCH** |
| --- | --- | --- | --- |
| **Data Collection** |  |  |  |
| EM equipment | FEI Titan Krios |  |  |
| Voltage (kV) | 300 |  |  |
| Detector | Gatan K2 Summit |  |  |
| Pixel size (Å) | 1.014 |  |  |
| Electron Dose (e<sup>-</sup>/Å<sup>2</sup>) | 51.2 | 51.2 | 51.2 |
| Defocus range (μm) | -2.5 ~ -1.5 |  |  |
| **Reconstruction** |  |  |  |
| Software | Relion 3.0/ CryoSPARC-2 |  | Relion 3.0 |
| Number of used particles | 12,671 | 56,801 | 44,894 |
| Map sharpening B-factors (Å<sup>2</sup>) | -102 | -105 | -75 |
| Final Resolution (Å) | 3.2 | 2.9 | 3.5 |
| **Model Building** |  |  |  |
| Software | COOT |  |  |
| **Refinement** |  |  |  |
| Software | PHENIX real_space_refinement |  |  |
| Resolution (Å) | 3.5 | 3.6 | 3.7 |
| **Model composition** |  |  |  |
| Protein Residues | 1645 | 1351 | 1528 |
| Ligands | 7 | 7 | 8 |
| **Validation** |  |  |  |
| RMS deviation |  |  |  |
| Bonds Length (Å) | 0.006 | 0.004 | 0.004 |
| Bonds angle (°) | 0.951 | 0.613 | 0.537 |
| Ramachandran plot statistics (%) |  |  |  |
| Favorite | 97.33 | 97.33 | 98.38 |
| Allowed | 2.67 | 2.67 | 1.68 |
| Outlier | 0.00 | 0.00 | 0.00 |
| Rotamer outliers (%) | 0.00 | 0.00 | 0.00 |
| C-beta outliers (%) | 0.00 | 0.00 | 0.00 |
| Mean isotropic B |  |  |  |
| Protein/Ligands | 160.97/187.33 | 134.74/172.07 | 106.96/147.68 |

## Reference

Abbracchio MP, Burnstock G, Verkhratsky A, Zimmermann H (2009) Purinergic signalling in the nervous system: an overview. Trends Neurosci 32: 19–29

Adams PD, Afonine PV, Bunkoczi G, Chen VB, Davis IW, Echols N, Headd JJ, Hung LW, Kapral GJ, Grosse-Kunstleve RW, McCoy AJ, Moriarty NW, Oeffner R, Read RJ, Richardson DC, Richardson JS, Terwilliger TC, Zwart PH (2010) PHENIX: a comprehensive Python-based system for macromolecular structure solution. *Acta crystallographica Section D*, Biological crystallography 66: 213–221

Bussi G, Donadio D, Parrinello M (2007) Canonical Sampling Through Velocity Rescaling. The Journal of chemical physics 126: 014101

Choi W, Clemente N, Sun W, Du J, Lu W (2019) The structures and gating mechanism of human calcium homeostasis modulator 2. Nature 576: 163–167

D.A. Case, I.Y. Ben-Shalom, S.R. Brozell, D.S. Cerutti, T.E. Cheatham, III, V.W.D. Cruzeiro, T.A. Darden, R.E. Duke, D. Ghoreishi, M.K. Gilson, H. Gohlke, A.W. Goetz, D. Greene, R Harris, N. Homeyer, S. Izadi, A. Kovalenko, T. Kurtzman, T.S. Lee, S. LeGrand, P. Li, C. Lin, J. Liu, T. Luchko, R. Luo, D.J. Mermelstein, K.M. Merz, Y. Miao, G. Monard, C. Nguyen, H. Nguyen, I. Omelyan, A. Onufriev, F. Pan, R. Qi, D.R. Roe, A. Roitberg, C. Sagui, S. Schott-Verdugo, J. Shen, C.L. Simmerling, J. Smith, R. Salomon-Ferrer, J. Swails, R.C. Walker, J. Wang, H. Wei, R.M. Wolf, X. Wu, L. Xiao, York DM, Kollman PA. (2018) AMBER 2018. University of California, San Francisco.

Darden T, York D, Pedersen L (1993) Particle mesh Ewald: An Nlog (N) method for Ewald sums in large systems. The Journal of Chemical Physics 98: 10089–10092

Deganutti G, Moro S, Reynolds C (2020) A Supervised Molecular Dynamics Approach to Unbiased Ligand-Protein Unbinding. Journal of Chemical Information and Modeling XXXX

Demura K, Kusakizako T, Shihoya W, Hiraizumi M, Nomura K, Shimada H, Yamashita K, Nishizawa T, Taruno A, Nureki O (2020) Cryo-EM structures of calcium homeostasis modulator channels in diverse oligomeric assemblies. Science advances 6: eaba8105

Dickson CJ, Madej BD, Skjevik AA, Betz RM, Teigen K, Gould IR, Walker RC (2014) Lipid14: The Amber Lipid Force Field. Journal of Chemical Theory and Computation 10: 865–879

Drozdzyk K, Sawicka M, Bahamonde-Santos MI, Jonas Z, Deneka D, Albrecht C, Dutzler R (2020) Cryo-EM structures and functional properties of CALHM channels of the human placenta. eLife 9

Fernandez-Leiro R, Scheres SHW (2017) A pipeline approach to single-particle processing in RELION. Acta Crystallogr D Struct Biol 73: 496–502

Foskett JK (2020) Structures of CALHM channels revealed. Nat Struct Mol Biol 27:227–228

Hess B, Bekker H, Berendsen H, Fraaije J (1998) LINCS: A Linear Constraint Solver for molecular simulations. Journal of Computational Chemistry 18

Humphrey W, Dalke A, Schulten K (1996) VMD: Visual molecular dynamics. journal of molecular graphics and modelling 14: 33–38

Jorgensen WL, Chandrasekhar J, Madura JD, Impey RW, Klein ML (1983) Comparison of simple potential functions for simulating liquid water. The Journal of Chemical Physics 79: 926–935

Ma Z, Siebert AP, Cheung KH, Lee RJ, Johnson B, Cohen AS, Vingtdeux V, Marambaud P, Foskett JK (2012) Calcium homeostasis modulator 1 (CALHM1) is the pore-forming subunit of an ion channel that mediates extracellular Ca2+ regulation of neuronal excitability. Proceedings of the National Academy of Sciences of the United States of America 109: E1963–1971

Ma Z, Tanis JE, Taruno A, Foskett JK (2016) Calcium homeostasis modulator (CALHM) ion channels. Pflugers Arch 468: 395–403

Ma Z, Taruno A, Ohmoto M, Jyotaki M, Lim JC, Miyazaki H, Niisato N, Marunaka Y, Lee RJ, Hoff H, Payne R, Demuro A, Parker I, Mitchell CH, Henao-Mejia J, Tanis JE, Matsumoto I, Tordoff MG, Foskett JK (2018) CALHM3 Is Essential for Rapid Ion Channel-Mediated Purinergic Neurotransmission of GPCR-Mediated Tastes. Neuron 98: 547–561 e510

Maier JA, Martinez C, Kasavajhala K, Wickstrom L, Hauser KE, Simmerling C (2015) ff14SB: Improving the Accuracy of Protein Side Chain and Backbone Parameters from ff99SB. Journal of Chemical Theory and Computation 11: 3696–3713

Marrink SJ, de Vries AH, Mark AE (2004) Coarse Grained Model for Semiquantitative Lipid Simulations. The Journal of Physical Chemistry B 108: 750–760

Marrink SJ, Risselada HJ, Yefimov S, Tieleman DP, de Vries AH (2007) The MARTINI force field: Coarse grained model for biomolecular simulations. Journal of Physical Chemistry B 111: 7812–7824

Monticelli L, Kandasamy SK, Periole X, Larson RG, Tieleman DP, Marrink S-J (2008) The MARTINI Coarse-Grained Force Field: Extension to Proteins. Journal of Chemical Theory and Computation 4: 819–834

Parrinello MRA, Rahman AJ (1982) Polymorphic Transitions in Single Crystals: A New Molecular Dynamics Method. Journal of Applied Physics 52: 7182–7190

Pastor RW, Brooks BR, Szabo A (1988) An analysis of the accuracy of Langevin and molecular dynamics algorithms. Molecular Physics 65: 1409–1419

Periole X, Cavalli M, Marrink SJ, Ceruso MA (2009) Combining an Elastic Network With a Coarse-Grained Molecular Force Field: Structure, Dynamics, and Intermolecular Recognition. Journal of Chemical Theory and Computation 5: 2531–2543

Ren Y, Wen T, Xi Z, Li S, Lu J, Zhang X, Yang X, Shen Y (2020) Cryo-EM structure of the calcium homeostasis modulator 1 channel. Science advances 6: eaba8161

Rohou A, Grigorieff N (2015) CTFFIND4: Fast and accurate defocus estimation from electron micrographs. J Struct Biol 192: 216–221

Romanov RA, Lasher RS, High B, Savidge LE, Lawson A, Rogachevskaja OA, Zhao H, Rogachevsky VV, Bystrova MF, Churbanov GD, Adameyko I, Harkany T, Yang R, Kidd GJ, Marambaud P, Kinnamon JC, Kolesnikov SS, Finger TE (2018) Chemical synapses without synaptic vesicles: Purinergic neurotransmission through a CALHM1 channel-mitochondrial signaling complex. Sci Signal 11

Ruan Z, Orozco IJ, Du J, Lu W (2020) Structures of human pannexin 1 reveal ion pathways and mechanism of gating. Nature 584: 646–651

Ryckaert J-P, Ciccotti G, Berendsen HJC (1977) Numerical Integration of the Cartesian Equations of Motion of a System with Constraints: Molecular Dynamics of N-alkanes. Journal of Computational Physics 23: 327–341

Salmaso V, Sturlese M, Cuzzolin A, Moro S (2017) Exploring Protein-Peptide Recognition Pathways Using a Supervised Molecular Dynamics Approach. Structure 25

Schlitter J, Engels M, Krüger P (1994) Targeted Molecular-Dynamics – a New Approach for Searching Pathways of Conformational Transitions. Journal of Molecular Graphics 12: 84–89

Siebert AP, Ma Z, Grevet JD, Demuro A, Parker I, Foskett JK (2013) Structural and functional similarities of calcium homeostasis modulator 1 (CALHM1) ion channel with connexins, pannexins, and innexins. J Biol Chem 288: 6140–6153

Swint-Kruse L, Brown CS (2005) Resmap: automated representation of macromolecular interfaces as two-dimensional networks. Bioinformatics 21: 3327–3328

Syrjanen JL, Michalski K, Chou TH, Grant T, Rao S, Simorowski N, Tucker SJ, Grigorieff N, Furukawa H (2020) Structure and assembly of calcium homeostasis modulator proteins. Nat Struct Mol Biol 27: 150–159

Tanis JE, Ma Z, Foskett JK (2017) The NH2 terminus regulates voltage-dependent gating of CALHM ion channels. American journal of physiology Cell physiology 313: C173–C186

Taruno A (2018) ATP Release Channels. Int J Mol Sci 19

Taruno A, Vingtdeux V, Ohmoto M, Ma Z, Dvoryanchikov G, Li A, Adrien L, Zhao H, Leung S, Abernethy M, Koppel J, Davies P, Civan MM, Chaudhari N, Matsumoto I, Hellekant G, Tordoff MG, Marambaud P, Foskett JK (2013) CALHM1 ion channel mediates purinergic neurotransmission of sweet, bitter and umami tastes. Nature 495: 223–226

Wassenaar TA, Ingolfsson HI, Boeckmann RA, Tieleman DP, Marrink SJ (2015) Computational Lipidomics with insane: A Versatile Tool for Generating Custom Membranes for Molecular Simulations. Journal of Chemical Theory and Computation 11: 2144–2155

Webb B, Sali A (2016) Comparative Protein Structure Modeling Using MODELLER. Current protocols in protein science 86: 2.9.1-2.9.37

Xiao X, Zeng X, Yuan Y, Gao N, Guo Y, Pu X, Li M (2015) Understanding the conformation transition in the activation pathway of β2 adrenergic receptor via a targeted molecular dynamics simulation. Physical Chemistry Chemical Physics Pccp 17

Zheng SQ, Palovcak E, Armache JP, Verba KA, Cheng Y, Agard DA (2017) MotionCor2: anisotropic correction of beam-induced motion for improved cryo- electron microscopy. Nat Methods 14: 331–332

